# An adaptive compromise - Conflicting evolutionary pressures on arthropod-borne Zika virus dinucleotide composition in mammalian hosts and mosquito vectors

**DOI:** 10.1101/2021.02.09.430415

**Authors:** Jelke J. Fros, Imke Visser, Bing Tang, Kexin Yan, Eri Nakayama, Tessa M. Visser, Constantianus J.M. Koenraadt, Monique M. van Oers, Gorben P. Pijlman, Andreas Suhrbier, Peter Simmonds

## Abstract

Most vertebrate RNA viruses show pervasive suppression of CpG and UpA dinucleotides, closely resembling the dinucleotide composition of host cell transcriptomes. In contrast, CpG suppression is absent in both invertebrate mRNA and RNA viruses that exclusively infect arthropods. Arthropod-borne (arbo) viruses are transmitted between vertebrate hosts by invertebrate vectors and thus encounter potentially conflicting evolutionary pressures in the different cytoplasmic environments. Using a newly developed Zika virus (ZIKV) model, we have investigated how demands for CpG suppression in vertebrate cells can be reconciled with potentially quite different compositional requirements in invertebrates, and how this affects ZIKV replication and transmission.

Mutant viruses with synonymously elevated CpG or UpA dinucleotide frequencies showed attenuated replication in vertebrate cell lines, which was rescued by knockout of the zinc-finger antiviral protein (ZAP). Conversely, in mosquito cells, ZIKV mutants with elevated CpG dinucleotide frequencies showed substantially enhanced replication compared to wildtype. Host-driven effects on virus replication attenuation and enhancement were even more apparent in mouse and mosquito models. Infections with CpG-or UpA-high ZIKV mutants in mice did not cause typical ZIKV-induced tissue damage and completely protected mice during subsequent challenge with wildtype virus, which demonstrates their potential as live-attenuated vaccines. In contrast, the CpG-high mutants displayed enhanced replication in *Aedes aegypti* mosquitoes and a larger proportion of mosquitoes carried infectious virus in their saliva.

These findings show that mosquito cells are also capable of discriminating RNA based on dinucleotide composition. However, the evolutionary pressure on the CpG dinucleotides of viral genomes in arthropod vectors directly opposes the pressure present in vertebrate host cells, which provides evidence that an adaptive compromise is required for arbovirus transmission. This suggests that the genome composition of arthropod-borne flaviviruses is crucial to maintain the balance between high-level replication in the vertebrate host and persistent replication in the mosquito vector.

## Introduction

Arthropod-borne viruses (arboviruses) are transmitted between vertebrate hosts by invertebrate arthropod vectors such as mosquitoes, ticks, biting midges and sand-flies. When hematophagous vectors feed on infected vertebrate hosts, the virus first needs to enter and pass through the vector’s midgut endothelial cells and persistently infect other tissues to disseminate into the vector’s salivary glands before the arbovirus is transmitted to the next vertebrate host during subsequent feeding events. Transmission of arboviruses thus requires active virus replication in two evolutionary distant hosts that are separated by more than 500 million years of evolution [1]. Hence, arboviruses encounter divergent cellular and molecular environments, exemplified by the different antiviral responses that vertebrates hosts and invertebrate vectors rely on to cope with viral infections. Vertebrate animals possess a plethora of innate and adaptive antiviral immune responses [2], while invertebrates largely rely on RNA interference (RNAi) to control viral infections [3, 4]. As intracellular pathogens, all viruses rely on cellular machinery for their replication, which exposes viruses to the intracellular milieu of the host cell. The transmission cycle of arboviruses places them in two distinct intracellular environments with dissimilar adaptive pressures for replication and evasion of host antiviral responses (reviewed in [5]). How arboviruses are able to thrive while continuously alternating between these distinct cellular environments remains an intriguing research topic [6].

One of the most striking molecular differences between vertebrate and invertebrate organisms is the dinucleotide usage in their genomic DNA and coding mRNA sequences. In the DNA of vertebrates the dinucleotide combination of a cytosine followed by a guanine (CpG) occurs less frequently than predicted, based on frequencies of C and G mononucleotides. The cytosine in CpG is prone to methylation, which can cause the cytosine to deaminate and mutate into a thymine [7, 8]. In contrast, the genome of most invertebrates contains largely unbiased frequencies of CpG dinucleotides, reflecting greatly reduced DNA methylation activity [9]. Both vertebrates and invertebrates have additionally evolved to reduce UpA dinucleotide frequencies in their mRNA through a range of mechanisms largely related to mRNA stability [10, 11]. Most viruses with RNA genomes have evolved to mirror the dinucleotide usage of their host’s mRNA, with invertebrate viruses solely suppressing UpA dinucleotides and vertebrate viruses underrepresenting both UpA and CpG dinucleotides, even though CpGs in viral RNA are not subjected to the same methylation and mutation pressure that acts on vertebrate host DNA [12–14].

Artificially increasing CpG or UpA dinucleotide frequencies through hundreds of synonymous mutations effectively attenuates the replication of multiple RNA viruses in vertebrate cell cultures [15–19]. Vertebrate cells identify unnaturally elevated levels of CpG dinucleotides as non-self, mediated by the zinc-finger antiviral protein (ZAP) [16, 20–22]. We previously showed that ZAP is also required for the attenuation of UpA-high mutants of echovirus 7 (E7) (genus *Enterovirus B*, family *Picornaviridae*) [21]. ZAP is interconnected with the interferon response and can modulate translation initiation through the sequestration of eukaryotic translation initiation factor 4A (eIF4A) [23–25]. ZAP is additionally associated with cellular RNA degradation, with links to stress granule and processing body components as well as to the endonuclease RNase L activated by 2’-5’-oligoadenylate synthetase 3 (OAS3), the 3’-5’ exosomal and the 5’-3’ XRN1-mediated RNA degradation pathways [20, 26–28]. Together, this indicates that vertebrate ZAP is involved in the detection of both UpA and CpG dinucleotides and recruits diverse host cell proteins to inhibit virus replication [21, 29, 30].

Whether invertebrate cells possess similar non-self RNA detection mechanisms and whether alternating between vertebrate hosts and arthropod vectors puts additional selective pressure on arboviruses is currently unknown. The lack of CpG suppression in invertebrates and their viruses suggests the absence of selection pressure against CpGs in arthropods, while UpA dinucleotides are clearly under-represented in both arthropods and their viruses [13, 14]. We therefore hypothesized that the selection pressures on UpA dinucleotides would be similar in vertebrate hosts and arthropod vectors, while the selection pressure on CpG dinucleotides may be absent in these vectors. To investigate the constraints that an arbovirus encounters during its transmission cycle we turned to the genus *Flavivirus* (Family *Flaviviridae*). This genus contains many clinically relevant arboviruses, which include mosquito-borne flaviviruses (MBFV), *e.g*. Zika virus (ZIKV), dengue virus, West Nile virus, Japanese encephalitis virus and yellow fever virus, as well as a number of tick-borne flaviviruses (TBFV), *e.g*. tick-borne encephalitis virus. In addition to arboviruses, the genus *Flavivirus* also contains members with no known vector (NKVF) and insect-specific flaviviruses (ISF). The latter are unable to infect vertebrates and exclusively replicate in insects and cultured insect cells [31]. To investigate whether arbovirus genome composition is subject to distinct evolutionary pressures in vertebrate host and arthropod vectors, we constructed ZIKV mutants with synonymously increased CpG or UpA dinucleotide frequencies and tested their replication kinetics in vertebrate and invertebrate cells, mice and mosquitoes.

## Results

### Flavivirus genome composition and mutant Zika virus design

To compare the CpG and UpA dinucleotide usage between different flaviviruses, the ratio of CpG and UpA dinucleotide frequencies from full length flavivirus genome sequences were plotted as the observed over the expected (O/E) frequencies (Fig 1A, large data points). Viral sequence data was superimposed onto large randomly selected sets of vertebrate (*H. sapiens*) and invertebrate mosquito (*Ae. aegypti*) mRNAs (Fig 1A, small data points). Both human host and mosquito vector mRNA suppress UpA dinucleotides (y-axis), while only human mRNA under-represents CpG dinucleotides (x-axis). All flaviviruses that infect vertebrates (*i.e*. MBFV, TBFV and NKVF) have clearly adapted to mimic the genome composition of the vertebrate host, with marked under-representation of both CpG and UpA dinucleotides. In contrast, the ISFs and particularly the phylogenetically most distant lineage 1 ISFs contain a much higher ratio of CpG dinucleotides frequencies, more closely resembling the unbiased CpG frequency found in their arthropod hosts. Lineage II ISFs display an insect-specific phenotype and contain more CpG dinucleotides than VIFs, while phylogenetically they cluster together with the VIFs (Fig 1A) [14, 31–34].

**Fig 1.**
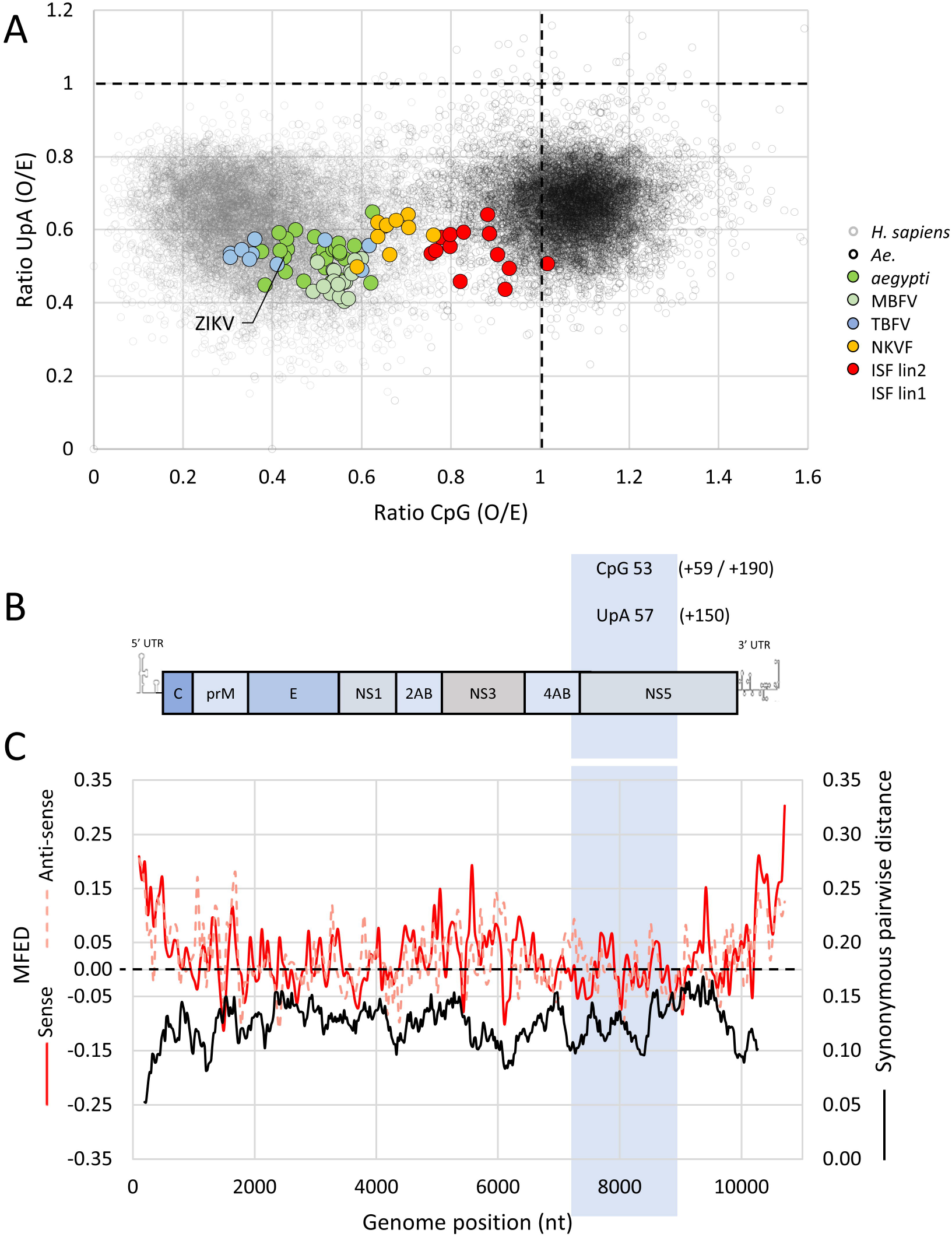
Flavivirus CpG and UpA dinucleotide usage, Zika virus genome organization and the position of the mutated region. **(A)** Data points represent the ratio of CpG (x-axis) and UpA (y-axis) dinucleotides in individual RNA sequences. Small data points denote human mRNAs (gray) and mosquito mRNAs (black). Large data points represent full length genome sequences of flavivirus species with mosquito-borne flaviviruses (MBFV) green, tick-borne flaviviruses (TBFV) light green, no known vector flaviviruses (NKVF) blue and the lineage 1 and 2 insect-specific flaviviruses (ISF) red and orange, respectively. The observed / expected (O/E) ratio of 1.0 (dotted lines) is the mathematically expected frequency of occurrence if the RNA’s mononucleotides were randomly distributed. **(B)** Schematic representation of the ZIKV genome, with the mutated region boxed and displaying the number of CpG and UpA dinucleotides in wild type ZIKV. Additional CpG and UpA dinucleotides of the CpG_1.0 (+59), CpG_max (+190) and UpA_max (+150) mutant viruses are listed in brackets. **(C)** Minimal free energy difference (MFED) and synonymous pairwise distance (black line) were calculated for 19 ZIKV isolates that were >1% dissimilar in nucleotide sequences using a fragment size of 200 with 48 nucleotide increments for MFED and sliding window of 21 codons for synonymous pairwise distance that were >1% dissimilar in nucleotide sequences using a fragment size of 200 with 48 nucleotide increments for MFED and sliding window of 21 codons for synonymous pairwise distance as used in previous analyses of echovirus 7 and murine norovirus [19, 59]. MFED values of both the sense (red line) and the anti-sense (dashed pink line) strand in the selected region are close to the null expectation (dotted black line). Values were calculated using SSE software v1.4.

To investigate the biological role of CpG and UpA suppression in a typical mosquito-borne flavivirus, ZIKV mutants were created with increased frequencies of either CpG or UpA dinucleotides by introducing many synonymous changes into a 1654 nucleotide long, coding region of the viral genome (Fig 1B). The complete genome of wildtype (WT) ZIKV has ratios of 0.42 and 0.52 for the O/E frequencies of CpG and UpA dinucleotides, respectively. The selected region contains 53 CpG and 57 UpA dinucleotides with comparable CpG and UpA suppression (detailed in S1 Table). Before introducing any mutations, we confirmed that the ZIKV open reading frame (ORF), except for the 5’–end which contains the cyclisation sequence [35], has a relatively uniform nucleotide variability at synonymous sites and low minimal free energy differences (MFED)(Fig 1C). This suggests that, apart from conserving amino acid sequence, there are no other conserved nucleotide sequences or critical RNA structures in the ZIKV ORF. To confirm the absence of critical RNA sequences experimentally, a scrambled control virus (SCR) was designed that contained extensive synonymous changes in the target region, but retained WT dinucleotide frequencies. Furthermore, we created mutants that maximally over-represented CpG or UpA dinucleotides, without altering the number of the other UpA or CpG dinucleotide or the encoded amino acid sequence. This resulted in the CpG_max ZIKV mutant with 190 additional CpG dinucleotides and an O/E ratio of 1.62 and the UpA_max ZIKV mutant (+ 150 UpA dinucleotides, O/E ratio 1.72). As the genome of *Aedes aegypti*, the main mosquito vector for ZIKV, shows no suppression of CpG dinucleotides (O/E ratio ~ 1.0) (Fig 1A), we additionally created a mutant virus with synonymously increased CpG dinucleotides to the unsuppressed (mosquito-like) ratio of 1.0 (termed CpG_1.0) by addition of 59 CpG dinucleotides. We made use of an infectious cDNA clone to generate virus stocks of these ZIKV mutants in Vero E6 cells. As the fitness between the ZIKV mutants was expected to vary substantially, we used equal numbers of viral RNA copies to infect cells.

### ZIKV with elevated CpG or UpA dinucleotide frequencies is attenuated in vertebrate cell cultures

Vertebrate cell lines were infected with the different mutant ZIKV stocks to investigate the replication kinetics of the mutant viruses. Samples were taken daily from cell culture supernatants and checked for the presence of infectious virus (Fig 2). In all cases the SCR control behaved very similarly to the WT, indicating that the introduced synonymous changes did not affect virus replication. In vertebrate Vero E6 cells and A549 cells, UpA_max ZIKV was clearly attenuated with ~10-fold lower titres of progeny virus at any time post infection. The CpG_max ZIKV mutant displayed 2-to 3-fold lower titres compared to WT, while the replication kinetics of the CpG_1.0 mutant virus were similar to that of the WT virus (Fig 2A, B). At two days post infection, CpG_max and UpA_max virus differed significantly from WT and the SCR control (Fig 2D) (two-way ANOVA, F (4, 30) = 6.315, p < 0.05). Knockout of ZAP rescued the attenuated phenotype of the CpG_max ZIKV mutant, compared to that in wildtype A549 cells (Fig 2C, D) (two-way ANOVA, F (1, 30) = 11.49, p < 0.05). Although replication of the UpA_max ZIKV mutant appeared similarly rescued by knockout of ZAP (Fig 2B-D), this was not significantly different from replication in wildtype A549 cells (p = 0.06). Similar to what was reported previously for CpG-high echovirus 7 mutants [21], knockout of RNaseL or OAS3 resulted in comparable replication rates between CpG_max and WT ZIKV to WT levels (Fig 2 and S1 Fig A-C).

**Fig 2.**
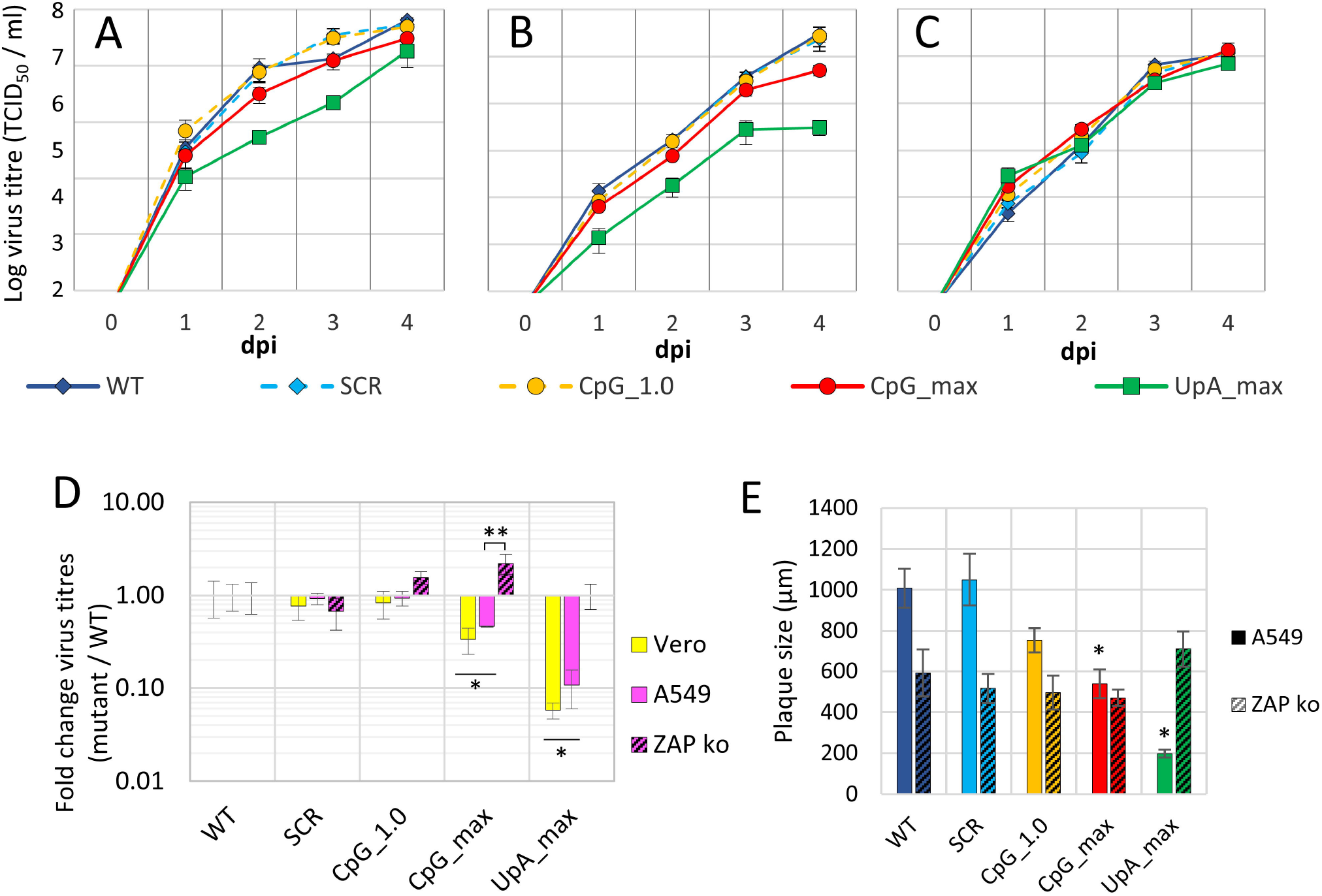
Replication of ZIKV with increased UpA and CpG dinucleotides is restricted in vertebrate cell culture. Vertebrate Vero E6 cells **(A)**, A549 cells **(B)** and A549 ZAP knockout cells **(C)** were infected with the mutant ZIKV viruses at 1 RNA / cell. At the indicated days post infection (dpi) the 50% tissue culture infectious dose was determined by end point dilution assays. **(D)** Relative replication of mutant viruses was calculated by dividing mutant over WT virus titres at 2dpi. Data points represent the average of a total of four independent experiments, compiled of two experiments performed for each of two independently rescued virus populations. The error bars indicate one standard error of the mean. Single asterisks highlight significant differences in wildtype vertebrate cells from WT ZIKV and two asterisks the difference between A549 cells and A549 ZAP knockout cells (two-way ANOVA, FDR adjusted P < 0.05). **(E)** Average plaque size measured in A549 and A549 ZAP knockout cells 3 dpi with ZIKV or indicated mutant viruses. After removal of the overlay, cells were fixed and stained for ZIKV E protein. Micrographs were taken from plaques from two independent experiments. The plaque diameter was measured using the Axiovision software. Data points represent the average plaque size, error bars indicate one standard error of the mean and asterisks highlight significant difference from WT (one-way ANOVA with Dunnett’s post Hoc test, P < 0.05).

The cytopathic effects (CPE) caused by ZIKV infections in A549 cells were mild; however, delays in CPE for the CpG_1.0, CpG_max and UpA_max viruses were observed. To investigate replication phenotypes further, a plaque assay was performed and plaque sizes were determined by immunostaining as a measure for the rate at which infectious virus spread through the culture. Three days post inoculation with WT ZIKV and SCR ZIKV, large plaques with an average diameter just over 1000 μm were observed in parental A549 cells (Fig 2E and S1 Fig D). Inoculation with CpG_1.0 ZIKV resulted in slightly smaller average plaque diameters (~750 μm), while the CpG_max and UpA_max ZIKV mutants displayed significantly smaller average plaque sizes (530 μm and 200 μm, respectively) (Fig 2E and S1 Fig D) (one-way ANOVA, F (4, 23) = 19.06, p <0.05). The same virus dilutions were used to infect A549 ZAP knockout cells in parallel. In these cells, plaques observed in response to WT ZIKV infection were smaller compared to WT infections in ZAP expressing A549 cells. However, no further reduction in plaque sizes was observed when cells were infected with CpG-or UpA-high mutant ZIKV (Fig 2E and g D) (one-way ANOVA, F (4, 19) = 1.020, p=0.42). Plaque phenotypes were confirmed in Vero E6 in which also the average plaque diameters of the CpG_1.0 mutant were significantly smaller than those of WT (S1 Fig E) (F (4, 25) = 65.29, p < 0.05). (S1 Fig E). The analysis of plaque diameters suggests that ZAP is responsible for the attenuation of both CpG_max and UpA_max mutant viruses. The results further suggest that the CpG_1.0 mutant is mildly attenuated in vertebrate cells that express ZAP, something that was not visible from the virus growth curves (Fig 2).

### Replication of ZIKV mutants is differentially affected by genomic CpG and UpA dinucleotide frequencies in mosquito cells

Cell lines originating from *Aedes spp*. mosquitoes were infected with WT ZIKV and the mutant viruses to investigate how changes in CpG and UpA dinucleotide frequencies affect ZIKV replication in invertebrate cells. The replication kinetics of WT ZIKV and the SCR control were highly similar. Interestingly, both mutants with increased CpG dinucleotide frequencies (Fig 3A, B and S2 Fig A-C, circles) displayed an increased fitness compared to WT in all *Aedes* mosquito cells tested, with a most consistent difference in the exponential growth phase at two days post infection. Two days post infection the mean virus titres of the CpG_1.0 and the CpG_max mutants across all five cell lines were 6- and 5-fold higher than WT, respectively (two-way ANOVA, F (4, 45) = 4.120, p = < 0.05). In contrast, virus titres of the UpA_max mutant were lower than WT in all cell lines tested, however this did not significantly differ from WT (Fig 3C).

**Fig 3.**
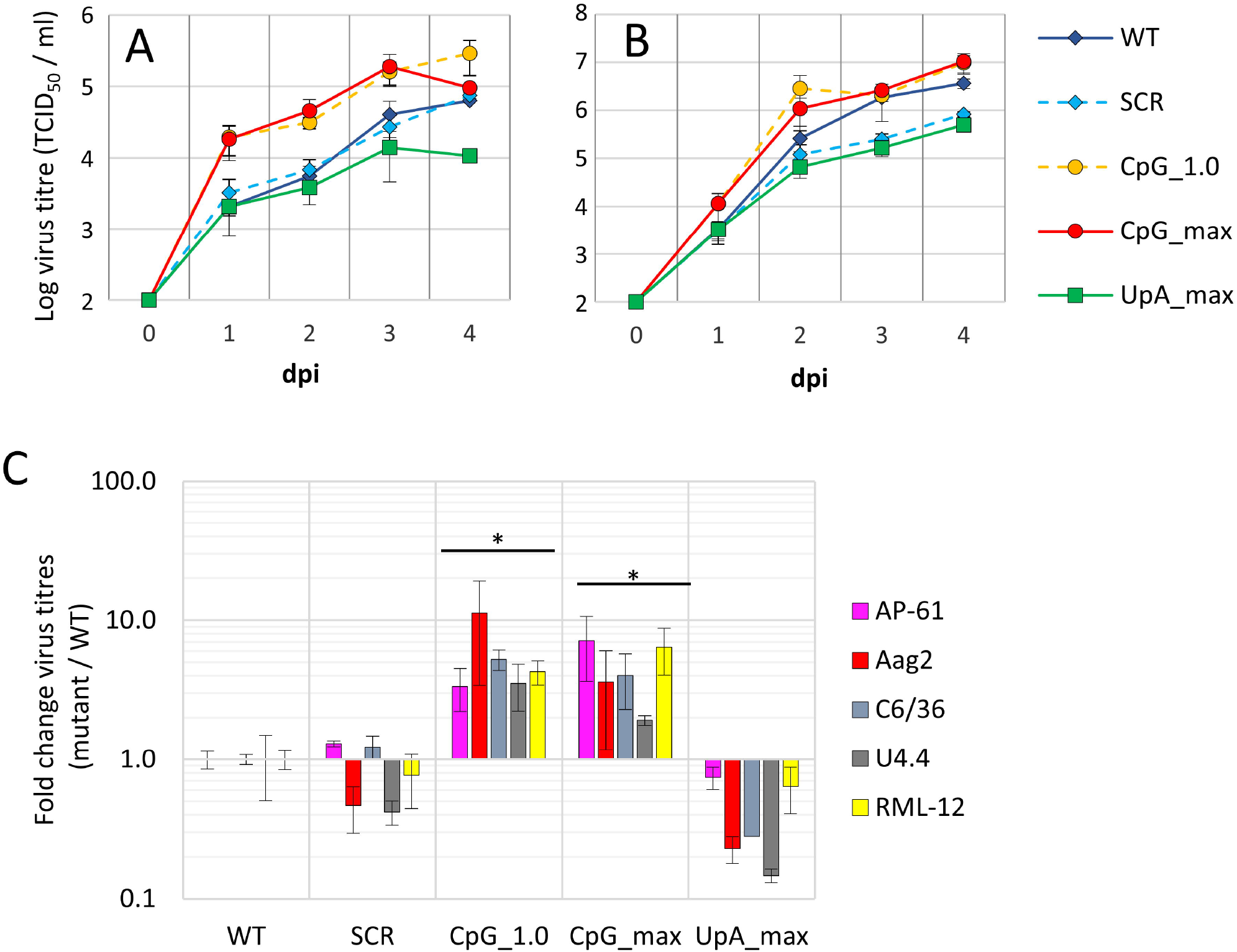
Replication of ZIKV with increased CpG dinucleotides has a replication advantage in mosquito cells. *Aedes* mosquito cells: *Ae*. *pseudoscutellaris* AP-61 **(A)**, *Ae. aegypti* Aag2 **(B)** were infected with the mutant ZIKV viruses at 10 RNA / cell. At the indicated days post infection (dpi) the 50% tissue culture infectious dose was determined by end point dilution assays. Data points represent the average of a total of four independent experiments, compiled of two experiments performed for each of two independently rescued virus populations. **(C)** Relative replication of mutant viruses in 5 distinct mosquito cell lines originating from *Ae*. *pseudoscutellaris* AP-61 **(A)**, *Ae. aegypti* Aag2 **(B)** and *Ae. albopictus* RML-12, C6/36 and U4.4 (growth curves in S2 Fig) were calculated by dividing mutant over WT virus titres at 2dpi. The error bars indicate one standard error of the mean. Asterisks indicate that over all five mosquito cell lines, mutant virus replication is significantly different from WT (two-way ANOVA, F (4, 45) = 4.120, FDR adjusted p < 0.05).

To confirm the opposing replicative fitness of the CpG-high mutant viruses in either vertebrate or mosquito cells in a more sensitive assay, a competitive fitness experiment was performed. Human A549 cells, A549 ZAP knockout cells and AP-61 mosquito cells were infected with two ZIKV variants. WT ZIKV was mixed with either the scrambled control virus (SCR), CpG-high viruses (CpG_1.0 and CpG_max) or UpA_max at equal RNA concentrations. Virus was passaged to new cells twice before total RNA was isolated and part of the mutated region within the ZIKV genome was reverse transcribed and PCR amplified. Amplicon DNA was digested to identify WT and mutant viruses. As expected, WT and SCR viruses were both detected in all three cell types (Fig 4A). In A549 cells, the CpG_1.0 mutant was almost eliminated, while the CpG_max mutant was completely outcompeted by WT (Fig 4B,C). In contrast, in AP-61 mosquito cells, more CpG_max ZIKV was detected compared to WT and the CpG_1.0 mutant had almost completely outcompeted WT (Fig 4B,C). The UpA_max mutant was outcompeted by WT in A549 cells, while in mosquito cells it displayed fitness equal to that of WT. Knockout of ZAP, strongly reduced any fitness advantages of WT over the mutants in A549 cells as all mutants were readily detected together with WT (Fig 4A-D). Together with the virus growth curves, this indicates that while vertebrate cells attenuate the replication of viruses with increased CpG dinucleotide frequencies, these viruses have a replication advantage in mosquito cells.

**Fig 4.**
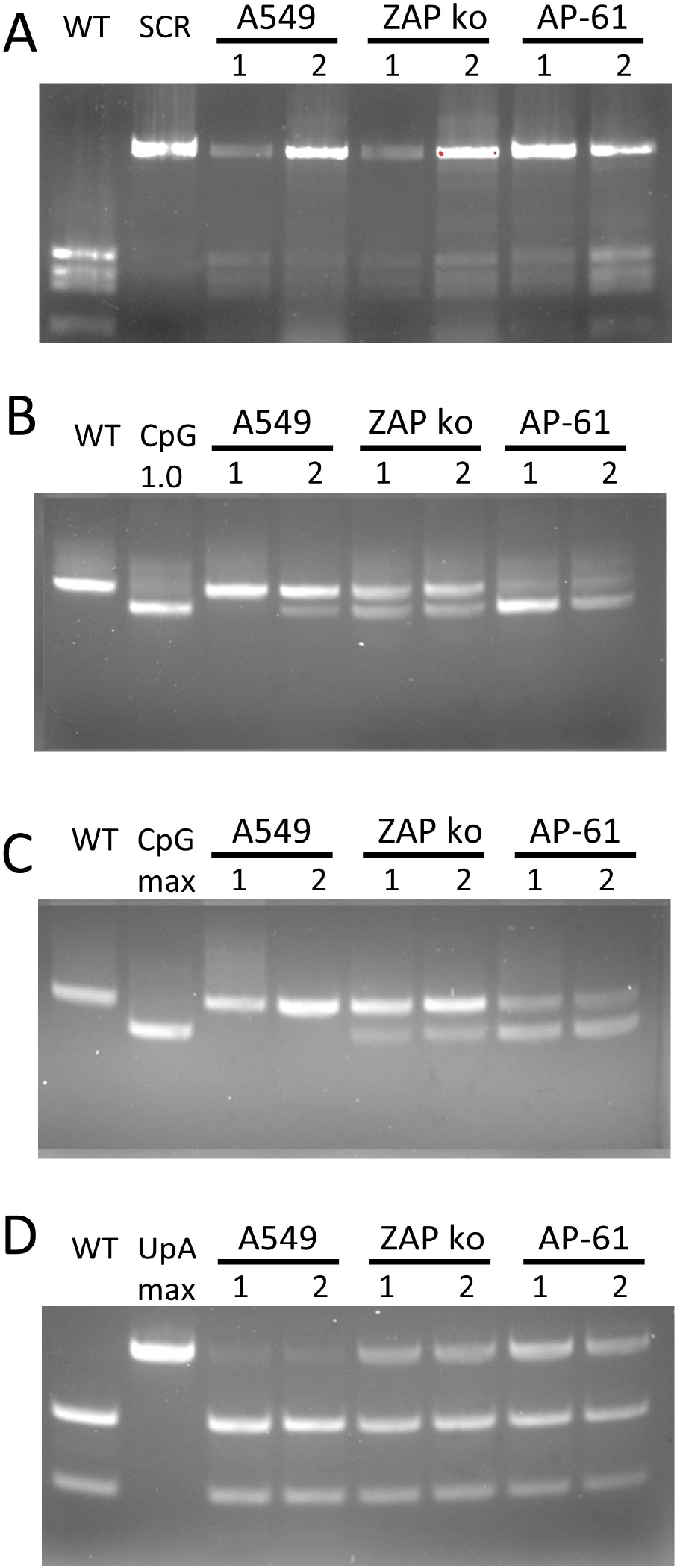
Fitness comparison between WT and mutant ZIKV in human and mosquito cells. Human A549 cells, A549 ZAP knockout cells and AP-61 mosquito cells were infected with equal RNA concentrations of WT ZIKV mixed with either the scrambled control virus (SCR) **(A)**, CpG-high viruses (CpG_1.0 and CpG_max) **(B,C)** or UpA_max **(D)**. Virus was passaged to new cells twice, before total RNA was isolated. ZIKV RNA was amplified with RT-PCR and digested with BsaI (A), SalI (B,C) or HeaIII (D) restriction enzymes to distinguish between WT and the respective mutant viruses. DNA gel images display the largest most distinctive products from each digestion. For more information see S3 Fig. From left to right images display the digested DNA of the indicated individual viruses, followed by two independent experimental repeats performed with separately rescued virus populations (numbered 1 and 2).

### CpG-high ZIKV is more infectious in *Aedes aegypti* mosquitoes

To investigate the biological role of dinucleotides frequencies in arbovirus RNA in living invertebrate vectors, *Aedes aegypti* mosquitoes were orally infected with WT ZIKV and the mutant viruses. Mosquitoes were fed an infectious blood meal containing a fixed number (10^8^/ml) of RNA copies, which for WT ZIKV equalled 5 x1 0^5^ TCID_50_/ml (Fig 5). Engorged females were selected and incubated for 14 days before mosquito saliva was obtained. Whole body mosquito homogenates and saliva samples were scored for the presence of ZIKV (Fig 5A). Strikingly, 70 % of the mosquito bodies became infected after ingesting CpG_1.0 ZIKV and 76 % after ingesting the CpG_max ZIKV mutant, while this was only between 26 and 37 % for mosquitoes that had ingested WT, SCR or UpA_max viruses (Fig 5B).

**Fig 5.**
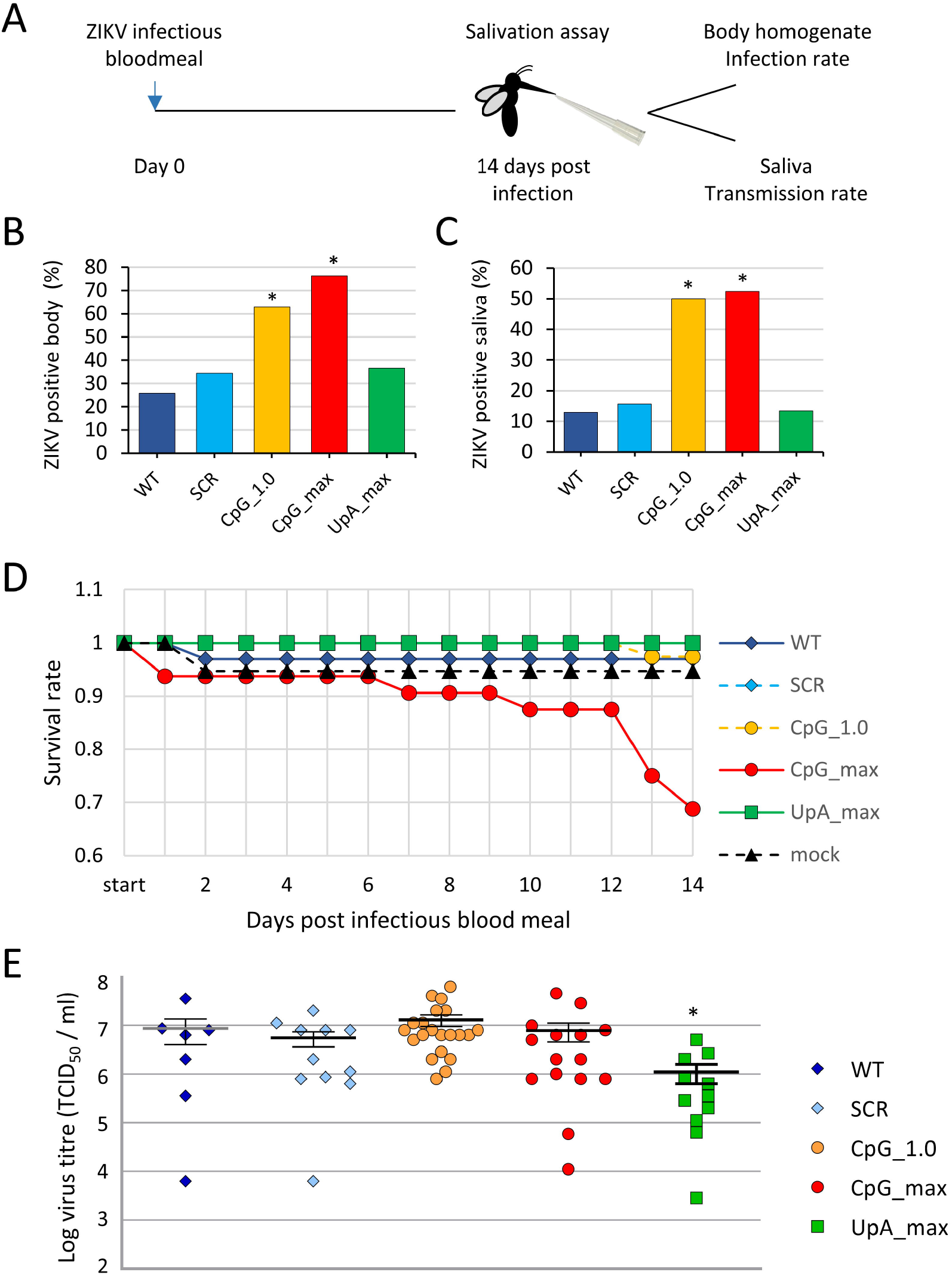
Infection of *Aedes aegypti* with ZIKV mutants. *Aedes* mosquitoes were offered an infectious bloodmeal containing equal RNA concentrations of each of the Zika viruses. Fully engorged females were selected and 14 days later subjected to a salivation assay **(A)**. Mosquito body homogenates **(B)** and mosquito saliva **(C)** were inoculated on Vero E6 cells to detect the presence of infectious virus. Bars represent the percentage of ZIKV positive samples with n between 21 and 36 engorged mosquitoes per experimental group. Asterisks represent significant difference from WT (Fisher exact test, P < 0.01). Survival rate of blood fed mosquitoes during the experiment was differentially affected by mutant viruses (Log-rank test, P < 0.001), with n between 31 and 39 mosquitoes that had engorged an infections blood meal at the start of the experiment **(D).** The infectious virus titre inside ZIKV positive mosquito bodies was determined by end point dilution assays. Data points indicate individual mosquito samples and the mean and SEM for each experimental group are displayed. Asterisk indicates significant difference from WT (Mann-Whitney U, P < 0.05) **(E)**.

Between 13 and 16 % of mosquitoes carried infectious WT, SCR or UpA_max ZIKV in their saliva. Significantly higher percentages of ZIKV positive saliva were detected among mosquitoes that fed on blood containing the CpG_1.0 (50 %) and CpG_max (52 %) ZIKV mutants (Fisher’s exact test, P < 0.01) (Fig 5C). This indicates that dissemination into the saliva also occurred more frequently when mosquitoes were infected with the CpG-high viruses. Interestingly, a notable mortality of mosquitoes that fed on the CpG_max ZIKV mutant was observed during the experiment (Log-rank test, P < 0.001) (Fig 5D). Despite the different infection rates, whole body homogenates of WT, SCR, CpG_1.0 and CpG_max ZIKV infected mosquitoes contained similar viral titers (approximately 10^7^ TCID_50_/ml) (Fig 5E). However, mosquitoes infected with UpA_max ZIKV displayed 10-fold lower viral titres (Mann-Whitney U, P < 0.05) (Fig 5E). In an independent experiment in which mosquitoes were fed blood meals containing equal infectious virus titres (5 x 10^5^ TCID_50_/ml) instead of equal RNA copy numbers, mosquitoes were again more often infected with the ZIKV mutants with elevated CpG frequencies, compared to WT and SCR ZIKV (S4 Fig) (Fisher exact test, P < 0.01).

### ZIKV mutants with elevated CpG and UpA dinucleotide frequencies show attenuated infection in IFNAR^−/−^ mice

To investigate the behaviour of ZIKV mutants in a mammalian host, groups of adult male IFNAR^−/−^ mice (defective in the interferon-α/β receptor) were infected with the WT and ZIKV mutants, or were injected with PBS as a negative control (Fig 6A). Weight loss and viremia were determined for each animal. During the first five days mice infected with WT or SCR ZIKV lost 6-8 % of their total body weight (Fig 6B). Infections with the CpG_1.0 and CpG_max ZIKV mutants resulted in a 5-6 % loss of total body weight. However, the mice infected with the CpG_1.0 and CpG_max ZIKV mutants recovered faster than mice infected with WT ZIKV and reached their respective initial body weights around day 29 and 23 post infection compared to 32 dpi for mice infected with WT ZIKV. Mice that were inoculated with UpA_max ZIKV or PBS showed no signs of disease and maintained a healthy body weight throughout the experiment (Fig 6B). Next, blood viremia levels in mice infected with both mutants that displayed reduced weight loss were compared to those infected with wildtype ZIKV. Mice infected with the CpG_max and UpA_max ZIKV mutants still developed substantial viremia, which was detected for seven days post infection. However, at peak viremia titres were on average 5-fold lower than for WT ZIKV (Fig 6C, day 3). All infections resulted in strong anti-ZIKV IgG antibody responses in week 4, though responses in CpG_max and UpA_max infected mice led to mildly lower (~2-fold) end-point ELISA titers compared to infections with WT, SCR and CpG_1.0 ZIKV (Fig 6D) (one-way ANOVA, F (4, 20) = 6.917, p < 0.05).

**Fig 6.**
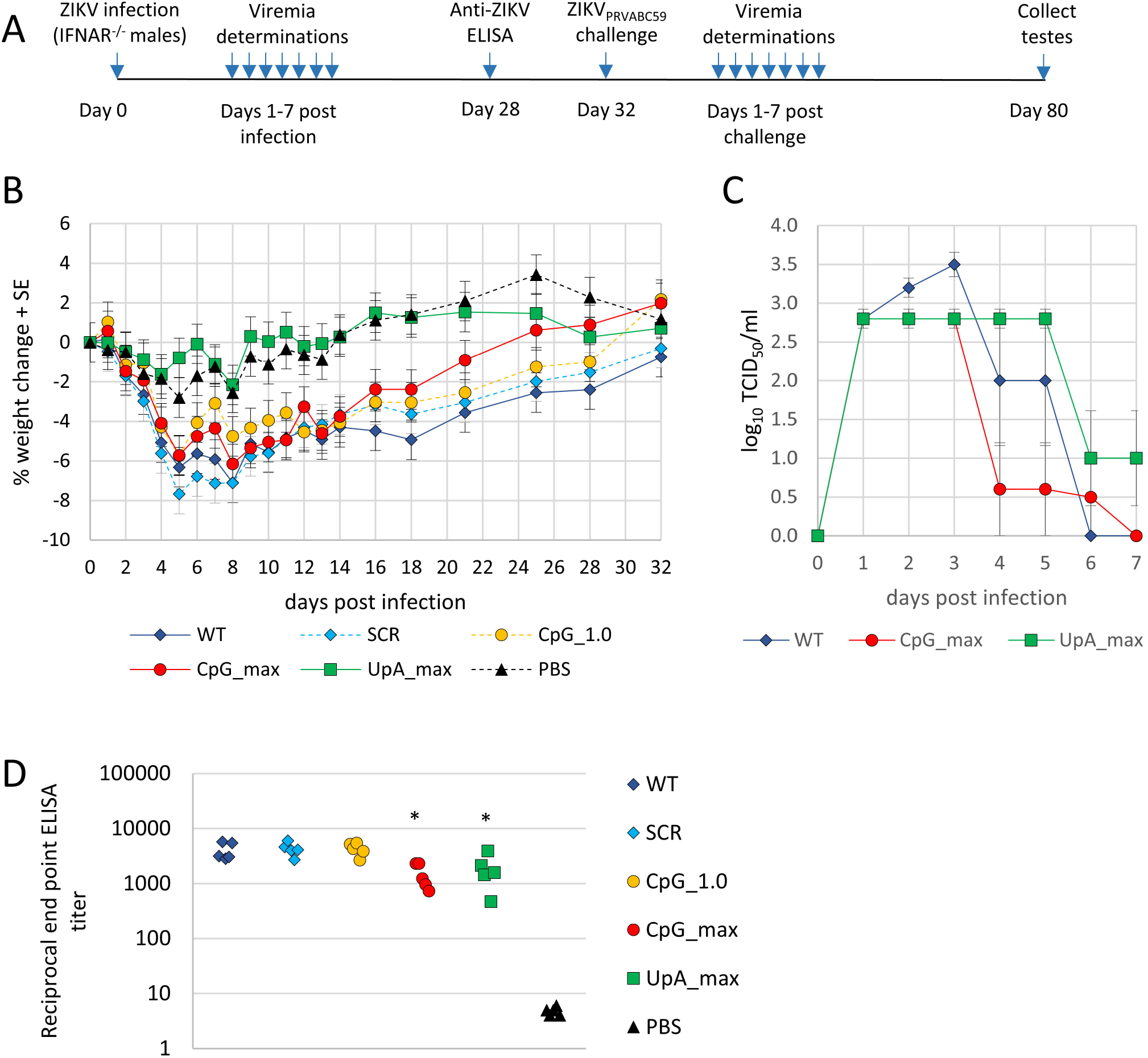
Weight loss, viremia and ZIKV antibody responses post primary ZIKV infection. **(A)** Timeline of experiment. **(B)** Weight loss of IFNAR^−/−^ mice post infection (n = 5 mice per group). **(C)** Viremia post infection of mice infected with WT ZIKV or CpG_max or UpA_max ZIKV. **(D)** 50% End-point of total IgG ELISA titers 4 weeks post-vaccination with indicated ZIKV mutants. Significant changes from WT are calculated by a One-way ANOVA with Dunett’s multiple comparisons test and indicated with asterisks (P < 0.05).

### Challenge with ZIKV_PRVABC59_

The mice were subsequently challenged with ZIKV_PRVABC59_ on day 32 after the immunising infection, with the ZIKV mutants (see Fig 6A). All the previously infected mice were protected against ZIKV_PRVABC59_-induced weight loss (Fig 7A). Only those mice that were mock-infected with PBS showed signs of weight loss and developed ZIKV viremia after the challenge (Fig 7A and B). In this IFNAR^−/−^ mouse model, ZIKV infection results in testicular damage, clearly seen as substantial overt reductions in testis size. The unvaccinated PBS control group displayed a marked reduction in testis size after challenge with ZIKV_PRVABC59_. Testis size reductions were also observed in mice immunised with WT ZIKV and the SCR and CpG_1.0 ZIKV mutants, likely as a result of the first infection rather than due to ZIKV_PRVABC59_ challenge, given that all mice seroconverted after the initial infection (Fig 6D). Mice that were first infected with the CpG_max and UpA_max ZIKV mutants retained normal testis size after challenge with ZIKV_PRVABC59_ (Fig 7C) (Kruskal-Wallis test, p < 0.05), illustrating that neither the first infection nor ZIKV_PRVABC59_ challenge resulted in detectable testes damage.

**Fig 7.**
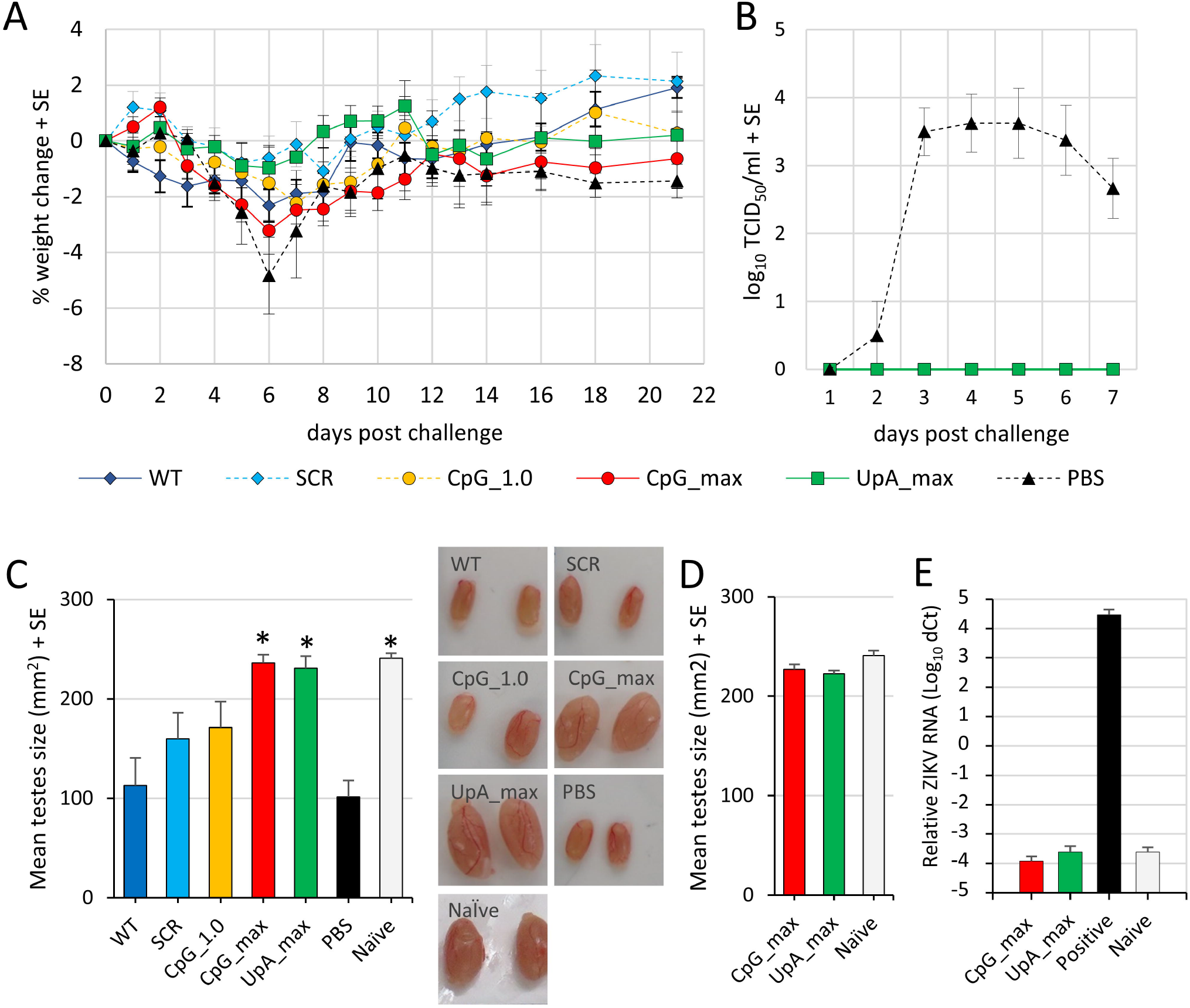
Weight loss, viremia and testes size in IFNAR−/−mice post ZIKV_PRVABC59_ challenge. **(A)** Weight loss of ZIKVPRVABC59-challenged IFNAR^−/−^ mice (n = 3-5 mice per group). **(B)** Viremia post-ZIKVPRVABC59 challenge (n = 4-5 mice per group). **(C)** Testes size (height x width) and representative images of testes from vaccinated mice post-ZIKV_PRVABC59_ challenge compared with testes of not vaccinated, PBS injected control mice and naïve (uninfected, healthy control) mice. Asterisk indicates significant difference from mice that received PBS injections instead of vaccination prior to ZIKV_PRVABC59_ challenge (Kruskal-wallis test, p < 0.05). **(D)** vaccine safety test displays the mean testis size of mice infected with CpG_max or UpA_max mutant viruses (n = 6 mice per group, testes harvested on day 49 post-vaccination) compared with testes size of naïve mice (n = 3) without subsequent challenge. **(E)** qRT-PCR of ZIKV RNA in testes of mice from D and mice infected with ZIKV_PRVABC59_ for 48 days as positive control. Bars represent the mean Ct values normalised for RPL13A housekeeping RNA.

The suitability of using these two mutants as a vaccine was further evaluated by infecting additional mice with either CpG_max or UpA_max ZIKV mutants, with the size of their testes determined 49 days post infection. No reduction in testes size was observed when compared to non-infected (naïve) mice (Fig 7D). qRT-PCR of the testes also illustrated that neither CpG_max nor UpA_max ZIKV mutants contained detectable levels of ZIKV RNA (Fig 7E). Together, these results indicate that the CpG_max and UpA_max ZIKV mutants do not cause testes damage and can protect against ZIKV challenge.

## Discussion

The motivation for this study was the evident difference in CpG dinucleotide representation in the genomes of RNA viruses infecting vertebrates compared to those infecting insects (Fig 1) [5, 13, 14, 31]. A recent study reported that ZIKV mutants with elevated CpG dinucleotide frequencies were attenuated in vertebrate cells and mice, while no significant effects on virus replication were observed in C6/36 *Aedes albopictus* cells [36]. This suggests that CpG suppression in arthropod-borne flaviviruses is required for sufficient replication in vertebrate hosts and that there is no selection on CpG dinucleotide frequencies in mosquitoes or other arthropod hosts. However, in our experiments ZIKV mutants with elevated CpG dinucleotide frequencies (CpG representations of 1.0 and 1.62) reproducibly showed enhanced replication rates in a range of different *Aedes spp*. mosquito cell lines (Fig 3). Importantly, live vector mosquitoes became more frequently infected with the CpG-high ZIKV mutants and dissemination into the saliva occurred more often compared to infections with WT ZIKV (Fig 4). In marked contrast to these pro-viral effects of increased CpG frequencies, elevated UpA dinucleotide frequencies resulted in reduced replication rates in mosquito cell cultures and lowered virus titers in infected *Aedes aegypti* mosquitoes (Figs 3 and 4). Taken together this study provides the first experimental evidence that dinucleotide composition of viral genomes is subject to substantial evolutionary pressures in invertebrate mosquito cells.

This new information sheds light on the biological basis for the compositional differences between flaviviruses that are arthropod-borne and those that replicate exclusively in arthropod hosts (ISFs). Both groups of flaviviruses persistently infect their arthropod hosts without any noticeable pathogenicity. The observation that elevated CpG frequencies enhance ZIKV replication in mosquito cells (Figs. 3 and 4) suggests that wildtype ZIKV infections in mosquitoes are normally attenuated. The substantial mortality in mosquitoes infected with the CpG_max ZIKV (Fig 4D) further illustrates how finely balanced CpG composition has to be maintained for flaviviruses to persistently infect their arthropod hosts, especially given that the mutated region represents only 15% of the ZIKV genome. Lineage 2 ISFs mildly suppress CpG dinucleotides (Fig 1) and are believed to have evolved from arboviruses that lost the ability to infect vertebrate cells [31]. Rather than drifting towards an unbiased CpG dinucleotide frequency due to a lack of selection pressure, the observed evolutionary pressures in mosquito cells suggest that lineage 2 ISFs accumulate CpG dinucleotides by natural selection. However, the accumulation of CpG dinucleotides in lineage 2 ISF genomes is potentially constrained as non-pathogenic persistent replication needs to be maintained.

This delicate balance additionally has to be sustained in the context of antiviral responses. In the arthropod host, the most potent antiviral responses are mediated through RNAi (reviewed in [37]). Suppression of RNAi in mosquitoes infected with Sindbis virus (arbovirus from family *Togaviridae*, genus *Alphavirus*) results in increased mosquito mortality [38]. However, replication of the ZIKV mutants with elevated CpG or UpA dinucleotide frequencies was similarly enhanced or suppressed, respectively, in all mosquito cells tested, including C6/36 cells (Fig 3), which are defective in the production of antiviral siRNAs [39, 40]. This suggests that other pathways or mechanisms are active in discriminating dinucleotide frequencies in mosquitoes.

Alternatively, synonymous changes in dinucleotide frequencies inevitably alter the codon usage of a nucleotide sequence and changes in translation efficiency may affect the bio-synthesis rate of the encoded viral polyprotein and as a consequence viral replication rates. However, comparison of the synonymous codon usage of the mutated region in ZIKV and the synonymous codon frequency of human and insect reference sets show no consistent correlation between the elevation of CpGs and similarities in codon usage with either host reference set (S1 Table). Indeed, previous studies have largely discounted effects of codon use or codon pair bias as the explanation for attenuation of CpG-high mutants of vertebrate viruses. For example, elevation of CpG and UpA dinucleotide frequencies in untranslated regions of viral RNA attenuated virus replication to equivalent degrees as similar dinucleotide modifications in coding regions of the viral genome [15, 16, 41]. Furthermore, it is now established that the vertebrate antiviral protein ZAP mediates the attenuated replication of viruses with high CpG frequencies.

In vertebrates, the N-terminal zinc-fingers of ZAP bind high CpG containing RNA sequences through a number of hydrogen bonds and stacking interactions [22] to initiate a plethora of downstream intracellular interactions with proteins involved in translation, RNA degradation and antiviral immunity [20, 23–28, 42]. Specifically, the host proteins RNaseL, activated by OAS3, and both TRIM25 and KHNYN are crucial for the ZAP-mediated attenuation of CpG-high E7 and HIV-1, respectively [16, 21, 29]. Correspondingly, knockout of ZAP or OAS3/RNaseL abolished the attenuated phenotype of ZIKV mutants with elevated CpG dinucleotide frequencies in vertebrate cells (Fig 2). Additionally, mice vaccinated with CpG_max ZIKV and subsequently challenged with ZIKV_PRVABC59_ did not show any signs of ZIKV induced testicular damage as measured by their testis size, whereas vaccination with CpG_1.0 ZIKV resulted in an intermediate reduction in average testis size (Figs 6 and 7). Together, these results indicate that the degree of virus attenuation is correlated to CpG composition and mediated by vertebrate ZAP.

While most recent investigations involving genome composition and virus replication have focussed on CpG dinucleotides, a growing body of evidence indicates a substantial additional effect of UpA dinucleotide frequencies on virus attenuation. UpA dinucleotides are suppressed in vertebrate and invertebrate animals and their viruses (Fig 1) [14]. The UpA_max ZIKV mutant used in this study showed strong attenuation in vertebrate cells and no signs of disease in mice (Figs 2 and 6). Moreover, the attenuated phenotype of UpA_max ZIKV was rescued by knockout of ZAP, suggesting that the interaction between ZAP and UpA dinucleotides is not exclusive for previously published E7 mutant viruses [21]. Whether ZAP directly binds UpA dinucleotides or requires additional host proteins for this interaction is unknown and remains a topic of active investigation [43]. Whether the selection against UpA dinucleotides in insect cells is equally strong as that observed in vertebrate cells remains unclear. Although flaviviruses suppress UpA dinucleotides in their RNA, UpA_max ZIKV replication and dissemination was not consistently reduced compared to WT ZIKV in mosquito cells and live mosquitoes, respectively (Figs 3–5).

The diverse selective pressures that arboviruses encounter have influenced arbovirus disease emergence on multiple levels (Reviewed in [44]). The current study identifies CpG frequencies as a key adaptive trait that enables ZIKV to replicate in two profoundly different hosts. Furthermore, the finding that increasing CpG frequencies enhanced the replication of ZIKV in mosquitoes provides evidence that CpG frequency differences between vector-borne flaviviruses and ISFs do not simply result from compositional constraints in one host and none in the other. Instead, the data indicates the existence of strongly conflicting evolutionary pressures on ZIKV to enable sufficient replication in both hosts to maintain a vector-borne life cycle. In contrast to flaviviruses, vector-borne viruses from the genus *Alphavirus* (Family *Togaviridae*) contain relatively high CpG dinucleotide frequencies, which may suggest these viruses have evolved a more optimal genome composition for replication in the mosquito vector. Although most alphaviruses effectively infect vertebrate cells, these viruses are highly sensitive to ZAP [45]. The same constraints may operate in other dual-host viruses. For example, we and others recently discovered that the replication of potato virus Y and plum pox virus (genus *Potyvirus*) in plants was similarly attenuated by the synonymous introduction of CpG and UpA dinucleotides [46, 47]. Plant transcriptomes also display CpG suppression that is also reflected in the majority of plant RNA viruses. This suggests that plant viruses which are maintained in transmission cycles involving persistent replication in both plant host and arthropod vectors [48] may encounter similarly conflicting evolutionary pressures as arboviruses. The findings can spark investigations into novel strategies to intervene with the transmission cycles of arboviruses. Attenuating vertebrate viruses by increasing CpG and UpA dinucleotide frequencies is a promising and universally applicable approach to generate live-attenuated virus vaccines (Figs 6 and 7) [17, 36]. The enhanced replication of CpG-high mutants of ZIKV in mosquito cells may serve to improve production of live attenuated or inactivated vaccines and can lead to novel immunisation strategies [49]. A radical, yet intriguing application is to release CpG-high infected mosquitoes into the environment to serve as vaccine vectors. At the moment this application remains purely hypothetical as such an approach raises many questions regarding ethics, vaccine safety and the spread of genetically modified viruses into the environment. However, theoretically mosquitoes could be highly effective (Fig 5) in not only vaccinating humans and livestock, but also animal reservoirs by transmitting high doses of flavivirus vaccines with the same attenuated replication in vertebrate animals as observed in the experimentally infected mice (Figs 6 and 7).

## Acknowledgements

The authors thank Prof. Andres Merits (University of Tartu) for sharing the ZIKV_BeH819015_ pCCI-SP6 cDNA clone, Prof. Roy Hall and Dr Jody Hobson-Peters (University of Queensland, Australia) for sharing RML-12 *Aedes albopictus* mosquito cells, 4G2 antibody and MR766 ZIKV antigen, and Dr Astrid Bryon, Dr Mark Sterken and Marleen Henkens for their support and technical assistance. A. Suhrbier is a leadership fellow of the National Health and Medical Research Council (NHMRC) of Australia (APP1173880) which in part funded this research via project grant APP1144950.

## Materials and methods

### *In silico* analyses

A single full length flavivirus genome was selected per flavivirus species. For analysis of host gene composition, mRNA sequences of human (GRCh38.p13i) and *Aedes aegypti* (AaegL5.0) were downloaded from RefSeq (https://www.ncbi.nlm.nih.gov/nuccore). From these datasets, representative selections of 10000 sequences were extracted at random using Perl with an in-house script. Sequences were filtered to exclude coding region sequences of length <250 bases. To avoid redundancy, similar or identical sequences (< 1% nucleotide divergence) were identified with SSE software (v1.4) [50] and removed from the dataset. This resulted in over 9000 human and mosquito sequences, which were used for composition analysis (all GenBank accession numbers available in S1 Appendix). Dinucleotide frequencies were calculated using the program Composition Scan within SSE (v.1.4). Synonymous site variability (SSV) and mean folding energy differences (MFED) of the ZIKV genome were calculated as previously described, using a fragment size of 200 and sliding window of 21 codons for SSV and a fragment size 200 with 48 nucleotide increments for MFED using a selection of ZIKV isolates [51]. The codon adaptation index (CAI) of full length ZIKV and the mutated regions was calculated using an online CAI calculator (www.biologicscorp.com) with the codon usage of *Homo sapiens* and *Drosophila melanogaster* as reference data sets (S1 Table).

### Cells and viruses

*Ae. albopictus* C6/36 (ATCC CRL-1660) and U4.4 were cultured at 28 °C in Leibovitz-L15 medium (Gibco, Carlsbad, CA, USA) supplemented with 10% heat-inactivated fetal bovine serum (FBS; Gibco), 2% tryptose phosphate broth (Gibco) and 1% nonessential amino acids (Gibco). *Ae. albopictus* RML-12 cells were cultured in Leibovitz’s L-15 medium supplemented with 10% FBS and 10% tryptose phosphate buffer at 28 °C [52]. *Ae. aegypti* Aag2 cells [53], *Ae. pseudoscutellaris* AP-61 cells [54] were cultured at 28 °C in Schneider’s Drosophila medium (Gibco) supplemented with 10% FBS. Human A549 cells including ZAP, Ribonuclease L (RNaseL), 2’-5’-oligoadenylate synthetase 1 (OAS1) and OAS3 knockout cells were described previously [21] and African green monkey kidney Vero E6 (ATCC CRL-1586) were all maintained at 37°C with 5% CO_2_ in Dulbecco’s modified Eagle medium (DMEM) (Gibco) supplemented with 10% heat-inactivated FBS, 100 U/ml penicillin and 100 μg/ml streptomycin (Sigma-Aldrich). Cell lines were routinely checked for mycoplasma using MycoAlert™ Mycoplasma Detection Kit (Lonza).

A pCC1-BAC–based and SP6 promoter-driven infectious clone of the ZIKVBeH81901 isolate from Brazil (GenBank KU365778.1) [55] was used for the construction of mutant viruses. A region of 1653 nucleotides was selected for the described synonymous mutations. Mutant regions with elevated CpG or UpA dinucleotide frequencies were designed using the ‘mutate sequence’ function in SSE, taking care not to alter the coding amino acid sequence or the other UpA or CpG dinucleotide, respectively. The SCR sequence was originally designed for a region of 4344 nucleotides that included the above mentioned mutated region using the ‘scramble sequence’ function in SSE with the CDLR algorithm. The amino acid sequence, codon usage and dinucleotide usage of the SCR sequence were kept identical to that of the corresponding WT sequence. Mutated regions were synthesized (Geneart) and incorporated into the viral genome using unique restriction sites RsrII and Bst BI (New England Biolabs). Nucleotide sequences of mutated regions are listed in S2 Appendix. Plasmid DNA was linearized with PmeI (New England Biolabs) and complete digestion was verified on agarose gel. Linearized DNA was purified by phenol/chloroform extraction and RNA was *in vitro* transcribed in the presence of G(5’)ppp(5’)G RNA Cap structure analog using SP6 polymerase (New England Biolabs) according to the manufacturer’s protocol. RNA was transfected into Vero cells with lipofectamine 2000 (Invitrogen) to launch virus replication. Between one and four days post transfection, cytopathic effects were observed and cell culture supernatant was transferred to a fresh 60% confluent monolayer of Vero cells to generate passage 1 (P1). After two hours, inoculum was removed and cells were washed with cell culture medium. Three days post infection cell culture supernatant was harvested. Newly generated mutant virus stocks were sequence verified by reverse transcriptase PCR (see competition assay below for details) followed by restriction analysis using DdeI, SalI and PstI restriction enzymes (New England Biolabs) and DNA sequencing (Macrogen Europe). Stocks were again verified by restriction digestion prior to the mouse experiments. At the end of the mosquito experiments, total RNA from pools of eight infected mosquitoes was isolated and ZIKV mutants were sequence verified by RNA-sequencing (BGI Genomics) as described previously [56].

### Virus quantification

Tissue culture infectious dose 50% (TCID_50_) was determined by end-point dilution assays (EPDA). Serial 10-fold dilutions of virus stocks and experimental samples were prepared in complete DMEM cell culture medium. Vero E6 cells from confluent tissue cultures were detached with Trypsin-EDTA (Gibco) and dissolved in a total volume equal to 1 ml / 1.67 cm^2^. Cell suspension was added to the virus dilutions in a 1: 1 ratio, mixed and plated in 6-fold onto micro-titer plates (Nunc). Virus titres were determined by cytopathic effects (CPE) at the end-point of infection. For each of the mutant viruses the viral titres determined by CPE and respective end-points of the EPDA were confirmed by immunofluorescence. Cells were washed with phosphate buffered saline (PBS) and fixed with 4% paraformaldehyde in PBS. ZIKV envelope was stained with 4G2, a pan-flavivirus primary α-E mouse monoclonal antibody diluted in PBS containing 2 % FBS (1:50) [57] for 1 hour at 37 °C. Samples were washed three times with PBS and stained with a secondary goat α-mouse Alexa Fluor 488 (1:2000; Invitrogen). RNA copy numbers in virus stocks were determined by comparing RNA from virus stocks with purified SP6 *in vitro* transcribed RNA. RNA isolation was performed with TRIzol reagent for liquid samples according to the manufacturer’s protocol. Reverse transcriptase and a real-time PCR were performed with two sets of ZIKV specific primers F1 5’-cctgatgaccatctgtggca-3’, R1 5’-cccatagagcaccactcctt-3’ and F2 5’-tcaggaggtggtgttgaagg-3’, R2 5’-gtgccctttctccatttggt-3’ using superscript III (Invitrogen) and SYBR green (Applied Biosystems) according to the respective manufacturer’s protocols with an annealing temperature of 58 °C during real-time PCR in a CFX96 real-time system (Bio-Rad) thermocycler.

### Virus growth curves

Cell monolayers were seeded in 6-well plates to a 50 – 60 % confluency. The cell culture fluid was removed and infections were performed by diluting virus stocks (P2) in standard culture media for each of the respective cell types in a total volume of 1 ml / well. After 2 hours the inoculum was removed and the monolayers were washed twice with 1 ml of PBS, before 2 ml of fresh culture medium was added. At indicated times post infection 50 μl of cell culture supernatant was taken per well and stored at −80°C until TCID_50_/ ml was determined by EPDA.

### Competition assay

Equal concentrations of WT and mutant virus (1 RNA / cell) were mixed in DMEM or XXX cell culture medium and added to A549, A549 ZAP knockout or AP-61 mosquito cells at 50% confluence. After two hours, the inoculum was replaced with culture medium. Ten μl of cell culture supernatant was passaged to fresh cells. After two passages total RNA was isolated using TRIzol reagent according to the manufacturers protocol. RNA was reverse transcribed and part of the mutated region within the ZIKV genome was PCR amplified using SuperScript III One-Step RT-PCR System with Platinum Taq Polymerase (Invitrogen) together with either the 5’-aagtggagaaaaagatggg-3’ (WT and SCR) or 5’-gaggaaaaagagtggaagac-3’ (WT, CpG_1.0, CpG_max and UpA_max) forward primers in combination with a 5’-gtgccctttctccatttggt-3’ reverse primer. Reverse transcription and PCR amplification were performed according to the manufacturers protocol with the reverse transcription at 50 °C for 30 minutes and an annealing temperature during PCR amplification of 58 °C in a in an Applied Biosystems 2720 thermocycler. The resulting DNA amplicons from the comparisons between WT and SRC were digested with BsaI-HFv2 (New England Biolabs), WT and CpG_1.0 or CpG_max with SalI-HF (New England Biolabs) and WT and UpA_max with HaeIII (Invitrogen) and ran on 2% agarose gels.

### Mosquito rearing

*Aedes aegypti* mosquitoes (Rockefeller strain, Bayer AG, Monheim, Germany) were maintained at 27±1°C with 12:12 light:dark cycle and 70% relative humidity. Adult mosquitoes were provided with 6% *ad libitum* glucose solution. Human blood (Sanquin Blood Supply Foundation, Nijmegen, The Netherlands) was provided through Parafilm using the Hemotek PS5 feeder (Discovery Workshops, Lancashire, United Kingdom). Adult mosquitoes were kept in Bugdorm-1 insect rearing cages (30 x 30 x 30 cm, Bugdorm, Taiwan, China) until they were 2 −6 days old, before females were transferred to buckets (diameter: 12.2 cm, height: 12.2 cm; Jokey, Wipperfürth, Germany). Buckets contained approximately 50 females each and were transported to the Biological Safety Level 3 facility for virus infection assays.

### Infectious blood meal

Virus solutions were made by diluting the virus stocks to equal concentrations in DMEM before mixing with human blood to their final concentrations. Infectious blood meals were offered through Parafilm using the Hemotek PS5 feeder. Mosquitoes from a single bucket were offered a single blood meal containing one of the ZIKV mutants and allowed to feed for 1 h *ad libitum* in light conditions, at 24°C and 70% relative humidity (RH). Mosquitoes were anesthetized with 100% CO_2_ and fully engorged females were selected. Engorged mosquitoes were kept for 14 days on glucose solution at 28°C in a 12:12 light:dark cycle.

### Salivation assay

Fourteen days post blood meal mosquitoes were anesthetized with 100% CO_2_, and placed on a CO_2_ pad. Mosquitoes that died within the 14 days incubation period were counted and discarded. Mosquitoes were immobilized by removing their legs and wings. The proboscis of each mosquito was inserted into a 200 μl yellow pipet tip (Greiner Bio-One) containing 5 μl of a 1:1 solution of 50% glucose solution and FBS, for 1 hour. After salivation, the mosquito bodies were added to 1.5 ml Safe-Seal micro tubes (Sarstedt, Nümbrecht, Germany) containing 0.5 mm zirconium beads (Next Advance, Averill Park, NY, United States). Each saliva sample was added to a 1.5 ml micro tube (Sarstedt) with 55 μl DMEM cell culture medium containing FBS, Penicillin and streptomycin supplemented with additional Fungizone (50 μg/ml; Invitrogen), and Gentamycin (50 μg/ml; Life technologies), hereafter referred to as DMEM-complete. Mosquito bodies and saliva samples were stored at −80°C until further processing.

### Infectivity assays

Frozen mosquito bodies were homogenized for 2 min at speed 10 in a Bullet Blender Storm (Next Advance). Homogenized bodies were centrifuged briefly and resuspended in 100 μl DMEM-complete medium. Homogenization was repeated and mixtures were centrifuged for 1 min at 14.500 rpm in a table top centrifuge. From the mosquito homogenates or thawed saliva samples 30 μl was used to inoculate a Vero cell monolayer in a 96 wells plate. After 2–3 h the inoculum was removed and replaced with 100 μl DMEM-complete. Wells were scored for virus induced CPE at 3 and 4 dpi. Bodies and saliva samples of infected mosquitoes were additionally titrated by EPDA.

### Immunostained plaque assays

An 80 % confluent monolayers of cells were infected with a series of 10-fold dilutions with equal RNA copies of each of the mutant viruses. After 2 hours of incubation the inoculum was removed and a 1 % carboxymethyl cellulose overlay was added. At the end of the experiment, the overlay was removed and cells were fixed and stained for ZIKV envelope protein using 4G2 primary antibodies and green alexafluor 488 secondary fluorophores. Micrographs were taken with an Axio Observer Z1m inverted microscope in combination with an X-Cite 120 series lamp and AxioVision software of all primary plaques at the lowest possible concentration.

### Infection of IFNAR^−/−^ male mice with mutant ZIKVs

All viruses were grown in C6/36 cells and titers were determined by immunostained TCID_50_ assays using C6/36 cells. Briefly, virus was titrated in 10-fold serial dilutions in 96 well plates with C6/36 cells (2×10^4^ cell/well). After 5 days plates were washed and fixed in 80% cold acetone for 1 hr at −20°C. Acetone was removed. Plates dried and ZIKVs were detected by indirect immunofluorescence using mouse anti-4G2 monoclonal primary antibody and polyclonal goat anti-mouse IgG conjugated IRDye 800CW secondary antibody (LiCor, 926-32210). Staining was visualized using the LI-COR Biosciences Odyssey Infrared Imaging System and application software. Any staining in any well was considered positive and TCID_50_ titers determined.

Male type I IFN receptors knockout (IFNAR^−/−^) mice (C57BL/6J background) (n=5 per group, age 2-6 months) were infected subcutaneously (s.c.) at the base of the tail with 10^3^ TCID_50_ (100 ul in Roswell Park Memorial Institute (RPMI) 1640 supplemented with 2% FCS) of each virus per mouse. Mice were weighed at the indicated times and viremias determined using immunostained TCID_50_ assays as above. Briefly, mouse serum was titrated in 10-fold serial dilutions into 96 well plates with C6/36 cells and ZIKVs were detected by indirect immunofluorescence as above, except background IgG was blocked prior to addition of the secondary antibody with Goat F(ab) antimouse IgG (H&L) (Abcam).

### Antigen-coated ELISA

Anti-ZIKV IgG titers in mouse sera were determined by antigen-coated ELISA. Plates were coated with MR766 antigen in BupH Carbonate-Bicarbonate buffer (ThermoFisher) and incubated O/N at 4°C. After incubation, plates were blocked with 5% skim milk in PBS. Next, the mouse sera were added in triplicates diluted 1:30 in PBS-Tween containing 1% skim milk and incubated for one hour at room temperature. After incubation, plates were washed with PBS-Tween. Subsequently, secondary antibody HRP (1:2000 in PBS-Tween with 1% skim milk) was added and incubated for 1 hour at room temperature. Samples were washed with PBS-Tween and scored after one-hour incubation in the dark with ABTS substrate (1:1000 H_2_O_2_).

### ZIKV_PRVABC59_ challenge

An infectious clone ZIKV_PRVABC59_ (GenBank: MH158237.1) was constructed as described previously [58] and was provided by S. Tajima (Department of Virology I, National Institute of Infectious Diseases, Tokyo, Japan). Infectious virus was recovered by transfection of Vero E6 cells. ZIKV _PRVABC59_ stocks were grown in sub-confluent C6/36 mosquito cells maintained in RPMI1640 supplemented with 2% FBS and stored in −80°C. Viral stock titres (TCID_50_) were determined by 10-fold serial titration in C6/36 cells (2×10^4^ cells/well of a 96 well plate) and incubated for 5 days. After incubation, 25 μl from each well was placed into parallel plates containing Vero E6 (2×10^4^ cells/well), and the cytopathic effect assessed on day 7.

On day 32 post primary injections, mice were challenged s.c. base of tail with ZIKV_PRVABC59_ (10^3^ TCID_50_ in 100 ul) as described previously [49]. Five mice were mock injected with 100 μl PBS and used as uninfected controls. Mice were weighed at the indicated times. Mouse serum was collected by tail vein bleed and ZIKV_PRVABC59_ titers determined by TCID_50_ assays as described above. Mice were euthanized on day 80 post-primary infection, testes were removed, photographed and measured (height x width).

### Vaccine safety

To illustrate that the CpG_max and UpA_max ZIKV mutants do not cause testes damage and are suitable for use as attenuated vaccines, two groups of male IFNAR^−/−^ mice (n = 6 mice per group) were infected with these ZIKV mutants (10^3^ TCID_50_) and testes harvested 49 days post-infection and compared to testes from uninfected mice. qRT-PCR for ZIKV RNA was also performed on the testes as described previously [49].

### Statistics

Statistical analyses were done with the software package Prism (v. 9.0.0) and Microsoft Excel 2016 and detailed in the main text and in the relevant Fig legend. All two-way analyses of variance (ANOVA) were performed to compare replication of ZIKV variants in multiple cell lines. Outcomes were corrected for multiple comparisons by controlling the false discovery rate (FDR) (Q = 0.05). Reported p-values for two-way ANOVAs are FDR adjusted p-values unless stated otherwise.

### Ethics statement for murine experiments

All mouse work was conducted in accordance with the “Australian Code for the care and use of animals for scientific purposes” as defined by the National Health and Medical Research Council of Australia. Mouse experiments and associated statistical treatments were reviewed and approved by the QIMR Berghofer Medical Research Institute animal ethics committee (P2195).

---

**S1 Fig. Antiviral pathways involved in replication of ZIKV mutants**. Growth curves on RNaseL knockout **(A)**, OAS1 knockout **(B)** and OAS3 knockout **(C)** A549 cells. Cells were infected with mutant ZIKV viruses at 1 RNA / cell. At the indicated days post infection (dpi) the 50% tissue culture infectious dose was determined by end point dilution assays. Data points represent the average of two biological experiments and the error bars indicate one standard error of the mean. **(D)** Representative images of ZIKV induced immunoplaques in A549 and A549 ZAP knockout cells. Cells were fixed and stained for ZIKV E protein with primary antibody 4G2 and secondary Alexafluor 488 (green). Results of an immunoplaque assay on Vero E6 cells **(E)**. Data points represent the average plaque size, error bars indicate one standard error of the mean and asterisks highlight significant differences from WT (P < 0.05, One-way ANOVA with Dunnett’s post Hoc test).

**S2 Fig. Replication of ZIKV with increased CpG dinucleotides has a replication advantage in mosquito cells**. *Aedes albopictus* mosquito cell lines: C6/36 **(A)**, U4.4 **(B)** and RML-12 **(C)** were infected with the mutant ZIKV viruses at 10 RNA / cell. At the indicated days post infection (dpi) the 50% tissue culture infectious dose was determined by end point dilution assays. Data points represent the average of two independent biological experiments and the error bars indicate one standard error of the mean.

**S3 Fig. Images from DNA gel electrophoresis**. Complete and unmodified gel images corresponding to Fig 4. WT ZIKV mixed with either the scrambled control virus (SCR) **(A)**, CpG-high viruses (CpG_1.0 and CpG_max) **(B,C)** or UpA_max **(D)**.

**S4 Fig. Replication of ZIKV mutants in *Aedes aegypti***. Aedes mosquitoes were offered an infectious bloodmeal containing equal infectious virus titres of each of the Zika viruses. Mosquito body homogenates **(A)** and mosquito saliva **(B)** were inoculated on Vero E6 cells to detect the presence of infectious virus. Bars represent the percentage of ZIKV positive samples with n varied between 13 and 32 engorged mosquitoes per experimental group. Asterisks represent significant difference from WT (Fisher exact test, P < 0.01). The infectious virus titre inside ZIKV positive mosquito bodies was determined by end point dilution assay **(C)**. Data points indicate individual mosquito samples and the mean and SEM for each experimental group are displayed.

## Notes

### Competing Interest Statement

The authors have declared no competing interest.

