## Supplementary figures and images for "An adaptive compromise - Conflicting evolutionary pressures on arthropod-borne Zika virus dinucleotide composition in mammalian hosts and mosquito vectors"

### Supplement Figure 1

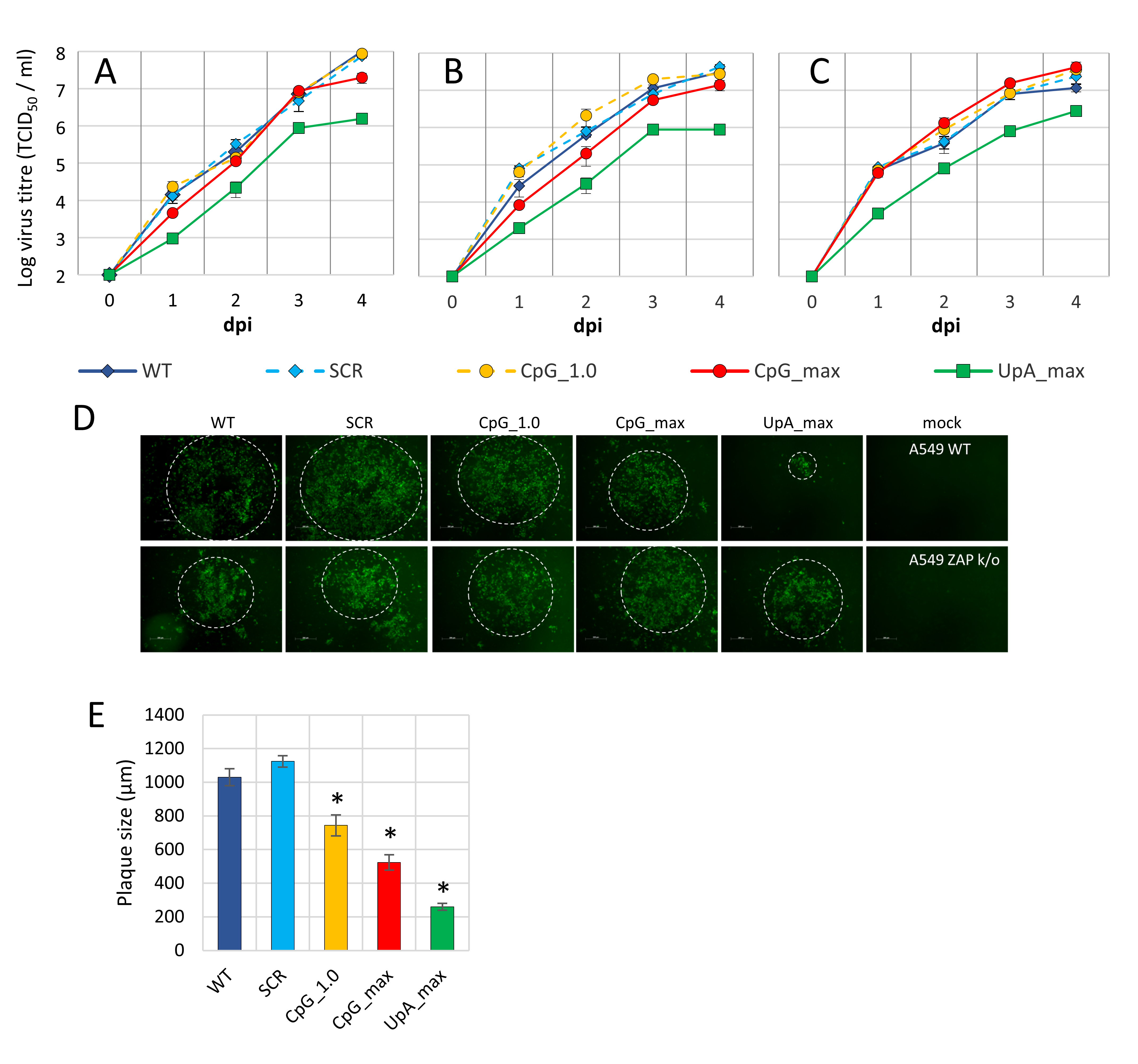

### Supplement Figure 2

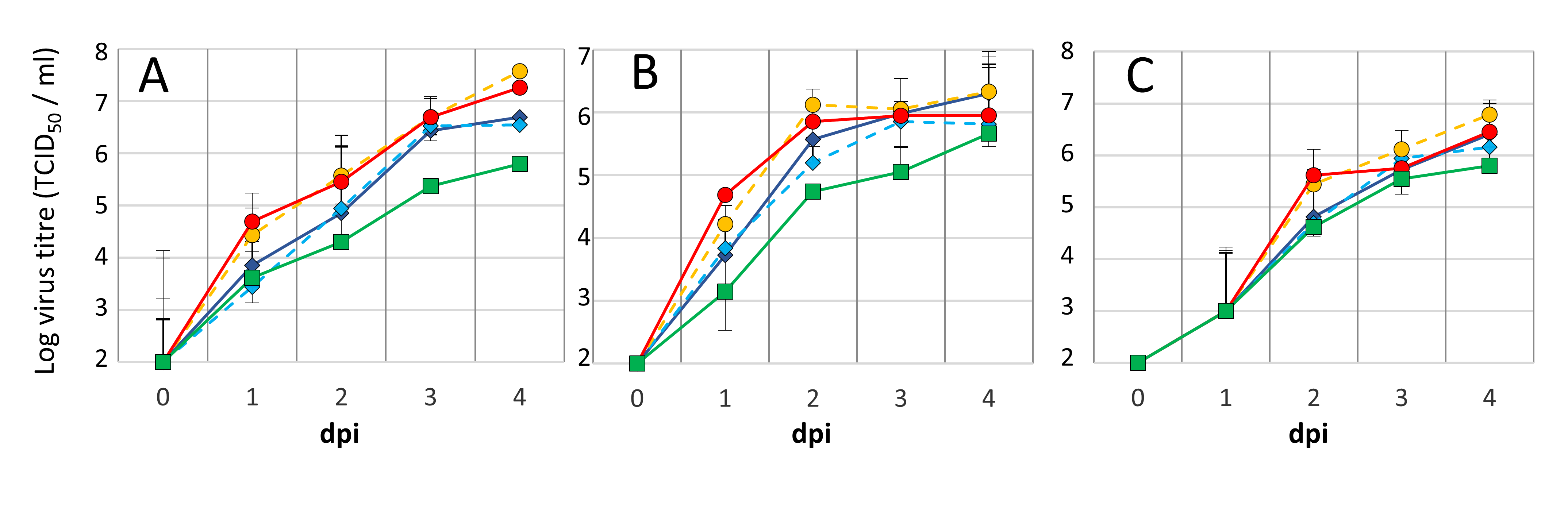

### Supplement Figure 3

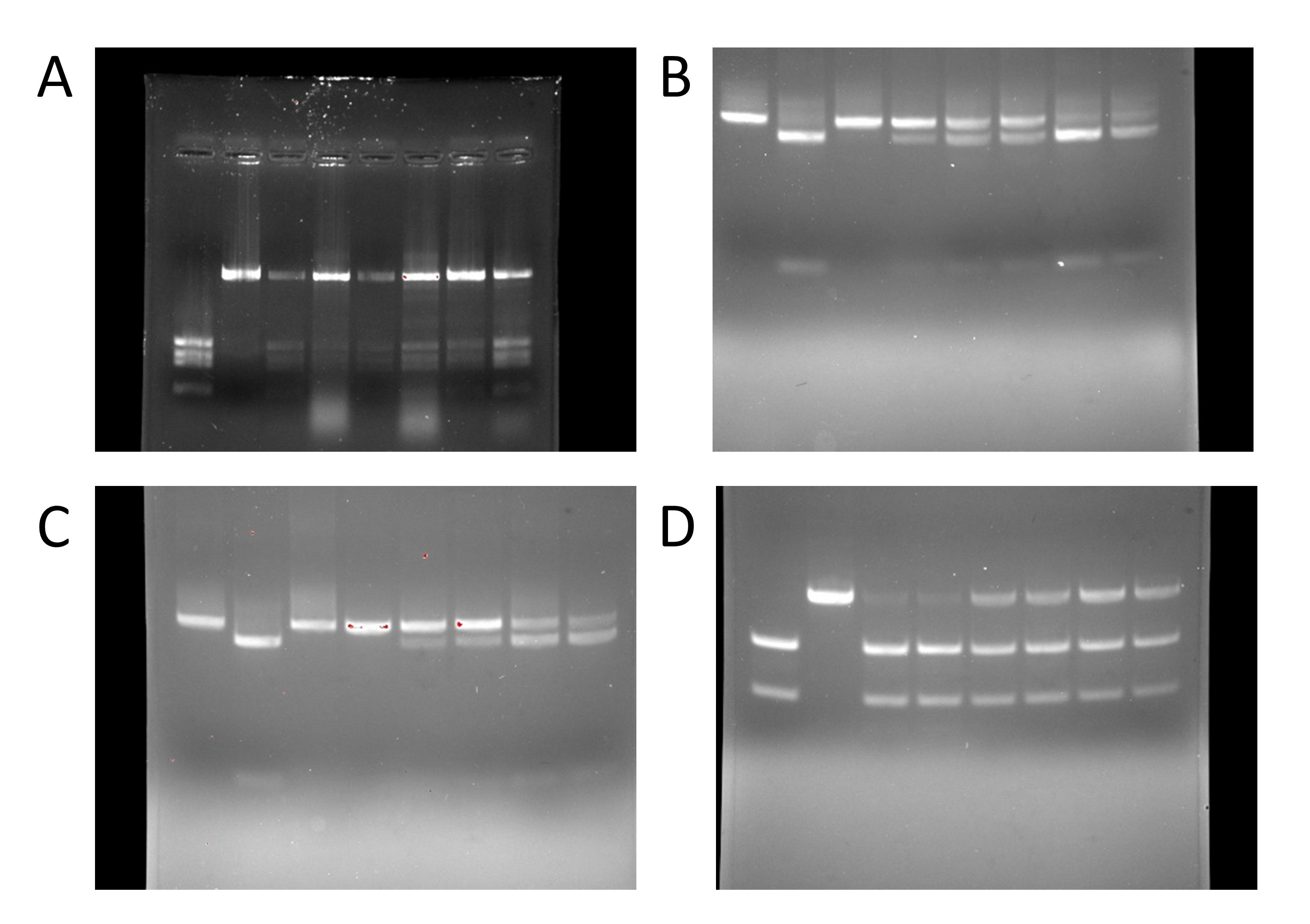

### Supplement Figure 4

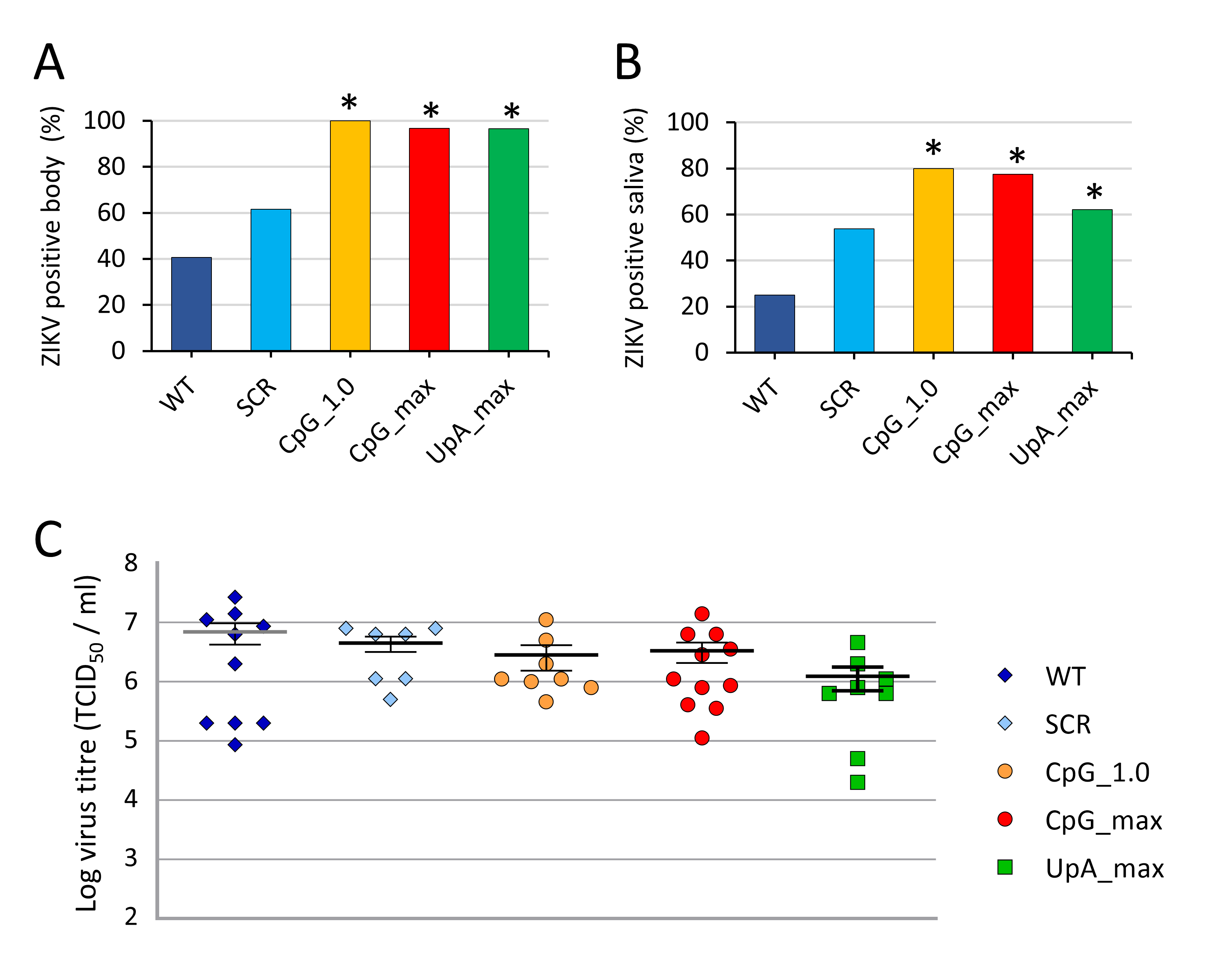
