## Supplement Table 1 for "An adaptive compromise - Conflicting evolutionary pressures on arthropod-borne Zika virus dinucleotide composition in mammalian hosts and mosquito vectors"

Supplement table 1 – Composition of mutated region

| | *Total nt* | *Freq G + C* | *Total CpG* | *Ratio O/E CpG* | *Total UpA* | *Ratio O/E UpA* | *CAI[a] Hs* | *CAI[b] Dm* |
|---|---|---|---|---|---|---|---|---|
| *Wildtype* | 10727 | 0.51 | 308 | 0.45 | 342 | 0.54 | 0.71 | 0.60 |
| *Wildtype* | 1654 | 0.53 | 55 | 0.49 | 57 | 0.64 | 0.77 | 0.64 |
| *SCR* | 1654 | 0.53 | 55 | 0.49 | 49 | 0.55 | 0.76 | 0.65 |
| *CpG_1.0* | 1654 | 0.46 | 114 | 1.01 | 57 | 0.64 | 0.70 | 0.63 |
| *CpG_max* | 1654 | 0.61 | 245 | 1.62 | 57 | 0.93 | 0.67 | 0.71 |
| *UpA_max* | 1654 | 0.53 | 55 | 0.68 | 207 | 1.71 | 0.63 | 0.51 |

[a] Codon adaptation index (CAI) calculated against the codon usage table of Homo sapiens (Hs)

[b] CAI calculated against the codon usage table of Drosophila melanogaster (Dm)
