## Supplement file 1 for "An adaptive compromise - Conflicting evolutionary pressures on arthropod-borne Zika virus dinucleotide composition in mammalian hosts and mosquito vectors"

| *H. Sapiens* | *Ae. Aegypti* | flaviviruses | ZIKV isolates |
|---|---|---|---|
| NM_000043.6 | XM_021837896.1 | DQ235145 | KU365778 |
| NM_000061.3 | XM_021838519.1 | AY323490 | KU497555.1 |
| NM_000068.4 | XM_021838576.1 | AF331718 | KU955595.1 |
| NM_000101.4 | XM_021856355.1 | AF253419 | KU955593.1 |
| NM_000116.5 | XM_021837651.1 | NC_001809 | KX247646.1 |
| NM_000117.3 | XR_002502975.1 | DQ235152 | KU744693.1 |
| NM_000148.4 | XM_021848511.1 | DQ235151 | MK241417.1 |
| NM_000151.4 | XR_002502976.1 | DQ235153 | MN100039.1 |
| NM_000184.3 | XR_002501108.1 | AY323489 | KX117076.1 |
| NM_000193.4 | XR_002500046.1 | PWARPT | MH513600.1 |
| NM_000195.5 | XR_002499437.1 | DQ235149 | KY785466.1 |
| NM_000221.3 | XR_002500378.1 | TEU27495 | KY014320.2 |
| NM_000233.4 | XM_021839360.1 | KU761576 | MK028857.1 |
| NM_000244.3 | XM_021855092.1 | L40361 | KU922960.1 |
| NM_000285.4 | XR_002499655.1 | DQ235144 | KY014319.2 |
| NM_000289.6 | XM_001660351.2 | DQ235150 | KY014300.2 |
| NM_000325.6 | XM_001653904.2 | DQ235148 | KY325476.1 |
| NM_000334.4 | XR_002499247.1 | DQ235146 | KX830930.1 |
| NM_000355.4 | XM_021843798.1 | KF917535 | KU527068 |
| NM_000360.4 | XM_021841335.1 | AY632536 | |
| NM_000380.4 | XM_021839798.1 | AY632538 | |
| NM_000420.3 | XR_002500965.1 | KF917538 | |
| NM_000442.5 | XM_001649659.2 | DVU88536 | |
| NM_000455.5 | XM_001653108.2 | DENRCG | |
| NM_000463.3 | XR_002498938.1 | DENCME | |
| NM_000486.6 | XM_001651609.2 | AF326573 | |
| NM_000487.6 | XM_001652798.2 | LN849009 | |
| NM_000514.4 | XM_001649658.2 | JEVCG | |
| NM_000531.6 | XR_002500826.1 | EU082200 | |
| NM_000603.5 | XR_002500705.1 | AY898809 | |
| NM_000661.5 | XR_002500569.1 | AF161266 | |
| NM_000666.3 | XR_002501273.1 | DQ525916 | |
| NM_000679.4 | XR_002500752.1 | AY453411 | |
| NM_000700.3 | XR_002499843.1 | KUNCG | |
| NM_000705.4 | XM_021853272.1 | WNFCG | |
| NM_000717.5 | XM_021847017.1 | EU082199 | |
| NM_000723.5 | XM_021839613.1 | NC_009029 | |
| NM_000742.4 | XM_021846941.1 | MF380434 | |
| NM_000748.3 | XR_002500629.1 | NC_009028 | |
| NM_000784.4 | XM_001661645.2 | MF461639 | |
| NM_000794.5 | XR_002501303.1 | KC734552 | |
| NM_000808.4 | XM_021857092.1 | NC_018705 | |
| NM_000809.4 | XM_001650132.2 | KF192951 | |
| NM_000842.5 | XR_002499034.1 | DQ859059 | |
| NM_000845.3 | XR_002500400.1 | DQ859063 | |
| NM_000848.4 | XM_001659562.2 | DQ859058 | |
| NM_000862.3 | XM_021844096.1 | X03700 | |
| NM_000869.6 | XM_021854409.1 | NC_008718 | |
| NM_000882.4 | XM_021839364.1 | NC_026624 | |

| | | |
|---|---|---|
| NM_000896.3 | XM_021857090.1 | NC_005039 |
| NM_000911.4 | XM_021847667.1 | AF160193 |
| NM_000932.5 | XM_021856647.1 | NC_026620 |
| NM_000936.4 | XM_021839537.1 | AJ242984 |
| NM_000952.5 | XM_021850851.1 | AJ299445 |
| NM_000954.6 | XM_001656983.2 | KJ469370 |
| NM_000961.4 | XM_021856340.1 | NC_034007 |
| NM_001001.5 | XM_021837967.1 | AF144692 |
| NM_001001349.2 | XM_021856059.1 | NC_005064 |
| NM_001001437.4 | XR_002499502.1 | JQ268258 |
| NM_001001479.4 | XR_002499293.1 | NC_024299 |
| NM_001001547.3 | XM_021850832.1 | NC_027817 |
| NM_001001563.5 | XR_002502609.1 | NC_001564 |
| NM_001001580.2 | XM_021856422.1 | GQ165809 |
| NM_001001694.3 | XR_002499302.1 | NC_012932 |
| NM_001001715.4 | XM_021850829.1 | KC505248 |
| NM_001001722.2 | XR_002499606.1 | NC_012671 |
| NM_001001913.2 | XM_021842531.1 | HE574574 |
| NM_001001918.1 | XR_002502037.1 | NC_008604 |
| NM_001001952.1 | XR_002499177.1 | KX669689 |
| NM_001001957.2 | XM_021839538.1 | DQ400858 |
| NM_001001958.1 | XR_002503231.1 | NC_033694 |
| NM_001001965.1 | XM_001649656.2 | NC_024017 |
| NM_001001991.3 | XM_021856243.1 | MF139576 |
| NM_001002006.3 | XR_002500221.1 | NC_017086 |
| NM_001002249.2 | XR_002500915.1 | NC_016997 |
| NM_001002251.3 | XM_021849176.1 | KC692068 |
| NM_001002261.4 | XM_001652840.2 | KC496020 |
| NM_001002262.4 | XM_021842437.1 | MF139575 |
| NM_001002294.3 | XR_002500263.1 | NC_024805 |
| NM_001002814.3 | XM_001653251.2 | EU159426 |
| NM_001002879.1 | XR_002502942.1 | MG587038 |
| NM_001002914.3 | XR_002499522.1 | MH158418 |
| NM_001003443.3 | XM_021838320.1 | MH158417 |
| NM_001003745.2 | XR_002502671.1 | MH158416 |
| NM_001003760.5 | XM_021842955.1 | MH158415 |
| NM_001003789.3 | XR_002500597.1 | |
| NM_001003845.3 | XM_001655319.2 | |
| NM_001003897.2 | XR_002498878.1 | |
| NM_001003937.3 | XR_002501094.1 | |
| NM_001003941.3 | XM_001649647.2 | |
| NM_001004.4 | XR_002500515.1 | |
| NM_001004056.2 | XR_002502072.1 | |
| NM_001004135.2 | XM_021843923.1 | |
| NM_001004298.4 | XR_002500009.1 | |
| NM_001004416.3 | XM_021848799.1 | |
| NM_001004463.2 | XR_002500395.1 | |
| NM_001004705.2 | XR_002499348.1 | |
| NM_001004723.3 | XR_002500819.1 | |
| NM_001004732.2 | XR_002500043.1 | |

| | |
|---|---|
| NM_001004738.2 | XM_021854178.1 |
| NM_001004750.1 | XR_002502353.1 |
| NM_001004752.2 | XR_002502492.1 |
| NM_001004757.2 | XM_021839628.1 |
| NM_001005160.3 | XR_002500971.1 |
| NM_001005163.2 | XM_021843332.1 |
| NM_001005165.2 | XM_001649672.2 |
| NM_001005166.5 | XM_021844475.1 |
| NM_001005172.2 | XM_001653220.2 |
| NM_001005198.2 | XM_001656328.2 |
| NM_001005241.4 | XM_021842532.1 |
| NM_001005279.3 | XR_002503024.1 |
| NM_001005284.1 | XM_021842460.1 |
| NM_001005289.5 | XR_002499049.1 |
| NM_001005369.1 | XM_021841437.1 |
| NM_001005404.4 | XM_021844264.1 |
| NM_001005416.2 | XM_021851628.1 |
| NM_001005466.2 | XR_002500997.1 |
| NM_001005473.3 | XM_021842795.1 |
| NM_001005489.2 | XR_002499861.1 |
| NM_001005527.3 | XR_002502233.1 |
| NM_001005781.2 | XM_021852829.1 |
| NM_001005853.1 | XM_001661458.2 |
| NM_001006109.1 | XM_021855606.1 |
| NM_001006946.2 | XM_001656229.2 |
| NM_001007025.2 | XM_021850900.1 |
| NM_001007089.4 | XM_021844310.1 |
| NM_001007226.1 | XR_002499834.1 |
| NM_001007234.3 | XR_002498801.1 |
| NM_001007464.3 | XR_002500163.1 |
| NM_001007530.3 | XR_002502932.1 |
| NM_001007537.2 | XR_002502345.1 |
| NM_001007559.3 | XM_001660308.2 |
| NM_001008215.3 | XM_001658818.2 |
| NM_001008273.2 | XM_001649665.2 |
| NM_001008387.3 | XM_021852850.1 |
| NM_001008740.4 | XM_021843268.1 |
| NM_001008781.3 | XM_001662067.2 |
| NM_001009566.3 | XR_002498673.1 |
| NM_001009584.2 | XM_021846071.1 |
| NM_001009614.3 | XM_001647908.2 |
| NM_001009992.1 | XM_001652936.2 |
| NM_001010000.3 | XR_002501908.1 |
| NM_001010844.4 | XM_021844314.1 |
| NM_001010866.4 | XM_001661544.2 |
| NM_001010877.5 | XM_021838751.1 |
| NM_001011.4 | XM_021850874.1 |
| NM_001011658.4 | XM_001659574.2 |
| NM_001011709.3 | XM_021857311.1 |
| NM_001012455.2 | XM_021848696.1 |

| | |
|---|---|
| NM_001012727.2 | XM_021851496.1 |
| NM_001012763.2 | XM_021844347.1 |
| NM_001013253.2 | XR_002498816.1 |
| NM_001013438.3 | XR_002502089.1 |
| NM_001013630.2 | XM_011494961.2 |
| NM_001013690.5 | XM_021857312.1 |
| NM_001013699.3 | XM_001659603.2 |
| NM_001013837.2 | XM_021842468.1 |
| NM_001014283.2 | XM_021853495.1 |
| NM_001014336.2 | XM_001658477.2 |
| NM_001014342.3 | XM_021852378.1 |
| NM_001014435.2 | XR_002501980.1 |
| NM_001014447.3 | XM_021837982.1 |
| NM_001014999.3 | XM_021850754.1 |
| NM_001015.5 | XM_021849319.1 |
| NM_001015049.3 | XR_002500809.1 |
| NM_001015050.3 | XM_021848995.1 |
| NM_001015072.4 | XM_021856296.1 |
| NM_001017395.5 | XM_001651099.2 |
| NM_001017416.2 | XM_021844340.1 |
| NM_001017926.3 | XM_021844346.1 |
| NM_001017970.3 | XR_002500570.1 |
| NM_001018100.5 | XM_021847572.1 |
| NM_001018136.3 | XR_002501642.1 |
| NM_001018160.2 | XM_021842896.1 |
| NM_001018161.2 | XM_021853485.1 |
| NM_001019.5 | XM_021850768.1 |
| NM_001020821.2 | XM_021844306.1 |
| NM_001024074.3 | XM_001651095.2 |
| NM_001024381.2 | XM_021855457.1 |
| NM_001024383.2 | XR_002499044.1 |
| NM_001024593.2 | XM_021842264.1 |
| NM_001024601.3 | XM_021841334.1 |
| NM_001024628.3 | XM_021844313.1 |
| NM_001024662.3 | XR_002503129.1 |
| NM_001024855.3 | XR_002503136.1 |
| NM_001024924.2 | XM_021844246.1 |
| NM_001025076.2 | XR_002500336.1 |
| NM_001025081.2 | XM_021853642.1 |
| NM_001025107.3 | XM_021839614.1 |
| NM_001025160.3 | XM_021850164.1 |
| NM_001025238.2 | XR_002502762.1 |
| NM_001029875.3 | XM_001659747.3 |
| NM_001029891.3 | XR_002499399.1 |
| NM_001029955.4 | XM_021839505.1 |
| NM_001031698.3 | XM_021841432.1 |
| NM_001031713.3 | XR_002501504.1 |
| NM_001031735.3 | XM_021849007.1 |
| NM_001031835.3 | XM_021857462.1 |
| NM_001032.5 | XM_001657977.2 |

| | |
|---|---|
| NM_001033117.3 | XM_001660597.2 |
| NM_001033602.4 | XM_001654891.2 |
| NM_001033855.3 | XR_002502700.1 |
| NM_001034172.4 | XM_021837394.1 |
| NM_001034957.3 | XM_001651695.2 |
| NM_001035003.2 | XR_002498611.1 |
| NM_001037132.4 | XM_021846196.1 |
| NM_001037165.2 | XR_002499975.1 |
| NM_001037232.4 | XR_002500384.1 |
| NM_001037330.3 | XM_021842231.1 |
| NM_001037331.3 | XM_021844343.1 |
| NM_001037442.4 | XR_002500401.1 |
| NM_001037499.2 | XR_002499402.1 |
| NM_001037553.2 | XM_021844365.1 |
| NM_001037582.3 | XR_002498598.1 |
| NM_001037671.4 | XM_001654389.2 |
| NM_001038707.2 | XM_021847927.1 |
| NM_001039130.2 | XR_002501977.1 |
| NM_001039211.3 | XM_021839673.1 |
| NM_001039213.4 | XR_002500499.1 |
| NM_001039348.3 | XM_021854226.1 |
| NM_001039374.5 | XR_002500473.1 |
| NM_001039656.1 | XM_021855298.1 |
| NM_001039664.2 | XR_002499666.1 |
| NM_001039780.4 | XM_001662134.2 |
| NM_001040025.3 | XR_002501421.1 |
| NM_001040105.2 | XR_002500397.1 |
| NM_001040107.2 | XR_002499170.1 |
| NM_001040110.2 | XM_001659590.2 |
| NM_001040126.2 | XM_021848608.1 |
| NM_001040272.6 | XR_002498935.1 |
| NM_001040284.3 | XR_002502283.1 |
| NM_001040443.3 | XM_021837869.1 |
| NM_001040647.2 | XM_021839369.1 |
| NM_001042351.3 | XM_021844624.1 |
| NM_001042361.5 | XM_001649652.2 |
| NM_001042370.2 | XR_002501342.1 |
| NM_001042422.3 | XR_002499384.1 |
| NM_001042423.3 | XM_001651978.2 |
| NM_001042428.2 | XM_021856855.1 |
| NM_001042442.3 | XM_001661646.2 |
| NM_001042450.4 | XR_002499112.1 |
| NM_001042459.3 | XR_002499931.1 |
| NM_001042469.3 | XM_021844219.1 |
| NM_001042574.3 | XR_002501535.1 |
| NM_001042616.3 | XR_002498794.1 |
| NM_001042705.3 | XM_021844363.1 |
| NM_001043353.2 | XM_021848994.1 |
| NM_001045477.4 | XR_002498918.1 |
| NM_001063.4 | XM_001660307.2 |

| | |
|---|---|
| NM_001076683.2 | XM_021843025.1 |
| NM_001077.4 | XM_021848999.1 |
| NM_001077188.2 | XM_021851716.1 |
| NM_001077194.2 | XR_002498936.1 |
| NM_001077471.3 | XM_021839357.1 |
| NM_001077477.3 | XR_002502028.1 |
| NM_001077480.3 | XM_001655174.2 |
| NM_001077526.3 | XM_021844469.1 |
| NM_001077619.2 | XM_021844368.1 |
| NM_001077653.2 | XM_001652515.2 |
| NM_001077664.3 | XM_021837887.1 |
| NM_001078170.2 | XR_002501641.1 |
| NM_001079817.3 | XM_001648290.2 |
| NM_001079882.2 | XM_001652439.1 |
| NM_001080121.3 | XR_002501234.1 |
| NM_001080446.3 | XM_021854213.1 |
| NM_001080482.4 | XR_002499813.1 |
| NM_001080503.3 | XM_001650651.2 |
| NM_001080534.3 | XR_002502882.1 |
| NM_001080744.2 | XM_001650627.2 |
| NM_001080998.2 | XM_001658763.2 |
| NM_001081492.2 | XM_021848027.1 |
| NM_001081573.3 | XM_001660306.2 |
| NM_001081675.3 | XM_021844356.1 |
| NM_001083592.2 | XM_021851489.1 |
| NM_001083600.3 | XM_021856403.1 |
| NM_001083956.2 | XM_021844736.1 |
| NM_001085426.3 | XM_021850730.1 |
| NM_001093729.2 | XM_021853349.1 |
| NM_001097605.2 | XR_002500998.1 |
| NM_001098212.2 | XR_002501864.1 |
| NM_001098405.2 | XM_021837888.1 |
| NM_001098412.2 | XM_021839564.1 |
| NM_001098418.2 | XM_021844309.1 |
| NM_001098504.2 | XM_021843047.1 |
| NM_001098519.2 | XM_021853610.1 |
| NM_001098525.3 | XR_002501503.1 |
| NM_001098535.1 | XR_002502990.1 |
| NM_001098670.2 | XR_002501749.1 |
| NM_001098818.4 | XR_002501620.1 |
| NM_001099403.2 | XR_002501587.1 |
| NM_001099650.2 | XM_001660303.2 |
| NM_001099669.2 | XM_001655670.2 |
| NM_001099670.3 | XM_021856701.1 |
| NM_001099681.2 | XR_002498829.1 |
| NM_001099733.2 | XR_002499945.1 |
| NM_001099735.2 | XM_001654902.2 |
| NM_001099746.2 | XM_011494663.2 |
| NM_001099783.2 | XM_021853577.1 |
| NM_001099857.5 | XM_021837745.1 |

| | |
|---|---|
| NM_001100407.3 | XM_001659051.2 |
| NM_001100411.3 | XM_021844233.1 |
| NM_001100620.3 | XM_001649537.2 |
| NM_001100621.3 | XR_002501422.1 |
| NM_001100625.3 | XM_021844241.1 |
| NM_001101311.2 | XM_001661485.2 |
| NM_001101362.3 | XM_021838675.1 |
| NM_001102470.2 | XM_001660722.2 |
| NM_001102559.2 | XM_001654006.2 |
| NM_001102562.3 | XM_021851408.1 |
| NM_001105.5 | XR_002500135.1 |
| NM_001105207.3 | XR_002499366.1 |
| NM_001105513.3 | XM_021847919.1 |
| NM_001105518.2 | XM_021844852.1 |
| NM_001105558.1 | XR_002500772.1 |
| NM_001105563.2 | XM_001654652.2 |
| NM_001105569.3 | XM_001655668.2 |
| NM_001105573.2 | XR_002501293.1 |
| NM_001109754.4 | XM_021845365.1 |
| NM_001110354.2 | XM_021855447.1 |
| NM_001111035.3 | XR_002498847.1 |
| NM_001111307.2 | XM_001661984.2 |
| NM_001111309.1 | XM_001648207.2 |
| NM_001113324.3 | XM_021845366.1 |
| NM_001113348.2 | XR_002502445.1 |
| NM_001113511.2 | XM_021851409.1 |
| NM_001113525.2 | XM_021855190.1 |
| NM_001113535.2 | XM_011494756.2 |
| NM_001113546.2 | XM_021837396.1 |
| NM_001114086.2 | XM_021839356.1 |
| NM_001114092.2 | XR_002499894.1 |
| NM_001114182.3 | XM_021839954.1 |
| NM_001114377.2 | XM_021849169.1 |
| NM_001114759.3 | XM_021853335.1 |
| NM_001117.5 | XM_001660301.2 |
| NM_001122630.2 | XR_002501618.1 |
| NM_001122679.2 | XM_021841219.1 |
| NM_001122825.2 | XR_002502799.1 |
| NM_001122847.3 | XR_002500899.1 |
| NM_001122853.3 | XM_021856183.1 |
| NM_001122898.3 | XM_021849333.1 |
| NM_001123366.2 | XM_021849097.1 |
| NM_001123377.2 | XM_021851594.1 |
| NM_001126129.2 | XM_021855455.1 |
| NM_001126339.3 | XM_001654900.2 |
| NM_001127223.1 | XM_001662456.2 |
| NM_001127255.1 | XR_002499500.1 |
| NM_001127361.3 | XM_021855286.1 |
| NM_001127362.2 | XM_021857197.1 |
| NM_001127392.3 | XM_001660623.2 |

| | |
|---|---|
| NM_001127493.3 | XR_002501976.1 |
| NM_001127585.2 | XM_001663928.2 |
| NM_001127644.2 | XM_001656568.2 |
| NM_001127670.4 | XM_021844167.1 |
| NM_001127698.2 | XM_001651203.2 |
| NM_001127700.2 | XM_001657679.2 |
| NM_001127895.2 | XM_021856850.1 |
| NM_001128310.3 | XM_021851424.1 |
| NM_001128612.2 | XM_001652656.2 |
| NM_001128631.3 | XM_001649339.2 |
| NM_001128633.2 | XR_002503107.1 |
| NM_001128635.1 | XM_021844753.1 |
| NM_001128847.4 | XM_021856122.1 |
| NM_001128852.2 | XR_002502691.1 |
| NM_001128932.2 | XM_021849171.1 |
| NM_001129729.3 | XM_021850780.1 |
| NM_001129830.3 | XR_002501746.1 |
| NM_001129834.2 | XR_002502713.1 |
| NM_001129837.2 | XR_002500879.1 |
| NM_001130066.2 | XM_001663845.2 |
| NM_001130087.2 | XM_021843046.1 |
| NM_001130091.2 | XM_001654004.2 |
| NM_001130410.2 | XM_021837649.1 |
| NM_001130455.2 | XM_021855580.1 |
| NM_001130514.3 | XM_001650449.2 |
| NM_001130831.2 | XM_021850537.1 |
| NM_001130861.1 | XM_021838725.1 |
| NM_001130864.2 | XM_021843888.1 |
| NM_001130957.2 | XM_021847795.1 |
| NM_001130964.2 | XM_021847666.1 |
| NM_001131009.2 | XM_011494845.2 |
| NM_001134296.2 | XR_002502881.1 |
| NM_001134373.3 | XR_002502471.1 |
| NM_001134432.2 | XR_002499370.1 |
| NM_001134451.2 | XR_002499583.1 |
| NM_001134649.3 | XR_002501238.1 |
| NM_001134672.2 | XR_002499730.1 |
| NM_001134673.4 | XM_021840542.1 |
| NM_001134759.2 | XM_021839373.1 |
| NM_001134771.2 | XR_002502651.1 |
| NM_001134774.2 | XM_021851738.1 |
| NM_001135054.2 | XM_021855291.1 |
| NM_001135191.2 | XM_001655010.2 |
| NM_001135599.4 | XM_021844133.1 |
| NM_001135664.2 | XM_001650203.2 |
| NM_001135669.2 | XM_021842667.1 |
| NM_001135671.3 | XM_021852016.1 |
| NM_001135675.2 | XM_021839525.1 |
| NM_001135697.3 | XM_001653180.2 |
| NM_001135701.2 | XM_021851066.1 |

| | |
|---|---|
| NM_001135707.3 | XM_021843082.1 |
| NM_001135955.3 | XR_002500073.1 |
| NM_001135998.3 | XM_021848386.1 |
| NM_001135999.1 | XM_021842130.1 |
| NM_001136020.3 | XM_021854203.1 |
| NM_001136109.3 | XM_001650025.2 |
| NM_001136130.3 | XM_021855773.1 |
| NM_001136214.3 | XM_001659702.2 |
| NM_001136263.2 | XM_001655178.2 |
| NM_001136493.3 | XR_002500273.1 |
| NM_001136497.3 | XM_021844159.1 |
| NM_001136534.3 | XM_021845707.1 |
| NM_001136558.2 | XM_021840544.1 |
| NM_001136575.2 | XM_021853620.1 |
| NM_001137675.4 | XM_021837676.1 |
| NM_001139459.2 | XM_021852244.1 |
| NM_001139518.1 | XR_002500600.1 |
| NM_001142310.2 | XR_002498881.1 |
| NM_001142370.2 | XM_021842137.1 |
| NM_001142434.2 | XM_001651303.2 |
| NM_001142463.3 | XM_001660714.2 |
| NM_001142473.2 | XM_021851011.1 |
| NM_001142536.2 | XM_021840603.1 |
| NM_001142540.2 | XM_001660213.2 |
| NM_001142545.2 | XM_011494649.2 |
| NM_001142550.2 | XM_021856473.1 |
| NM_001142576.2 | XR_002499485.1 |
| NM_001142625.2 | XM_021855770.1 |
| NM_001142643.3 | XM_021847664.1 |
| NM_001142673.3 | XM_021856987.1 |
| NM_001142678.2 | XM_021853575.1 |
| NM_001142799.3 | XM_021851607.1 |
| NM_001142800.2 | XM_021845066.1 |
| NM_001142935.2 | XM_021840541.1 |
| NM_001142961.1 | XM_001654459.3 |
| NM_001143773.1 | XR_002500728.1 |
| NM_001143788.2 | XR_002498927.1 |
| NM_001143812.2 | XR_002501772.1 |
| NM_001143823.3 | XM_021855836.1 |
| NM_001143942.2 | XM_001655657.2 |
| NM_001144012.3 | XR_002502915.1 |
| NM_001144060.2 | XR_002500851.1 |
| NM_001144831.2 | XM_001660202.2 |
| NM_001144875.2 | XR_002498767.1 |
| NM_001144879.2 | XM_001660604.2 |
| NM_001144903.3 | XM_021852964.1 |
| NM_001144916.2 | XM_021838257.1 |
| NM_001144920.3 | XM_001655303.2 |
| NM_001144924.2 | XM_021848578.1 |
| NM_001144984.3 | XR_002499555.1 |

| | |
|---|---|
| NM_001145059.2 | XR_002499734.1 |
| NM_001145102.2 | XM_021839876.1 |
| NM_001145104.2 | XM_021844095.1 |
| NM_001145118.1 | XR_002501686.1 |
| NM_001145124.1 | XM_021854208.1 |
| NM_001145144.2 | XM_021842582.1 |
| NM_001145146.2 | XM_021839543.1 |
| NM_001145345.1 | XM_001663374.2 |
| NM_001145445.1 | XM_021841635.1 |
| NM_001145475.3 | XM_001651495.2 |
| NM_001145548.2 | XM_001663015.2 |
| NM_001145640.2 | XR_002501458.1 |
| NM_001145650.2 | XM_001658827.2 |
| NM_001145659.1 | XR_002501262.1 |
| NM_001145668.2 | XM_001657903.2 |
| NM_001145713.2 | XM_001660414.2 |
| NM_001145717.1 | XR_002499433.1 |
| NM_001145734.2 | XM_021855626.1 |
| NM_001145829.2 | XM_021846359.1 |
| NM_001145928.2 | XR_002500618.1 |
| NM_001146055.2 | XM_001656485.3 |
| NM_001146158.2 | XR_002501440.1 |
| NM_001146171.2 | XM_021847573.1 |
| NM_001146191.2 | XR_002502754.1 |
| NM_001146333.3 | XM_021848953.1 |
| NM_001146339.2 | XM_021849088.1 |
| NM_001146683.2 | XR_002500632.1 |
| NM_001146686.3 | XR_002503020.1 |
| NM_001148.6 | XM_021845608.1 |
| NM_001159286.2 | XR_002502650.1 |
| NM_001159322.2 | XM_021855628.1 |
| NM_001159379.2 | XM_021839670.1 |
| NM_001159554.1 | XM_021850915.1 |
| NM_001159601.2 | XM_021847038.1 |
| NM_001159740.2 | XM_001660730.2 |
| NM_001159773.2 | XR_002503181.1 |
| NM_001160002.2 | XM_011495066.2 |
| NM_001160116.2 | XM_021849087.1 |
| NM_001160125.2 | XM_001657764.2 |
| NM_001160166.2 | XR_002499642.1 |
| NM_001160176.4 | XM_021852576.1 |
| NM_001160224.2 | XM_001659180.1 |
| NM_001160244.2 | XR_002502949.1 |
| NM_001160300.2 | XM_021846536.1 |
| NM_001160302.1 | XM_021849405.1 |
| NM_001161330.2 | XM_021854866.1 |
| NM_001161368.3 | XM_021852995.1 |
| NM_001161454.1 | XM_001652365.2 |
| NM_001161527.2 | XM_021849170.1 |
| NM_001161562.3 | XM_001649750.2 |

| | |
|---|---|
| NM_001161584.2 | XM_021845159.1 |
| NM_001161662.2 | XM_021856425.1 |
| NM_001162407.1 | XM_021853208.1 |
| NM_001162862.2 | XM_001652313.2 |
| NM_001162936.2 | XR_002498855.1 |
| NM_001163335.2 | XM_021841531.1 |
| NM_001163484.2 | XM_001661348.3 |
| NM_001163560.3 | XM_021845131.1 |
| NM_001163735.2 | XM_021856683.1 |
| NM_001164161.2 | XM_021844171.1 |
| NM_001164283.3 | XM_001661007.2 |
| NM_001164319.2 | XM_021857597.1 |
| NM_001164381.2 | XM_021848580.1 |
| NM_001164399.2 | XM_021843277.1 |
| NM_001164442.2 | XM_001661784.2 |
| NM_001164538.2 | XR_002501499.1 |
| NM_001164542.2 | XR_002503273.1 |
| NM_001164619.2 | XR_002503071.1 |
| NM_001164638.3 | XM_021843979.1 |
| NM_001164690.2 | XM_021837968.1 |
| NM_001164694.2 | XR_002502498.1 |
| NM_001164738.1 | XM_021848396.1 |
| NM_001164752.2 | XR_002499278.1 |
| NM_001164778.2 | XR_002498804.1 |
| NM_001164781.2 | XM_021849025.1 |
| NM_001164831.3 | XM_021844116.1 |
| NM_001164835.2 | XM_021852697.1 |
| NM_001165255.2 | XM_021856590.1 |
| NM_001165412.2 | XM_021837995.1 |
| NM_001165885.2 | XM_021847325.1 |
| NM_001165928.3 | XM_001663026.2 |
| NM_001165945.2 | XM_001649671.2 |
| NM_001166208.2 | XM_011495224.2 |
| NM_001166280.2 | XM_021843763.1 |
| NM_001166287.2 | XM_001648972.2 |
| NM_001166387.4 | XM_021853336.1 |
| NM_001166393.2 | XM_021850949.1 |
| NM_001166533.2 | XM_021839541.1 |
| NM_001166703.2 | XM_001661166.2 |
| NM_001166704.2 | XM_021850960.1 |
| NM_001167605.2 | XR_002502270.1 |
| NM_001167608.3 | XM_001663498.2 |
| NM_001167890.2 | XM_021841326.1 |
| NM_001167915.3 | XR_002500373.1 |
| NM_001168323.2 | XM_001652071.2 |
| NM_001168390.2 | XM_021850955.1 |
| NM_001170330.1 | XR_002503092.1 |
| NM_001170423.2 | XM_001659755.2 |
| NM_001170670.2 | XM_021844190.1 |
| NM_001170687.4 | XM_021843079.1 |

| | |
|---|---|
| NM_001170741.3 | XM_021844324.1 |
| NM_001170760.2 | XM_001648225.2 |
| NM_001170881.2 | XM_021857502.1 |
| NM_001171025.2 | XM_001650968.2 |
| NM_001171653.2 | XR_002501970.1 |
| NM_001171743.3 | XR_002502948.1 |
| NM_001171934.1 | XR_002501813.1 |
| NM_001172568.2 | XM_021850749.1 |
| NM_001172640.2 | XM_021851897.1 |
| NM_001172671.2 | XM_021850115.1 |
| NM_001172771.2 | XM_021845376.1 |
| NM_001172772.2 | XM_021855629.1 |
| NM_001172815.3 | XM_001651216.2 |
| NM_001172818.1 | XM_001658820.2 |
| NM_001173129.2 | XM_021847574.1 |
| NM_001174070.3 | XM_021848671.1 |
| NM_001174085.2 | XM_021854864.1 |
| NM_001174104.2 | XM_001661747.2 |
| NM_001174105.2 | XR_002502454.1 |
| NM_001174127.2 | XM_021842675.1 |
| NM_001174166.2 | XR_002500097.1 |
| NM_001177310.2 | XM_021844712.1 |
| NM_001177704.3 | XM_001664182.2 |
| NM_001177800.2 | XR_002500196.1 |
| NM_001177801.2 | XR_002503135.1 |
| NM_001177998.2 | XM_001656650.2 |
| NM_001178030.2 | XM_021857081.1 |
| NM_001178094.2 | XM_021849163.1 |
| NM_001178106.1 | XR_002503255.1 |
| NM_001178140.2 | XR_002500520.1 |
| NM_001184693.2 | XM_001655691.2 |
| NM_001184696.1 | XR_002502340.1 |
| NM_001184721.2 | XM_001655646.2 |
| NM_001184751.2 | XM_001661338.2 |
| NM_001184760.2 | XR_002500561.1 |
| NM_001184792.2 | XR_002501955.1 |
| NM_001184866.2 | XM_021853605.1 |
| NM_001184872.2 | XM_021843602.1 |
| NM_001184903.1 | XM_001662189.3 |
| NM_001184970.3 | XM_001660967.2 |
| NM_001184974.2 | XM_021857066.1 |
| NM_001184993.2 | XM_001658441.2 |
| NM_001185054.2 | XM_021850172.1 |
| NM_001185119.1 | XM_021844521.1 |
| NM_001185149.1 | XR_002500902.1 |
| NM_001190158.1 | XR_002500194.1 |
| NM_001190447.2 | XM_021839739.1 |
| NM_001190462.2 | XM_021846542.1 |
| NM_001190482.2 | XM_021848401.1 |
| NM_001190789.2 | XM_001663679.2 |

| | |
|---|---|
| NM_001190791.2 | XM_021855299.1 |
| NM_001190801.2 | XM_021843729.1 |
| NM_001190811.2 | XM_021856205.1 |
| NM_001190848.2 | XM_021857546.1 |
| NM_001190882.3 | XM_001651206.2 |
| NM_001190983.2 | XM_021839639.1 |
| NM_001191029.2 | XR_002503257.1 |
| NM_001191037.2 | XM_001657417.2 |
| NM_001191059.4 | XR_002501586.1 |
| NM_001193272.2 | XR_002503021.1 |
| NM_001193360.2 | XM_021840501.1 |
| NM_001193461.2 | XM_021856559.1 |
| NM_001193487.3 | XM_021845705.1 |
| NM_001193503.2 | XM_021840487.1 |
| NM_001193557.2 | XM_001656559.2 |
| NM_001194958.2 | XR_002501801.1 |
| NM_001195035.2 | XM_021846137.1 |
| NM_001195076.2 | XR_002501597.1 |
| NM_001195104.3 | XM_021856937.1 |
| NM_001195203.4 | XM_021854860.1 |
| NM_001195249.2 | XM_021844090.1 |
| NM_001195296.2 | XM_021847596.1 |
| NM_001195303.3 | XR_002501581.1 |
| NM_001195307.2 | XM_021852226.1 |
| NM_001195578.2 | XR_002498810.1 |
| NM_001195803.2 | XM_021838741.1 |
| NM_001198541.3 | XM_021845943.1 |
| NM_001198621.4 | XR_002501889.1 |
| NM_001198689.2 | XM_021842372.1 |
| NM_001198756.1 | XM_001654990.2 |
| NM_001198855.1 | XM_001661988.2 |
| NM_001198898.2 | XM_021843294.1 |
| NM_001198909.2 | XM_001648723.2 |
| NM_001199041.2 | XM_021837694.1 |
| NM_001199042.2 | XM_001662155.2 |
| NM_001199081.3 | XM_001660041.2 |
| NM_001199135.3 | XM_021837678.1 |
| NM_001199148.1 | XM_021840612.1 |
| NM_001199255.3 | XM_001660277.2 |
| NM_001199262.3 | XR_002500072.1 |
| NM_001199379.2 | XM_021845792.1 |
| NM_001199388.3 | XM_001657239.3 |
| NM_001199427.2 | XR_002503133.1 |
| NM_001199463.2 | XM_021850453.1 |
| NM_001199570.2 | XM_011495347.2 |
| NM_001199636.2 | XM_021852727.1 |
| NM_001199637.2 | XM_021843291.1 |
| NM_001199642.1 | XM_021845023.1 |
| NM_001199661.1 | XR_002500803.1 |
| NM_001199708.2 | XM_001660760.2 |

| | |
|---|---|
| NM_001199752.3 | XR_002498798.1 |
| NM_001199771.3 | XM_021852381.1 |
| NM_001199805.1 | XM_021846144.1 |
| NM_001199837.3 | XM_021847389.1 |
| NM_001199851.1 | XM_001653669.2 |
| NM_001199859.3 | XM_021843293.1 |
| NM_001199876.1 | XM_001657906.2 |
| NM_001199987.2 | XM_001655635.2 |
| NM_001200051.2 | XR_002501799.1 |
| NM_001201365.2 | XM_021848395.1 |
| NM_001201473.2 | XR_002501064.1 |
| NM_001201480.2 | XM_001663652.2 |
| NM_001202409.2 | XM_021856348.1 |
| NM_001202410.2 | XM_021857637.1 |
| NM_001202498.2 | XM_021845008.1 |
| NM_001202555.2 | XM_001657908.2 |
| NM_001203261.2 | XM_021843286.1 |
| NM_001204066.2 | XM_001658890.2 |
| NM_001204078.2 | XM_021853959.1 |
| NM_001204145.3 | XR_002498591.1 |
| NM_001204148.3 | XM_021845693.1 |
| NM_001204175.2 | XM_021848676.1 |
| NM_001204190.2 | XM_001650008.2 |
| NM_001204212.2 | XM_021840237.1 |
| NM_001204221.2 | XM_021844087.1 |
| NM_001204290.2 | XM_021847088.1 |
| NM_001204293.2 | XM_021847395.1 |
| NM_001204408.2 | XM_001654903.2 |
| NM_001204505.3 | XM_021847672.1 |
| NM_001204514.2 | XM_001661499.2 |
| NM_001204698.2 | XM_021840503.1 |
| NM_001204803.2 | XM_021843987.1 |
| NM_001204887.2 | XM_021844500.1 |
| NM_001205301.2 | XM_001652627.2 |
| NM_001206482.2 | XR_002501432.1 |
| NM_001206488.3 | XM_001650755.2 |
| NM_001206627.2 | XR_002500560.1 |
| NM_001206671.4 | XM_021853006.1 |
| NM_001206924.2 | XR_002500807.1 |
| NM_001206994.2 | XM_001651466.2 |
| NM_001207037.2 | XR_002499852.1 |
| NM_001207042.3 | XM_021849174.1 |
| NM_001207051.2 | XM_001657418.2 |
| NM_001207053.2 | XM_001652760.2 |
| NM_001214907.1 | XM_001658243.3 |
| NM_001226.4 | XM_021848786.1 |
| NM_001228.4 | XM_021857248.1 |
| NM_001235.5 | XM_021843728.1 |
| NM_001242375.1 | XM_021856224.1 |
| NM_001242376.3 | XM_021845689.1 |

| | |
|---|---|
| NM_001242399.2 | XM_021853873.1 |
| NM_001242560.2 | XR_002498978.1 |
| NM_001242589.3 | XM_001647950.2 |
| NM_001242597.2 | XM_021851961.1 |
| NM_001242598.2 | XM_021847323.1 |
| NM_001242630.2 | XM_001653229.2 |
| NM_001242639.2 | XM_021855068.1 |
| NM_001242692.2 | XM_021856681.1 |
| NM_001242763.2 | XM_001652384.2 |
| NM_001242798.2 | XM_021851567.1 |
| NM_001242818.2 | XM_021845075.1 |
| NM_001242820.2 | XM_021845789.1 |
| NM_001242847.2 | XM_021841454.1 |
| NM_001242848.2 | XM_001649759.2 |
| NM_001242851.1 | XR_002502780.1 |
| NM_001242858.3 | XR_002500034.1 |
| NM_001242868.2 | XR_002498946.1 |
| NM_001242875.3 | XM_021849999.1 |
| NM_001243042.1 | XM_021843324.1 |
| NM_001243108.2 | XR_002500096.1 |
| NM_001243116.2 | XM_001655235.2 |
| NM_001243137.2 | XM_021851745.1 |
| NM_001243177.4 | XM_021848500.1 |
| NM_001243237.2 | XM_001660369.2 |
| NM_001243256.2 | XM_001648309.2 |
| NM_001243271.2 | XR_002502328.1 |
| NM_001243423.2 | XM_021840502.1 |
| NM_001243597.2 | XM_021845448.1 |
| NM_001243754.2 | XM_021855103.1 |
| NM_001243782.2 | XM_021853044.1 |
| NM_001243784.2 | XM_021842664.1 |
| NM_001243798.2 | XM_021843415.1 |
| NM_001243799.1 | XM_021847262.1 |
| NM_001243925.2 | XM_021853718.1 |
| NM_001243942.1 | XM_001657893.2 |
| NM_001243998.2 | XM_001662691.2 |
| NM_001244.4 | XM_021844497.1 |
| NM_001244015.2 | XM_021853865.1 |
| NM_001244192.2 | XM_021850783.1 |
| NM_001245005.2 | XM_021853908.1 |
| NM_001247997.2 | XM_001648004.2 |
| NM_001248002.2 | XM_021856439.1 |
| NM_001251962.2 | XR_002500550.1 |
| NM_001251963.2 | XR_002499567.1 |
| NM_001251973.2 | XM_021857598.1 |
| NM_001251974.2 | XM_021855986.1 |
| NM_001252007.1 | XM_021838555.1 |
| NM_001252120.2 | XR_002501085.1 |
| NM_001252226.2 | XM_021844151.1 |
| NM_001252269.2 | XM_001662058.2 |

| | |
|---|---|
| NM_001253723.2 | XM_001656300.2 |
| NM_001253750.1 | XM_021842665.1 |
| NM_001253793.2 | XM_021851943.1 |
| NM_001253826.2 | XM_021839693.1 |
| NM_001253846.2 | XR_002499498.1 |
| NM_001253853.3 | XM_021851611.1 |
| NM_001253891.2 | XM_011495241.2 |
| NM_001254.4 | XR_002502586.1 |
| NM_001256047.2 | XR_002500610.1 |
| NM_001256054.3 | XM_021856685.1 |
| NM_001256092.2 | XM_001651573.2 |
| NM_001256155.2 | XM_001654825.3 |
| NM_001256165.1 | XM_001662721.2 |
| NM_001256293.2 | XM_001654683.2 |
| NM_001256304.3 | XM_001658422.3 |
| NM_001256328.2 | XM_021850491.1 |
| NM_001256337.3 | XM_021856443.1 |
| NM_001256348.2 | XR_002501117.1 |
| NM_001256371.2 | XM_021847995.1 |
| NM_001256412.2 | XM_021843833.1 |
| NM_001256423.2 | XR_002502269.1 |
| NM_001256468.2 | XR_002502606.1 |
| NM_001256494.2 | XM_001660777.2 |
| NM_001256496.2 | XM_001654663.2 |
| NM_001256503.2 | XM_001649169.2 |
| NM_001256549.2 | XM_021839552.1 |
| NM_001256601.1 | XM_021857554.1 |
| NM_001256605.2 | XM_021856624.1 |
| NM_001256619.2 | XM_001652103.2 |
| NM_001256630.1 | XM_001658226.2 |
| NM_001256643.1 | XR_002499161.1 |
| NM_001256677.1 | XR_002502195.1 |
| NM_001256688.2 | XR_002502099.1 |
| NM_001256701.1 | XR_002502041.1 |
| NM_001256725.2 | XR_002503095.1 |
| NM_001256727.2 | XM_001661421.2 |
| NM_001256753.2 | XM_021844756.1 |
| NM_001256758.1 | XM_001664031.2 |
| NM_001256860.1 | XM_021846816.1 |
| NM_001256864.2 | XM_001653428.2 |
| NM_001256873.1 | XM_021854109.1 |
| NM_001256887.3 | XM_021841327.1 |
| NM_001256917.2 | XR_002502159.1 |
| NM_001257.5 | XM_001663229.2 |
| NM_001257230.2 | XM_021849076.1 |
| NM_001257282.2 | XM_021844201.1 |
| NM_001257414.2 | XM_021856440.1 |
| NM_001258028.2 | XM_001651455.2 |
| NM_001258208.2 | XM_011495262.2 |
| NM_001258216.1 | XR_002502801.1 |

| | |
|---|---|
| NM_001258252.1 | XM_001654081.2 |
| NM_001258264.1 | XM_021839740.1 |
| NM_001258269.1 | XM_001648040.2 |
| NM_001258298.2 | XM_001655170.2 |
| NM_001258324.1 | XM_021852551.1 |
| NM_001258325.1 | XR_002502458.1 |
| NM_001258331.2 | XR_002500591.1 |
| NM_001258355.2 | XM_021845678.1 |
| NM_001258373.2 | XM_001651222.2 |
| NM_001258379.2 | XM_021848879.1 |
| NM_001258395.2 | XM_021843891.1 |
| NM_001258399.2 | XM_021854123.1 |
| NM_001258416.2 | XM_021838603.1 |
| NM_001258418.2 | XR_002503274.1 |
| NM_001258447.1 | XM_021843892.1 |
| NM_001258450.2 | XM_021842659.1 |
| NM_001258453.2 | XM_001650241.2 |
| NM_001260510.2 | XR_002502091.1 |
| NM_001261409.1 | XR_002499822.1 |
| NM_001261818.2 | XR_002501784.1 |
| NM_001261837.2 | XM_021851052.1 |
| NM_001261838.2 | XM_021850427.1 |
| NM_001265578.2 | XM_021843507.1 |
| NM_001265608.2 | XM_001663168.2 |
| NM_001267040.1 | XM_021854001.1 |
| NM_001267064.2 | XM_021842663.1 |
| NM_001267571.2 | XM_021856216.1 |
| NM_001267574.2 | XM_021843658.1 |
| NM_001269048.3 | XM_021841218.1 |
| NM_001270375.2 | XM_021846729.1 |
| NM_001270400.2 | XM_021843894.1 |
| NM_001270411.2 | XR_002503277.1 |
| NM_001270420.2 | XM_011494840.2 |
| NM_001270437.2 | XM_021852553.1 |
| NM_001270453.2 | XM_021854043.1 |
| NM_001270492.2 | XR_002502790.1 |
| NM_001270501.2 | XM_021844458.1 |
| NM_001270530.2 | XM_021852416.1 |
| NM_001270532.2 | XM_001651293.2 |
| NM_001270601.2 | XM_001660399.2 |
| NM_001270623.2 | XM_021846498.1 |
| NM_001270695.1 | XM_021846767.1 |
| NM_001270709.2 | XM_021846440.1 |
| NM_001270733.2 | XM_021843509.1 |
| NM_001270766.2 | XM_001662208.2 |
| NM_001270841.2 | XM_021848488.1 |
| NM_001270895.2 | XM_011495374.2 |
| NM_001270941.2 | XM_021842660.1 |
| NM_001271012.2 | XM_021852552.1 |
| NM_001271023.2 | XM_021856720.1 |

| | |
|---|---|
| NM_001271213.2 | XR_002501739.1 |
| NM_001271286.2 | XM_021842516.1 |
| NM_001271427.2 | XM_021843510.1 |
| NM_001271442.1 | XM_021848487.1 |
| NM_001271519.2 | XM_001659290.2 |
| NM_001271560.2 | XM_021853071.1 |
| NM_001271573.2 | XM_021843837.1 |
| NM_001271696.3 | XM_021844839.1 |
| NM_001271719.2 | XM_001654236.2 |
| NM_001271734.2 | XM_021843033.1 |
| NM_001271780.2 | XM_021846924.1 |
| NM_001271823.2 | XR_002499785.1 |
| NM_001271825.2 | XM_021844160.1 |
| NM_001271830.2 | XM_021857013.1 |
| NM_001271834.2 | XM_021856711.1 |
| NM_001271872.3 | XR_002501742.1 |
| NM_001271900.2 | XR_002501514.1 |
| NM_001271918.2 | XR_002502374.1 |
| NM_001271926.3 | XM_001661608.2 |
| NM_001271970.2 | XM_021853604.1 |
| NM_001272042.2 | XR_002500530.1 |
| NM_001272053.2 | XM_021840918.1 |
| NM_001272077.2 | XR_002499263.1 |
| NM_001272104.2 | XM_021851123.1 |
| NM_001273.5 | XM_021848162.1 |
| NM_001276252.2 | XM_001662946.2 |
| NM_001276271.2 | XR_002502637.1 |
| NM_001276375.2 | XM_001657456.2 |
| NM_001276395.2 | XM_021856714.1 |
| NM_001276437.2 | XM_021845103.1 |
| NM_001276453.2 | XR_002499367.1 |
| NM_001276501.2 | XM_021850823.1 |
| NM_001276723.2 | XR_002499280.1 |
| NM_001276726.2 | XM_021842508.1 |
| NM_001277075.3 | XM_021843670.1 |
| NM_001277081.1 | XR_002501804.1 |
| NM_001277083.2 | XM_021848640.1 |
| NM_001277227.2 | XR_002500478.1 |
| NM_001277403.2 | XM_021844185.1 |
| NM_001277971.2 | XM_021843834.1 |
| NM_001278067.1 | XR_002503244.1 |
| NM_001278162.2 | XM_021846791.1 |
| NM_001278210.2 | XM_001654001.2 |
| NM_001278214.2 | XM_001658199.2 |
| NM_001278215.2 | XM_001652124.2 |
| NM_001278267.1 | XM_001662720.2 |
| NM_001278296.2 | XM_001659087.2 |
| NM_001278300.2 | XM_021848501.1 |
| NM_001278309.2 | XM_021845690.1 |
| NM_001278398.2 | XM_001658579.2 |

| | |
|---|---|
| NM_001278428.3 | XM_021845355.1 |
| NM_001278432.1 | XM_021838747.1 |
| NM_001278464.2 | XM_021844156.1 |
| NM_001278471.2 | XR_002498691.1 |
| NM_001278499.2 | XM_021843681.1 |
| NM_001278500.2 | XR_002500391.1 |
| NM_001278503.2 | XM_021857106.1 |
| NM_001278512.2 | XM_021844836.1 |
| NM_001278524.2 | XM_001661870.2 |
| NM_001278546.2 | XM_021843656.1 |
| NM_001278548.1 | XR_002499824.1 |
| NM_001278549.2 | XM_021845287.1 |
| NM_001278559.2 | XM_001660321.2 |
| NM_001278592.2 | XR_002500057.1 |
| NM_001278599.2 | XM_001653625.3 |
| NM_001278603.2 | XM_021848638.1 |
| NM_001278606.2 | XM_001659405.2 |
| NM_001278610.2 | XR_002498945.1 |
| NM_001278618.2 | XM_021856315.1 |
| NM_001278629.2 | XM_021838855.1 |
| NM_001278639.2 | XM_021838860.1 |
| NM_001278641.2 | XM_001652025.2 |
| NM_001278645.2 | XM_021843718.1 |
| NM_001278678.2 | XM_001658777.2 |
| NM_001278715.2 | XM_001652492.2 |
| NM_001278794.2 | XM_001658036.3 |
| NM_001278928.2 | XM_021844400.1 |
| NM_001279.4 | XM_021838854.1 |
| NM_001279349.2 | XM_001656551.2 |
| NM_001280550.2 | XR_002499342.1 |
| NM_001280795.2 | XM_021847660.1 |
| NM_001280799.2 | XM_021843716.1 |
| NM_001281518.3 | XR_002499823.1 |
| NM_001281520.2 | XR_002502953.1 |
| NM_001282068.2 | XM_001662289.2 |
| NM_001282113.2 | XM_021842507.1 |
| NM_001282117.2 | XR_002498731.1 |
| NM_001282190.2 | XM_021838091.1 |
| NM_001282195.2 | XM_021845679.1 |
| NM_001282215.2 | XM_021838862.1 |
| NM_001282237.2 | XM_021840238.1 |
| NM_001282305.1 | XM_021856773.1 |
| NM_001282310.2 | XM_001659866.2 |
| NM_001282356.2 | XM_021856164.1 |
| NM_001282398.2 | XM_021845076.1 |
| NM_001282418.2 | XM_021851047.1 |
| NM_001282419.3 | XM_001654676.2 |
| NM_001282456.4 | XM_021854029.1 |
| NM_001282466.2 | XM_021844261.1 |
| NM_001282597.3 | XM_021846764.1 |

| | |
|---|---|
| NM_001282624.2 | XM_021848026.1 |
| NM_001282706.2 | XM_021849061.1 |
| NM_001282715.2 | XM_021851069.1 |
| NM_001282719.2 | XM_021848351.1 |
| NM_001282753.2 | XM_021856289.1 |
| NM_001282755.2 | XR_002501131.1 |
| NM_001282790.2 | XM_021854033.1 |
| NM_001282792.2 | XR_002499707.1 |
| NM_001282796.2 | XM_001658039.2 |
| NM_001282800.2 | XM_021854055.1 |
| NM_001282803.2 | XM_021838861.1 |
| NM_001282806.2 | XM_021847333.1 |
| NM_001282851.2 | XM_021850916.1 |
| NM_001282854.2 | XM_021843822.1 |
| NM_001282885.2 | XM_001653951.2 |
| NM_001282914.2 | XM_021844197.1 |
| NM_001282920.1 | XM_021846549.1 |
| NM_001282921.2 | XR_002501849.1 |
| NM_001282933.2 | XM_001652622.2 |
| NM_001282959.2 | XM_021844381.1 |
| NM_001283004.3 | XM_021839480.1 |
| NM_001283023.2 | XM_021841341.1 |
| NM_001283027.1 | XM_021854014.1 |
| NM_001283060.2 | XM_021851048.1 |
| NM_001283235.3 | XM_021849826.1 |
| NM_001284210.2 | XM_021854981.1 |
| NM_001284502.2 | XM_021846770.1 |
| NM_001284511.2 | XR_002500281.1 |
| NM_001284528.2 | XM_021850430.1 |
| NM_001285.4 | XM_021851125.1 |
| NM_001285526.2 | XM_021853547.1 |
| NM_001286.5 | XM_001654897.2 |
| NM_001286035.2 | XM_021846562.1 |
| NM_001286051.2 | XM_001660659.2 |
| NM_001286094.2 | XM_021844163.1 |
| NM_001286107.2 | XM_001663935.2 |
| NM_001286139.2 | XR_002498993.1 |
| NM_001286197.2 | XR_002503138.1 |
| NM_001286206.2 | XM_021845548.1 |
| NM_001286210.2 | XM_021845414.1 |
| NM_001286354.1 | XM_021842384.1 |
| NM_001286399.2 | XM_001650789.2 |
| NM_001286427.1 | XM_021840096.1 |
| NM_001286536.2 | XM_021844333.1 |
| NM_001286537.2 | XM_001663454.2 |
| NM_001286550.2 | XM_021857047.1 |
| NM_001286563.3 | XM_021838761.1 |
| NM_001286583.2 | XM_021838863.1 |
| NM_001286613.2 | XM_021848233.1 |
| NM_001286639.2 | XM_021857485.1 |

| | |
|---|---|
| NM_001286655.1 | XM_021854145.1 |
| NM_001286688.2 | XM_021840811.1 |
| NM_001286695.1 | XM_021844378.1 |
| NM_001286700.2 | XM_021846759.1 |
| NM_001286702.2 | XM_021852609.1 |
| NM_001286749.2 | XM_021844382.1 |
| NM_001286751.2 | XM_001661138.2 |
| NM_001286756.2 | XR_002501424.1 |
| NM_001286760.1 | XM_021838234.1 |
| NM_001286810.2 | XM_021857254.1 |
| NM_001286828.2 | XR_002500694.1 |
| NM_001286830.2 | XM_021843992.1 |
| NM_001286957.2 | XR_002499118.1 |
| NM_001287005.2 | XM_001660027.2 |
| NM_001287060.2 | XM_021856407.1 |
| NM_001287136.1 | XM_021846773.1 |
| NM_001287190.2 | XM_021844383.1 |
| NM_001287262.2 | XR_002500486.1 |
| NM_001287390.3 | XM_001659691.2 |
| NM_001287394.2 | XM_021844003.1 |
| NM_001287422.2 | XM_021851196.1 |
| NM_001287430.1 | XM_021849663.1 |
| NM_001287432.2 | XM_021850694.1 |
| NM_001287509.2 | XM_021844663.1 |
| NM_001287513.2 | XR_002501488.1 |
| NM_001287525.2 | XM_001651785.2 |
| NM_001287601.1 | XM_021847361.1 |
| NM_001288566.2 | XM_001657123.2 |
| NM_001288568.2 | XM_021847338.1 |
| NM_001288582.2 | XM_001650767.2 |
| NM_001288643.2 | XR_002498889.1 |
| NM_001288653.2 | XM_021851600.1 |
| NM_001288662.1 | XM_021850690.1 |
| NM_001288705.3 | XM_021851886.1 |
| NM_001288713.1 | XR_002499069.1 |
| NM_001288732.2 | XR_002500252.1 |
| NM_001288746.2 | XR_002499987.1 |
| NM_001288752.2 | XR_002499817.1 |
| NM_001288836.1 | XM_021846782.1 |
| NM_001288951.2 | XM_021839661.1 |
| NM_001288967.2 | XM_021843197.1 |
| NM_001288975.2 | XM_021847104.1 |
| NM_001288980.2 | XM_021852814.1 |
| NM_001288989.2 | XM_021853980.1 |
| NM_001289035.2 | XM_021856847.1 |
| NM_001289037.2 | XM_021843159.1 |
| NM_001289057.2 | XR_002499786.1 |
| NM_001289129.2 | XM_021857238.1 |
| NM_001289152.2 | XM_021856678.1 |
| NM_001289173.2 | XR_002500469.1 |

| | |
|---|---|
| NM_001289180.2 | XM_021838251.1 |
| NM_001289184.2 | XR_002500222.1 |
| NM_001289189.2 | XM_021849703.1 |
| NM_001289396.1 | XM_021846804.1 |
| NM_001289399.1 | XM_011494934.2 |
| NM_001289911.2 | XM_021843703.1 |
| NM_001289975.1 | XR_002501820.1 |
| NM_001290030.2 | XM_021839624.1 |
| NM_001290059.2 | XM_001653429.2 |
| NM_001290184.2 | XR_002498937.1 |
| NM_001290194.2 | XM_021838571.1 |
| NM_001290233.2 | XM_021853989.1 |
| NM_001290258.2 | XR_002501491.1 |
| NM_001290360.3 | XM_001660770.2 |
| NM_001291281.3 | XM_021854010.1 |
| NM_001291308.3 | XM_021852890.1 |
| NM_001291328.2 | XM_021856408.1 |
| NM_001291454.2 | XR_002501306.1 |
| NM_001291491.2 | XM_001650901.3 |
| NM_001291527.2 | XM_001657703.2 |
| NM_001291588.2 | XM_021851568.1 |
| NM_001291700.1 | XM_021852363.1 |
| NM_001291716.2 | XR_002501474.1 |
| NM_001291719.2 | XM_001650661.2 |
| NM_001291732.2 | XM_021842364.1 |
| NM_001291738.1 | XM_001647871.2 |
| NM_001291813.2 | XM_021842410.1 |
| NM_001291826.2 | XM_021842587.1 |
| NM_001291837.2 | XM_021841546.1 |
| NM_001291860.2 | XR_002503030.1 |
| NM_001291873.3 | XM_001650333.2 |
| NM_001291879.2 | XM_021854930.1 |
| NM_001291896.2 | XM_001647936.2 |
| NM_001291918.2 | XM_021841407.1 |
| NM_001291931.2 | XM_021842381.1 |
| NM_001291956.3 | XM_021851140.1 |
| NM_001291979.2 | XM_021843126.1 |
| NM_001292004.2 | XM_001652207.2 |
| NM_001292018.2 | XM_021853839.1 |
| NM_001292048.2 | XM_001659191.3 |
| NM_001293213.2 | XM_001652668.2 |
| NM_001293628.2 | XR_002499419.1 |
| NM_001293810.1 | XM_001649463.2 |
| NM_001294332.2 | XM_021853112.1 |
| NM_001294335.2 | XR_002502296.1 |
| NM_001294338.2 | XM_001657698.2 |
| NM_001297424.2 | XM_021852558.1 |
| NM_001297435.1 | XM_021839577.1 |
| NM_001297440.2 | XM_021845523.1 |
| NM_001297569.2 | XM_001650453.2 |

| | |
|---|---|
| NM_001297592.2 | XM_021856310.1 |
| NM_001297610.2 | XR_002501445.1 |
| NM_001297643.2 | XR_002501588.1 |
| NM_001297647.2 | XM_021847809.1 |
| NM_001297648.2 | XM_001654094.2 |
| NM_001297654.2 | XM_021846288.1 |
| NM_001297664.1 | XM_021854105.1 |
| NM_001297669.3 | XM_021851650.1 |
| NM_001297710.2 | XM_021853413.1 |
| NM_001297723.2 | XR_002502372.1 |
| NM_001297724.2 | XM_001663951.2 |
| NM_001297759.2 | XM_021841448.1 |
| NM_001297774.2 | XM_001654289.2 |
| NM_001297776.2 | XM_021852923.1 |
| NM_001300756.2 | XM_021847541.1 |
| NM_001300786.2 | XR_002502740.1 |
| NM_001300787.2 | XM_021846438.1 |
| NM_001300856.2 | XM_021840924.1 |
| NM_001300859.2 | XM_021846478.1 |
| NM_001300867.2 | XM_021848859.1 |
| NM_001300885.2 | XR_002500027.1 |
| NM_001300887.2 | XM_001661874.2 |
| NM_001300898.2 | XM_021857483.1 |
| NM_001300911.2 | XR_002502197.1 |
| NM_001300919.1 | XM_021850685.1 |
| NM_001300926.2 | XR_002503190.1 |
| NM_001300938.2 | XR_002502999.1 |
| NM_001300946.3 | XM_021857110.1 |
| NM_001300960.2 | XM_021846854.1 |
| NM_001300982.2 | XM_021837317.1 |
| NM_001300990.2 | XM_021839485.1 |
| NM_001300992.2 | XM_001657621.2 |
| NM_001301021.2 | XM_011494795.2 |
| NM_001301028.3 | XM_021849961.1 |
| NM_001301031.1 | XM_001659105.2 |
| NM_001301097.1 | XM_021855772.1 |
| NM_001301134.2 | XR_002500219.1 |
| NM_001301244.2 | XM_001659476.2 |
| NM_001301253.2 | XM_001658646.2 |
| NM_001301409.2 | XM_001652481.2 |
| NM_001301645.2 | XM_021839738.1 |
| NM_001301648.2 | XM_021837951.1 |
| NM_001301775.2 | XM_021853956.1 |
| NM_001301828.2 | XM_021844966.1 |
| NM_001301830.2 | XM_021842951.1 |
| NM_001302509.2 | XM_021843803.1 |
| NM_001302554.2 | XM_021850134.1 |
| NM_001302622.2 | XM_021843526.1 |
| NM_001302688.2 | XR_002500729.1 |
| NM_001302695.2 | XM_021846883.1 |

| | |
|---|---|
| NM_001302817.3 | XM_021843001.1 |
| NM_001302836.2 | XM_001656173.2 |
| NM_001303000.2 | XM_001653370.2 |
| NM_001303030.2 | XR_002499134.1 |
| NM_001303037.2 | XM_021841596.1 |
| NM_001303047.2 | XM_011495175.2 |
| NM_001303235.2 | XM_021857168.1 |
| NM_001303255.3 | XM_001655409.2 |
| NM_001303276.2 | XM_021854172.1 |
| NM_001303459.3 | XM_001649686.2 |
| NM_001303475.1 | XM_021848054.1 |
| NM_001303477.2 | XM_021844447.1 |
| NM_001303487.3 | XM_001655924.2 |
| NM_001303498.3 | XM_021844461.1 |
| NM_001303615.2 | XM_021842377.1 |
| NM_001303619.2 | XM_001654548.2 |
| NM_001304331.2 | XM_001656508.2 |
| NM_001304345.2 | XM_001649188.2 |
| NM_001304360.2 | XM_021849300.1 |
| NM_001304488.1 | XM_021851138.1 |
| NM_001304489.1 | XM_021856894.1 |
| NM_001304500.2 | XM_021853405.1 |
| NM_001304507.1 | XM_021848615.1 |
| NM_001304513.2 | XR_002502564.1 |
| NM_001304549.2 | XM_021845107.1 |
| NM_001304569.2 | XR_002500382.1 |
| NM_001304763.2 | XM_021852372.1 |
| NM_001304808.3 | XM_021847718.1 |
| NM_001304814.2 | XM_021851750.1 |
| NM_001304819.2 | XM_021853279.1 |
| NM_001304834.2 | XM_021837528.1 |
| NM_001304942.2 | XM_021856154.1 |
| NM_001304947.3 | XR_002500493.1 |
| NM_001304957.2 | XM_001656281.2 |
| NM_001304993.2 | XM_021854506.1 |
| NM_001305080.2 | XR_002502366.1 |
| NM_001305108.1 | XR_002500983.1 |
| NM_001305208.2 | XM_021850975.1 |
| NM_001305457.2 | XM_021854332.1 |
| NM_001305545.1 | XM_021850808.1 |
| NM_001305625.2 | XM_021841547.1 |
| NM_001306070.2 | XM_021842990.1 |
| NM_001306092.2 | XM_021838740.1 |
| NM_001306151.2 | XR_002502097.1 |
| NM_001306160.3 | XM_011494945.2 |
| NM_001306172.2 | XM_021851061.1 |
| NM_001306197.2 | XM_021838684.1 |
| NM_001306200.2 | XM_021841902.1 |
| NM_001306214.2 | XM_021844071.1 |
| NM_001307976.2 | XM_021842484.1 |

| | |
|---|---|
| NM_001308023.2 | XR_002502014.1 |
| NM_001308028.2 | XM_021853447.1 |
| NM_001308075.2 | XM_021848645.1 |
| NM_001308080.2 | XM_021852913.1 |
| NM_001308097.2 | XM_021850182.1 |
| NM_001308105.1 | XM_021838341.1 |
| NM_001308119.2 | XM_021856589.1 |
| NM_001308161.1 | XM_021842916.1 |
| NM_001308169.2 | XM_001660482.2 |
| NM_001308185.2 | XM_021837693.1 |
| NM_001308203.2 | XM_021843896.1 |
| NM_001308237.2 | XM_001657229.2 |
| NM_001308242.2 | XR_002501766.1 |
| NM_001308264.2 | XM_021855639.1 |
| NM_001308342.2 | XM_021843865.1 |
| NM_001308430.2 | XM_001661471.2 |
| NM_001308469.3 | XR_002498875.1 |
| NM_001308954.2 | XM_021840545.1 |
| NM_001309193.2 | XM_021851136.1 |
| NM_001309427.2 | XM_021853484.1 |
| NM_001309450.2 | XM_021855888.1 |
| NM_001310137.3 | XM_021843788.1 |
| NM_001310160.2 | XR_002499812.1 |
| NM_001311197.2 | XM_021842376.1 |
| NM_001311311.2 | XM_021852910.1 |
| NM_001312921.2 | XM_011494754.2 |
| NM_001313741.1 | XM_021844074.1 |
| NM_001313943.2 | XR_002502603.1 |
| NM_001313953.3 | XM_021851139.1 |
| NM_001314052.2 | XM_021850657.1 |
| NM_001315.3 | XM_001660731.2 |
| NM_001316314.3 | XM_021850835.1 |
| NM_001316317.2 | XR_002502679.1 |
| NM_001316348.2 | XM_021851137.1 |
| NM_001316358.2 | XM_021841339.1 |
| NM_001316374.2 | XR_002500901.1 |
| NM_001316943.2 | XM_021845506.1 |
| NM_001316965.2 | XM_001656167.2 |
| NM_001317019.1 | XM_021851824.1 |
| NM_001317034.1 | XR_002499590.1 |
| NM_001317062.2 | XM_021842563.1 |
| NM_001317103.2 | XR_002500827.1 |
| NM_001317120.2 | XM_001662114.2 |
| NM_001317161.2 | XM_021844504.1 |
| NM_001317184.2 | XR_002498975.1 |
| NM_001317406.3 | XM_021841474.1 |
| NM_001317721.2 | XM_001660709.2 |
| NM_001317740.2 | XM_001657255.3 |
| NM_001317774.1 | XR_002499599.1 |
| NM_001317781.2 | XM_021849557.1 |

| | |
|---|---|
| NM_001317788.2 | XM_021841123.1 |
| NM_001317789.1 | XR_002503215.1 |
| NM_001317804.2 | XM_021857050.1 |
| NM_001317891.1 | XM_021845196.1 |
| NM_001317895.2 | XM_001648213.2 |
| NM_001317919.2 | XM_021842472.1 |
| NM_001317935.2 | XM_021843007.1 |
| NM_001317943.2 | XM_001662940.2 |
| NM_001317992.2 | XM_021851965.1 |
| NM_001318021.1 | XM_001663913.2 |
| NM_001318022.2 | XM_021842475.1 |
| NM_001318036.2 | XM_001659128.2 |
| NM_001318042.2 | XM_021839379.1 |
| NM_001318068.1 | XM_001654523.2 |
| NM_001318082.2 | XM_001648138.2 |
| NM_001318091.2 | XM_001657015.2 |
| NM_001318154.2 | XM_001652433.2 |
| NM_001318199.3 | XM_021853134.1 |
| NM_001318226.2 | XM_001647622.2 |
| NM_001318238.2 | XM_021848336.1 |
| NM_001318239.2 | XR_002502621.1 |
| NM_001318500.2 | XM_021855469.1 |
| NM_001318710.2 | XM_021857141.1 |
| NM_001318715.2 | XM_001663442.2 |
| NM_001318726.2 | XM_021849829.1 |
| NM_001318761.2 | XR_002503012.1 |
| NM_001318781.2 | XM_021847329.1 |
| NM_001318787.2 | XM_011495397.2 |
| NM_001318800.2 | XM_001658372.2 |
| NM_001318815.2 | XR_002498748.1 |
| NM_001318830.2 | XM_001659903.2 |
| NM_001318841.2 | XR_002499356.1 |
| NM_001318855.1 | XM_021842564.1 |
| NM_001318870.2 | XM_001657774.2 |
| NM_001318873.2 | XM_021856900.1 |
| NM_001318909.2 | XM_021841084.1 |
| NM_001318928.2 | XM_021857108.1 |
| NM_001318942.1 | XR_002502872.1 |
| NM_001318945.2 | XM_021847086.1 |
| NM_001319039.2 | XM_021850042.1 |
| NM_001319062.2 | XM_021855047.1 |
| NM_001319092.1 | XM_001653496.2 |
| NM_001319110.2 | XM_021847802.1 |
| NM_001319198.2 | XM_021845932.1 |
| NM_001319217.2 | XM_021857071.1 |
| NM_001319220.2 | XM_021842782.1 |
| NM_001319228.2 | XM_021851967.1 |
| NM_001319229.2 | XM_021856951.1 |
| NM_001319242.1 | XM_021850097.1 |
| NM_001319295.2 | XM_021854944.1 |

| | |
|---|---|
| NM_001319297.2 | XM_021856715.1 |
| NM_001319658.2 | XR_002500678.1 |
| NM_001319664.2 | XM_021848764.1 |
| NM_001319952.2 | XM_021852470.1 |
| NM_001319960.2 | XM_021855633.1 |
| NM_001319988.2 | XM_001654710.2 |
| NM_001320000.2 | XM_001659028.2 |
| NM_001320050.2 | XM_001663670.2 |
| NM_001320134.1 | XM_021848707.1 |
| NM_001320210.2 | XM_011495042.2 |
| NM_001320219.2 | XM_021847031.1 |
| NM_001320321.2 | XM_021847428.1 |
| NM_001320351.2 | XM_021849998.1 |
| NM_001320365.2 | XM_001657243.3 |
| NM_001320458.2 | XM_021854057.1 |
| NM_001320488.2 | XM_021840643.1 |
| NM_001320511.2 | XM_001654262.2 |
| NM_001320546.3 | XM_021853690.1 |
| NM_001320555.2 | XM_021855041.1 |
| NM_001320557.2 | XM_021846765.1 |
| NM_001320585.1 | XR_002498800.1 |
| NM_001320592.2 | XM_011494887.2 |
| NM_001320597.2 | XM_021842405.1 |
| NM_001320608.2 | XM_021848885.1 |
| NM_001320624.1 | XM_021851287.1 |
| NM_001320714.2 | XM_021845029.1 |
| NM_001320715.2 | XM_021838512.1 |
| NM_001320722.3 | XM_001656686.2 |
| NM_001320746.3 | XM_021850775.1 |
| NM_001320750.3 | XM_021851073.1 |
| NM_001320830.2 | XM_021838010.1 |
| NM_001320840.2 | XM_021841085.1 |
| NM_001320849.2 | XM_021856541.1 |
| NM_001320851.2 | XR_002500421.1 |
| NM_001320865.2 | XM_001653939.2 |
| NM_001320870.2 | XM_001659150.2 |
| NM_001320872.2 | XM_021856892.1 |
| NM_001320883.2 | XM_021851477.1 |
| NM_001320933.2 | XM_021844287.1 |
| NM_001320948.2 | XM_021852024.1 |
| NM_001320953.2 | XM_001661599.2 |
| NM_001320971.2 | XM_001654839.2 |
| NM_001321007.2 | XM_001648211.2 |
| NM_001321015.2 | XM_001652006.2 |
| NM_001321018.2 | XR_002499649.1 |
| NM_001321072.1 | XM_021856460.1 |
| NM_001321107.2 | XM_021853424.1 |
| NM_001321144.2 | XM_021838340.1 |
| NM_001321224.2 | XM_001651325.2 |
| NM_001321234.2 | XR_002500909.1 |

| | |
|---|---|
| NM_001321266.2 | XM_021840091.1 |
| NM_001321285.2 | XM_001659467.2 |
| NM_001321356.2 | XM_021848040.1 |
| NM_001321363.2 | XM_021846481.1 |
| NM_001321365.2 | XM_021852505.1 |
| NM_001321369.2 | XM_021854512.1 |
| NM_001321381.3 | XR_002502935.1 |
| NM_001321458.2 | XM_001656432.2 |
| NM_001321470.1 | XM_021850938.1 |
| NM_001321501.2 | XR_002498704.1 |
| NM_001321525.1 | XM_001659473.2 |
| NM_001321592.2 | XM_021850720.1 |
| NM_001321612.2 | XR_002500160.1 |
| NM_001321617.2 | XM_021843198.1 |
| NM_001321677.2 | XR_002503261.1 |
| NM_001321747.2 | XM_001650139.2 |
| NM_001321808.2 | XR_002500018.1 |
| NM_001321868.2 | XM_021857564.1 |
| NM_001321881.2 | XM_021856331.1 |
| NM_001321907.3 | XM_001662070.2 |
| NM_001321908.3 | XM_021856975.1 |
| NM_001321923.2 | XM_021846585.1 |
| NM_001321927.1 | XM_021846993.1 |
| NM_001321929.1 | XM_021852621.1 |
| NM_001321935.1 | XM_001658594.3 |
| NM_001321953.2 | XM_021854513.1 |
| NM_001321972.2 | XR_002500077.1 |
| NM_001321983.2 | XM_001664214.2 |
| NM_001321993.3 | XM_001660375.2 |
| NM_001322012.2 | XR_002502951.1 |
| NM_001322028.2 | XR_002501124.1 |
| NM_001322029.2 | XM_021845873.1 |
| NM_001322085.2 | XM_021857088.1 |
| NM_001322147.2 | XM_021838562.1 |
| NM_001322164.2 | XM_001658061.2 |
| NM_001322180.2 | XM_021842480.1 |
| NM_001322215.2 | XM_021845190.1 |
| NM_001322221.2 | XM_021844471.1 |
| NM_001322252.2 | XM_021852397.1 |
| NM_001322256.2 | XR_002502005.1 |
| NM_001322296.2 | XM_021854818.1 |
| NM_001322367.1 | XM_021848380.1 |
| NM_001322391.2 | XM_021837265.1 |
| NM_001322402.3 | XM_021845088.1 |
| NM_001322494.1 | XM_021852447.1 |
| NM_001322840.2 | XM_021851702.1 |
| NM_001322861.2 | XM_001662825.2 |
| NM_001322889.2 | XM_021855088.1 |
| NM_001322902.2 | XM_021854300.1 |
| NM_001322920.1 | XM_001653224.2 |

| | |
|---|---|
| NM_001322929.2 | XM_001647730.2 |
| NM_001322930.2 | XM_021854819.1 |
| NM_001322963.2 | XM_021848231.1 |
| NM_001322971.2 | XM_021839797.1 |
| NM_001323022.1 | XR_002500313.1 |
| NM_001323067.2 | XM_021841462.1 |
| NM_001323073.1 | XM_021842938.1 |
| NM_001323255.2 | XM_021842382.1 |
| NM_001323304.2 | XM_021851536.1 |
| NM_001323329.2 | XM_021843002.1 |
| NM_001323342.2 | XM_001654503.2 |
| NM_001323360.2 | XM_001648559.2 |
| NM_001323367.2 | XM_021855119.1 |
| NM_001323377.2 | XM_021845198.1 |
| NM_001323382.2 | XM_021840727.1 |
| NM_001323389.2 | XM_021849889.1 |
| NM_001323395.2 | XM_021852028.1 |
| NM_001323543.2 | XM_001663586.2 |
| NM_001323563.2 | XM_021838195.1 |
| NM_001323575.2 | XM_021851537.1 |
| NM_001323578.2 | XR_002499262.1 |
| NM_001323644.2 | XM_021856069.1 |
| NM_001323657.2 | XM_001654882.2 |
| NM_001323674.2 | XM_001659207.2 |
| NM_001323681.2 | XM_021837858.1 |
| NM_001323906.2 | XM_001648230.3 |
| NM_001323907.2 | XM_021849187.1 |
| NM_001323911.2 | XM_021844141.1 |
| NM_001323930.2 | XR_002503226.1 |
| NM_001323948.2 | XM_021839729.1 |
| NM_001323964.2 | XM_021852067.1 |
| NM_001323972.2 | XM_021845703.1 |
| NM_001323973.2 | XM_021852546.1 |
| NM_001324002.1 | XM_001657709.2 |
| NM_001324003.1 | XM_021842311.1 |
| NM_001324025.2 | XM_021854104.1 |
| NM_001324029.3 | XM_021843255.1 |
| NM_001324067.2 | XR_002503097.1 |
| NM_001324092.2 | XM_021843501.1 |
| NM_001324093.2 | XR_002500417.1 |
| NM_001324094.2 | XM_001654213.2 |
| NM_001324109.2 | XM_021839567.1 |
| NM_001324116.3 | XM_021857437.1 |
| NM_001324119.2 | XM_001659086.2 |
| NM_001324138.2 | XM_001655261.2 |
| NM_001324169.2 | XM_021844172.1 |
| NM_001324171.2 | XR_002503134.1 |
| NM_001324178.2 | XM_001652473.2 |
| NM_001324186.2 | XM_021853067.1 |
| NM_001324187.1 | XR_002502537.1 |

| | |
|---|---|
| NM_001324198.2 | XM_021848762.1 |
| NM_001324201.2 | XM_021839043.1 |
| NM_001324202.2 | XM_001658100.2 |
| NM_001324220.2 | XM_021849726.1 |
| NM_001324230.2 | XM_021842388.1 |
| NM_001324242.2 | XM_021843708.1 |
| NM_001324283.2 | XM_021854553.1 |
| NM_001324294.2 | XM_021853233.1 |
| NM_001324296.2 | XM_001650039.2 |
| NM_001324303.2 | XM_001662227.2 |
| NM_001324304.2 | XM_021845034.1 |
| NM_001324343.2 | XM_021852674.1 |
| NM_001324353.2 | XM_001663607.2 |
| NM_001324354.2 | XR_002502859.1 |
| NM_001324357.2 | XM_021849648.1 |
| NM_001324387.2 | XM_021843721.1 |
| NM_001324390.1 | XM_021848536.1 |
| NM_001324415.2 | XM_021857073.1 |
| NM_001324446.2 | XM_021842596.1 |
| NM_001326314.2 | XM_021845197.1 |
| NM_001326332.2 | XR_002502965.1 |
| NM_001328620.2 | XM_021850038.1 |
| NM_001328627.1 | XM_001654943.2 |
| NM_001328646.3 | XM_021839037.1 |
| NM_001328654.2 | XM_021851900.1 |
| NM_001328674.2 | XR_002499032.1 |
| NM_001329137.2 | XM_001659994.3 |
| NM_001329144.2 | XM_021849145.1 |
| NM_001329201.2 | XR_002502238.1 |
| NM_001329442.2 | XM_001650048.2 |
| NM_001329586.2 | XM_001657486.2 |
| NM_001329615.2 | XR_002501018.1 |
| NM_001329633.2 | XR_002500411.1 |
| NM_001329634.1 | XM_021853795.1 |
| NM_001329718.2 | XM_021856887.1 |
| NM_001329723.2 | XM_021856208.1 |
| NM_001329763.2 | XM_021842085.1 |
| NM_001329775.2 | XM_001656698.2 |
| NM_001329803.2 | XM_021837992.1 |
| NM_001329911.2 | XM_001654504.2 |
| NM_001329919.2 | XM_001663991.2 |
| NM_001329920.2 | XM_021844370.1 |
| NM_001329974.2 | XM_021843502.1 |
| NM_001330000.2 | XM_001655288.2 |
| NM_001330059.2 | XM_021845822.1 |
| NM_001330062.2 | XM_021851241.1 |
| NM_001330094.2 | XM_021848400.1 |
| NM_001330113.2 | XM_001655801.2 |
| NM_001330115.2 | XM_001656755.2 |
| NM_001330125.1 | XM_021850727.1 |

| | |
|---|---|
| NM_001330128.2 | XM_021843649.1 |
| NM_001330130.2 | XR_002502355.1 |
| NM_001330132.2 | XM_021840371.1 |
| NM_001330148.2 | XM_021837315.1 |
| NM_001330157.2 | XM_001653262.2 |
| NM_001330170.2 | XM_001660969.2 |
| NM_001330183.2 | XM_001650004.2 |
| NM_001330187.2 | XM_021850811.1 |
| NM_001330214.1 | XM_021845007.1 |
| NM_001330227.3 | XR_002501470.1 |
| NM_001330250.1 | XM_021850773.1 |
| NM_001330252.2 | XM_021837307.1 |
| NM_001330285.2 | XM_001661936.2 |
| NM_001330303.2 | XM_021837320.1 |
| NM_001330310.2 | XM_021841675.1 |
| NM_001330316.2 | XR_002499059.1 |
| NM_001330326.2 | XM_021852711.1 |
| NM_001330354.2 | XM_021842302.1 |
| NM_001330373.2 | XM_021842308.1 |
| NM_001330380.2 | XM_021855513.1 |
| NM_001330388.2 | XM_021842303.1 |
| NM_001330404.2 | XM_021854650.1 |
| NM_001330406.2 | XM_021856721.1 |
| NM_001330420.2 | XM_021841356.1 |
| NM_001330442.2 | XM_021846281.1 |
| NM_001330447.2 | XM_001657901.2 |
| NM_001330475.2 | XM_021845560.1 |
| NM_001330514.2 | XM_021857246.1 |
| NM_001330521.2 | XR_002498699.1 |
| NM_001330537.2 | XM_021856324.1 |
| NM_001330570.3 | XM_021845933.1 |
| NM_001330613.2 | XM_021851419.1 |
| NM_001330645.2 | XM_001657197.2 |
| NM_001330713.2 | XM_021854464.1 |
| NM_001330721.2 | XR_002499859.1 |
| NM_001330723.2 | XM_021843375.1 |
| NM_001330728.1 | XM_001657341.2 |
| NM_001330729.2 | XM_021856415.1 |
| NM_001330750.2 | XM_001649107.2 |
| NM_001330759.2 | XM_021841674.1 |
| NM_001330996.3 | XM_021842470.1 |
| NM_001331066.2 | XM_021837860.1 |
| NM_001331071.3 | XM_021843024.1 |
| NM_001331158.2 | XM_001658754.2 |
| NM_001331211.2 | XM_021842228.1 |
| NM_001344.4 | XM_021853747.1 |
| NM_001345881.2 | XR_002499098.1 |
| NM_001345882.2 | XR_002503002.1 |
| NM_001345888.2 | XR_002500582.1 |
| NM_001345909.2 | XM_021844106.1 |

| | |
|---|---|
| NM_001345949.3 | XM_021844351.1 |
| NM_001345977.1 | XR_002499612.1 |
| NM_001346004.2 | XM_021842511.1 |
| NM_001346070.2 | XR_002501380.1 |
| NM_001346103.2 | XM_001650120.2 |
| NM_001346116.2 | XM_021846390.1 |
| NM_001346139.1 | XM_021849423.1 |
| NM_001346223.2 | XR_002498846.1 |
| NM_001346227.2 | XM_021842895.1 |
| NM_001346229.2 | XM_021850572.1 |
| NM_001346253.2 | XM_001648666.2 |
| NM_001346254.2 | XM_021852619.1 |
| NM_001346268.2 | XM_021844403.1 |
| NM_001346311.2 | XM_021843252.1 |
| NM_001346341.2 | XM_021847805.1 |
| NM_001346342.2 | XM_021840604.1 |
| NM_001346353.2 | XM_001658075.2 |
| NM_001346382.2 | XM_021851428.1 |
| NM_001346404.1 | XM_021854339.1 |
| NM_001346413.2 | XM_021851812.1 |
| NM_001346450.2 | XM_021845875.1 |
| NM_001346457.2 | XR_002498602.1 |
| NM_001346474.2 | XM_001657900.2 |
| NM_001346477.2 | XM_021852029.1 |
| NM_001346482.1 | XM_021843254.1 |
| NM_001346543.2 | XR_002502356.1 |
| NM_001346548.2 | XM_021840011.1 |
| NM_001346549.2 | XM_001649597.2 |
| NM_001346560.3 | XM_021843210.1 |
| NM_001346603.2 | XM_021852579.1 |
| NM_001346719.2 | XM_001652301.2 |
| NM_001346818.1 | XM_021844986.1 |
| NM_001346862.2 | XM_011495227.2 |
| NM_001346932.2 | XM_021844769.1 |
| NM_001346934.2 | XM_021852309.1 |
| NM_001346946.2 | XR_002499982.1 |
| NM_001346955.2 | XM_001650094.2 |
| NM_001346972.2 | XM_021845701.1 |
| NM_001347015.2 | XM_021851214.1 |
| NM_001347018.2 | XM_021841599.1 |
| NM_001347031.2 | XM_021844671.1 |
| NM_001347096.2 | XM_001653925.2 |
| NM_001347219.2 | XM_001655139.2 |
| NM_001347664.1 | XM_021857247.1 |
| NM_001347703.2 | XM_021840823.1 |
| NM_001347738.2 | XM_001651818.2 |
| NM_001347828.2 | XR_002502343.1 |
| NM_001347849.2 | XM_021843650.1 |
| NM_001347864.2 | XM_001650929.3 |
| NM_001347893.2 | XM_021852031.1 |

| | |
|---|---|
| NM_001347895.2 | XM_021843646.1 |
| NM_001347898.2 | XR_002501948.1 |
| NM_001347946.2 | XM_021839581.1 |
| NM_001347947.2 | XM_001656413.2 |
| NM_001347967.2 | XM_001652696.2 |
| NM_001347997.2 | XR_002499715.1 |
| NM_001348013.2 | XM_001656592.2 |
| NM_001348129.2 | XM_021847359.1 |
| NM_001348136.2 | XM_021843161.1 |
| NM_001348144.2 | XM_021856693.1 |
| NM_001348189.2 | XM_021844291.1 |
| NM_001348203.1 | XM_021842943.1 |
| NM_001348233.2 | XM_021857032.1 |
| NM_001348239.2 | XR_002499757.1 |
| NM_001348300.1 | XM_021848063.1 |
| NM_001348304.1 | XM_001653085.2 |
| NM_001348322.1 | XM_021839829.1 |
| NM_001348323.3 | XM_001651837.2 |
| NM_001348372.2 | XR_002502833.1 |
| NM_001348397.2 | XR_002499499.1 |
| NM_001348424.1 | XM_021846862.1 |
| NM_001348460.2 | XM_021856757.1 |
| NM_001348466.2 | XM_001656961.2 |
| NM_001348509.2 | XM_021839728.1 |
| NM_001348542.2 | XM_021839727.1 |
| NM_001348547.2 | XM_021857165.1 |
| NM_001348702.2 | XM_001653703.2 |
| NM_001348735.2 | XM_021842616.1 |
| NM_001348756.2 | XR_002499417.1 |
| NM_001348764.2 | XM_021846792.1 |
| NM_001348768.2 | XM_021848486.1 |
| NM_001348781.2 | XM_021843342.1 |
| NM_001348986.1 | XM_021850742.1 |
| NM_001348988.1 | XM_001654139.2 |
| NM_001349012.1 | XR_002502715.1 |
| NM_001349041.2 | XM_021853864.1 |
| NM_001349044.2 | XM_021842723.1 |
| NM_001349099.2 | XM_001648710.2 |
| NM_001349104.2 | XM_001659763.2 |
| NM_001349112.3 | XM_001648209.2 |
| NM_001349149.2 | XM_021854061.1 |
| NM_001349157.2 | XM_001649791.2 |
| NM_001349158.2 | XM_021843315.1 |
| NM_001349168.2 | XM_021842724.1 |
| NM_001349191.2 | XM_021844149.1 |
| NM_001349193.2 | XM_021854559.1 |
| NM_001349222.2 | XM_001660997.2 |
| NM_001349230.2 | XM_001653828.2 |
| NM_001349242.2 | XM_021844278.1 |
| NM_001349258.2 | XM_001651312.2 |

| | |
|---|---|
| NM_001349261.2 | XM_021853810.1 |
| NM_001349263.2 | XM_021839546.1 |
| NM_001349297.2 | XM_021856614.1 |
| NM_001349312.2 | XM_021846109.1 |
| NM_001349320.2 | XM_021842324.1 |
| NM_001349350.2 | XM_021851746.1 |
| NM_001349359.2 | XM_021842725.1 |
| NM_001349362.2 | XM_021840889.1 |
| NM_001349378.2 | XM_001648797.2 |
| NM_001349379.2 | XM_001651301.2 |
| NM_001349383.1 | XM_021847999.1 |
| NM_001349385.1 | XM_021838774.1 |
| NM_001349390.2 | XM_001662630.2 |
| NM_001349394.2 | XM_021855278.1 |
| NM_001349395.2 | XM_021848072.1 |
| NM_001349402.2 | XM_021853615.1 |
| NM_001349413.1 | XM_021849164.1 |
| NM_001349450.1 | XM_021842727.1 |
| NM_001349457.2 | XM_001662758.2 |
| NM_001349460.2 | XM_001655850.2 |
| NM_001349464.2 | XM_021844275.1 |
| NM_001349476.1 | XR_002502130.1 |
| NM_001349479.1 | XM_021847370.1 |
| NM_001349498.1 | XM_021844050.1 |
| NM_001349551.2 | XM_021847679.1 |
| NM_001349623.1 | XM_021843112.1 |
| NM_001349635.1 | XM_021844640.1 |
| NM_001349661.2 | XM_021851429.1 |
| NM_001349711.2 | XM_021838057.1 |
| NM_001349712.2 | XR_002502017.1 |
| NM_001349735.2 | XR_002501375.1 |
| NM_001349743.2 | XR_002499925.1 |
| NM_001349776.2 | XM_021837308.1 |
| NM_001349803.3 | XM_021842290.1 |
| NM_001349853.2 | XR_002498709.1 |
| NM_001349866.2 | XM_021852382.1 |
| NM_001349921.2 | XM_021842528.1 |
| NM_001349935.2 | XM_001659672.2 |
| NM_001349969.2 | XM_021839630.1 |
| NM_001349990.2 | XM_021850740.1 |
| NM_001349991.2 | XM_021837353.1 |
| NM_001350017.2 | XM_021849531.1 |
| NM_001350022.2 | XM_001658963.2 |
| NM_001350043.2 | XM_021850802.1 |
| NM_001350044.2 | XM_021839719.1 |
| NM_001350115.2 | XM_021855951.1 |
| NM_001350123.2 | XM_001650904.2 |
| NM_001350234.2 | XM_021850353.1 |
| NM_001350251.2 | XM_021838545.1 |
| NM_001350262.2 | XM_021844876.1 |

| | |
|---|---|
| NM_001350303.2 | XM_001663217.2 |
| NM_001350310.2 | XM_021851529.1 |
| NM_001350327.2 | XM_021839430.1 |
| NM_001350331.2 | XM_011495002.2 |
| NM_001350394.2 | XM_021848907.1 |
| NM_001350398.2 | XM_021844996.1 |
| NM_001350399.2 | XM_021839717.1 |
| NM_001350402.1 | XM_021844659.1 |
| NM_001350442.2 | XM_001657927.2 |
| NM_001350446.2 | XM_021852355.1 |
| NM_001350456.2 | XM_021848053.1 |
| NM_001350506.2 | XM_001650110.3 |
| NM_001350514.2 | XM_001651706.3 |
| NM_001350565.2 | XM_001654953.2 |
| NM_001350570.2 | XM_001650995.2 |
| NM_001350581.2 | XM_001648775.2 |
| NM_001350602.2 | XM_021852904.1 |
| NM_001350614.2 | XM_001660506.2 |
| NM_001350629.2 | XM_021856308.1 |
| NM_001350635.3 | XM_021848700.1 |
| NM_001350645.2 | XM_001655624.2 |
| NM_001350681.1 | XR_002500903.1 |
| NM_001350714.2 | XM_021851427.1 |
| NM_001350745.2 | XM_021843220.1 |
| NM_001350769.2 | XM_021843031.1 |
| NM_001350781.1 | XM_001655171.2 |
| NM_001350829.2 | XM_001650404.2 |
| NM_001350894.1 | XM_021844617.1 |
| NM_001350898.2 | XM_021854336.1 |
| NM_001350913.2 | XM_021853926.1 |
| NM_001350922.2 | XM_021846189.1 |
| NM_001350936.2 | XM_021838972.1 |
| NM_001350955.2 | XR_002501675.1 |
| NM_001351023.2 | XM_001660792.2 |
| NM_001351044.2 | XM_001658809.2 |
| NM_001351059.2 | XM_001654174.2 |
| NM_001351068.2 | XM_021842500.1 |
| NM_001351076.2 | XM_001660501.2 |
| NM_001351087.2 | XM_021842086.1 |
| NM_001351100.2 | XR_002502162.1 |
| NM_001351114.2 | XM_001660509.2 |
| NM_001351123.2 | XM_001653591.2 |
| NM_001351127.2 | XM_021842707.1 |
| NM_001351136.2 | XM_021845146.1 |
| NM_001351140.2 | XM_001654018.2 |
| NM_001351149.2 | XM_021842469.1 |
| NM_001351163.2 | XM_021841422.1 |
| NM_001351222.2 | XR_002501932.1 |
| NM_001351242.2 | XM_021850906.1 |
| NM_001351243.2 | XM_021843011.1 |

| | |
|---|---|
| NM_001351263.2 | XM_021845065.1 |
| NM_001351266.2 | XM_021842986.1 |
| NM_001351271.2 | XM_021839792.1 |
| NM_001351293.2 | XR_002501991.1 |
| NM_001351302.3 | XM_001654417.2 |
| NM_001351336.2 | XR_002502530.1 |
| NM_001351357.2 | XR_002500622.1 |
| NM_001351367.2 | XM_001651675.2 |
| NM_001351392.2 | XM_001663531.2 |
| NM_001351408.2 | XM_021840373.1 |
| NM_001351418.2 | XM_021845649.1 |
| NM_001351462.2 | XR_002503160.1 |
| NM_001351488.2 | XM_021837733.1 |
| NM_001351509.2 | XM_021837340.1 |
| NM_001351521.2 | XM_021841100.1 |
| NM_001351531.2 | XM_021845564.1 |
| NM_001351532.2 | XM_021846197.1 |
| NM_001351568.3 | XM_021856145.1 |
| NM_001351587.2 | XM_021839722.1 |
| NM_001351589.2 | XM_021844492.1 |
| NM_001351609.2 | XM_021854130.1 |
| NM_001351615.2 | XR_002502867.1 |
| NM_001351616.1 | XM_021838203.1 |
| NM_001351624.2 | XM_021838756.1 |
| NM_001351644.2 | XM_021842087.1 |
| NM_001351835.2 | XR_002502511.1 |
| NM_001351942.2 | XM_021850020.1 |
| NM_001351959.2 | XR_002500813.1 |
| NM_001351970.2 | XM_021845856.1 |
| NM_001352027.3 | XM_021838614.1 |
| NM_001352053.2 | XM_021847406.1 |
| NM_001352057.2 | XR_002500090.1 |
| NM_001352081.2 | XM_021854505.1 |
| NM_001352098.2 | XR_002500900.1 |
| NM_001352102.2 | XM_021848796.1 |
| NM_001352106.2 | XM_021838895.1 |
| NM_001352108.2 | XM_021844406.1 |
| NM_001352139.2 | XM_011495019.2 |
| NM_001352161.2 | XM_001652122.2 |
| NM_001352162.2 | XM_001655974.2 |
| NM_001352173.2 | XM_001656494.2 |
| NM_001352210.2 | XM_021837953.1 |
| NM_001352242.2 | XM_011495049.2 |
| NM_001352308.2 | XM_001653694.2 |
| NM_001352411.2 | XM_021843247.1 |
| NM_001352442.2 | XM_001653285.2 |
| NM_001352498.2 | XM_001653189.2 |
| NM_001352516.2 | XM_021847497.1 |
| NM_001352517.1 | XM_021849736.1 |
| NM_001352529.2 | XM_021842936.1 |

| | |
|---|---|
| NM_001352539.2 | XM_021851333.1 |
| NM_001352542.2 | XR_002499379.1 |
| NM_001352615.2 | XM_021842855.1 |
| NM_001352617.2 | XM_001657204.2 |
| NM_001352618.2 | XM_021845321.1 |
| NM_001352670.2 | XM_001662842.2 |
| NM_001352671.2 | XM_021844444.1 |
| NM_001352684.2 | XR_002501420.1 |
| NM_001352695.2 | XM_001654505.2 |
| NM_001352718.2 | XR_002500176.1 |
| NM_001352750.2 | XR_002501413.1 |
| NM_001352764.2 | XM_021843119.1 |
| NM_001352825.2 | XM_021856178.1 |
| NM_001352826.2 | XM_001648139.3 |
| NM_001352849.2 | XM_021843846.1 |
| NM_001352881.1 | XM_021848305.1 |
| NM_001352910.2 | XR_002502364.1 |
| NM_001352912.1 | XM_021839040.1 |
| NM_001352934.1 | XM_021857729.1 |
| NM_001352949.1 | XR_002502761.1 |
| NM_001352961.1 | XM_021841598.1 |
| NM_001352988.2 | XM_021851071.1 |
| NM_001352995.2 | XM_001654153.2 |
| NM_001353006.2 | XM_021851467.1 |
| NM_001353011.2 | XM_001648229.2 |
| NM_001353018.2 | XM_021843335.1 |
| NM_001353023.2 | XM_021852335.1 |
| NM_001353064.2 | XM_001647784.2 |
| NM_001353075.1 | XM_021853568.1 |
| NM_001353098.2 | XM_001663447.2 |
| NM_001353124.2 | XM_021853046.1 |
| NM_001353154.1 | XM_001656742.2 |
| NM_001353207.1 | XM_001655673.2 |
| NM_001353222.2 | XR_002500817.1 |
| NM_001353229.2 | XM_021842848.1 |
| NM_001353232.1 | XM_021847117.1 |
| NM_001353237.1 | XM_021839944.1 |
| NM_001353248.2 | XM_001651300.2 |
| NM_001353301.2 | XM_021852310.1 |
| NM_001353302.1 | XR_002500770.1 |
| NM_001353313.2 | XM_001656603.2 |
| NM_001353320.2 | XM_021856241.1 |
| NM_001353340.1 | XM_021855736.1 |
| NM_001353486.2 | XM_001663333.2 |
| NM_001353533.2 | XR_002498915.1 |
| NM_001353551.1 | XM_021847768.1 |
| NM_001353602.2 | XR_002498898.1 |
| NM_001353658.2 | XM_021843123.1 |
| NM_001353666.2 | XM_001649999.2 |
| NM_001353676.2 | XM_021850796.1 |

| | |
|---|---|
| NM_001353695.2 | XM_001656215.2 |
| NM_001353715.2 | XM_021847649.1 |
| NM_001353717.2 | XR_002502288.1 |
| NM_001353734.2 | XM_001657501.3 |
| NM_001353743.2 | XM_021851470.1 |
| NM_001353776.2 | XM_021848767.1 |
| NM_001353780.2 | XM_001649372.2 |
| NM_001353824.2 | XM_021856618.1 |
| NM_001353827.2 | XM_021843454.1 |
| NM_001353829.2 | XM_021839025.1 |
| NM_001353837.2 | XM_001657842.2 |
| NM_001353881.2 | XM_021855269.1 |
| NM_001353904.1 | XM_001656094.2 |
| NM_001353924.2 | XM_021850515.1 |
| NM_001353951.2 | XM_021844301.1 |
| NM_001353985.2 | XM_021844256.1 |
| NM_001353994.1 | XM_021848456.1 |
| NM_001354001.3 | XM_001652868.2 |
| NM_001354011.2 | XM_021843569.1 |
| NM_001354024.2 | XM_021851622.1 |
| NM_001354047.1 | XM_021846486.1 |
| NM_001354096.2 | XM_021841646.1 |
| NM_001354173.2 | XR_002500681.1 |
| NM_001354260.2 | XM_001660941.2 |
| NM_001354300.2 | XM_021854252.1 |
| NM_001354315.2 | XM_001653648.2 |
| NM_001354346.2 | XM_021851990.1 |
| NM_001354392.2 | XM_021854761.1 |
| NM_001354397.2 | XM_021852005.1 |
| NM_001354398.2 | XM_021843595.1 |
| NM_001354423.2 | XM_021840022.1 |
| NM_001354424.2 | XM_021838209.1 |
| NM_001354427.2 | XM_021843319.1 |
| NM_001354460.2 | XM_001650621.2 |
| NM_001354473.2 | XR_002500853.1 |
| NM_001354476.2 | XM_021843051.1 |
| NM_001354567.2 | XM_021843524.1 |
| NM_001354579.2 | XM_021840602.1 |
| NM_001354608.2 | XR_002499630.1 |
| NM_001354619.2 | XM_021842728.1 |
| NM_001354640.2 | XM_021843862.1 |
| NM_001354690.3 | XM_021838760.1 |
| NM_001354693.3 | XR_002498926.1 |
| NM_001354698.2 | XM_021846296.1 |
| NM_001354712.2 | XM_021843857.1 |
| NM_001354727.2 | XM_001652893.2 |
| NM_001354751.2 | XM_021839540.1 |
| NM_001354779.2 | XM_001663017.2 |
| NM_001354817.1 | XM_001656310.2 |
| NM_001354932.2 | XR_002500946.1 |

| | |
|---|---|
| NM_001354934.2 | XM_021849153.1 |
| NM_001354974.2 | XM_001656130.2 |
| NM_001355001.2 | XR_002502329.1 |
| NM_001355006.2 | XM_001658603.2 |
| NM_001355276.2 | XM_021851324.1 |
| NM_001355278.1 | XM_021846515.1 |
| NM_001355465.2 | XM_021844428.1 |
| NM_001356372.2 | XM_021843353.1 |
| NM_001357943.2 | XR_002499142.1 |
| NM_001358263.1 | XM_021851188.1 |
| NM_001358345.2 | XM_021852478.1 |
| NM_001359651.2 | XM_021849659.1 |
| NM_001361.5 | XM_021851466.1 |
| NM_001362481.2 | XM_021853750.1 |
| NM_001362777.2 | XM_001651903.2 |
| NM_001362787.2 | XM_021853047.1 |
| NM_001362848.1 | XM_021839668.1 |
| NM_001362872.2 | XM_021851233.1 |
| NM_001362873.2 | XM_001661089.2 |
| NM_001362926.2 | XM_001652894.2 |
| NM_001362965.2 | XM_001650765.3 |
| NM_001362988.2 | XM_021843053.1 |
| NM_001363041.2 | XM_021852605.1 |
| NM_001363046.2 | XR_002499338.1 |
| NM_001363083.1 | XM_021856965.1 |
| NM_001363104.2 | XM_021852623.1 |
| NM_001363138.1 | XM_021852628.1 |
| NM_001363343.2 | XM_021846024.1 |
| NM_001363345.2 | XM_021850774.1 |
| NM_001363366.1 | XR_002502946.1 |
| NM_001363413.2 | XM_021839502.1 |
| NM_001363447.2 | XM_021853797.1 |
| NM_001363460.1 | XM_021844277.1 |
| NM_001363514.2 | XM_001658576.2 |
| NM_001363516.2 | XM_001650095.2 |
| NM_001363518.2 | XM_021849715.1 |
| NM_001363522.2 | XM_001654272.2 |
| NM_001363539.2 | XM_021844412.1 |
| NM_001363543.2 | XM_021844187.1 |
| NM_001363578.1 | XM_021855734.1 |
| NM_001363651.3 | XM_021838001.1 |
| NM_001363655.2 | XM_021841703.1 |
| NM_001363657.3 | XM_001661492.2 |
| NM_001363699.2 | XM_001656055.2 |
| NM_001363707.2 | XM_001659438.2 |
| NM_001363737.2 | XM_021853786.1 |
| NM_001363766.1 | XM_021843571.1 |
| NM_001363794.2 | XM_021848107.1 |
| NM_001363802.1 | XR_002502668.1 |
| NM_001363816.2 | XR_002498912.1 |

| | |
|---|---|
| NM_001363875.2 | XR_002501626.1 |
| NM_001363900.1 | XM_021843452.1 |
| NM_001363914.1 | XM_001651230.2 |
| NM_001363916.1 | XM_021846293.1 |
| NM_001363964.1 | XM_021846105.1 |
| NM_001363988.1 | XM_001653686.2 |
| NM_001364021.1 | XM_001648644.2 |
| NM_001364084.2 | XM_001654240.2 |
| NM_001364159.2 | XR_002498828.1 |
| NM_001364228.2 | XM_021857688.1 |
| NM_001364245.2 | XM_001657215.2 |
| NM_001364255.2 | XM_021846492.1 |
| NM_001364393.2 | XM_001662049.3 |
| NM_001364486.2 | XM_021850339.1 |
| NM_001364520.2 | XM_021844533.1 |
| NM_001364573.2 | XM_021852185.1 |
| NM_001364681.2 | XR_002501627.1 |
| NM_001364682.1 | XM_021846493.1 |
| NM_001364818.2 | XM_021844616.1 |
| NM_001365125.2 | XM_001650949.2 |
| NM_001365156.1 | XM_021847383.1 |
| NM_001365202.1 | XM_021844485.1 |
| NM_001365203.1 | XM_021845596.1 |
| NM_001365204.1 | XR_002500056.1 |
| NM_001365208.1 | XM_021844833.1 |
| NM_001365250.1 | XM_021850039.1 |
| NM_001365252.1 | XM_021838151.1 |
| NM_001365264.1 | XM_021839656.1 |
| NM_001365373.1 | XM_021845473.1 |
| NM_001365408.1 | XM_001656781.2 |
| NM_001365496.2 | XR_002499005.1 |
| NM_001365520.2 | XM_021847364.1 |
| NM_001365598.3 | XM_021853553.1 |
| NM_001365603.3 | XM_021855479.1 |
| NM_001365634.1 | XM_021854471.1 |
| NM_001365713.1 | XM_021842759.1 |
| NM_001365717.1 | XM_001656935.2 |
| NM_001365743.2 | XM_021856639.1 |
| NM_001365768.2 | XM_001650920.2 |
| NM_001365774.2 | XM_021846760.1 |
| NM_001365779.1 | XM_021846556.1 |
| NM_001365805.1 | XM_021856484.1 |
| NM_001365835.2 | XM_021842632.1 |
| NM_001365837.1 | XR_002499775.1 |
| NM_001365855.1 | XM_021850923.1 |
| NM_001365891.2 | XM_021842761.1 |
| NM_001365902.3 | XM_001650881.2 |
| NM_001365951.3 | XM_021856491.1 |
| NM_001366086.1 | XR_002500464.1 |
| NM_001366088.2 | XR_002499596.1 |

| | |
|---|---|
| NM_001366213.1 | XR_002501022.1 |
| NM_001366253.2 | XM_021849692.1 |
| NM_001366283.2 | XM_001655351.2 |
| NM_001366361.1 | XM_001657307.2 |
| NM_001366371.2 | XM_021837711.1 |
| NM_001366473.2 | XM_021856227.1 |
| NM_001366479.2 | XM_001648306.2 |
| NM_001366501.2 | XM_021849766.1 |
| NM_001366532.1 | XM_021845073.1 |
| NM_001366545.2 | XM_001661314.2 |
| NM_001366556.2 | XM_001664108.2 |
| NM_001366570.1 | XM_021850539.1 |
| NM_001366583.1 | XR_002500249.1 |
| NM_001366586.1 | XM_001662998.2 |
| NM_001366618.1 | XM_021843256.1 |
| NM_001366682.2 | XR_002501143.1 |
| NM_001366686.3 | XM_011494703.2 |
| NM_001366779.1 | XM_021839027.1 |
| NM_001366803.2 | XM_021849481.1 |
| NM_001366993.1 | XR_002502386.1 |
| NM_001367064.1 | XM_021852087.1 |
| NM_001367360.2 | XM_021854102.1 |
| NM_001367404.1 | XM_021850677.1 |
| NM_001367424.1 | XM_021840853.1 |
| NM_001367460.1 | XM_021841072.1 |
| NM_001367461.1 | XM_021843908.1 |
| NM_001367462.1 | XR_002500084.1 |
| NM_001367510.1 | XM_001648408.2 |
| NM_001367517.1 | XM_001654349.2 |
| NM_001367561.1 | XM_001653701.2 |
| NM_001367569.1 | XM_021850748.1 |
| NM_001367570.1 | XM_001663625.2 |
| NM_001367600.2 | XM_001661587.2 |
| NM_001367645.1 | XM_001659844.2 |
| NM_001367651.1 | XM_021844369.1 |
| NM_001367691.1 | XM_021851347.1 |
| NM_001367737.1 | XM_001652340.2 |
| NM_001367747.1 | XR_002499400.1 |
| NM_001367753.1 | XR_002500236.1 |
| NM_001367791.1 | XM_021848037.1 |
| NM_001367797.1 | XM_021850734.1 |
| NM_001367810.1 | XM_001648803.2 |
| NM_001367842.1 | XR_002502653.1 |
| NM_001367850.1 | XM_021843253.1 |
| NM_001367851.1 | XM_021842915.1 |
| NM_001367856.1 | XM_021854321.1 |
| NM_001367860.1 | XM_001662654.2 |
| NM_001367872.1 | XM_021855622.1 |
| NM_001367873.1 | XM_021843677.1 |
| NM_001367895.1 | XM_021850701.1 |

| | |
|---|---|
| NM_001367897.1 | XR_002502415.1 |
| NM_001367904.1 | XM_021837318.1 |
| NM_001367915.1 | XM_001651037.2 |
| NM_001367968.1 | XM_001653891.2 |
| NM_001367975.1 | XM_001663743.2 |
| NM_001368066.1 | XM_021846871.1 |
| NM_001368275.1 | XM_021843695.1 |
| NM_001368324.2 | XM_021852433.1 |
| NM_001368888.2 | XR_002500643.1 |
| NM_001368900.2 | XR_002503178.1 |
| NM_001368906.2 | XM_021838793.1 |
| NM_001368921.2 | XM_021845858.1 |
| NM_001369044.1 | XM_021853769.1 |
| NM_001369110.1 | XR_002501817.1 |
| NM_001369256.1 | XM_021856662.1 |
| NM_001369401.1 | XM_021838013.1 |
| NM_001369406.2 | XM_001648419.2 |
| NM_001369468.1 | XM_021852386.1 |
| NM_001369486.1 | XM_021850063.1 |
| NM_001369495.1 | XM_021839609.1 |
| NM_001369512.1 | XM_001662350.2 |
| NM_001369523.1 | XM_021841425.1 |
| NM_001369542.1 | XR_002501512.1 |
| NM_001369551.1 | XM_021850209.1 |
| NM_001369578.1 | XM_021846436.1 |
| NM_001369579.1 | XM_001655075.3 |
| NM_001369580.1 | XM_021845860.1 |
| NM_001369583.1 | XM_021850362.1 |
| NM_001369600.1 | XM_001648795.2 |
| NM_001369639.1 | XM_001661594.3 |
| NM_001369644.1 | XR_002501983.1 |
| NM_001369672.1 | XM_021842880.1 |
| NM_001369703.1 | XM_021839802.1 |
| NM_001369788.1 | XM_001657539.2 |
| NM_001369799.1 | XM_021843387.1 |
| NM_001369821.1 | XM_001650334.2 |
| NM_001369864.1 | XM_001660596.2 |
| NM_001369917.1 | XM_001658114.2 |
| NM_001369920.1 | XM_021844102.1 |
| NM_001369930.1 | XM_021849237.1 |
| NM_001370076.1 | XM_021846020.1 |
| NM_001370085.1 | XM_001655165.2 |
| NM_001370104.2 | XM_021848197.1 |
| NM_001370113.2 | XM_021839563.1 |
| NM_001370126.1 | XM_021848102.1 |
| NM_001370136.1 | XM_021838446.1 |
| NM_001370153.1 | XM_021844449.1 |
| NM_001370167.1 | XR_002498845.1 |
| NM_001370202.1 | XM_021852647.1 |
| NM_001370221.1 | XM_021845637.1 |

| | |
|---|---|
| NM_001370237.1 | XM_021854579.1 |
| NM_001370260.1 | XM_001654324.2 |
| NM_001370261.1 | XM_001654170.2 |
| NM_001370271.1 | XM_021849228.1 |
| NM_001370317.1 | XM_021847650.1 |
| NM_001370348.2 | XM_021853273.1 |
| NM_001370367.1 | XM_021845643.1 |
| NM_001370403.1 | XM_001651395.2 |
| NM_001370426.1 | XM_001650877.2 |
| NM_001370432.1 | XM_001658309.2 |
| NM_001370461.1 | XM_001654871.2 |
| NM_001370467.1 | XR_002500145.1 |
| NM_001370501.1 | XM_021852648.1 |
| NM_001370532.1 | XM_001652650.2 |
| NM_001370533.1 | XM_021839795.1 |
| NM_001370554.1 | XM_001662670.2 |
| NM_001370567.1 | XM_021845138.1 |
| NM_001370588.1 | XM_021847268.1 |
| NM_001370608.1 | XM_021843172.1 |
| NM_001370639.1 | XR_002499865.1 |
| NM_001370649.1 | XM_021837331.1 |
| NM_001370652.1 | XM_021850056.1 |
| NM_001370687.1 | XM_021839188.1 |
| NM_001371009.1 | XM_021845647.1 |
| NM_001371075.1 | XR_002502440.1 |
| NM_001371128.1 | XM_001654718.2 |
| NM_001371137.1 | XM_021849229.1 |
| NM_001371138.1 | XM_021852105.1 |
| NM_001371168.1 | XM_001656037.2 |
| NM_001371206.1 | XM_021839584.1 |
| NM_001371210.1 | XM_021842062.1 |
| NM_001371215.1 | XR_002502634.1 |
| NM_001371216.1 | XM_021857428.1 |
| NM_001371217.1 | XR_002501222.1 |
| NM_001371237.1 | XM_001662175.2 |
| NM_001371251.1 | XM_001655598.2 |
| NM_001371328.1 | XR_002500067.1 |
| NM_001371332.1 | XM_021849473.1 |
| NM_001371333.1 | XM_001653265.2 |
| NM_001371335.1 | XM_021848848.1 |
| NM_001371350.1 | XM_021847029.1 |
| NM_001371355.1 | XM_021844936.1 |
| NM_001371362.1 | XM_001651704.2 |
| NM_001371389.2 | XM_021857206.1 |
| NM_001371410.1 | XM_001663709.2 |
| NM_001371427.1 | XM_021850629.1 |
| NM_001371445.1 | XM_001655806.2 |
| NM_001371452.1 | XM_021843386.1 |
| NM_001371589.1 | XM_021849243.1 |
| NM_001371592.1 | XM_021845570.1 |

| | |
|---|---|
| NM_001371740.1 | XR_002501674.1 |
| NM_001371774.1 | XR_002499206.1 |
| NM_001371858.3 | XM_021852203.1 |
| NM_001371923.1 | XM_001650831.2 |
| NM_001371936.1 | XM_021854438.1 |
| NM_001372024.1 | XM_021853567.1 |
| NM_001372025.1 | XM_001648677.3 |
| NM_001372070.1 | XM_021854519.1 |
| NM_001372080.1 | XM_001663110.2 |
| NM_001372167.1 | XM_021838447.1 |
| NM_001372189.1 | XM_001660365.2 |
| NM_001374256.1 | XM_021837943.1 |
| NM_001374264.2 | XM_001649766.2 |
| NM_001374269.1 | XM_021846715.1 |
| NM_001374357.1 | XM_001652321.2 |
| NM_001374523.1 | XM_021851570.1 |
| NM_001374596.1 | XR_002502950.1 |
| NM_001374781.1 | XM_021845661.1 |
| NM_001374824.1 | XM_021854627.1 |
| NM_001374847.1 | XM_021843465.1 |
| NM_001374868.1 | XR_002499086.1 |
| NM_001375286.1 | XM_021845620.1 |
| NM_001375291.1 | XM_021850389.1 |
| NM_001375296.1 | XM_021857204.1 |
| NM_001375299.1 | XM_001661561.2 |
| NM_001375321.1 | XM_021843986.1 |
| NM_001375330.1 | XR_002498802.1 |
| NM_001375334.1 | XM_021840130.1 |
| NM_001375353.1 | XR_002502827.1 |
| NM_001375360.1 | XM_021843373.1 |
| NM_001375411.1 | XR_002499265.1 |
| NM_001375444.1 | XM_001654388.2 |
| NM_001375463.1 | XR_002498914.1 |
| NM_001375467.1 | XM_001648892.3 |
| NM_001375468.1 | XM_021848560.1 |
| NM_001375477.1 | XM_021848031.1 |
| NM_001375491.1 | XM_001655100.3 |
| NM_001375492.1 | XM_021850891.1 |
| NM_001375554.1 | XM_021843618.1 |
| NM_001375575.1 | XM_001658031.2 |
| NM_001375583.1 | XM_021844279.1 |
| NM_001375585.1 | XM_021852679.1 |
| NM_001375589.2 | XM_001660720.2 |
| NM_001375618.1 | XM_001659751.2 |
| NM_001375630.1 | XM_001648227.2 |
| NM_001375651.1 | XM_021856517.1 |
| NM_001375655.1 | XM_021846590.1 |
| NM_001375673.1 | XM_001663136.2 |
| NM_001375697.1 | XM_021850442.1 |
| NM_001375705.1 | XR_002500702.1 |

| | |
|---|---|
| NM_001375720.1 | XM_021851346.1 |
| NM_001375725.1 | XM_001649342.2 |
| NM_001375741.1 | XM_021849762.1 |
| NM_001375756.1 | XM_001651334.2 |
| NM_001375780.1 | XR_002500555.1 |
| NM_001375845.1 | XM_001648782.2 |
| NM_001375897.1 | XM_001654325.2 |
| NM_001375908.1 | XM_021853373.1 |
| NM_001375939.1 | XM_021843336.1 |
| NM_001375968.1 | XM_001659499.2 |
| NM_001376068.1 | XM_021847145.1 |
| NM_001376085.1 | XR_002500039.1 |
| NM_001376112.1 | XM_021847010.1 |
| NM_001376119.1 | XM_011495451.2 |
| NM_001376131.1 | XM_001655795.2 |
| NM_001376132.1 | XM_021843453.1 |
| NM_001376150.1 | XM_021856870.1 |
| NM_001376153.1 | XM_001652759.2 |
| NM_001376157.1 | XM_021856852.1 |
| NM_001376161.1 | XM_021843693.1 |
| NM_001376248.1 | XM_021845124.1 |
| NM_001376251.1 | XM_021848744.1 |
| NM_001376253.1 | XM_021840576.1 |
| NM_001376323.2 | XM_021848252.1 |
| NM_001376342.1 | XM_021838107.1 |
| NM_001376351.1 | XM_001660504.2 |
| NM_001376354.1 | XM_021844704.1 |
| NM_001376364.1 | XR_002501247.1 |
| NM_001376379.1 | XR_002501660.1 |
| NM_001376443.1 | XR_002500388.1 |
| NM_001376446.1 | XM_021848610.1 |
| NM_001376450.1 | XM_021849257.1 |
| NM_001376483.1 | XM_021850383.1 |
| NM_001376523.1 | XR_002501014.1 |
| NM_001376529.1 | XM_001649558.2 |
| NM_001376543.1 | XR_002503139.1 |
| NM_001376580.1 | XM_021848704.1 |
| NM_001376591.1 | XM_021837390.1 |
| NM_001376616.1 | XM_021843756.1 |
| NM_001376619.1 | XM_021855577.1 |
| NM_001376625.1 | XR_002499947.1 |
| NM_001376639.1 | XM_021839197.1 |
| NM_001376654.1 | XR_002502770.1 |
| NM_001376682.1 | XM_021844427.1 |
| NM_001376690.1 | XM_021852256.1 |
| NM_001376696.1 | XM_021839593.1 |
| NM_001376725.1 | XM_021843839.1 |
| NM_001376726.1 | XM_021840825.1 |
| NM_001376728.1 | XM_001648038.3 |
| NM_001376783.1 | XM_021850635.1 |

| | |
|---|---|
| NM_001376793.1 | XM_001652352.2 |
| NM_001376817.1 | XM_021848030.1 |
| NM_001376821.1 | XM_021856853.1 |
| NM_001376844.1 | XM_021847998.1 |
| NM_001376887.1 | XR_002500317.1 |
| NM_001376932.1 | XM_021843326.1 |
| NM_001377062.1 | XR_002499339.1 |
| NM_001377129.1 | XM_021842066.1 |
| NM_001377130.1 | XM_001662711.2 |
| NM_001377173.1 | XM_001652497.2 |
| NM_001377174.1 | XM_001653655.2 |
| NM_001377178.1 | XM_021846386.1 |
| NM_001377198.1 | XM_021841419.1 |
| NM_001377203.1 | XM_001662420.2 |
| NM_001377204.1 | XM_001658588.2 |
| NM_001377208.1 | XM_001659727.2 |
| NM_001377212.1 | XM_001649902.2 |
| NM_001377228.1 | XM_021848253.1 |
| NM_001377238.1 | XM_001648586.2 |
| NM_001377291.1 | XM_001652734.2 |
| NM_001377317.2 | XM_021837654.1 |
| NM_001377358.1 | XM_001659414.2 |
| NM_001377362.1 | XM_001652002.2 |
| NM_001377377.1 | XR_002500921.1 |
| NM_001377379.1 | XR_002503191.1 |
| NM_001377394.1 | XM_021847972.1 |
| NM_001377405.1 | XM_021844177.1 |
| NM_001377408.1 | XM_001651067.2 |
| NM_001377411.1 | XM_001651278.2 |
| NM_001377418.1 | XM_021844350.1 |
| NM_001377427.1 | XM_001659705.2 |
| NM_001377470.1 | XM_001659534.2 |
| NM_001377491.1 | XM_021849191.1 |
| NM_001377492.1 | XM_021849226.1 |
| NM_001377496.1 | XR_002498596.1 |
| NM_001377931.1 | XR_002503221.1 |
| NM_001377961.1 | XM_021839381.1 |
| NM_001377970.1 | XM_001659407.2 |
| NM_001377974.1 | XM_001659597.2 |
| NM_001378029.1 | XM_021851874.1 |
| NM_001378034.1 | XM_001659623.2 |
| NM_001378074.1 | XM_021842607.1 |
| NM_001378077.1 | XM_021844432.1 |
| NM_001378092.1 | XM_021846584.1 |
| NM_001378094.1 | XM_021838159.1 |
| NM_001378102.1 | XM_021839947.1 |
| NM_001378107.1 | XM_021846615.1 |
| NM_001378132.1 | XM_021844818.1 |
| NM_001378144.1 | XM_021854439.1 |
| NM_001378152.1 | XM_021840035.1 |

| | |
|---|---|
| NM_001378157.1 | XM_001659263.2 |
| NM_001378178.1 | XM_021851787.1 |
| NM_001378181.1 | XM_021848237.1 |
| NM_001378254.1 | XM_021851747.1 |
| NM_001378257.1 | XR_002502338.1 |
| NM_001378270.1 | XM_021847213.1 |
| NM_001378408.1 | XM_021855655.1 |
| NM_001378456.1 | XM_021840906.1 |
| NM_001378485.1 | XR_002498653.1 |
| NM_001378486.1 | XR_002499682.1 |
| NM_001378533.1 | XM_021848562.1 |
| NM_001378538.1 | XM_021854763.1 |
| NM_001378546.1 | XM_021838015.1 |
| NM_001378851.1 | XM_001651595.2 |
| NM_001379104.1 | XR_002500928.1 |
| NM_001379331.1 | XM_021849474.1 |
| NM_001379461.1 | XM_021850567.1 |
| NM_001379515.1 | XM_001649607.2 |
| NM_001379551.1 | XM_001651347.2 |
| NM_001379566.1 | XR_002499839.1 |
| NM_001379610.1 | XM_021853753.1 |
| NM_001379614.1 | XM_021850839.1 |
| NM_001381865.2 | XM_021845566.1 |
| NM_001381886.1 | XR_002498655.1 |
| NM_001381939.1 | XM_021844174.1 |
| NM_001381941.1 | XM_021854798.1 |
| NM_001381949.1 | XR_002498677.1 |
| NM_001382242.1 | XR_002502961.1 |
| NM_001382287.1 | XM_021843352.1 |
| NM_001382288.1 | XM_021854799.1 |
| NM_001382310.1 | XM_001651499.2 |
| NM_001382339.1 | XM_021844803.1 |
| NM_001382396.1 | XM_001654171.2 |
| NM_001382408.1 | XM_021840590.1 |
| NM_001382424.1 | XM_021850233.1 |
| NM_001382429.1 | XM_001656688.2 |
| NM_001382467.1 | XM_021850680.1 |
| NM_001382470.1 | XR_002503204.1 |
| NM_001382527.1 | XM_001663502.2 |
| NM_001382528.1 | XM_021854444.1 |
| NM_001382554.1 | XM_021855572.1 |
| NM_001382573.1 | XR_002503177.1 |
| NM_001382611.1 | XM_001653206.2 |
| NM_001382616.1 | XM_021857284.1 |
| NM_001382619.1 | XM_021855728.1 |
| NM_001382628.1 | XM_021852373.1 |
| NM_001382634.1 | XM_021855018.1 |
| NM_001382639.1 | XM_021841708.1 |
| NM_001382645.1 | XM_021839504.1 |
| NM_001382652.1 | XM_021849392.1 |

| | |
|---|---|
| NM_001382674.1 | XM_001654441.2 |
| NM_001382696.1 | XM_021850385.1 |
| NM_001382707.1 | XM_021839246.1 |
| NM_001382722.1 | XM_021843173.1 |
| NM_001382730.1 | XR_002499063.1 |
| NM_001382740.1 | XM_001653663.2 |
| NM_001382767.1 | XM_001660717.2 |
| NM_001382787.1 | XM_021842604.1 |
| NM_001384133.1 | XM_021844614.1 |
| NM_001384268.1 | XM_021852516.1 |
| NM_001384339.1 | XM_021838841.1 |
| NM_001384350.1 | XM_021843356.1 |
| NM_001384356.1 | XM_021847118.1 |
| NM_001384369.1 | XM_001654856.2 |
| NM_001384387.1 | XM_021842051.1 |
| NM_001384390.1 | XM_001656810.2 |
| NM_001384401.1 | XM_021843234.1 |
| NM_001384404.1 | XM_001653623.3 |
| NM_001384414.1 | XR_002501132.1 |
| NM_001384420.1 | XM_021851901.1 |
| NM_001384445.1 | XR_002501148.1 |
| NM_001384459.1 | XM_001653684.2 |
| NM_001384464.1 | XR_002502133.1 |
| NM_001384488.1 | XM_021851810.1 |
| NM_001384508.1 | XR_002499449.1 |
| NM_001384513.1 | XM_001655688.2 |
| NM_001384526.1 | XM_021842760.1 |
| NM_001384534.1 | XM_021843694.1 |
| NM_001384537.1 | XM_021842021.1 |
| NM_001384538.1 | XR_002503163.1 |
| NM_001384554.1 | XM_001660818.2 |
| NM_001384572.1 | XM_001651802.3 |
| NM_001384576.1 | XM_021851876.1 |
| NM_001384582.1 | XM_021849161.1 |
| NM_001384682.1 | XM_001660729.2 |
| NM_001384704.1 | XM_021849343.1 |
| NM_001384707.1 | XR_002501137.1 |
| NM_001384708.1 | XM_021845005.1 |
| NM_001384713.1 | XM_001655480.2 |
| NM_001384734.1 | XR_002502748.1 |
| NM_001384745.1 | XM_001660649.2 |
| NM_001384778.1 | XM_021850395.1 |
| NM_001384809.1 | XR_002500984.1 |
| NM_001384851.1 | XM_001658593.3 |
| NM_001384866.1 | XM_001657780.2 |
| NM_001384867.1 | XM_021847435.1 |
| NM_001384882.1 | XM_021840391.1 |
| NM_001384892.1 | XM_021845253.1 |
| NM_001384898.1 | XM_021853833.1 |
| NM_001384900.1 | XR_002501215.1 |

| | |
|---|---|
| NM_001384936.1 | XM_021852649.1 |
| NM_001384988.1 | XM_021848032.1 |
| NM_001384993.1 | XM_021850426.1 |
| NM_001384995.1 | XM_001660837.2 |
| NM_001385010.1 | XM_021843475.1 |
| NM_001385017.1 | XR_002499787.1 |
| NM_001385028.1 | XM_021837825.1 |
| NM_001385029.1 | XM_001648655.2 |
| NM_001385072.1 | XM_021842100.1 |
| NM_001385084.1 | XR_002502470.1 |
| NM_001385086.1 | XM_021840732.1 |
| NM_001385093.1 | XM_021842902.1 |
| NM_001385110.1 | XM_001661245.2 |
| NM_001385111.1 | XM_021855180.1 |
| NM_001385116.1 | XR_002503216.1 |
| NM_001385146.1 | XM_021852476.1 |
| NM_001385177.1 | XM_021857155.1 |
| NM_001385195.1 | XM_021856159.1 |
| NM_001385217.1 | XM_001650214.2 |
| NM_001385218.1 | XM_021853681.1 |
| NM_001385245.1 | XM_021844103.1 |
| NM_001385251.1 | XM_001656041.2 |
| NM_001385258.1 | XM_021839589.1 |
| NM_001385259.1 | XM_021844411.1 |
| NM_001385274.1 | XM_001658056.2 |
| NM_001385292.1 | XM_001657034.2 |
| NM_001385321.1 | XR_002501773.1 |
| NM_001385341.1 | XR_002501794.1 |
| NM_001385348.1 | XM_021856523.1 |
| NM_001385351.1 | XM_021840915.1 |
| NM_001385354.1 | XM_021853148.1 |
| NM_001385361.1 | XM_001654123.2 |
| NM_001385368.1 | XR_002500172.1 |
| NM_001385374.1 | XM_021853678.1 |
| NM_001385426.1 | XM_021849975.1 |
| NM_001385429.1 | XM_021855692.1 |
| NM_001385444.1 | XM_021848429.1 |
| NM_001385463.1 | XM_001660758.2 |
| NM_001385478.1 | XM_021848225.1 |
| NM_001385491.1 | XM_001652606.2 |
| NM_001385509.1 | XM_021846156.1 |
| NM_001385531.1 | XM_021851186.1 |
| NM_001385551.1 | XM_021848743.1 |
| NM_001385558.1 | XM_011495301.2 |
| NM_001385618.1 | XM_021854227.1 |
| NM_001385621.1 | XM_021851399.1 |
| NM_001385673.1 | XM_001661400.2 |
| NM_001385676.1 | XR_002499119.1 |
| NM_001385689.1 | XM_001663611.2 |
| NM_001385693.1 | XM_021847437.1 |

| | |
|---|---|
| NM_001385703.1 | XM_021847520.1 |
| NM_001385705.1 | XM_001663921.2 |
| NM_001385730.1 | XM_001653240.2 |
| NM_001385735.1 | XM_021838211.1 |
| NM_001385748.1 | XM_021840261.1 |
| NM_001385757.1 | XR_002501515.1 |
| NM_001385769.1 | XM_001651775.2 |
| NM_001385778.1 | XM_021852847.1 |
| NM_001385784.1 | XM_021850994.1 |
| NM_001385792.1 | XM_001656277.2 |
| NM_001385809.1 | XM_021850441.1 |
| NM_001385812.1 | XM_021848688.1 |
| NM_001385821.1 | XM_021848322.1 |
| NM_001385883.1 | XM_001653985.2 |
| NM_001385905.1 | XM_021854498.1 |
| NM_001385913.1 | XM_021850354.1 |
| NM_001385947.1 | XM_021841963.1 |
| NM_001385967.1 | XR_002500955.1 |
| NM_001385997.1 | XM_021842236.1 |
| NM_001386004.1 | XM_021843753.1 |
| NM_001386015.1 | XM_021852670.1 |
| NM_001386019.1 | XM_021844992.1 |
| NM_001386027.1 | XM_001653727.2 |
| NM_001386032.1 | XM_021854142.1 |
| NM_001386084.1 | XM_001648445.2 |
| NM_001386095.1 | XR_002500627.1 |
| NM_001386139.1 | XM_021854370.1 |
| NM_001386141.1 | XM_021844178.1 |
| NM_001386146.1 | XM_021846184.1 |
| NM_001386191.1 | XM_021850175.1 |
| NM_001386193.1 | XM_001652150.2 |
| NM_001386198.1 | XM_001659354.2 |
| NM_001386208.1 | XR_002502887.1 |
| NM_001386290.1 | XM_021844813.1 |
| NM_001386425.1 | XM_021846895.1 |
| NM_001386430.1 | XM_001652293.2 |
| NM_001386441.1 | XM_021855978.1 |
| NM_001386462.1 | XR_002498737.1 |
| NM_001386585.1 | XM_021851144.1 |
| NM_001386608.1 | XM_001663038.2 |
| NM_001386609.1 | XM_001655895.2 |
| NM_001386677.1 | XM_021839662.1 |
| NM_001386682.1 | XM_001661866.2 |
| NM_001386793.1 | XM_021854022.1 |
| NM_001386901.1 | XM_001652130.2 |
| NM_001386929.1 | XM_021845599.1 |
| NM_001386934.1 | XM_021841637.1 |
| NM_001386968.1 | XM_021857440.1 |
| NM_001386978.1 | XR_002498738.1 |
| NM_001387002.1 | XM_021842336.1 |

| | |
|---|---|
| NM_001387009.1 | XM_021846142.1 |
| NM_001387053.1 | XM_021844990.1 |
| NM_001387063.1 | XM_021844988.1 |
| NM_001387067.1 | XR_002502188.1 |
| NM_001387096.1 | XM_021838669.1 |
| NM_001387097.1 | XM_001653912.2 |
| NM_001387102.1 | XM_001664122.2 |
| NM_001387112.1 | XM_021844179.1 |
| NM_001387119.1 | XR_002500539.1 |
| NM_001387138.1 | XM_021848751.1 |
| NM_001387195.1 | XR_002503263.1 |
| NM_001387206.1 | XM_021854358.1 |
| NM_001387230.1 | XM_021851148.1 |
| NM_001387260.1 | XM_001649850.2 |
| NM_001387272.1 | XM_021841961.1 |
| NM_001387302.1 | XM_021852846.1 |
| NM_001387303.1 | XM_021847018.1 |
| NM_001387308.1 | XM_001651473.2 |
| NM_001387317.1 | XM_021850806.1 |
| NM_001387318.1 | XR_002499340.1 |
| NM_001387320.1 | XM_001657890.2 |
| NM_001387347.1 | XM_021844334.1 |
| NM_001387379.1 | XM_001655370.2 |
| NM_001387401.1 | XM_021843707.1 |
| NM_001387412.1 | XM_001654136.2 |
| NM_001387418.1 | XM_021852068.1 |
| NM_001387446.1 | XM_021848111.1 |
| NM_001387481.1 | XM_001654210.2 |
| NM_001387490.1 | XM_001652234.2 |
| NM_001387501.1 | XM_021845022.1 |
| NM_001387505.1 | XM_001651929.2 |
| NM_001387526.1 | XM_001648519.2 |
| NM_001387562.1 | XM_001661204.2 |
| NM_001387569.1 | XM_021844046.1 |
| NM_001387598.1 | XM_021855139.1 |
| NM_001387623.1 | XM_021854125.1 |
| NM_001387650.1 | XM_021837914.1 |
| NM_001387652.1 | XR_002502150.1 |
| NM_001387663.1 | XM_021850328.1 |
| NM_001387697.1 | XM_021853606.1 |
| NM_001387762.1 | XM_021849693.1 |
| NM_001387777.1 | XM_021853455.1 |
| NM_001387780.1 | XM_021842342.1 |
| NM_001387794.1 | XR_002500740.1 |
| NM_001387813.1 | XR_002501271.1 |
| NM_001387821.1 | XR_002499261.1 |
| NM_001387856.1 | XM_001659225.2 |
| NM_001387862.1 | XM_021837921.1 |
| NM_001387919.1 | XM_021842739.1 |
| NM_001387940.1 | XM_021857104.1 |

| | |
|---|---|
| NM_001387960.1 | XM_001649090.2 |
| NM_001387968.1 | XM_021852093.1 |
| NM_001387971.1 | XM_021857459.1 |
| NM_001387976.1 | XM_021838767.1 |
| NM_001387984.1 | XM_021854357.1 |
| NM_001388001.1 | XM_021844101.1 |
| NM_001388009.1 | XR_002500098.1 |
| NM_001388013.1 | XM_011495296.2 |
| NM_001388042.1 | XM_021840327.1 |
| NM_001388043.1 | XM_001657189.2 |
| NM_001388083.1 | XM_021857315.1 |
| NM_001388091.1 | XM_021857501.1 |
| NM_001388109.1 | XM_021854027.1 |
| NM_001388111.1 | XM_021849297.1 |
| NM_001388118.1 | XM_001654951.2 |
| NM_001388127.1 | XM_001655001.2 |
| NM_001388133.1 | XM_021846235.1 |
| NM_001388137.1 | XM_021849851.1 |
| NM_001388151.1 | XM_021855689.1 |
| NM_001388165.1 | XM_021838213.1 |
| NM_001388167.1 | XR_002501070.1 |
| NM_001388171.1 | XM_021847517.1 |
| NM_001388188.1 | XM_001655954.2 |
| NM_001388191.1 | XM_021857280.1 |
| NM_001388200.1 | XM_021854554.1 |
| NM_001388215.1 | XM_001651906.2 |
| NM_001388223.1 | XM_021855694.1 |
| NM_001388235.1 | XM_021848314.1 |
| NM_001388256.1 | XM_021845477.1 |
| NM_001388265.1 | XM_021841116.1 |
| NM_001388269.1 | XM_021840416.1 |
| NM_001388285.1 | XM_001650260.2 |
| NM_001388288.1 | XR_002502722.1 |
| NM_001388300.1 | XM_001660397.2 |
| NM_001388306.1 | XR_002500575.1 |
| NM_001388358.1 | XM_021847556.1 |
| NM_001388380.1 | XM_001654726.2 |
| NM_001388391.1 | XM_001657155.3 |
| NM_001388401.1 | XM_021845006.1 |
| NM_001388410.1 | XM_021840098.1 |
| NM_001388427.1 | XM_021841319.1 |
| NM_001388496.1 | XM_021843527.1 |
| NM_001396.5 | XM_021849152.1 |
| NM_001399.5 | XR_002502744.1 |
| NM_001405.4 | XM_001653709.2 |
| NM_001458.5 | XM_021856679.1 |
| NM_001464.5 | XR_002501427.1 |
| NM_001487.3 | XM_021846537.1 |
| NM_001521.4 | XM_021856274.1 |
| NM_001535.5 | XM_021854631.1 |

| | |
|---|---|
| NM_001545.3 | XM_001663633.2 |
| NM_001546.4 | XM_001659878.2 |
| NM_001556.3 | XM_021857643.1 |
| NM_001605.3 | XM_001661756.2 |
| NM_001618.4 | XM_001656424.2 |
| NM_001619.5 | XR_002500970.1 |
| NM_001640.4 | XM_021850903.1 |
| NM_001669.4 | XR_002501098.1 |
| NM_001674.4 | XM_021839941.1 |
| NM_001718.6 | XM_021844978.1 |
| NM_001729.4 | XM_001648346.2 |
| NM_001731.3 | XM_001661212.2 |
| NM_001741.3 | XM_001658813.2 |
| NM_001745.4 | XM_021845038.1 |
| NM_001834.5 | XM_001660393.2 |
| NM_001858.6 | XM_021837698.1 |
| NM_001876.4 | XM_001651070.2 |
| NM_001895.4 | XM_021847655.1 |
| NM_001939.3 | XM_021842284.1 |
| NM_001951.4 | XM_001649124.2 |
| NM_001966.4 | XM_001648252.2 |
| NM_001988.4 | XM_021850926.1 |
| NM_001992.5 | XM_021844020.1 |
| NM_001997.5 | XM_001659901.2 |
| NM_002040.4 | XM_001662483.2 |
| NM_002086.5 | XM_021842560.1 |
| NM_002092.4 | XM_021857105.1 |
| NM_002122.5 | XM_021840100.1 |
| NM_002127.6 | XM_021849661.1 |
| NM_002188.3 | XM_021849980.1 |
| NM_002235.4 | XM_021849483.1 |
| NM_002241.5 | XM_021844941.1 |
| NM_002258.3 | XM_021848315.1 |
| NM_002262.5 | XM_021854835.1 |
| NM_002264.4 | XM_021848822.1 |
| NM_002279.5 | XM_001648840.2 |
| NM_002317.7 | XM_001651584.3 |
| NM_002346.3 | XM_001647885.2 |
| NM_002348.4 | XM_021849274.1 |
| NM_002360.4 | XM_021844335.1 |
| NM_002362.4 | XM_001658173.2 |
| NM_002376.7 | XM_021846037.1 |
| NM_002383.4 | XM_021850461.1 |
| NM_002390.6 | XM_011494854.2 |
| NM_002409.5 | XM_001655197.2 |
| NM_002420.6 | XM_021852786.1 |
| NM_002441.5 | XM_021848049.1 |
| NM_002446.4 | XM_021845929.1 |
| NM_002449.5 | XM_021848547.1 |
| NM_002470.4 | XM_021844974.1 |

| | |
|---|---|
| NM_002477.1 | XR_002502584.1 |
| NM_002478.5 | XR_002499986.1 |
| NM_002483.7 | XM_021843089.1 |
| NM_002514.4 | XM_021840211.1 |
| NM_002522.4 | XR_002500712.1 |
| NM_002525.3 | XM_021839940.1 |
| NM_002537.3 | XM_021854258.1 |
| NM_002597.5 | XM_001647887.2 |
| NM_002599.5 | XM_021848050.1 |
| NM_002612.4 | XM_021839752.1 |
| NM_002622.5 | XR_002501953.1 |
| NM_002638.4 | XM_021849001.1 |
| NM_002653.5 | XM_001648946.2 |
| NM_002666.5 | XM_021848036.1 |
| NM_002674.4 | XM_021838507.1 |
| NM_002691.4 | XM_021852291.1 |
| NM_002715.4 | XM_021855122.1 |
| NM_002783.3 | XM_021843166.1 |
| NM_002795.4 | XM_021839789.1 |
| NM_002803.4 | XM_001655221.2 |
| NM_002813.7 | XR_002498905.1 |
| NM_002841.4 | XM_021855690.1 |
| NM_002853.4 | XM_021840213.1 |
| NM_002885.4 | XM_001663117.2 |
| NM_002951.5 | XM_001658160.2 |
| NM_002956.3 | XM_011495461.2 |
| NM_003035.2 | XM_021843174.1 |
| NM_003053.4 | XM_021856963.1 |
| NM_003096.4 | XM_021848526.1 |
| NM_003124.5 | XR_002501917.1 |
| NM_003141.4 | XM_021838160.1 |
| NM_003149.3 | XM_001657791.2 |
| NM_003155.3 | XM_021840234.1 |
| NM_003171.5 | XM_021856061.1 |
| NM_003203.5 | XM_001662026.2 |
| NM_003238.6 | XM_021845255.1 |
| NM_003250.6 | XM_021853464.1 |
| NM_003254.3 | XM_021855511.1 |
| NM_003265.3 | XM_011494866.2 |
| NM_003274.5 | XM_001663294.2 |
| NM_003278.3 | XM_001655092.2 |
| NM_003313.4 | XM_021850166.1 |
| NM_003319.4 | XM_021840380.1 |
| NM_003324.5 | XM_021850927.1 |
| NM_003331.5 | XM_021842777.1 |
| NM_003347.4 | XM_021845688.1 |
| NM_003377.5 | XM_001663929.2 |
| NM_003392.7 | XM_021837287.1 |
| NM_003407.5 | XM_021837257.1 |
| NM_003409.5 | XM_021846026.1 |

| | |
|---|---|
| NM_003445.4 | XM_021845090.1 |
| NM_003446.4 | XM_001648386.2 |
| NM_003490.4 | XM_001648254.2 |
| NM_003503.4 | XR_002502127.1 |
| NM_003504.5 | XM_001658214.2 |
| NM_003510.3 | XM_001654682.3 |
| NM_003514.2 | XM_021844520.1 |
| NM_003569.3 | XM_001648152.2 |
| NM_003610.4 | XM_021852929.1 |
| NM_003617.4 | XM_021849976.1 |
| NM_003633.4 | XM_001656237.2 |
| NM_003639.4 | XM_021843938.1 |
| NM_003645.4 | XM_021842793.1 |
| NM_003654.6 | XR_002502045.1 |
| NM_003670.3 | XM_021854445.1 |
| NM_003702.4 | XM_021854748.1 |
| NM_003710.4 | XM_021857290.1 |
| NM_003735.3 | XM_021847937.1 |
| NM_003799.3 | XM_021846351.1 |
| NM_003812.4 | XM_021847944.1 |
| NM_003854.4 | XM_021848375.1 |
| NM_003884.5 | XM_021852876.1 |
| NM_003907.3 | XR_002499941.1 |
| NM_003948.5 | XM_001648247.2 |
| NM_003959.3 | XM_021844104.1 |
| NM_003963.3 | XM_021857521.1 |
| NM_003974.4 | XM_021853352.1 |
| NM_003981.4 | XR_002502802.1 |
| NM_003989.5 | XM_021848388.1 |
| NM_003992.5 | XM_001655130.2 |
| NM_004000.3 | XM_001649911.2 |
| NM_004010.3 | XM_001662529.2 |
| NM_004039.3 | XM_021851683.1 |
| NM_004041.5 | XM_021851941.1 |
| NM_004052.3 | XM_021856774.1 |
| NM_004061.5 | XR_002500429.1 |
| NM_004091.4 | XM_021854834.1 |
| NM_004094.5 | XM_021857647.1 |
| NM_004111.6 | XM_021847938.1 |
| NM_004112.4 | XM_021855252.1 |
| NM_004137.4 | XR_002499494.1 |
| NM_004145.4 | XM_001662763.2 |
| NM_004196.7 | XM_021854247.1 |
| NM_004197.2 | XM_021854790.1 |
| NM_004206.4 | XM_001652376.2 |
| NM_004208.4 | XM_021850402.1 |
| NM_004213.5 | XM_021845847.1 |
| NM_004214.5 | XM_021839510.1 |
| NM_004236.4 | XM_021844626.1 |
| NM_004246.3 | XM_011495283.2 |

| | |
|---|---|
| NM_004254.4 | XM_021845230.1 |
| NM_004257.6 | XM_001655560.2 |
| NM_004279.3 | XM_021848446.1 |
| NM_004316.4 | XM_021845577.1 |
| NM_004326.4 | XM_021841197.1 |
| NM_004354.3 | XM_021847003.1 |
| NM_004359.2 | XM_021848447.1 |
| NM_004362.3 | XM_001663682.2 |
| NM_004403.3 | XM_021851371.1 |
| NM_004418.4 | XR_002502992.1 |
| NM_004448.4 | XM_021853648.1 |
| NM_004452.4 | XM_021840099.1 |
| NM_004459.7 | XM_021855695.1 |
| NM_004492.3 | XM_021842026.1 |
| NM_004507.4 | XM_021843761.1 |
| NM_004508.4 | XM_001653075.2 |
| NM_004524.3 | XM_021854789.1 |
| NM_004532.6 | XM_021842931.1 |
| NM_004535.3 | XR_002500139.1 |
| NM_004544.4 | XM_001656305.2 |
| NM_004606.5 | XM_021848935.1 |
| NM_004660.5 | XR_002499027.1 |
| NM_004663.5 | XR_002501645.1 |
| NM_004688.3 | XM_021848567.1 |
| NM_004690.4 | XM_001654020.2 |
| NM_004720.7 | XM_001652913.2 |
| NM_004764.5 | XM_021857441.1 |
| NM_004769.4 | XM_021853587.1 |
| NM_004821.3 | XM_021843522.1 |
| NM_004840.3 | XM_021844250.1 |
| NM_004854.5 | XM_021850697.1 |
| NM_004861.3 | XM_021855578.1 |
| NM_004863.4 | XM_001652970.2 |
| NM_004896.5 | XM_021856581.1 |
| NM_004901.5 | XM_021855646.1 |
| NM_004902.4 | XM_021840293.1 |
| NM_004904.4 | XM_021847635.1 |
| NM_004942.4 | XM_001653335.2 |
| NM_004945.4 | XM_021855870.1 |
| NM_004953.5 | XM_021844293.1 |
| NM_004959.5 | XR_002499163.1 |
| NM_004972.4 | XR_002498866.1 |
| NM_004981.2 | XM_021855618.1 |
| NM_004994.3 | XM_001654003.2 |
| NM_005014.3 | XM_021837341.1 |
| NM_005020.5 | XM_001656408.2 |
| NM_005028.5 | XM_021849534.1 |
| NM_005032.7 | XM_001654543.2 |
| NM_005045.4 | XM_001659155.2 |
| NM_005052.3 | XR_002502414.1 |

| | |
|---|---|
| NM_005061.3 | XM_021852058.1 |
| NM_005103.5 | XM_021855516.1 |
| NM_005131.3 | XM_021839629.1 |
| NM_005156.7 | XM_021845224.1 |
| NM_005175.3 | XR_002501624.1 |
| NM_005186.4 | XM_001654160.2 |
| NM_005189.3 | XM_021853143.1 |
| NM_005191.4 | XM_021841737.1 |
| NM_005230.4 | XM_021843191.1 |
| NM_005235.3 | XM_001662404.2 |
| NM_005258.3 | XM_021845467.1 |
| NM_005262.3 | XM_021848319.1 |
| NM_005278.5 | XM_021857463.1 |
| NM_005294.3 | XM_021849075.1 |
| NM_005313.5 | XM_021855687.1 |
| NM_005319.4 | XM_021849500.1 |
| NM_005327.7 | XM_001650932.2 |
| NM_005360.5 | XM_021846295.1 |
| NM_005365.5 | XM_021842794.1 |
| NM_005372.1 | XM_001653158.2 |
| NM_005374.5 | XR_002499323.1 |
| NM_005387.7 | XR_002498692.1 |
| NM_005398.7 | XM_001658094.2 |
| NM_005426.3 | XM_001650248.2 |
| NM_005460.4 | XM_021857074.1 |
| NM_005469.4 | XM_001659913.2 |
| NM_005486.3 | XM_001658884.2 |
| NM_005496.3 | XM_001655116.2 |
| NM_005530.3 | XM_021856973.1 |
| NM_005534.4 | XM_021846930.1 |
| NM_005537.5 | XM_001650071.2 |
| NM_005554.4 | XM_021838685.1 |
| NM_005568.5 | XM_021848621.1 |
| NM_005600.3 | XM_021850947.1 |
| NM_005640.3 | XM_021852842.1 |
| NM_005658.5 | XR_002500962.1 |
| NM_005670.4 | XM_021840629.1 |
| NM_005691.4 | XM_021856013.1 |
| NM_005720.4 | XM_021841927.1 |
| NM_005729.4 | XM_021843458.1 |
| NM_005741.5 | XM_021845383.1 |
| NM_005744.5 | XM_001660975.2 |
| NM_005745.8 | XM_021844874.1 |
| NM_005802.5 | XM_021847814.1 |
| NM_005803.4 | XM_021855338.1 |
| NM_005817.5 | XM_021843878.1 |
| NM_005819.6 | XM_021847943.1 |
| NM_005824.3 | XM_021840369.1 |
| NM_005838.4 | XR_002502382.1 |
| NM_005894.3 | XR_002502163.1 |

| | |
|---|---|
| NM_005902.4 | XM_021847054.1 |
| NM_005914.4 | XM_021854333.1 |
| NM_005920.4 | XM_001658008.2 |
| NM_005928.4 | XM_021844627.1 |
| NM_005968.5 | XM_001653812.2 |
| NM_005987.4 | XM_021843270.1 |
| NM_006003.3 | XM_021851129.1 |
| NM_006009.4 | XM_021849912.1 |
| NM_006016.6 | XM_021838966.1 |
| NM_006035.4 | XR_002501370.1 |
| NM_006080.3 | XM_021839703.1 |
| NM_006092.4 | XM_021839601.1 |
| NM_006134.7 | XM_021849190.1 |
| NM_006140.6 | XM_021852849.1 |
| NM_006156.3 | XM_021846176.1 |
| NM_006177.5 | XM_021854800.1 |
| NM_006192.5 | XR_002499060.1 |
| NM_006237.4 | XM_021854475.1 |
| NM_006242.4 | XM_021854381.1 |
| NM_006246.5 | XM_021843194.1 |
| NM_006301.4 | XM_001660080.2 |
| NM_006315.6 | XM_001655013.2 |
| NM_006331.8 | XM_021854555.1 |
| NM_006341.4 | XM_021856117.1 |
| NM_006345.4 | XM_011495344.2 |
| NM_006350.5 | XM_021839228.1 |
| NM_006391.3 | XM_021853635.1 |
| NM_006408.4 | XM_001648071.2 |
| NM_006424.3 | XM_021848105.1 |
| NM_006456.3 | XM_021857466.1 |
| NM_006460.3 | XM_021856515.1 |
| NM_006462.6 | XM_021846075.1 |
| NM_006467.3 | XM_021847567.1 |
| NM_006480.5 | XM_021857640.1 |
| NM_006495.4 | XR_002500665.1 |
| NM_006536.7 | XM_021847175.1 |
| NM_006560.4 | XM_021854386.1 |
| NM_006567.5 | XM_021850980.1 |
| NM_006573.5 | XM_001657334.2 |
| NM_006585.4 | XM_021845479.1 |
| NM_006615.3 | XM_021854264.1 |
| NM_006653.5 | XM_021849502.1 |
| NM_006657.3 | XM_001664068.2 |
| NM_006668.2 | XM_021856640.1 |
| NM_006700.3 | XR_002500053.1 |
| NM_006721.4 | XM_021856641.1 |
| NM_006734.4 | XM_021838648.1 |
| NM_006755.2 | XR_002500010.1 |
| NM_006790.3 | XM_021842252.1 |
| NM_006808.3 | XM_021848626.1 |

| | |
|---|---|
| NM_006818.4 | XM_021843035.1 |
| NM_006828.4 | XM_021856532.1 |
| NM_006829.3 | XM_021852431.1 |
| NM_006833.5 | XM_021842762.1 |
| NM_006868.4 | XM_021845373.1 |
| NM_006875.4 | XM_021842545.1 |
| NM_006888.6 | XM_021844184.1 |
| NM_006928.5 | XM_001663342.2 |
| NM_006932.5 | XM_001658735.2 |
| NM_006980.5 | XM_001654145.2 |
| NM_006999.6 | XR_002501443.1 |
| NM_007006.3 | XM_021846932.1 |
| NM_007007.3 | XM_021842589.1 |
| NM_007010.5 | XM_021845697.1 |
| NM_007108.4 | XM_021842366.1 |
| NM_007112.5 | XM_001656937.2 |
| NM_007118.4 | XM_001650437.2 |
| NM_007134.1 | XM_021848316.1 |
| NM_007192.4 | XM_021854174.1 |
| NM_007194.4 | XM_001661402.2 |
| NM_007196.4 | XM_001653671.3 |
| NM_007210.4 | XM_021854259.1 |
| NM_007211.5 | XM_021857336.1 |
| NM_007219.5 | XM_021843455.1 |
| NM_007223.3 | XM_001661474.2 |
| NM_007267.7 | XM_001648244.2 |
| NM_007276.5 | XM_021851280.1 |
| NM_007278.2 | XM_021844082.1 |
| NM_007294.4 | XR_002502844.1 |
| NM_007333.2 | XR_002502721.1 |
| NM_009589.5 | XM_021853768.1 |
| NM_012088.3 | XM_021846177.1 |
| NM_012116.4 | XM_021847940.1 |
| NM_012130.4 | XM_021841960.1 |
| NM_012133.6 | XM_001659167.2 |
| NM_012138.4 | XM_021855515.1 |
| NM_012154.5 | XM_021840480.1 |
| NM_012164.4 | XM_021851338.1 |
| NM_012182.3 | XM_021857642.1 |
| NM_012207.3 | XM_021846790.1 |
| NM_012257.4 | XM_021856737.1 |
| NM_012265.3 | XM_001653004.2 |
| NM_012282.4 | XM_021852404.1 |
| NM_012294.5 | XM_001661916.2 |
| NM_012402.5 | XM_021849231.1 |
| NM_012413.4 | XM_021845100.1 |
| NM_012423.4 | XM_021848622.1 |
| NM_012427.5 | XM_001661687.2 |
| NM_012461.3 | XM_021841749.1 |
| NM_013239.5 | XM_021851614.1 |

| | |
|---|---|
| NM_013246.3 | XM_021849604.1 |
| NM_013249.4 | XM_001657750.2 |
| NM_013252.3 | XM_021856015.1 |
| NM_013264.5 | XM_021847578.1 |
| NM_013306.5 | XM_001654327.2 |
| NM_013308.4 | XM_021842419.1 |
| NM_013363.4 | XM_001663344.2 |
| NM_013365.5 | XM_021850449.1 |
| NM_013410.4 | XR_002501987.1 |
| NM_013444.4 | XM_021847997.1 |
| NM_013449.4 | XM_021842259.1 |
| NM_013450.4 | XM_001660021.2 |
| NM_013975.4 | XM_021839793.1 |
| NM_014015.4 | XM_021842645.1 |
| NM_014017.4 | XM_021848605.1 |
| NM_014045.5 | XM_021843927.1 |
| NM_014053.4 | XM_021839936.1 |
| NM_014058.4 | XM_021850526.1 |
| NM_014060.3 | XM_021855166.1 |
| NM_014064.4 | XM_021850082.1 |
| NM_014071.5 | XM_021853141.1 |
| NM_014203.3 | XM_021845758.1 |
| NM_014205.4 | XM_021855914.1 |
| NM_014264.5 | XM_021856214.1 |
| NM_014276.4 | XM_001653563.2 |
| NM_014284.3 | XM_021840976.1 |
| NM_014287.4 | XM_021857550.1 |
| NM_014339.7 | XM_001654331.2 |
| NM_014343.3 | XM_001660376.2 |
| NM_014349.3 | XR_002501678.1 |
| NM_014480.4 | XM_021843653.1 |
| NM_014482.3 | XM_001660208.2 |
| NM_014488.5 | XM_021841588.1 |
| NM_014563.6 | XM_021838207.1 |
| NM_014596.6 | XR_002499194.1 |
| NM_014598.4 | XM_001652416.2 |
| NM_014602.3 | XM_021857579.1 |
| NM_014633.5 | XR_002502317.1 |
| NM_014639.4 | XM_021849852.1 |
| NM_014640.5 | XM_021848418.1 |
| NM_014709.4 | XM_021838711.1 |
| NM_014757.5 | XM_021847188.1 |
| NM_014812.3 | XM_021852699.1 |
| NM_014813.3 | XM_021850463.1 |
| NM_014822.4 | XM_001663572.2 |
| NM_014848.7 | XM_021852810.1 |
| NM_014861.4 | XM_021845384.1 |
| NM_014872.3 | XM_021850046.1 |
| NM_014882.3 | XM_001655952.2 |
| NM_014886.6 | XM_021842135.1 |

| | |
|---|---|
| NM_014903.6 | XR_002502676.1 |
| NM_014912.5 | XM_021855418.1 |
| NM_014917.4 | XM_021847636.1 |
| NM_014944.4 | XM_021844970.1 |
| NM_014961.5 | XM_001661027.2 |
| NM_014977.3 | XM_021845013.1 |
| NM_015000.4 | XR_002500800.1 |
| NM_015025.4 | XM_021853767.1 |
| NM_015082.2 | XM_021843064.1 |
| NM_015114.3 | XM_021847714.1 |
| NM_015147.3 | XM_021849292.1 |
| NM_015151.4 | XM_001650570.2 |
| NM_015189.3 | XM_021847733.1 |
| NM_015196.4 | XM_001657619.2 |
| NM_015205.3 | XM_021855676.1 |
| NM_015232.2 | XM_021851898.1 |
| NM_015241.3 | XR_002502203.1 |
| NM_015245.3 | XM_021854749.1 |
| NM_015253.2 | XM_021856334.1 |
| NM_015264.2 | XM_021857291.1 |
| NM_015266.3 | XM_021848217.1 |
| NM_015270.5 | XM_001651977.2 |
| NM_015284.4 | XR_002499372.1 |
| NM_015319.2 | XM_021839201.1 |
| NM_015323.5 | XM_021844968.1 |
| NM_015331.3 | XM_021838716.1 |
| NM_015367.4 | XM_021837653.1 |
| NM_015395.3 | XM_021851476.1 |
| NM_015425.6 | XR_002502860.1 |
| NM_015434.4 | XM_021837786.1 |
| NM_015436.4 | XM_021850178.1 |
| NM_015471.4 | XM_021840726.1 |
| NM_015513.6 | XM_021845481.1 |
| NM_015566.3 | XM_001662749.2 |
| NM_015567.2 | XR_002501845.1 |
| NM_015575.4 | XM_021857148.1 |
| NM_015596.3 | XM_021854875.1 |
| NM_015633.3 | XM_021849240.1 |
| NM_015646.6 | XM_021843827.1 |
| NM_015657.4 | XM_021855517.1 |
| NM_015692.5 | XM_021838805.1 |
| NM_015693.4 | XM_021854883.1 |
| NM_015716.5 | XM_011495463.2 |
| NM_015719.4 | XM_001656380.2 |
| NM_015722.4 | XM_021842862.1 |
| NM_015723.5 | XM_021850017.1 |
| NM_015853.5 | XM_021856171.1 |
| NM_015967.7 | XM_021838524.1 |
| NM_015986.4 | XM_021843883.1 |
| NM_016003.4 | XM_021855419.1 |

| | |
|---|---|
| NM_016021.3 | XR_002500079.1 |
| NM_016026.4 | XM_001661783.2 |
| NM_016033.3 | XM_021850465.1 |
| NM_016083.6 | XM_021839230.1 |
| NM_016085.5 | XM_021851400.1 |
| NM_016111.4 | XM_021844415.1 |
| NM_016148.5 | XM_001649704.2 |
| NM_016161.3 | XM_001650744.2 |
| NM_016170.5 | XM_021856453.1 |
| NM_016178.2 | XM_021853789.1 |
| NM_016184.4 | XM_021857332.1 |
| NM_016235.3 | XM_021851757.1 |
| NM_016275.5 | XM_021840812.1 |
| NM_016284.5 | XM_021848158.1 |
| NM_016287.5 | XM_001659479.2 |
| NM_016315.4 | XM_021838390.1 |
| NM_016321.3 | XM_021838503.1 |
| NM_016322.4 | XM_021843517.1 |
| NM_016323.4 | XM_021848060.1 |
| NM_016360.4 | XM_021844285.1 |
| NM_016379.4 | XM_021840850.1 |
| NM_016464.5 | XM_021841323.1 |
| NM_016541.3 | XR_002502449.1 |
| NM_016553.5 | XM_021847872.1 |
| NM_016562.4 | XM_021856176.1 |
| NM_016568.3 | XR_002501877.1 |
| NM_016605.3 | XM_001660101.3 |
| NM_016617.4 | XM_021843945.1 |
| NM_016628.5 | XM_021854050.1 |
| NM_016643.4 | XM_021843243.1 |
| NM_016652.6 | XM_021845247.1 |
| NM_016734.3 | XM_021838391.1 |
| NM_016829.3 | XM_021851004.1 |
| NM_017514.5 | XM_021854524.1 |
| NM_017516.3 | XM_021848635.1 |
| NM_017522.5 | XM_021847553.1 |
| NM_017527.4 | XM_021844300.1 |
| NM_017533.2 | XM_021844522.1 |
| NM_017548.5 | XM_021839663.1 |
| NM_017576.4 | XM_021848002.1 |
| NM_017620.3 | XM_021841114.1 |
| NM_017659.4 | XM_021851630.1 |
| NM_017661.4 | XM_021857478.1 |
| NM_017676.2 | XM_021855673.1 |
| NM_017677.4 | XM_021854585.1 |
| NM_017686.4 | XM_021844708.1 |
| NM_017705.4 | XM_021848455.1 |
| NM_017714.3 | XM_021851758.1 |
| NM_017719.5 | XM_021850630.1 |
| NM_017789.5 | XM_021857193.1 |

| | |
|---|---|
| NM_017816.3 | XM_021854266.1 |
| NM_017831.4 | XR_002501574.1 |
| NM_017832.4 | XM_021851413.1 |
| NM_017867.3 | XR_002499304.1 |
| NM_017872.5 | XM_021854250.1 |
| NM_017891.5 | XM_001651387.2 |
| NM_017899.4 | XR_002502253.1 |
| NM_017905.6 | XM_001653355.2 |
| NM_017954.11 | XM_021854385.1 |
| NM_017956.4 | XM_021840914.1 |
| NM_018003.4 | XR_002501500.1 |
| NM_018047.3 | XM_021843976.1 |
| NM_018059.5 | XR_002503045.1 |
| NM_018062.4 | XM_021855840.1 |
| NM_018095.6 | XR_002500504.1 |
| NM_018099.5 | XM_021846788.1 |
| NM_018120.6 | XM_001660197.2 |
| NM_018166.3 | XM_021844749.1 |
| NM_018188.5 | XM_021846737.1 |
| NM_018223.2 | XM_021855142.1 |
| NM_018242.3 | XM_021853295.1 |
| NM_018270.6 | XM_001658851.2 |
| NM_018313.5 | XM_001658130.2 |
| NM_018314.6 | XM_021852851.1 |
| NM_018318.5 | XM_021844750.1 |
| NM_018324.3 | XM_021853253.1 |
| NM_018330.7 | XM_001658604.2 |
| NM_018340.3 | XM_021845341.1 |
| NM_018342.5 | XM_021845714.1 |
| NM_018374.4 | XM_021840674.1 |
| NM_018383.5 | XM_021839626.1 |
| NM_018400.4 | XM_001658442.2 |
| NM_018404.3 | XM_001660361.2 |
| NM_018451.5 | XM_001660346.2 |
| NM_018455.6 | XM_001650902.2 |
| NM_018480.7 | XM_021844620.1 |
| NM_018487.3 | XM_001654332.2 |
| NM_018561.5 | XM_021852989.1 |
| NM_018648.4 | XM_001663218.2 |
| NM_018651.4 | XM_021850267.1 |
| NM_018669.6 | XM_021838110.1 |
| NM_018724.4 | XM_001658613.2 |
| NM_018837.4 | XM_001657980.3 |
| NM_018838.5 | XM_021844862.1 |
| NM_018842.5 | XM_021856771.1 |
| NM_018898.5 | XM_001651960.2 |
| NM_018946.4 | XM_001660751.2 |
| NM_018952.5 | XM_021849634.1 |
| NM_018956.5 | XM_021849444.1 |
| NM_018963.5 | XM_021838977.1 |

| | |
|---|---|
| NM_018970.7 | XM_021846735.1 |
| NM_018981.4 | XM_021842396.1 |
| NM_018990.4 | XM_021837620.1 |
| NM_018999.4 | XM_021853616.1 |
| NM_019046.3 | XM_021842617.1 |
| NM_019112.4 | XM_021840611.1 |
| NM_019116.3 | XR_002502491.1 |
| NM_019120.5 | XM_021853601.1 |
| NM_019601.4 | XM_021843820.1 |
| NM_019844.4 | XM_021847451.1 |
| NM_019849.3 | XM_021848661.1 |
| NM_020133.3 | XM_001654035.2 |
| NM_020154.3 | XM_001659808.2 |
| NM_020164.5 | XM_021844115.1 |
| NM_020166.5 | XR_002500787.1 |
| NM_020171.2 | XM_021838061.1 |
| NM_020203.5 | XM_021839226.1 |
| NM_020207.7 | XM_021845394.1 |
| NM_020228.3 | XR_002499903.1 |
| NM_020244.3 | XM_021855165.1 |
| NM_020248.3 | XM_021850702.1 |
| NM_020337.3 | XM_021845136.1 |
| NM_020376.4 | XR_002500838.1 |
| NM_020377.5 | XM_021842398.1 |
| NM_020409.3 | XM_021844006.1 |
| NM_020422.6 | XM_021856691.1 |
| NM_020425.6 | XM_001661052.2 |
| NM_020469.3 | XM_021838960.1 |
| NM_020485.6 | XM_021856749.1 |
| NM_020528.3 | XM_021855584.1 |
| NM_020677.6 | XM_021850043.1 |
| NM_020692.3 | XM_001660707.2 |
| NM_020741.3 | XM_001655106.2 |
| NM_020760.4 | XM_021847486.1 |
| NM_020765.3 | XM_021855867.1 |
| NM_020781.4 | XR_002502926.1 |
| NM_020784.3 | XM_021838342.1 |
| NM_020824.4 | XM_021857266.1 |
| NM_020827.3 | XM_021847455.1 |
| NM_020830.5 | XM_011495178.2 |
| NM_020839.4 | XM_021844113.1 |
| NM_020853.2 | XM_021848937.1 |
| NM_020886.4 | XM_001659822.2 |
| NM_020911.2 | XM_021848122.1 |
| NM_020920.4 | XM_021837546.1 |
| NM_020959.3 | XM_021857019.1 |
| NM_020962.3 | XM_001657957.3 |
| NM_020984.4 | XM_001654924.2 |
| NM_020989.4 | XM_021853360.1 |
| NM_021033.7 | XR_002502777.1 |

| | |
|---|---|
| NM_021072.4 | XM_021845515.1 |
| NM_021104.2 | XM_021853014.1 |
| NM_021110.4 | XR_002499151.1 |
| NM_021134.4 | XM_021840325.1 |
| NM_021158.5 | XM_001648222.2 |
| NM_021168.5 | XM_021856111.1 |
| NM_021186.5 | XM_021850047.1 |
| NM_021192.3 | XM_021849253.1 |
| NM_021200.3 | XM_021852402.1 |
| NM_021202.3 | XM_021846896.1 |
| NM_021225.5 | XM_021853391.1 |
| NM_021245.4 | XR_002499979.1 |
| NM_021252.5 | XM_021852889.1 |
| NM_021574.3 | XM_021846846.1 |
| NM_021603.4 | XM_021840851.1 |
| NM_021623.2 | XM_021838672.1 |
| NM_021634.4 | XM_001651377.2 |
| NM_021643.4 | XM_021843975.1 |
| NM_021806.4 | XM_001650372.2 |
| NM_021983.4 | XM_021848770.1 |
| NM_022053.4 | XM_001653662.2 |
| NM_022058.4 | XM_021846446.1 |
| NM_022064.5 | XR_002503146.1 |
| NM_022098.4 | XR_002501635.1 |
| NM_022119.4 | XM_021841985.1 |
| NM_022135.4 | XM_021850049.1 |
| NM_022142.5 | XM_021845770.1 |
| NM_022170.2 | XM_021840907.1 |
| NM_022355.4 | XM_021841754.1 |
| NM_022437.3 | XM_001647598.2 |
| NM_022464.5 | XM_021843467.1 |
| NM_022468.5 | XM_021852239.1 |
| NM_022568.4 | XM_021855854.1 |
| NM_022645.2 | XR_002501197.1 |
| NM_022727.6 | XM_021845395.1 |
| NM_022757.5 | XM_021851795.1 |
| NM_022764.3 | XM_021844507.1 |
| NM_022765.4 | XR_002499855.1 |
| NM_022774.3 | XM_021840059.1 |
| NM_022784.3 | XM_021847833.1 |
| NM_022817.3 | XM_001657987.2 |
| NM_022818.5 | XM_001649803.2 |
| NM_022823.3 | XM_021849442.1 |
| NM_022902.5 | XM_021846867.1 |
| NM_023000.3 | XR_002503086.1 |
| NM_023915.4 | XR_002500641.1 |
| NM_023918.3 | XM_001663680.2 |
| NM_023937.4 | XM_021848147.1 |
| NM_023940.3 | XM_021846327.1 |
| NM_023946.5 | XM_021839977.1 |

| | |
|---|---|
| NM_024011.4 | XM_021844131.1 |
| NM_024059.3 | XM_021840111.1 |
| NM_024061.4 | XM_021852698.1 |
| NM_024067.4 | XM_021853252.1 |
| NM_024120.5 | XM_001652178.2 |
| NM_024121.3 | XM_021852245.1 |
| NM_024299.4 | XM_021840049.1 |
| NM_024324.5 | XM_001648798.2 |
| NM_024504.4 | XM_021853901.1 |
| NM_024512.5 | XM_021849630.1 |
| NM_024525.5 | XM_021842037.1 |
| NM_024528.4 | XM_001648314.2 |
| NM_024541.3 | XM_021839171.1 |
| NM_024556.4 | XM_001656578.2 |
| NM_024568.4 | XM_021842374.1 |
| NM_024577.4 | XM_021843973.1 |
| NM_024594.4 | XM_021841308.1 |
| NM_024598.4 | XM_001656954.3 |
| NM_024613.4 | XM_021850371.1 |
| NM_024617.4 | XM_021848146.1 |
| NM_024644.5 | XM_021838825.1 |
| NM_024687.4 | XM_021837983.1 |
| NM_024729.4 | XM_021848713.1 |
| NM_024754.5 | XM_001656969.2 |
| NM_024770.5 | XM_021845615.1 |
| NM_024783.4 | XM_001656664.2 |
| NM_024792.3 | XM_021848139.1 |
| NM_024798.3 | XR_002500274.1 |
| NM_024806.4 | XM_021850374.1 |
| NM_024830.5 | XM_021841699.1 |
| NM_024869.3 | XM_021845638.1 |
| NM_024872.4 | XM_021845727.1 |
| NM_024877.4 | XM_021854419.1 |
| NM_024894.4 | XM_021854797.1 |
| NM_024895.5 | XM_021855856.1 |
| NM_024899.4 | XM_021844586.1 |
| NM_024913.5 | XM_001663357.2 |
| NM_024939.3 | XM_001655462.2 |
| NM_024940.8 | XM_021850887.1 |
| NM_024955.6 | XM_021850521.1 |
| NM_024956.4 | XM_021848144.1 |
| NM_025052.5 | XM_021845612.1 |
| NM_025058.5 | XM_001657178.2 |
| NM_025142.1 | XM_021855113.1 |
| NM_025149.6 | XM_001650856.2 |
| NM_025160.7 | XM_021841797.1 |
| NM_025205.5 | XM_021850434.1 |
| NM_025233.7 | XM_021851872.1 |
| NM_025241.3 | XM_001650649.2 |
| NM_025258.3 | XM_021841028.1 |

| | |
|---|---|
| NM_025261.3 | XM_021853176.1 |
| NM_025264.5 | XM_021855032.1 |
| NM_030592.3 | XM_021843236.1 |
| NM_030631.4 | XR_002503270.1 |
| NM_030671.3 | XM_021843974.1 |
| NM_030763.3 | XR_002500148.1 |
| NM_030772.5 | XR_002500829.1 |
| NM_030779.4 | XM_021852780.1 |
| NM_030798.5 | XR_002501172.1 |
| NM_030820.4 | XM_021842399.1 |
| NM_030904.1 | XM_021850276.1 |
| NM_030912.3 | XM_021843828.1 |
| NM_030920.5 | XM_021851441.1 |
| NM_030927.4 | XM_021851101.1 |
| NM_030941.3 | XM_021845046.1 |
| NM_030943.4 | XM_021850368.1 |
| NM_030956.4 | XM_001653621.3 |
| NM_030961.3 | XM_001652357.2 |
| NM_030969.5 | XM_021842283.1 |
| NM_031231.4 | XM_021852092.1 |
| NM_031243.3 | XM_021845488.1 |
| NM_031268.6 | XM_021845838.1 |
| NM_031280.4 | XM_021842403.1 |
| NM_031294.4 | XM_021846348.1 |
| NM_031371.4 | XM_001654644.2 |
| NM_031444.4 | XR_002500510.1 |
| NM_031455.4 | XM_021855764.1 |
| NM_031495.2 | XM_021856949.1 |
| NM_031500.3 | XM_021842524.1 |
| NM_031852.2 | XM_021850924.1 |
| NM_031860.3 | XM_001648903.2 |
| NM_031922.5 | XM_021851368.1 |
| NM_031961.3 | XM_001659147.2 |
| NM_031965.2 | XM_021853299.1 |
| NM_032044.4 | XM_021851326.1 |
| NM_032049.4 | XM_021847735.1 |
| NM_032109.3 | XM_001656631.2 |
| NM_032111.4 | XM_021839066.1 |
| NM_032118.4 | XM_021854722.1 |
| NM_032125.3 | XM_021842346.1 |
| NM_032127.4 | XR_002498687.1 |
| NM_032145.5 | XM_021850290.1 |
| NM_032149.3 | XM_001659309.2 |
| NM_032164.4 | XM_001653664.2 |
| NM_032181.3 | XM_001647738.2 |
| NM_032268.5 | XM_021838387.1 |
| NM_032271.3 | XM_021842957.1 |
| NM_032360.4 | XM_021844495.1 |
| NM_032373.5 | XM_001654255.2 |
| NM_032375.5 | XM_021842157.1 |

| | |
|---|---|
| NM_032408.4 | XM_001655976.2 |
| NM_032414.3 | XM_001662958.2 |
| NM_032477.3 | XR_002498923.1 |
| NM_032523.4 | XM_011494635.2 |
| NM_032524.2 | XM_021852216.1 |
| NM_032536.4 | XM_021847173.1 |
| NM_032539.5 | XM_001659698.2 |
| NM_032550.4 | XM_021841640.1 |
| NM_032554.4 | XM_001656410.2 |
| NM_032630.3 | XM_021853265.1 |
| NM_032681.4 | XM_001657060.2 |
| NM_032796.4 | XM_021847918.1 |
| NM_032816.5 | XM_021849592.1 |
| NM_032831.4 | XM_021852328.1 |
| NM_032843.5 | XM_021839530.1 |
| NM_032866.5 | XM_021839301.1 |
| NM_032869.4 | XM_021843446.1 |
| NM_032881.3 | XM_001655217.2 |
| NM_032980.4 | XM_001661051.2 |
| NM_033011.4 | XM_021839263.1 |
| NM_033056.4 | XM_001663673.2 |
| NM_033070.3 | XM_001651810.2 |
| NM_033092.4 | XR_002501058.1 |
| NM_033110.3 | XM_001664039.2 |
| NM_033113.3 | XM_021838839.1 |
| NM_033123.4 | XR_002498651.1 |
| NM_033169.2 | XM_021845031.1 |
| NM_033172.3 | XR_002498644.1 |
| NM_033180.5 | XR_002499503.1 |
| NM_033181.4 | XM_021854312.1 |
| NM_033191.3 | XM_001655475.2 |
| NM_033253.4 | XM_021846579.1 |
| NM_033289.2 | XM_001649708.2 |
| NM_033293.4 | XM_021851681.1 |
| NM_033295.4 | XM_001662718.2 |
| NM_033313.3 | XM_021844621.1 |
| NM_033342.4 | XM_021840656.1 |
| NM_033394.3 | XM_021847719.1 |
| NM_033413.4 | XM_001661603.2 |
| NM_033423.5 | XM_021843644.1 |
| NM_033503.4 | XM_021854877.1 |
| NM_033670.4 | XM_021840725.1 |
| NM_052868.6 | XM_021837624.1 |
| NM_052870.4 | XM_021841752.1 |
| NM_052879.5 | XR_002499564.1 |
| NM_052902.4 | XM_021839178.1 |
| NM_052904.4 | XM_021857180.1 |
| NM_052926.3 | XM_021839857.1 |
| NM_052934.4 | XM_021840492.1 |
| NM_052951.3 | XM_021845188.1 |

| | |
|---|---|
| NM_052955.3 | XM_021849311.1 |
| NM_052959.3 | XM_001658229.2 |
| NM_052961.4 | XM_001654719.2 |
| NM_053016.6 | XM_001654979.2 |
| NM_053023.5 | XM_021854114.1 |
| NM_053051.5 | XM_021847815.1 |
| NM_054021.2 | XM_021846519.1 |
| NM_057090.3 | XM_021837755.1 |
| NM_057164.5 | XM_001652683.2 |
| NM_057179.3 | XM_001658137.2 |
| NM_058163.3 | XM_021845901.1 |
| NM_058169.6 | XM_001664064.2 |
| NM_058176.2 | XM_001658134.2 |
| NM_058187.5 | XM_021840296.1 |
| NM_058197.5 | XM_021853463.1 |
| NM_078481.4 | XM_021852791.1 |
| NM_078485.4 | XM_021839861.1 |
| NM_078626.3 | XM_011495173.2 |
| NM_078628.2 | XM_001660666.2 |
| NM_080282.4 | XM_021851948.1 |
| NM_080423.3 | XM_021854327.1 |
| NM_080590.4 | XR_002500631.1 |
| NM_080659.3 | XR_002501973.1 |
| NM_080672.5 | XR_002501944.1 |
| NM_080679.3 | XM_021851062.1 |
| NM_080737.3 | XM_021840860.1 |
| NM_080833.3 | XM_021852965.1 |
| NM_080836.4 | XM_021849716.1 |
| NM_080838.3 | XM_001655358.2 |
| NM_080841.3 | XM_021845259.1 |
| NM_080862.3 | XM_021842917.1 |
| NM_080868.3 | XM_001650797.2 |
| NM_080872.4 | XM_021844730.1 |
| NM_080879.3 | XM_001649364.2 |
| NM_080913.3 | XM_021854616.1 |
| NM_130759.4 | XM_021843213.1 |
| NM_130762.3 | XM_021852220.1 |
| NM_130787.3 | XM_001656421.2 |
| NM_130852.3 | XM_001663220.2 |
| NM_133277.4 | XM_021853472.1 |
| NM_133344.3 | XM_021844337.1 |
| NM_133373.5 | XM_021841644.1 |
| NM_133506.3 | XM_021842496.1 |
| NM_133635.6 | XM_021837639.1 |
| NM_134425.4 | XM_021849960.1 |
| NM_134440.3 | XM_001662875.2 |
| NM_134442.5 | XM_021856817.1 |
| NM_138286.3 | XM_001655960.2 |
| NM_138324.3 | XM_021850612.1 |
| NM_138348.6 | XM_021844773.1 |

| | |
|---|---|
| NM_138355.4 | XM_021840788.1 |
| NM_138373.5 | XM_021845480.1 |
| NM_138391.6 | XR_002501709.1 |
| NM_138413.4 | XM_001657749.2 |
| NM_138431.3 | XM_021844720.1 |
| NM_138461.4 | XR_002501527.1 |
| NM_138476.4 | XM_021852214.1 |
| NM_138499.4 | XM_001655434.2 |
| NM_138700.4 | XM_021847545.1 |
| NM_138702.1 | XM_021845632.1 |
| NM_138705.4 | XM_001663174.2 |
| NM_138711.6 | XM_021853467.1 |
| NM_138714.4 | XM_021837507.1 |
| NM_138736.3 | XM_021842859.1 |
| NM_138771.4 | XM_021842413.1 |
| NM_138782.3 | XM_021840791.1 |
| NM_138788.5 | XM_001650091.2 |
| NM_138804.5 | XM_021840191.1 |
| NM_138810.4 | XM_021839732.1 |
| NM_138927.4 | XM_001655012.2 |
| NM_138940.3 | XM_021838180.1 |
| NM_138973.4 | XR_002501286.1 |
| NM_139052.3 | XM_021841102.1 |
| NM_139074.4 | XM_021852707.1 |
| NM_139075.4 | XR_002502883.1 |
| NM_139136.4 | XM_021844569.1 |
| NM_139137.4 | XM_021855004.1 |
| NM_139209.3 | XM_001656632.2 |
| NM_139246.5 | XM_001656511.3 |
| NM_139314.3 | XM_021838157.1 |
| NM_144569.6 | XM_001649542.2 |
| NM_144608.2 | XM_021855590.1 |
| NM_144617.3 | XM_021841152.1 |
| NM_144632.5 | XM_021857043.1 |
| NM_144666.3 | XM_021853139.1 |
| NM_144672.4 | XM_001651364.2 |
| NM_144691.4 | XM_021840481.1 |
| NM_144707.4 | XM_001652727.2 |
| NM_144720.4 | XM_021849915.1 |
| NM_144725.4 | XM_021838784.1 |
| NM_144777.3 | XM_001648546.2 |
| NM_144778.4 | XM_001649235.2 |
| NM_144976.4 | XM_021841307.1 |
| NM_144989.3 | XM_001654512.2 |
| NM_145071.4 | XM_021852198.1 |
| NM_145166.4 | XR_002501843.1 |
| NM_145177.3 | XM_021853156.1 |
| NM_145178.4 | XM_021844729.1 |
| NM_145186.3 | XM_021846372.1 |
| NM_145241.5 | XM_021838179.1 |

| | |
|---|---|
| NM_145248.5 | XM_021857591.1 |
| NM_145257.5 | XM_021849514.1 |
| NM_145278.5 | XM_021850804.1 |
| NM_145333.3 | XM_021850805.1 |
| NM_145658.4 | XM_021855441.1 |
| NM_145808.4 | XM_021837756.1 |
| NM_145888.3 | XM_021851856.1 |
| NM_147127.5 | XM_001663647.2 |
| NM_147185.3 | XM_021855851.1 |
| NM_147686.4 | XM_001663698.2 |
| NM_148963.4 | XM_021843054.1 |
| NM_152244.2 | XM_021840790.1 |
| NM_152246.3 | XM_021840493.1 |
| NM_152274.5 | XM_021844575.1 |
| NM_152280.5 | XM_001651378.2 |
| NM_152282.5 | XM_021841383.1 |
| NM_152317.4 | XM_021847836.1 |
| NM_152318.3 | XM_001660068.2 |
| NM_152331.4 | XM_021850275.1 |
| NM_152344.4 | XM_021847075.1 |
| NM_152365.3 | XM_021851372.1 |
| NM_152369.5 | XM_001658043.2 |
| NM_152373.4 | XM_021839903.1 |
| NM_152374.2 | XM_021847721.1 |
| NM_152417.3 | XM_021852436.1 |
| NM_152432.4 | XM_021841905.1 |
| NM_152467.5 | XM_021852843.1 |
| NM_152496.3 | XR_002501695.1 |
| NM_152498.3 | XM_001649226.2 |
| NM_152512.4 | XR_002499022.1 |
| NM_152613.3 | XM_021841479.1 |
| NM_152614.3 | XM_021847989.1 |
| NM_152628.4 | XM_001648272.2 |
| NM_152641.4 | XM_021845047.1 |
| NM_152654.3 | XM_001654410.2 |
| NM_152682.4 | XM_021843520.1 |
| NM_152694.3 | XM_021839320.1 |
| NM_152724.3 | XM_021850772.1 |
| NM_152731.3 | XM_001651858.2 |
| NM_152753.4 | XR_002500917.1 |
| NM_152779.4 | XM_001657533.2 |
| NM_152792.4 | XM_021841901.1 |
| NM_152795.4 | XM_021851313.1 |
| NM_152830.3 | XM_021841746.1 |
| NM_152841.2 | XM_001662376.2 |
| NM_152869.4 | XM_001651874.2 |
| NM_152926.3 | XM_021856096.1 |
| NM_152990.4 | XR_002500447.1 |
| NM_153006.3 | XM_021856525.1 |
| NM_153047.4 | XM_021842717.1 |

| | |
|---|---|
| NM_153211.4 | XM_021842202.1 |
| NM_153273.4 | XM_001648690.2 |
| NM_153332.4 | XM_021849305.1 |
| NM_153437.3 | XM_021839231.1 |
| NM_153439.1 | XM_021839001.1 |
| NM_153456.4 | XM_021840450.1 |
| NM_153478.3 | XM_021853483.1 |
| NM_153480.2 | XM_021840322.1 |
| NM_153608.4 | XM_001659703.2 |
| NM_153694.5 | XM_001657875.2 |
| NM_153695.4 | XM_001648980.3 |
| NM_153712.5 | XM_021843442.1 |
| NM_153713.3 | XM_021853716.1 |
| NM_170750.3 | XM_021845051.1 |
| NM_171829.3 | XM_001661135.2 |
| NM_172107.4 | XM_001658859.2 |
| NM_172108.5 | XR_002500189.1 |
| NM_172173.3 | XR_002500002.1 |
| NM_172225.2 | XM_001651537.2 |
| NM_172316.3 | XR_002502928.1 |
| NM_172341.4 | XR_002502009.1 |
| NM_173081.5 | XM_021854738.1 |
| NM_173156.3 | XM_021848445.1 |
| NM_173163.3 | XM_021854515.1 |
| NM_173208.3 | XM_021844734.1 |
| NM_173216.2 | XR_002501122.1 |
| NM_173491.4 | XM_021850951.1 |
| NM_173497.4 | XM_021841909.1 |
| NM_173512.2 | XM_021856435.1 |
| NM_173515.4 | XM_021837515.1 |
| NM_173591.7 | XR_002500329.1 |
| NM_173595.4 | XR_002502914.1 |
| NM_173619.4 | XM_001660857.2 |
| NM_173626.4 | XM_021838568.1 |
| NM_173637.4 | XM_001662997.2 |
| NM_173653.4 | XM_021839760.1 |
| NM_173678.3 | XM_021840662.1 |
| NM_173689.7 | XM_001657506.2 |
| NM_173701.2 | XM_021849734.1 |
| NM_173803.4 | XM_021844655.1 |
| NM_173808.3 | XR_002501329.1 |
| NM_173826.4 | XM_021852407.1 |
| NM_173831.4 | XM_021856660.1 |
| NM_174882.3 | XM_021856197.1 |
| NM_174894.3 | XM_021856180.1 |
| NM_174931.4 | XM_021847555.1 |
| NM_174934.4 | XR_002502681.1 |
| NM_174969.4 | XM_021850156.1 |
| NM_175039.4 | XM_021850413.1 |
| NM_175066.4 | XR_002500265.1 |

| | |
|---|---|
| NM_175569.3 | XM_001649085.2 |
| NM_175739.4 | XM_021850053.1 |
| NM_175861.3 | XM_001656542.2 |
| NM_176787.5 | XM_021845582.1 |
| NM_177452.4 | XM_001661566.2 |
| NM_177536.4 | XM_021843723.1 |
| NM_177937.3 | XM_021851130.1 |
| NM_177972.3 | XM_001660445.2 |
| NM_177990.4 | XR_002498656.1 |
| NM_178151.3 | XM_021837691.1 |
| NM_178174.3 | XM_021839109.1 |
| NM_178177.5 | XM_021841108.1 |
| NM_178232.4 | XM_021856743.1 |
| NM_178332.2 | XM_021855194.1 |
| NM_178353.2 | XR_002499310.1 |
| NM_178354.3 | XM_021848828.1 |
| NM_178356.3 | XM_001660561.2 |
| NM_178452.6 | XM_021839637.1 |
| NM_178517.5 | XM_021846190.1 |
| NM_178543.5 | XR_002499496.1 |
| NM_178569.4 | XM_001661479.2 |
| NM_178587.3 | XM_021852993.1 |
| NM_178812.4 | XM_021842867.1 |
| NM_178813.6 | XR_002499347.1 |
| NM_178862.3 | XR_002499147.1 |
| NM_180982.3 | XM_021843554.1 |
| NM_181078.3 | XM_021851435.1 |
| NM_181359.3 | XM_021852059.1 |
| NM_181523.3 | XR_002501290.1 |
| NM_181538.3 | XM_021857490.1 |
| NM_181644.5 | XM_001663205.2 |
| NM_181673.3 | XM_001652914.2 |
| NM_181676.3 | XM_021855456.1 |
| NM_181715.3 | XM_021855216.1 |
| NM_181790.1 | XM_021853481.1 |
| NM_181892.4 | XM_001659792.2 |
| NM_182489.2 | XM_001653554.2 |
| NM_182491.4 | XM_021848788.1 |
| NM_182497.4 | XM_001661558.2 |
| NM_182498.4 | XM_021855208.1 |
| NM_182513.3 | XM_001662835.2 |
| NM_182581.4 | XM_021855390.1 |
| NM_182616.4 | XM_001652280.2 |
| NM_182628.3 | XM_021837294.1 |
| NM_182691.3 | XM_001655751.2 |
| NM_182719.2 | XM_001661243.2 |
| NM_182724.2 | XM_021852395.1 |
| NM_182767.6 | XM_001658318.2 |
| NM_183059.3 | XM_021840572.1 |
| NM_183060.3 | XM_001648695.2 |

| | |
|---|---|
| NM_183239.2 | XM_021845360.1 |
| NM_183323.3 | XM_021844758.1 |
| NM_183352.3 | XM_021841822.1 |
| NM_183361.3 | XM_021854662.1 |
| NM_183372.9 | XM_001655821.2 |
| NM_183419.4 | XR_002500081.1 |
| NM_187841.3 | XM_021854053.1 |
| NM_194272.3 | XM_021853264.1 |
| NM_194278.3 | XM_001654448.2 |
| NM_194294.2 | XM_021838365.1 |
| NM_194332.3 | XM_001663041.2 |
| NM_194447.3 | XM_001661650.2 |
| NM_197954.3 | XM_021856192.1 |
| NM_197966.3 | XM_021843722.1 |
| NM_197970.3 | XM_021852763.1 |
| NM_198057.3 | XM_021856110.1 |
| NM_198098.4 | XM_021854858.1 |
| NM_198186.3 | XM_021853050.1 |
| NM_198205.2 | XM_021845532.1 |
| NM_198269.3 | XM_021855761.1 |
| NM_198273.2 | XM_021851693.1 |
| NM_198316.2 | XM_021845787.1 |
| NM_198383.3 | XM_001648703.2 |
| NM_198404.3 | XM_021852787.1 |
| NM_198476.5 | XM_021848453.1 |
| NM_198493.3 | XM_001650093.2 |
| NM_198498.3 | XM_021853077.1 |
| NM_198530.4 | XR_002502772.1 |
| NM_198531.5 | XM_001663703.2 |
| NM_198686.3 | XM_021854299.1 |
| NM_198687.2 | XM_021846191.1 |
| NM_198699.1 | XM_021843304.1 |
| NM_198977.2 | XM_021843698.1 |
| NM_198996.4 | XM_021841784.1 |
| NM_198999.3 | XM_001660340.2 |
| NM_199123.2 | XM_021846548.1 |
| NM_199127.2 | XM_001648185.2 |
| NM_199162.3 | XM_021850139.1 |
| NM_199177.4 | XR_002500886.1 |
| NM_199184.2 | XM_021842204.1 |
| NM_199231.2 | XM_021842504.1 |
| NM_199245.3 | XM_021841305.1 |
| NM_199263.3 | XM_021838332.1 |
| NM_199321.3 | XM_001658373.2 |
| NM_199334.5 | XM_021841781.1 |
| NM_199344.3 | XM_021855855.1 |
| NM_199350.3 | XM_021845816.1 |
| NM_199363.3 | XM_021848127.1 |
| NM_199478.3 | XM_021852330.1 |
| NM_199487.3 | XM_001647941.2 |

| | |
|---|---|
| NM_201262.2 | XM_021852437.1 |
| NM_201380.4 | XR_002499828.1 |
| NM_201402.3 | XM_021840445.1 |
| NM_201414.3 | XM_001651231.2 |
| NM_201440.2 | XR_002499858.1 |
| NM_201441.3 | XM_021850624.1 |
| NM_201525.4 | XM_001650195.2 |
| NM_201557.4 | XM_021854661.1 |
| NM_201575.4 | XM_021851521.1 |
| NM_201632.5 | XM_021839265.1 |
| NM_203295.2 | XM_021848831.1 |
| NM_203344.3 | XM_021841482.1 |
| NM_203374.2 | XM_001656365.2 |
| NM_203453.5 | XM_021839851.1 |
| NM_203459.3 | XR_002499350.1 |
| NM_205768.3 | XM_001658649.2 |
| NM_205862.2 | XM_021848220.1 |
| NM_206825.2 | XM_001648022.2 |
| NM_206890.3 | XM_021848825.1 |
| NM_206909.3 | XM_021853570.1 |
| NM_206920.3 | XM_021840065.1 |
| NM_207128.3 | XM_001657185.2 |
| NM_207334.3 | XM_021851950.1 |
| NM_207335.4 | XM_001652445.2 |
| NM_207352.4 | XM_021854096.1 |
| NM_207364.2 | XM_021840523.1 |
| NM_207372.2 | XM_021838377.1 |
| NM_207417.3 | XM_021847564.1 |
| NM_207426.3 | XM_021839557.1 |
| NM_207519.2 | XM_021855410.1 |
| NM_212472.2 | XM_001660839.2 |
| NM_212478.3 | XM_021844203.1 |
| NM_212543.2 | XR_002500143.1 |
| NM_213603.3 | XM_001653037.2 |
| NM_213674.1 | XM_021848295.1 |
| NR_001574.2 | XR_002499002.1 |
| NR_002190.1 | XM_021855579.1 |
| NR_002311.1 | XM_001657874.2 |
| NR_002717.2 | XR_002499927.1 |
| NR_002720.2 | XM_001662534.2 |
| NR_002724.2 | XM_021845223.1 |
| NR_002807.4 | XM_021844731.1 |
| NR_002832.2 | XR_002500243.1 |
| NR_002836.2 | XM_001648188.2 |
| NR_002923.2 | XM_021852329.1 |
| NR_003040.2 | XM_021842363.1 |
| NR_003090.3 | XM_021842249.1 |
| NR_003187.3 | XM_021847816.1 |
| NR_003189.2 | XM_001653596.2 |
| NR_003228.1 | XM_021839519.1 |

| | |
|---|---|
| NR_003260.1 | XM_021843725.1 |
| NR_015358.2 | XM_021845617.1 |
| NR_015419.2 | XM_001664115.2 |
| NR_015436.2 | XM_021844810.1 |
| NR_015440.1 | XM_021840447.1 |
| NR_015448.1 | XM_021849940.1 |
| NR_016021.2 | XM_021851989.1 |
| NR_023349.3 | XM_021842165.1 |
| NR_023385.1 | XM_021845283.1 |
| NR_024027.2 | XM_021840660.1 |
| NR_024035.2 | XM_001655373.2 |
| NR_024056.2 | XM_021843870.1 |
| NR_024122.2 | XM_001659683.2 |
| NR_024159.1 | XR_002502268.1 |
| NR_024162.1 | XM_021852320.1 |
| NR_024188.3 | XM_021841971.1 |
| NR_024237.2 | XM_001656411.2 |
| NR_024248.1 | XM_001661062.2 |
| NR_024256.1 | XR_002500489.1 |
| NR_024269.1 | XM_021848914.1 |
| NR_024274.1 | XM_001655435.2 |
| NR_024345.1 | XM_001651868.2 |
| NR_024397.1 | XM_021854733.1 |
| NR_024435.2 | XM_021855552.1 |
| NR_024448.2 | XM_021845216.1 |
| NR_024452.1 | XM_021842166.1 |
| NR_024455.1 | XM_001662902.2 |
| NR_024457.2 | XM_001660023.2 |
| NR_024511.2 | XR_002499099.1 |
| NR_024515.2 | XM_021841981.1 |
| NR_024531.1 | XM_001652000.2 |
| NR_024533.1 | XM_001662372.2 |
| NR_024542.1 | XM_001663156.2 |
| NR_024603.1 | XM_021840665.1 |
| NR_024606.2 | XM_021847205.1 |
| NR_026542.1 | XM_021852675.1 |
| NR_026564.1 | XR_002503047.1 |
| NR_026597.2 | XM_021857142.1 |
| NR_026656.1 | XM_021852086.1 |
| NR_026708.2 | XM_021845839.1 |
| NR_026803.2 | XM_001649793.2 |
| NR_026818.1 | XM_021843764.1 |
| NR_026825.2 | XM_001651811.2 |
| NR_026857.1 | XM_021855504.1 |
| NR_026873.1 | XM_021851652.1 |
| NR_026878.1 | XM_021857264.1 |
| NR_026929.1 | XM_001653873.2 |
| NR_026933.2 | XM_001654852.2 |
| NR_026943.1 | XR_002502630.1 |
| NR_026963.1 | XM_021848827.1 |

| | |
|---|---|
| NR_026971.1 | XM_021841223.1 |
| NR_026988.1 | XM_021848350.1 |
| NR_027055.1 | XM_001662273.2 |
| NR_027072.2 | XM_021848273.1 |
| NR_027073.1 | XM_021845289.1 |
| NR_027078.1 | XM_001656905.2 |
| NR_027097.2 | XR_002502782.1 |
| NR_027111.1 | XR_002499143.1 |
| NR_027148.1 | XR_002500337.1 |
| NR_027163.1 | XM_021845115.1 |
| NR_027176.1 | XM_021838643.1 |
| NR_027232.1 | XM_021847509.1 |
| NR_027242.1 | XM_021845483.1 |
| NR_027277.2 | XM_021841608.1 |
| NR_027279.1 | XR_002500231.1 |
| NR_027354.2 | XM_021856307.1 |
| NR_027355.2 | XM_021857583.1 |
| NR_027365.3 | XM_021837288.1 |
| NR_027368.2 | XM_021847508.1 |
| NR_027379.1 | XM_021851228.1 |
| NR_027413.2 | XM_001654515.2 |
| NR_027418.1 | XM_021841639.1 |
| NR_027451.1 | XM_021838101.1 |
| NR_027454.2 | XM_001648518.2 |
| NR_027514.3 | XM_021857265.1 |
| NR_027632.1 | XM_021853783.1 |
| NR_027645.1 | XM_021846371.1 |
| NR_027684.2 | XM_001655596.2 |
| NR_027711.1 | XM_021852992.1 |
| NR_027749.3 | XM_021856722.1 |
| NR_027754.2 | XM_021849579.1 |
| NR_027763.2 | XM_021854618.1 |
| NR_027855.2 | XM_001654176.2 |
| NR_027946.3 | XM_021846955.1 |
| NR_027948.3 | XR_002499157.1 |
| NR_028025.3 | XM_021843933.1 |
| NR_028035.4 | XM_001654966.2 |
| NR_028272.1 | XM_001652239.2 |
| NR_028415.1 | XM_021849794.1 |
| NR_028436.3 | XM_021848513.1 |
| NR_028477.2 | XM_021845960.1 |
| NR_029392.1 | XM_021846623.1 |
| NR_029403.2 | XM_021848880.1 |
| NR_029408.1 | XM_021840378.1 |
| NR_030697.1 | XM_021855076.1 |
| NR_030770.2 | XM_001661095.2 |
| NR_033186.1 | XM_021857250.1 |
| NR_033267.1 | XM_021842420.1 |
| NR_033359.1 | XM_021849936.1 |
| NR_033658.1 | XM_021852974.1 |

| | |
|---|---|
| NR_033660.2 | XM_001654095.2 |
| NR_033702.2 | XM_021855077.1 |
| NR_033757.3 | XM_021840505.1 |
| NR_033807.3 | XM_021844882.1 |
| NR_033818.2 | XM_021848936.1 |
| NR_033835.1 | XM_021846720.1 |
| NR_033838.1 | XM_021850815.1 |
| NR_033839.1 | XM_021846936.1 |
| NR_033857.1 | XM_021854018.1 |
| NR_033866.1 | XM_001662650.2 |
| NR_033876.1 | XM_021843700.1 |
| NR_033894.1 | XM_001656587.2 |
| NR_033906.2 | XM_021849991.1 |
| NR_033938.1 | XM_021853477.1 |
| NR_033955.2 | XM_001652977.2 |
| NR_033965.1 | XM_001654776.2 |
| NR_033995.2 | XM_021852282.1 |
| NR_034054.1 | XM_021851357.1 |
| NR_034095.1 | XM_021856359.1 |
| NR_034105.4 | XM_001651361.2 |
| NR_034106.3 | XM_021852973.1 |
| NR_034137.1 | XR_002498640.1 |
| NR_036433.2 | XM_021843797.1 |
| NR_036437.2 | XM_021854927.1 |
| NR_036438.1 | XM_021848915.1 |
| NR_036485.1 | XM_021857021.1 |
| NR_036489.1 | XM_021852292.1 |
| NR_036490.1 | XM_001655290.2 |
| NR_036496.1 | XM_021855998.1 |
| NR_036498.1 | XM_021845778.1 |
| NR_036503.1 | XM_021840522.1 |
| NR_036569.1 | XM_021838840.1 |
| NR_036623.2 | XM_021846431.1 |
| NR_036677.1 | XM_021846367.1 |
| NR_037141.1 | XM_021841555.1 |
| NR_037160.1 | XM_021851857.1 |
| NR_037175.1 | XM_021849704.1 |
| NR_037179.1 | XM_021849908.1 |
| NR_037602.1 | XM_021854098.1 |
| NR_037611.1 | XM_001654056.2 |
| NR_037641.2 | XM_021851416.1 |
| NR_037645.2 | XM_021850010.1 |
| NR_037646.1 | XR_002500268.1 |
| NR_037703.2 | XM_021847479.1 |
| NR_037709.1 | XM_001656496.2 |
| NR_037771.2 | XM_021842619.1 |
| NR_037801.2 | XM_021853172.1 |
| NR_037805.1 | XM_021847155.1 |
| NR_037839.1 | XR_002500633.1 |
| NR_037863.1 | XM_021842876.1 |

| | |
|---|---|
| NR_037878.1 | XM_021855199.1 |
| NR_037891.1 | XM_021843074.1 |
| NR_037916.2 | XM_001647977.2 |
| NR_037925.1 | XM_021844897.1 |
| NR_037932.1 | XM_001653200.2 |
| NR_037946.1 | XM_021848991.1 |
| NR_038103.1 | XM_021842313.1 |
| NR_038159.2 | XM_021845330.1 |
| NR_038204.1 | XM_021851802.1 |
| NR_038220.1 | XM_021847377.1 |
| NR_038223.1 | XM_021848910.1 |
| NR_038228.1 | XM_001655415.2 |
| NR_038238.1 | XM_021845078.1 |
| NR_038246.1 | XM_021838748.1 |
| NR_038319.1 | XM_001660460.2 |
| NR_038322.2 | XM_021855448.1 |
| NR_038327.2 | XM_001647815.3 |
| NR_038392.2 | XM_021856112.1 |
| NR_038413.1 | XM_021837924.1 |
| NR_038435.1 | XM_021840122.1 |
| NR_038439.1 | XM_021851597.1 |
| NR_038449.1 | XM_021843043.1 |
| NR_038877.1 | XM_021837612.1 |
| NR_038916.1 | XM_001658356.2 |
| NR_038924.1 | XR_002501236.1 |
| NR_038930.1 | XM_021838782.1 |
| NR_038962.1 | XM_001655911.2 |
| NR_038963.1 | XM_021854890.1 |
| NR_038981.1 | XM_021842131.1 |
| NR_038996.1 | XR_002501729.1 |
| NR_040009.2 | XM_021843014.1 |
| NR_040030.1 | XM_001654830.2 |
| NR_040040.1 | XM_021838114.1 |
| NR_040044.1 | XR_002499984.1 |
| NR_040047.1 | XM_021843433.1 |
| NR_040049.1 | XM_021857691.1 |
| NR_040067.1 | XR_002502161.1 |
| NR_040070.1 | XM_021852413.1 |
| NR_040073.1 | XM_021842708.1 |
| NR_040243.2 | XM_001657934.2 |
| NR_040244.1 | XM_021848161.1 |
| NR_040718.2 | XM_021840456.1 |
| NR_045029.1 | XM_021838012.1 |
| NR_045072.2 | XM_021849774.1 |
| NR_045114.1 | XM_021849547.1 |
| NR_045118.1 | XM_001660753.2 |
| NR_045214.1 | XM_001661235.2 |
| NR_045216.1 | XR_002500006.1 |
| NR_045481.1 | XM_001650275.2 |
| NR_045555.2 | XM_001647943.2 |

| | |
|---|---|
| NR_045582.2 | XM_021848826.1 |
| NR_045637.1 | XM_021837664.1 |
| NR_045639.2 | XM_021846582.1 |
| NR_045663.4 | XM_001650834.2 |
| NR_045684.2 | XM_001654096.2 |
| NR_045697.1 | XM_021846106.1 |
| NR_045770.2 | XM_021840477.1 |
| NR_046012.1 | XM_021840228.1 |
| NR_046056.2 | XR_002500601.1 |
| NR_046082.2 | XR_002501751.1 |
| NR_046085.1 | XM_021841195.1 |
| NR_046093.2 | XM_021844478.1 |
| NR_046099.1 | XM_021841293.1 |
| NR_046115.2 | XM_021843311.1 |
| NR_046188.5 | XM_021841627.1 |
| NR_046223.2 | XM_021853486.1 |
| NR_046231.1 | XM_001653395.2 |
| NR_046269.1 | XM_001655556.2 |
| NR_046351.1 | XM_021857063.1 |
| NR_046354.1 | XM_021845470.1 |
| NR_046407.1 | XM_021846369.1 |
| NR_046411.2 | XR_002499107.1 |
| NR_046447.1 | XM_001651898.2 |
| NR_046465.2 | XM_001651281.2 |
| NR_046536.1 | XM_001647557.2 |
| NR_046574.1 | XM_021854815.1 |
| NR_046579.1 | XM_001654662.2 |
| NR_046630.1 | XM_001656645.2 |
| NR_046668.1 | XM_021837868.1 |
| NR_046672.1 | XM_021844129.1 |
| NR_046681.1 | XM_001655738.2 |
| NR_046708.1 | XM_001661810.2 |
| NR_046743.1 | XM_021842874.1 |
| NR_046748.1 | XM_021854529.1 |
| NR_046766.1 | XM_021856182.1 |
| NR_046769.1 | XM_021840582.1 |
| NR_046848.1 | XM_021847826.1 |
| NR_047011.1 | XR_002502734.1 |
| NR_047013.1 | XM_001655372.2 |
| NR_047493.1 | XM_021842383.1 |
| NR_047499.1 | XM_021852293.1 |
| NR_047521.1 | XM_021851546.1 |
| NR_047542.1 | XR_002499665.1 |
| NR_047550.1 | XM_021853171.1 |
| NR_047565.3 | XM_021851221.1 |
| NR_047577.2 | XM_021848136.1 |
| NR_047592.2 | XM_021854958.1 |
| NR_047607.1 | XM_021840959.1 |
| NR_047618.1 | XM_011494872.2 |
| NR_047622.1 | XM_021843625.1 |

| | |
|---|---|
| NR_047654.2 | XM_021852479.1 |
| NR_047658.2 | XM_021842063.1 |
| NR_047666.1 | XM_001650273.2 |
| NR_047669.2 | XM_001657475.2 |
| NR_047689.2 | XM_021849966.1 |
| NR_048537.2 | XM_001657357.2 |
| NR_048539.2 | XM_021841726.1 |
| NR_051954.3 | XM_021850260.1 |
| NR_051955.3 | XM_001656927.3 |
| NR_051960.1 | XM_001652205.3 |
| NR_051983.1 | XM_021849796.1 |
| NR_052015.4 | XM_021851457.1 |
| NR_072987.1 | XM_021845110.1 |
| NR_073020.3 | XM_001657372.2 |
| NR_073069.1 | XM_021843544.1 |
| NR_073084.1 | XM_021842853.1 |
| NR_073108.2 | XM_001651063.2 |
| NR_073136.2 | XR_002502032.1 |
| NR_073155.1 | XM_001663562.2 |
| NR_073156.2 | XM_021847592.1 |
| NR_073179.1 | XM_001660230.2 |
| NR_073366.2 | XM_001661844.2 |
| NR_073418.1 | XM_021838938.1 |
| NR_073444.2 | XM_001662980.2 |
| NR_073488.2 | XM_021854848.1 |
| NR_073504.2 | XM_001656771.2 |
| NR_073512.1 | XM_021847552.1 |
| NR_073515.3 | XM_001651631.2 |
| NR_073516.2 | XM_021846366.1 |
| NR_073543.2 | XM_021841900.1 |
| NR_073544.2 | XM_021851953.1 |
| NR_073553.3 | XM_021842357.1 |
| NR_073560.2 | XM_001654249.2 |
| NR_073582.2 | XM_001653267.2 |
| NR_074074.2 | XM_001654234.2 |
| NR_077215.1 | XM_021857725.1 |
| NR_077225.1 | XM_021848618.1 |
| NR_077236.1 | XM_021846374.1 |
| NR_102265.2 | XM_021849540.1 |
| NR_102336.1 | XM_021845219.1 |
| NR_102347.2 | XM_021838023.1 |
| NR_102713.1 | XM_001652819.2 |
| NR_102735.1 | XM_001650092.2 |
| NR_102747.1 | XM_001659423.2 |
| NR_102753.1 | XM_021854918.1 |
| NR_102761.1 | XM_021855213.1 |
| NR_102762.1 | XM_021837436.1 |
| NR_103441.2 | XM_021845345.1 |
| NR_103456.1 | XR_002498757.1 |
| NR_103460.2 | XM_001657182.2 |

| | |
|---|---|
| NR_103462.1 | XM_001661655.2 |
| NR_103470.1 | XM_021840472.1 |
| NR_103501.2 | XM_001650043.2 |
| NR_103523.1 | XM_021853824.1 |
| NR_103739.2 | XM_021847500.1 |
| NR_103757.2 | XM_021844670.1 |
| NR_103772.1 | XM_021849062.1 |
| NR_103776.1 | XM_021839347.1 |
| NR_103782.2 | XM_001650005.2 |
| NR_103805.1 | XM_021852865.1 |
| NR_103812.1 | XM_001663554.2 |
| NR_103828.1 | XM_021852155.1 |
| NR_103829.1 | XM_001663953.2 |
| NR_103833.1 | XM_001660752.2 |
| NR_103867.2 | XM_021840036.1 |
| NR_103873.1 | XM_021840580.1 |
| NR_104037.2 | XM_021843237.1 |
| NR_104089.2 | XM_001658753.2 |
| NR_104090.2 | XM_001662900.2 |
| NR_104091.1 | XM_021855412.1 |
| NR_104104.3 | XM_001651232.2 |
| NR_104110.1 | XM_021850604.1 |
| NR_104112.1 | XM_021849687.1 |
| NR_104116.1 | XM_021849978.1 |
| NR_104128.2 | XM_021852217.1 |
| NR_104129.1 | XR_002499736.1 |
| NR_104130.1 | XR_002500106.1 |
| NR_104169.3 | XM_001659282.2 |
| NR_104176.1 | XM_001660635.2 |
| NR_104195.2 | XM_001661236.2 |
| NR_104220.2 | XM_021840156.1 |
| NR_104222.2 | XM_021851366.1 |
| NR_104234.2 | XR_002500930.1 |
| NR_104242.2 | XM_021855847.1 |
| NR_104267.2 | XM_001656048.2 |
| NR_104442.2 | XM_021856571.1 |
| NR_104454.1 | XR_002502054.1 |
| NR_104455.2 | XR_002500170.1 |
| NR_104456.2 | XM_021855022.1 |
| NR_104457.2 | XM_021855409.1 |
| NR_104584.2 | XM_001651474.3 |
| NR_104585.2 | XM_001661374.2 |
| NR_104592.2 | XM_021854847.1 |
| NR_104607.1 | XM_021837957.1 |
| NR_104612.2 | XM_021849063.1 |
| NR_104632.1 | XM_021854691.1 |
| NR_104639.2 | XM_021843967.1 |
| NR_104675.1 | XM_021840177.1 |
| NR_104676.2 | XM_001664154.2 |
| NR_105009.1 | XR_002502908.1 |

| | |
|---|---|
| NR_105016.1 | XM_001655632.2 |
| NR_105056.2 | XM_021840675.1 |
| NR_108022.1 | XM_021843553.1 |
| NR_108058.1 | XM_021852485.1 |
| NR_108063.1 | XM_021849539.1 |
| NR_108070.1 | XM_001659543.2 |
| NR_108094.1 | XM_021840581.1 |
| NR_108097.1 | XM_021849014.1 |
| NR_109780.1 | XM_001653704.2 |
| NR_109805.1 | XM_001652271.2 |
| NR_109807.2 | XM_021852936.1 |
| NR_109808.2 | XM_021839004.1 |
| NR_109826.2 | XM_021846414.1 |
| NR_109836.1 | XM_021839174.1 |
| NR_109840.2 | XM_021857072.1 |
| NR_109846.1 | XM_021840805.1 |
| NR_109850.1 | XM_021840588.1 |
| NR_109859.1 | XM_021855257.1 |
| NR_109865.1 | XM_021845906.1 |
| NR_109871.2 | XM_001647652.2 |
| NR_109875.1 | XM_021845769.1 |
| NR_109881.1 | XM_021841789.1 |
| NR_109901.2 | XM_021857126.1 |
| NR_109917.1 | XM_021843824.1 |
| NR_109933.2 | XM_021848052.1 |
| NR_109950.1 | XM_021837931.1 |
| NR_109953.1 | XM_021841686.1 |
| NR_109978.2 | XM_021855013.1 |
| NR_109997.1 | XM_021854995.1 |
| NR_110032.1 | XM_021854152.1 |
| NR_110033.1 | XM_021850199.1 |
| NR_110034.2 | XM_021842798.1 |
| NR_110043.1 | XM_021844537.1 |
| NR_110052.1 | XM_021848793.1 |
| NR_110085.1 | XM_021839528.1 |
| NR_110088.1 | XM_001664144.2 |
| NR_110120.1 | XM_011495023.2 |
| NR_110139.1 | XM_001650603.2 |
| NR_110173.2 | XM_021853462.1 |
| NR_110183.1 | XM_021842849.1 |
| NR_110190.2 | XM_021849967.1 |
| NR_110195.1 | XM_021840053.1 |
| NR_110201.1 | XM_021839252.1 |
| NR_110278.1 | XM_001652614.2 |
| NR_110280.3 | XM_021838461.1 |
| NR_110303.1 | XM_021845296.1 |
| NR_110305.1 | XM_001648580.2 |
| NR_110328.3 | XM_021840270.1 |
| NR_110433.1 | XR_002501990.1 |
| NR_110434.1 | XM_021855916.1 |

| | |
|---|---|
| NR_110591.1 | XM_021856628.1 |
| NR_110606.1 | XM_021857538.1 |
| NR_110622.1 | XM_021842031.1 |
| NR_110651.1 | XM_021856181.1 |
| NR_110654.1 | XM_021856433.1 |
| NR_110718.1 | XM_021844421.1 |
| NR_110723.1 | XM_021840473.1 |
| NR_110729.1 | XM_021854153.1 |
| NR_110755.1 | XM_021844482.1 |
| NR_110770.2 | XM_021844643.1 |
| NR_110796.1 | XR_002499808.1 |
| NR_110812.1 | XM_021856752.1 |
| NR_110824.1 | XM_021854692.1 |
| NR_110852.1 | XM_021837381.1 |
| NR_110867.1 | XM_021845086.1 |
| NR_110874.1 | XM_021840474.1 |
| NR_110880.1 | XM_021850489.1 |
| NR_110883.1 | XM_001658443.2 |
| NR_110917.1 | XM_021845725.1 |
| NR_110945.1 | XM_021853267.1 |
| NR_110946.1 | XM_021857183.1 |
| NR_110948.1 | XM_021857174.1 |
| NR_110951.1 | XM_021850548.1 |
| NR_110991.2 | XM_021846232.1 |
| NR_110997.1 | XM_021843235.1 |
| NR_111001.1 | XM_001662198.2 |
| NR_117097.1 | XM_021857302.1 |
| NR_120308.1 | XR_002500325.1 |
| NR_120311.1 | XM_021846748.1 |
| NR_120326.1 | XM_001655244.2 |
| NR_120331.1 | XM_021840054.1 |
| NR_120343.1 | XM_021839170.1 |
| NR_120363.1 | XM_021841846.1 |
| NR_120392.1 | XR_002499755.1 |
| NR_120470.1 | XM_021849068.1 |
| NR_120495.1 | XM_021837379.1 |
| NR_120508.1 | XM_021845108.1 |
| NR_120513.1 | XM_001655633.2 |
| NR_120520.1 | XM_001663551.2 |
| NR_120527.1 | XM_001663544.2 |
| NR_120531.2 | XM_021839935.1 |
| NR_120539.1 | XM_021855258.1 |
| NR_120546.1 | XM_021852991.1 |
| NR_120547.1 | XM_021841697.1 |
| NR_120574.1 | XM_021846849.1 |
| NR_120580.1 | XM_021849969.1 |
| NR_120599.1 | XM_001651950.2 |
| NR_120609.1 | XM_021852456.1 |
| NR_120670.1 | XM_021845292.1 |
| NR_121187.1 | XM_021840192.1 |

| | |
|---|---|
| NR_121192.1 | XM_001652160.2 |
| NR_121564.1 | XM_021843404.1 |
| NR_121568.2 | XM_021848374.1 |
| NR_121577.1 | XM_001663538.2 |
| NR_121605.1 | XM_021837590.1 |
| NR_121612.1 | XM_021850202.1 |
| NR_121635.1 | XM_021843964.1 |
| NR_121649.1 | XM_001658838.2 |
| NR_121654.1 | XM_021852668.1 |
| NR_121661.1 | XM_021839280.1 |
| NR_121668.1 | XM_021845392.1 |
| NR_122112.1 | XM_021855544.1 |
| NR_123738.2 | XM_021855237.1 |
| NR_125330.2 | XM_001661510.2 |
| NR_125363.1 | XM_021845156.1 |
| NR_125367.1 | XM_001658073.2 |
| NR_125384.1 | XM_001661714.2 |
| NR_125426.1 | XM_001648398.2 |
| NR_125724.1 | XM_021856533.1 |
| NR_125729.1 | XM_001660113.2 |
| NR_125732.2 | XM_021852455.1 |
| NR_125763.1 | XM_021852306.1 |
| NR_125810.1 | XM_021857175.1 |
| NR_125821.1 | XM_021843741.1 |
| NR_125826.1 | XM_021854297.1 |
| NR_125848.1 | XM_021857340.1 |
| NR_125849.1 | XM_001659911.2 |
| NR_125858.1 | XM_001654567.2 |
| NR_125859.1 | XM_021840340.1 |
| NR_125860.1 | XM_021842925.1 |
| NR_125880.1 | XM_021842428.1 |
| NR_125881.1 | XM_021837830.1 |
| NR_125921.1 | XM_021847503.1 |
| NR_125932.1 | XM_021843440.1 |
| NR_125962.1 | XM_021838552.1 |
| NR_125970.1 | XM_021849060.1 |
| NR_125976.1 | XM_021849777.1 |
| NR_125995.1 | XM_001651422.2 |
| NR_126020.1 | XM_021842907.1 |
| NR_126025.1 | XM_021841861.1 |
| NR_126037.2 | XM_021840782.1 |
| NR_126054.1 | XR_002499913.1 |
| NR_126062.1 | XM_021837628.1 |
| NR_126168.1 | XM_021855436.1 |
| NR_126331.1 | XM_021854957.1 |
| NR_126333.1 | XM_021848283.1 |
| NR_126374.1 | XM_021846225.1 |
| NR_126406.1 | XM_021843140.1 |
| NR_126453.1 | XM_021855668.1 |
| NR_126500.1 | XM_001657363.2 |

| | |
|---|---|
| NR_126523.1 | XM_021838421.1 |
| NR_126561.1 | XM_021837632.1 |
| NR_130126.2 | XM_021853589.1 |
| NR_130130.2 | XM_021850595.1 |
| NR_130149.2 | XM_021841107.1 |
| NR_130728.1 | XM_021841088.1 |
| NR_130761.2 | XM_021849028.1 |
| NR_130766.1 | XM_021845674.1 |
| NR_130771.1 | XM_021855433.1 |
| NR_130772.1 | XM_001663302.2 |
| NR_130773.1 | XM_001661701.2 |
| NR_130927.2 | XM_001651268.2 |
| NR_130940.1 | XM_021852453.1 |
| NR_131184.1 | XR_002502046.1 |
| NR_131340.2 | XM_021841569.1 |
| NR_131753.2 | XM_021849545.1 |
| NR_131774.2 | XM_001651496.2 |
| NR_132102.1 | XM_021841118.1 |
| NR_132273.1 | XM_021852450.1 |
| NR_132338.1 | XM_021857299.1 |
| NR_132373.1 | XM_021838849.1 |
| NR_132380.1 | XM_021845062.1 |
| NR_132408.1 | XM_021844644.1 |
| NR_132415.1 | XR_002499788.1 |
| NR_132647.2 | XM_001657915.2 |
| NR_132740.2 | XM_021856089.1 |
| NR_132780.1 | XM_021857594.1 |
| NR_132970.1 | XM_021857702.1 |
| NR_132987.1 | XM_021843796.1 |
| NR_133566.1 | XR_002502830.1 |
| NR_133636.2 | XM_001647913.2 |
| NR_133637.2 | XR_002500924.1 |
| NR_133914.1 | XR_002502264.1 |
| NR_133940.1 | XM_021840465.1 |
| NR_133947.1 | XR_002499756.1 |
| NR_134238.1 | XM_001658099.2 |
| NR_134254.1 | XM_021856066.1 |
| NR_134266.1 | XM_021842624.1 |
| NR_134273.1 | XM_021843023.1 |
| NR_134275.1 | XM_021839068.1 |
| NR_134295.1 | XR_002502490.1 |
| NR_134299.2 | XM_021841620.1 |
| NR_134305.1 | XM_021846172.1 |
| NR_134332.1 | XM_021850646.1 |
| NR_134459.1 | XM_001658379.2 |
| NR_134464.2 | XM_001660215.2 |
| NR_134475.2 | XM_021840883.1 |
| NR_134529.2 | XM_021844154.1 |
| NR_134563.3 | XM_021843743.1 |
| NR_134577.1 | XM_001662574.2 |

| | |
|---|---|
| NR_134583.1 | XM_021838166.1 |
| NR_134598.1 | XM_001657919.2 |
| NR_134611.1 | XM_021840858.1 |
| NR_134615.1 | XM_021854160.1 |
| NR_134632.1 | XM_001656434.2 |
| NR_134637.1 | XR_002500208.1 |
| NR_134669.1 | XM_021838978.1 |
| NR_134872.2 | XM_001654840.2 |
| NR_134873.1 | XM_021844933.1 |
| NR_134878.1 | XM_021848353.1 |
| NR_134881.1 | XM_021846192.1 |
| NR_134893.2 | XM_021838399.1 |
| NR_134896.2 | XM_021854831.1 |
| NR_134924.1 | XM_001660507.2 |
| NR_134931.1 | XM_021846067.1 |
| NR_134976.3 | XM_001650806.2 |
| NR_134989.2 | XM_021847047.1 |
| NR_135049.1 | XM_021837579.1 |
| NR_135068.2 | XM_021846229.1 |
| NR_135076.1 | XM_021846171.1 |
| NR_135096.1 | XM_021845905.1 |
| NR_135108.1 | XM_001659080.2 |
| NR_135127.1 | XM_001657986.2 |
| NR_135132.1 | XM_001652414.2 |
| NR_135157.2 | XM_021843417.1 |
| NR_135166.2 | XM_001651335.2 |
| NR_135170.1 | XM_021855067.1 |
| NR_135174.1 | XM_021847985.1 |
| NR_135176.1 | XM_001650166.2 |
| NR_135203.1 | XR_002499232.1 |
| NR_135224.1 | XM_021843071.1 |
| NR_135238.1 | XM_001662279.2 |
| NR_135267.3 | XM_021854501.1 |
| NR_135481.1 | XM_021847374.1 |
| NR_135486.2 | XM_021846041.1 |
| NR_135504.1 | XM_001649177.2 |
| NR_135527.1 | XM_001663340.2 |
| NR_135532.1 | XM_021853310.1 |
| NR_135537.1 | XM_021843261.1 |
| NR_135551.1 | XM_021847052.1 |
| NR_135557.1 | XM_021853884.1 |
| NR_135616.1 | XM_021846948.1 |
| NR_135643.1 | XR_002499516.1 |
| NR_135653.2 | XM_001652763.2 |
| NR_135666.1 | XM_021857114.1 |
| NR_135674.1 | XM_021857717.1 |
| NR_135682.1 | XM_021838427.1 |
| NR_135687.1 | XM_021838168.1 |
| NR_135758.2 | XM_021847931.1 |
| NR_135764.1 | XM_021849799.1 |

| | |
|---|---|
| NR_135807.2 | XM_021855852.1 |
| NR_135828.1 | XM_021843748.1 |
| NR_135847.2 | XM_001651427.2 |
| NR_135855.2 | XM_001650246.2 |
| NR_136174.2 | XR_002501952.1 |
| NR_136178.1 | XM_021857719.1 |
| NR_136195.1 | XM_021851505.1 |
| NR_136201.1 | XM_001660350.2 |
| NR_136230.2 | XM_021841672.1 |
| NR_136242.2 | XM_021840294.1 |
| NR_136245.1 | XM_021846745.1 |
| NR_136253.1 | XM_021843794.1 |
| NR_136256.1 | XR_002503246.1 |
| NR_136272.1 | XM_001655872.3 |
| NR_136276.2 | XM_021849963.1 |
| NR_136290.1 | XM_021854990.1 |
| NR_136300.1 | XM_021852192.1 |
| NR_136301.1 | XR_002502101.1 |
| NR_136318.1 | XR_002499965.1 |
| NR_136331.3 | XM_021856688.1 |
| NR_136403.1 | XM_021855919.1 |
| NR_136518.1 | XM_021837919.1 |
| NR_136552.1 | XM_021850766.1 |
| NR_136565.2 | XM_021845726.1 |
| NR_136569.2 | XM_021845323.1 |
| NR_136586.2 | XR_002499531.1 |
| NR_136635.2 | XM_001663091.2 |
| NR_136644.1 | XM_001649286.2 |
| NR_136645.2 | XM_001649197.2 |
| NR_136700.2 | XM_021857330.1 |
| NR_136708.2 | XM_021839061.1 |
| NR_136719.2 | XM_021844128.1 |
| NR_136721.2 | XM_001661898.2 |
| NR_136746.2 | XM_021845503.1 |
| NR_136748.1 | XM_021839891.1 |
| NR_137185.1 | XM_021846330.1 |
| NR_137189.1 | XM_001663326.2 |
| NR_137194.1 | XM_021845388.1 |
| NR_137281.2 | XM_001658958.2 |
| NR_137425.1 | XM_021842631.1 |
| NR_137431.1 | XM_021850607.1 |
| NR_138035.2 | XM_021842982.1 |
| NR_138070.2 | XM_021846422.1 |
| NR_138073.2 | XM_021853912.1 |
| NR_138076.2 | XM_021851801.1 |
| NR_138077.2 | XM_021837965.1 |
| NR_138091.1 | XM_001660119.2 |
| NR_138116.2 | XR_002500293.1 |
| NR_138121.2 | XM_001649105.2 |
| NR_138123.2 | XM_021853913.1 |

| | |
|---|---|
| NR_138130.2 | XM_001652027.2 |
| NR_138256.2 | XM_011494848.2 |
| NR_138259.1 | XM_001654687.2 |
| NR_138426.2 | XM_001656014.2 |
| NR_138467.2 | XM_021857384.1 |
| NR_138609.2 | XM_001651600.2 |
| NR_138612.2 | XM_001658187.2 |
| NR_144335.2 | XM_001654383.2 |
| NR_144339.2 | XM_021843622.1 |
| NR_144380.2 | XM_021855430.1 |
| NR_144399.2 | XM_021842870.1 |
| NR_144524.2 | XM_021849733.1 |
| NR_144530.1 | XM_021841568.1 |
| NR_144540.1 | XM_021839859.1 |
| NR_144635.2 | XM_001653367.2 |
| NR_144759.2 | XM_021848790.1 |
| NR_144943.2 | XM_021851711.1 |
| NR_145129.1 | XM_021849561.1 |
| NR_145441.2 | XM_021855402.1 |
| NR_145449.1 | XM_021853822.1 |
| NR_145469.1 | XM_021855328.1 |
| NR_145502.2 | XM_021850999.1 |
| NR_145677.1 | XM_021855401.1 |
| NR_145683.2 | XR_002500107.1 |
| NR_145684.2 | XM_021847487.1 |
| NR_145685.2 | XM_001663392.2 |
| NR_145697.1 | XM_021844302.1 |
| NR_145704.1 | XM_001653015.2 |
| NR_145706.1 | XM_011495220.2 |
| NR_145819.1 | XM_021846552.1 |
| NR_145823.1 | XM_001660882.2 |
| NR_146091.2 | XM_021851712.1 |
| NR_146100.2 | XM_001660575.2 |
| NR_146109.1 | XM_021855426.1 |
| NR_146119.1 | XM_001649843.2 |
| NR_146128.2 | XM_021842961.1 |
| NR_146134.2 | XM_011494916.2 |
| NR_146143.3 | XM_021839711.1 |
| NR_146145.2 | XM_021848310.1 |
| NR_146167.2 | XM_021857671.1 |
| NR_146174.2 | XM_021853210.1 |
| NR_146176.2 | XM_021844272.1 |
| NR_146223.1 | XR_002501800.1 |
| NR_146233.2 | XM_021854223.1 |
| NR_146275.1 | XM_021844863.1 |
| NR_146279.1 | XM_021845917.1 |
| NR_146285.2 | XM_001649679.2 |
| NR_146312.1 | XM_021844653.1 |
| NR_146323.1 | XR_002498996.1 |
| NR_146341.2 | XM_021840063.1 |

| | |
|---|---|
| NR_146342.2 | XM_021852253.1 |
| NR_146384.1 | XM_001648269.2 |
| NR_146438.1 | XM_001654837.2 |
| NR_146439.1 | XM_021839892.1 |
| NR_146442.1 | XM_021845282.1 |
| NR_146456.1 | XM_021850943.1 |
| NR_146465.2 | XR_002498746.1 |
| NR_146476.1 | XM_021851556.1 |
| NR_146504.1 | XM_021844455.1 |
| NR_146507.1 | XR_002500980.1 |
| NR_146512.2 | XM_021846676.1 |
| NR_146588.1 | XM_001658961.2 |
| NR_146592.1 | XM_021840888.1 |
| NR_146598.2 | XM_021851363.1 |
| NR_146647.1 | XM_021849033.1 |
| NR_146717.1 | XM_021849778.1 |
| NR_146720.2 | XM_021838327.1 |
| NR_146733.1 | XM_001657543.2 |
| NR_146735.1 | XM_021850252.1 |
| NR_146736.1 | XM_021850257.1 |
| NR_146740.2 | XM_021844316.1 |
| NR_146763.2 | XM_001658078.2 |
| NR_146769.1 | XM_021849429.1 |
| NR_146771.1 | XM_021844900.1 |
| NR_146778.2 | XM_021855428.1 |
| NR_146866.1 | XM_021839786.1 |
| NR_146868.1 | XM_021857373.1 |
| NR_146879.2 | XM_021843932.1 |
| NR_146896.1 | XM_021839710.1 |
| NR_146905.1 | XM_021845162.1 |
| NR_146908.1 | XM_021847378.1 |
| NR_146957.1 | XM_021846677.1 |
| NR_146960.1 | XM_021855484.1 |
| NR_146965.2 | XM_021847287.1 |
| NR_146980.2 | XM_021845053.1 |
| NR_146984.3 | XM_021838323.1 |
| NR_147004.1 | XM_021851713.1 |
| NR_147021.2 | XM_021846679.1 |
| NR_147052.2 | XM_021845218.1 |
| NR_147060.2 | XM_001653111.2 |
| NR_147061.1 | XM_021853316.1 |
| NR_147081.2 | XM_021850418.1 |
| NR_147085.2 | XM_001663525.2 |
| NR_147095.2 | XM_021839856.1 |
| NR_147132.2 | XR_002499404.1 |
| NR_147136.2 | XM_021846202.1 |
| NR_147170.1 | XM_001653553.2 |
| NR_147171.2 | XM_001661539.2 |
| NR_147175.1 | XM_001652665.2 |
| NR_147186.1 | XM_001656168.2 |

| | |
|---|---|
| NR_147255.2 | XM_021851849.1 |
| NR_147408.1 | XM_021849911.1 |
| NR_147452.2 | XM_001650224.2 |
| NR_147506.1 | XM_021840886.1 |
| NR_147507.1 | XM_001659537.2 |
| NR_147791.2 | XM_021845325.1 |
| NR_147844.1 | XM_021842225.1 |
| NR_147910.2 | XM_001662162.2 |
| NR_147924.2 | XM_021838344.1 |
| NR_147928.2 | XM_021849438.1 |
| NR_147942.2 | XM_021848523.1 |
| NR_147986.1 | XM_021837689.1 |
| NR_148002.2 | XM_001653080.2 |
| NR_148203.1 | XM_021857661.1 |
| NR_148241.1 | XM_001655945.2 |
| NR_148367.1 | XM_001654671.2 |
| NR_148373.1 | XM_021846751.1 |
| NR_148384.2 | XM_021849218.1 |
| NR_148413.2 | XM_021853311.1 |
| NR_148440.2 | XM_001658397.2 |
| NR_148447.1 | XM_021847933.1 |
| NR_148469.2 | XM_021852179.1 |
| NR_148478.2 | XM_021838610.1 |
| NR_148508.1 | XM_021844806.1 |
| NR_148511.2 | XM_021845000.1 |
| NR_148921.2 | XM_001652876.3 |
| NR_148977.1 | XM_021856829.1 |
| NR_148984.2 | XM_021854441.1 |
| NR_148987.2 | XM_021847634.1 |
| NR_148995.1 | XM_021841532.1 |
| NR_149033.1 | XM_021850610.1 |
| NR_149038.1 | XM_001649562.2 |
| NR_149039.1 | XM_001656886.2 |
| NR_149061.2 | XM_001659323.2 |
| NR_149074.1 | XR_002500933.1 |
| NR_149086.1 | XM_001657661.2 |
| NR_149093.2 | XM_001656371.2 |
| NR_149106.1 | XM_001658843.2 |
| NR_149109.1 | XM_021853946.1 |
| NR_149165.2 | XM_021843144.1 |
| NR_151483.1 | XM_001655388.2 |
| NR_152570.1 | XR_002502226.1 |
| NR_152576.1 | XM_001657559.2 |
| NR_152746.1 | XM_021843898.1 |
| NR_152799.1 | XM_021847818.1 |
| NR_152811.1 | XR_002501025.1 |
| NR_152813.1 | XM_021848530.1 |
| NR_152870.1 | XM_021845037.1 |
| NR_156490.1 | XM_001656162.2 |
| NR_156732.1 | XM_021843072.1 |

| | |
|---|---|
| NR_157086.1 | XM_021844274.1 |
| NR_157112.2 | XM_021845533.1 |
| NR_157587.1 | XM_021847288.1 |
| NR_157804.1 | XM_021839989.1 |
| NR_157807.1 | XM_001658951.2 |
| NR_157824.1 | XM_021846840.1 |
| NR_157826.1 | XM_021850498.1 |
| NR_157842.1 | XM_021853725.1 |
| NR_158184.1 | XM_021843770.1 |
| NR_158193.1 | XM_021844016.1 |
| NR_158197.1 | XM_021837907.1 |
| NR_158215.1 | XM_021854685.1 |
| NR_158567.1 | XM_021843636.1 |
| NR_158629.1 | XM_021843934.1 |
| NR_158768.1 | XR_002502055.1 |
| NR_158770.1 | XM_021840149.1 |
| NR_158772.1 | XM_021845324.1 |
| NR_158778.1 | XM_021848437.1 |
| NR_159362.2 | XM_021841300.1 |
| NR_159369.2 | XM_021844525.1 |
| NR_159371.1 | XM_021847453.1 |
| NR_159376.2 | XM_021853137.1 |
| NR_159386.1 | XM_021840508.1 |
| NR_159387.1 | XM_001650559.2 |
| NR_159403.1 | XM_021851855.1 |
| NR_159737.1 | XM_021846935.1 |
| NR_159738.1 | XM_021846345.1 |
| NR_159806.1 | XM_001663167.2 |
| NR_159941.1 | XM_001650162.2 |
| NR_159966.1 | XM_021852500.1 |
| NR_160296.2 | XM_021846495.1 |
| NR_160413.1 | XM_021856074.1 |
| NR_160426.1 | XR_002500245.1 |
| NR_160551.1 | XM_021845344.1 |
| NR_160657.1 | XM_021840896.1 |
| NR_160705.1 | XM_021857300.1 |
| NR_160708.1 | XM_001663119.2 |
| NR_160718.1 | XM_021854075.1 |
| NR_160743.1 | XM_001657747.2 |
| NR_160745.1 | XR_002500815.1 |
| NR_160777.1 | XM_011494841.2 |
| NR_160778.1 | XM_021841862.1 |
| NR_160783.1 | XM_021838467.1 |
| NR_160915.1 | XR_002499038.1 |
| NR_160979.1 | XR_002499811.1 |
| NR_160980.1 | XM_021853109.1 |
| NR_161186.1 | XM_021845959.1 |
| NR_161203.1 | XM_021853258.1 |
| NR_161233.1 | XM_001657351.2 |
| NR_161292.1 | XM_001653290.2 |

| | |
|---|---|
| NR_161371.1 | XM_021849428.1 |
| NR_161435.2 | XM_021853950.1 |
| NR_161466.1 | XM_021854801.1 |
| NR_161467.1 | XM_021841061.1 |
| NR_163149.1 | XM_001655108.2 |
| NR_163195.1 | XM_021851619.1 |
| NR_163241.1 | XM_021838095.1 |
| NR_163266.1 | XM_021837438.1 |
| NR_163392.1 | XR_002498665.1 |
| NR_163410.1 | XM_021855765.1 |
| NR_163466.1 | XM_021841283.1 |
| NR_163481.1 | XM_021850559.1 |
| NR_163889.1 | XR_002500979.1 |
| NR_163917.1 | XM_021847796.1 |
| NR_163981.1 | XM_021847767.1 |
| NR_163993.1 | XM_021847665.1 |
| NR_164009.1 | XM_021847504.1 |
| NR_164155.1 | XR_002502193.1 |
| NR_164337.1 | XM_021843164.1 |
| NR_164338.1 | XR_002499922.1 |
| NR_164659.1 | XM_001659258.2 |
| NR_164700.2 | XM_021839995.1 |
| NR_164706.1 | XM_001656401.2 |
| NR_164715.1 | XM_001660646.2 |
| NR_164732.1 | XM_021839712.1 |
| NR_164747.1 | XM_021841813.1 |
| NR_164753.1 | XM_021851436.1 |
| NR_164767.1 | XM_021837754.1 |
| NR_164774.1 | XM_021849817.1 |
| NR_164779.1 | XM_021837278.1 |
| NR_164783.1 | XM_021857704.1 |
| NR_164787.1 | XM_001659861.2 |
| NR_164790.1 | XM_021852960.1 |
| NR_164810.1 | XM_021840182.1 |
| NR_164822.1 | XM_001662731.1 |
| NR_165108.1 | XR_002500884.1 |
| NR_165111.1 | XR_002500542.1 |
| NR_165115.1 | XM_001653675.2 |
| NR_165140.1 | XM_021843746.1 |
| NR_165235.1 | XM_021844470.1 |
| NR_165295.1 | XM_021842851.1 |
| NR_165336.1 | XM_001663729.2 |
| NR_165344.1 | XM_021845471.1 |
| NR_165367.1 | XM_021839249.1 |
| NR_165400.1 | XM_001660345.2 |
| NR_165401.1 | XM_021844825.1 |
| NR_165409.1 | XM_021843769.1 |
| NR_165431.1 | XR_002500024.1 |
| NR_165439.1 | XM_021840806.1 |
| NR_165630.1 | XM_021842879.1 |

| | |
|---|---|
| NR_165650.1 | XM_001654242.2 |
| NR_166069.1 | XM_021838796.1 |
| NR_166196.1 | XM_021848452.1 |
| NR_166825.1 | XM_021839000.1 |
| NR_166844.1 | XM_021851013.1 |
| NR_167706.1 | XM_021846430.1 |
| NR_167709.1 | XM_021841651.1 |
| NR_167744.1 | XM_021853731.1 |
| NR_167745.1 | XM_021849066.1 |
| NR_167888.1 | XM_021850913.1 |
| NR_167907.1 | XM_021845030.1 |
| NR_167910.1 | XM_021847870.1 |
| NR_167931.1 | XM_001648285.2 |
| NR_167974.1 | XM_021856023.1 |
| NR_167991.1 | XM_021853243.1 |
| NR_168050.1 | XM_001650796.2 |
| NR_168136.1 | XM_021839469.1 |
| NR_168365.1 | XM_021843432.1 |
| NR_168385.1 | XM_021849217.1 |
| NR_168386.1 | XM_021838017.1 |
| NR_168437.1 | XM_021851617.1 |
| NR_168445.1 | XM_001658196.2 |
| NR_168454.1 | XM_001661563.2 |
| NR_168520.1 | XM_021844472.1 |
| NR_169221.1 | XM_021856787.1 |
| NR_169231.1 | XM_021841576.1 |
| NR_169511.1 | XR_002499182.1 |
| NR_169524.1 | XM_001660542.2 |
| NR_169541.1 | XM_021856912.1 |
| NR_169542.1 | XM_021839841.1 |
| NR_169543.1 | XM_021857721.1 |
| NR_169570.1 | XM_021845021.1 |
| NR_169578.1 | XM_021857563.1 |
| NR_169585.1 | XM_021846046.1 |
| NR_169586.1 | XM_021857240.1 |
| NR_169640.1 | XM_021839813.1 |
| NR_169733.1 | XM_021854856.1 |
| NR_169740.1 | XM_021838712.1 |
| NR_169752.1 | XM_021856811.1 |
| NR_169768.1 | XM_021852501.1 |
| NR_169790.1 | XM_021846332.1 |
| NR_169835.1 | XM_021840348.1 |
| NR_169839.1 | XM_021843611.1 |
| NR_169846.1 | XM_001661082.2 |
| NR_169859.1 | XM_021857433.1 |
| NR_169880.1 | XM_021839956.1 |
| NR_170172.1 | XM_001651860.2 |
| NR_170230.1 | XM_021839451.1 |
| NR_170244.1 | XM_021849577.1 |
| NR_170268.1 | XM_021854699.1 |

| | |
|---|---|
| NR_170274.1 | XM_021841924.1 |
| NR_170318.1 | XM_021845947.1 |
| NR_170357.1 | XM_021842046.1 |
| NR_170364.1 | XM_021837874.1 |
| NR_170571.1 | XM_021840135.1 |
| NR_170584.1 | XM_021844541.1 |
| NR_170613.1 | XM_021838649.1 |
| NR_170627.1 | XM_021850618.1 |
| NR_170680.1 | XM_001660494.2 |
| NR_170691.1 | XM_021840772.1 |
| NR_170709.1 | XR_002501689.1 |
| NR_170867.1 | XM_021839483.1 |
| NR_170886.1 | XM_021845399.1 |
| NR_170903.1 | XM_021843091.1 |
| NR_170907.1 | XM_021843403.1 |
| NR_170928.1 | XM_021851367.1 |
| NR_170930.1 | XM_021840292.1 |
| NR_170960.1 | XM_021848241.1 |
| NR_170966.1 | XM_021854074.1 |
| NR_170970.1 | XM_001658848.3 |
| NR_170978.1 | XM_001658253.2 |
| NR_170982.1 | XM_001655641.2 |
| NR_171035.1 | XM_001649981.2 |
| NR_171164.1 | XM_021851732.1 |
| XM_005245005.2 | XM_021840865.1 |
| XM_005245024.2 | XM_021839225.1 |
| XM_005245048.3 | XM_021845054.1 |
| XM_005245267.4 | XR_002501237.1 |
| XM_005245314.2 | XM_021848813.1 |
| XM_005245325.4 | XM_001647527.2 |
| XM_005245348.4 | XM_001650229.2 |
| XM_005245453.1 | XM_021854905.1 |
| XM_005245508.3 | XM_021844511.1 |
| XM_005245532.4 | XM_021849802.1 |
| XM_005246141.4 | XM_021846877.1 |
| XM_005246212.3 | XM_021838098.1 |
| XM_005246255.2 | XM_021854294.1 |
| XM_005246446.3 | XM_021844651.1 |
| XM_005246456.1 | XM_001662815.3 |
| XM_005246489.4 | XM_021840595.1 |
| XM_005246505.2 | XR_002502118.1 |
| XM_005246593.2 | XM_021847949.1 |
| XM_005246812.2 | XM_001654563.2 |
| XM_005247248.5 | XM_021853173.1 |
| XM_005247284.3 | XM_021842497.1 |
| XM_005247309.2 | XM_021850190.1 |
| XM_005247347.4 | XM_021847442.1 |
| XM_005247372.4 | XM_001654939.2 |
| XM_005247458.5 | XM_001654938.2 |
| XM_005247512.1 | XM_001654024.2 |

| | |
|---|---|
| XM_005247538.3 | XM_021840885.1 |
| XM_005247594.5 | XM_021844123.1 |
| XM_005247982.3 | XM_021848058.1 |
| XM_005248028.5 | XM_001654401.2 |
| XM_005248060.1 | XM_001657332.2 |
| XM_005248231.3 | XM_021843150.1 |
| XM_005248376.4 | XM_021856056.1 |
| XM_005248504.4 | XM_021845286.1 |
| XM_005248612.3 | XR_002501564.1 |
| XM_005248700.3 | XM_021841149.1 |
| XM_005248843.4 | XM_001656857.2 |
| XM_005248893.3 | XM_021843930.1 |
| XM_005248901.3 | XM_001663312.2 |
| XM_005248951.4 | XM_021843163.1 |
| XM_005248976.1 | XM_021842696.1 |
| XM_005248999.2 | XM_001655768.2 |
| XM_005249010.2 | XM_001652794.2 |
| XM_005249079.2 | XM_001659206.2 |
| XM_005249184.5 | XR_002499135.1 |
| XM_005249218.1 | XM_001654379.2 |
| XM_005249295.1 | XM_001649170.2 |
| XM_005249315.3 | XM_021838392.1 |
| XM_005249353.4 | XM_021854168.1 |
| XM_005249403.3 | XM_021840813.1 |
| XM_005249531.1 | XM_021842080.1 |
| XM_005249584.3 | XM_021857505.1 |
| XM_005249729.1 | XM_021838138.1 |
| XM_005249883.5 | XM_021853250.1 |
| XM_005249916.1 | XM_021844524.1 |
| XM_005249952.4 | XM_001650451.2 |
| XM_005250129.4 | XM_011494761.2 |
| XM_005250405.2 | XM_021847156.1 |
| XM_005250493.1 | XM_021846640.1 |
| XM_005250502.2 | XM_021846625.1 |
| XM_005250509.4 | XM_021849043.1 |
| XM_005250574.3 | XM_021839293.1 |
| XM_005250772.3 | XM_021840520.1 |
| XM_005250773.3 | XM_021846341.1 |
| XM_005250978.3 | XM_021857210.1 |
| XM_005251025.5 | XM_021854518.1 |
| XM_005251040.4 | XM_001659611.2 |
| XM_005251121.2 | XM_021853073.1 |
| XM_005251135.4 | XM_001656668.2 |
| XM_005251590.1 | XM_021839454.1 |
| XM_005251783.3 | XM_001655274.2 |
| XM_005251826.2 | XM_001657991.2 |
| XM_005251973.4 | XM_021851973.1 |
| XM_005251987.4 | XM_021841098.1 |
| XM_005252067.4 | XM_021838165.1 |
| XM_005252151.1 | XM_021851805.1 |

| | |
|---|---|
| XM_005252208.2 | XM_021843929.1 |
| XM_005252369.3 | XM_021849614.1 |
| XM_005252627.3 | XM_011494970.2 |
| XM_005252669.3 | XM_021855759.1 |
| XM_005252787.2 | XM_021841302.1 |
| XM_005252788.2 | XM_021842642.1 |
| XM_005252921.3 | XM_021845887.1 |
| XM_005253138.5 | XM_001655520.3 |
| XM_005253168.3 | XM_001657469.2 |
| XM_005253196.3 | XM_021843145.1 |
| XM_005253229.2 | XR_002502336.1 |
| XM_005253334.3 | XM_021856426.1 |
| XM_005253698.4 | XM_001657918.2 |
| XM_005253824.3 | XM_021847490.1 |
| XM_005253834.4 | XR_002499409.1 |
| XM_005253908.4 | XM_021852100.1 |
| XM_005253944.4 | XM_021848814.1 |
| XM_005254235.3 | XM_021848750.1 |
| XM_005254430.5 | XM_021852815.1 |
| XM_005254481.3 | XM_021841230.1 |
| XM_005254529.4 | XM_021838873.1 |
| XM_005254646.2 | XR_002501692.1 |
| XM_005254671.2 | XM_001648890.2 |
| XM_005254685.4 | XM_021856005.1 |
| XM_005254795.5 | XM_001647909.2 |
| XM_005254916.3 | XM_021854058.1 |
| XM_005254934.4 | XM_021854807.1 |
| XM_005255077.1 | XM_001657110.2 |
| XM_005255182.3 | XM_001657964.2 |
| XM_005255210.2 | XM_021846340.1 |
| XM_005255304.4 | XM_021849484.1 |
| XM_005255539.3 | XM_021854377.1 |
| XM_005255558.2 | XM_021845964.1 |
| XM_005255725.5 | XM_021857593.1 |
| XM_005255741.4 | XM_021841811.1 |
| XM_005255952.5 | XM_021852659.1 |
| XM_005256255.2 | XM_021852445.1 |
| XM_005256399.5 | XM_001655822.2 |
| XM_005256424.2 | XM_001660628.2 |
| XM_005256629.1 | XM_021838872.1 |
| XM_005256678.5 | XM_021841355.1 |
| XM_005256770.1 | XM_021851593.1 |
| XM_005256784.4 | XR_002499353.1 |
| XM_005256856.3 | XM_021856553.1 |
| XM_005256866.5 | XM_021844322.1 |
| XM_005256905.2 | XM_001652148.2 |
| XM_005257019.1 | XM_001661173.2 |
| XM_005257076.3 | XM_001654141.2 |
| XM_005257098.3 | XM_021856918.1 |
| XM_005257126.4 | XM_001649603.2 |

| | |
|---|---|
| XM_005257161.3 | XR_002501855.1 |
| XM_005257187.4 | XM_021843643.1 |
| XM_005257365.4 | XR_002499205.1 |
| XM_005257370.4 | XM_021846044.1 |
| XM_005257373.4 | XM_001656630.2 |
| XM_005257468.5 | XR_002499626.1 |
| XM_005257512.2 | XM_021852990.1 |
| XM_005257703.1 | XM_021840131.1 |
| XM_005257774.4 | XM_021850647.1 |
| XM_005257859.4 | XM_021841723.1 |
| XM_005258315.5 | XM_021852595.1 |
| XM_005258361.3 | XM_001662106.2 |
| XM_005258459.4 | XR_002502177.1 |
| XM_005258620.2 | XM_001653393.2 |
| XM_005258710.5 | XM_001647886.2 |
| XM_005258746.3 | XM_011494796.2 |
| XM_005258883.2 | XR_002500387.1 |
| XM_005259153.3 | XM_021840622.1 |
| XM_005259215.4 | XM_021849409.1 |
| XM_005259323.1 | XM_001648284.2 |
| XM_005259327.3 | XM_021848320.1 |
| XM_005259467.1 | XM_021845755.1 |
| XM_005259813.4 | XM_021848086.1 |
| XM_005259834.1 | XR_002502274.1 |
| XM_005259913.2 | XM_001658819.2 |
| XM_005259941.4 | XM_021852033.1 |
| XM_005259951.4 | XM_021838485.1 |
| XM_005260183.2 | XM_021854528.1 |
| XM_005260192.2 | XM_021854495.1 |
| XM_005260195.4 | XM_001652422.2 |
| XM_005260197.4 | XM_001663850.2 |
| XM_005260382.4 | XM_001648250.2 |
| XM_005260407.4 | XM_021840015.1 |
| XM_005260465.3 | XM_021844946.1 |
| XM_005260467.4 | XM_001654923.2 |
| XM_005260501.5 | XM_021847076.1 |
| XM_005260749.4 | XM_021841894.1 |
| XM_005260978.4 | XM_021839923.1 |
| XM_005260979.2 | XM_001651872.2 |
| XM_005260997.5 | XM_021856058.1 |
| XM_005261068.3 | XM_021851708.1 |
| XM_005261121.3 | XM_021846972.1 |
| XM_005261135.3 | XM_021841359.1 |
| XM_005261168.4 | XM_021855398.1 |
| XM_005261171.3 | XM_021847310.1 |
| XM_005261415.3 | XM_001653958.2 |
| XM_005261430.2 | XR_002500025.1 |
| XM_005261525.4 | XM_021857228.1 |
| XM_005261695.1 | XM_001653029.2 |
| XM_005261705.4 | XM_021850713.1 |

| | |
|---|---|
| XM_005261986.4 | XM_021847632.1 |
| XM_005262155.4 | XM_021857674.1 |
| XM_005262310.3 | XM_021842960.1 |
| XM_005262315.3 | XM_021857519.1 |
| XM_005262384.4 | XR_002500751.1 |
| XM_005262472.2 | XM_001656487.3 |
| XM_005262489.5 | XR_002502186.1 |
| XM_005262750.4 | XM_021857375.1 |
| XM_005262772.3 | XM_021843413.1 |
| XM_005262774.4 | XM_001655947.2 |
| XM_005262937.4 | XM_021852147.1 |
| XM_005263031.4 | XM_001656585.2 |
| XM_005263077.4 | XM_001653975.2 |
| XM_005263166.4 | XM_021844418.1 |
| XM_005263372.3 | XM_021844008.1 |
| XM_005263403.3 | XM_021840623.1 |
| XM_005263449.5 | XM_021843768.1 |
| XM_005263462.4 | XM_021852585.1 |
| XM_005263547.3 | XM_021854703.1 |
| XM_005263548.3 | XM_021842230.1 |
| XM_005263629.3 | XM_021851194.1 |
| XM_005263710.2 | XM_021847906.1 |
| XM_005263862.4 | XM_021837875.1 |
| XM_005263921.4 | XM_021852117.1 |
| XM_005264203.3 | XM_021850603.1 |
| XM_005264230.4 | XM_021856134.1 |
| XM_005264283.2 | XM_021840578.1 |
| XM_005264318.3 | XM_021854975.1 |
| XM_005264421.3 | XM_021840785.1 |
| XM_005264459.2 | XM_021847991.1 |
| XM_005264487.2 | XM_021855444.1 |
| XM_005264843.4 | XM_021847174.1 |
| XM_005264921.5 | XM_021852764.1 |
| XM_005264940.4 | XM_021845965.1 |
| XM_005264968.1 | XM_021841493.1 |
| XM_005265098.4 | XR_002501047.1 |
| XM_005265379.3 | XM_021846945.1 |
| XM_005265587.5 | XM_021853326.1 |
| XM_005265653.4 | XM_021854400.1 |
| XM_005265673.4 | XM_011495202.2 |
| XM_005265967.2 | XM_021847293.1 |
| XM_005266050.4 | XR_002499905.1 |
| XM_005266419.1 | XM_021839862.1 |
| XM_005266460.2 | XM_021853030.1 |
| XM_005266461.3 | XM_021850761.1 |
| XM_005266464.3 | XR_002499617.1 |
| XM_005266583.5 | XM_021838092.1 |
| XM_005266642.3 | XM_021839314.1 |
| XM_005266970.1 | XM_001657600.2 |
| XM_005266983.4 | XM_021850327.1 |

| | |
|---|---|
| XM_005266984.4 | XM_001652859.2 |
| XM_005267135.3 | XM_001658478.3 |
| XM_005267179.4 | XM_001656854.2 |
| XM_005267325.5 | XR_002499915.1 |
| XM_005267406.5 | XM_021837242.1 |
| XM_005267510.1 | XM_001653593.2 |
| XM_005267537.4 | XM_021841417.1 |
| XM_005267709.3 | XM_001662893.2 |
| XM_005267833.5 | XM_001663446.2 |
| XM_005267854.1 | XM_021848829.1 |
| XM_005267890.5 | XM_001652292.2 |
| XM_005267988.3 | XM_021839154.1 |
| XM_005267991.3 | XM_001655512.2 |
| XM_005268111.3 | XM_001659699.2 |
| XM_005268128.1 | XM_001652543.3 |
| XM_005268372.4 | XM_021839999.1 |
| XM_005268377.4 | XM_001652075.2 |
| XM_005268447.4 | XM_021837702.1 |
| XM_005268477.1 | XR_002500305.1 |
| XM_005268493.2 | XM_021840205.1 |
| XM_005268508.4 | XM_021842343.1 |
| XM_005268556.2 | XM_011495423.2 |
| XM_005268707.4 | XM_001657252.2 |
| XM_005268730.3 | XM_021847183.1 |
| XM_005268738.3 | XR_002502086.1 |
| XM_005268739.4 | XM_021841631.1 |
| XM_005268757.4 | XM_021852584.1 |
| XM_005268927.2 | XM_021838463.1 |
| XM_005268997.2 | XM_001650331.2 |
| XM_005269011.3 | XM_001649612.2 |
| XM_005269051.3 | XM_001647841.2 |
| XM_005269125.2 | XM_021850959.1 |
| XM_005269133.2 | XM_021844910.1 |
| XM_005269207.1 | XM_001655529.2 |
| XM_005269224.5 | XM_021856048.1 |
| XM_005269476.4 | XM_021847739.1 |
| XM_005269478.4 | XM_001660865.2 |
| XM_005269509.3 | XM_021837938.1 |
| XM_005269655.3 | XM_021837863.1 |
| XM_005269742.1 | XM_021846043.1 |
| XM_005269890.1 | XM_001661198.2 |
| XM_005270026.4 | XM_021855798.1 |
| XM_005270148.2 | XM_021845320.1 |
| XM_005270231.3 | XM_001663007.2 |
| XM_005270241.4 | XM_021840934.1 |
| XM_005270360.2 | XM_021837822.1 |
| XM_005270562.3 | XM_001658325.2 |
| XM_005270777.2 | XM_021853633.1 |
| XM_005270779.4 | XM_001653309.2 |
| XM_005271103.4 | XM_021850125.1 |

| | |
|---|---|
| XM_005271107.2 | XM_001653339.2 |
| XM_005271110.2 | XM_021841832.1 |
| XM_005271113.5 | XM_021857590.1 |
| XM_005271149.4 | XR_002501219.1 |
| XM_005271223.3 | XM_021845334.1 |
| XM_005271235.3 | XR_002500689.1 |
| XM_005271260.5 | XM_021841481.1 |
| XM_005271274.4 | XM_021839812.1 |
| XM_005271355.3 | XM_021856011.1 |
| XM_005271471.3 | XM_021839576.1 |
| XM_005271485.3 | XM_021840518.1 |
| XM_005271721.5 | XM_021842056.1 |
| XM_005271745.4 | XR_002501100.1 |
| XM_005271793.4 | XM_001657971.2 |
| XM_005271835.4 | XR_002498710.1 |
| XM_005271894.3 | XM_021851658.1 |
| XM_005272016.4 | XM_001661163.2 |
| XM_005272048.4 | XM_001662594.2 |
| XM_005272105.4 | XM_021845149.1 |
| XM_005272109.5 | XM_001663115.2 |
| XM_005272118.4 | XM_021848059.1 |
| XM_005272572.4 | XM_001658034.2 |
| XM_005273165.4 | XM_021841050.1 |
| XM_005273322.4 | XM_001658079.2 |
| XM_005273481.3 | XM_021844499.1 |
| XM_005273585.4 | XM_021850958.1 |
| XM_005273611.4 | XM_021839416.1 |
| XM_005273613.4 | XM_021849505.1 |
| XM_005273801.4 | XM_001650263.2 |
| XM_005273976.2 | XM_021852784.1 |
| XM_005274161.1 | XR_002502497.1 |
| XM_005274177.3 | XM_021852396.1 |
| XM_005274368.2 | XR_002499623.1 |
| XM_005274521.4 | XR_002499167.1 |
| XM_005274578.2 | XM_021843500.1 |
| XM_005274716.3 | XM_001656134.2 |
| XM_005274742.2 | XM_021847051.1 |
| XM_005274778.3 | XM_001658072.2 |
| XM_005276753.5 | XM_021844902.1 |
| XM_005276755.5 | XM_021841199.1 |
| XM_005276975.5 | XM_021848343.1 |
| XM_005278250.5 | XR_002500178.1 |
| XM_005278279.2 | XR_002500150.1 |
| XM_006710346.3 | XM_021842623.1 |
| XM_006710394.4 | XM_021839651.1 |
| XM_006710426.3 | XM_021843167.1 |
| XM_006710474.3 | XM_021839010.1 |
| XM_006710477.3 | XM_021838656.1 |
| XM_006710585.3 | XM_001661849.2 |
| XM_006710618.3 | XM_021855814.1 |

| | |
|---|---|
| XM_006710631.3 | XM_001664161.2 |
| XM_006710731.1 | XM_021853057.1 |
| XM_006710859.1 | XM_021852115.1 |
| XM_006710922.1 | XM_001661280.2 |
| XM_006710995.2 | XM_021839907.1 |
| XM_006711174.2 | XM_021839768.1 |
| XM_006711221.3 | XR_002502937.1 |
| XM_006711253.2 | XM_021853452.1 |
| XM_006711356.3 | XM_021857083.1 |
| XM_006711567.4 | XM_021850573.1 |
| XM_006711643.3 | XM_021838636.1 |
| XM_006711929.3 | XM_001657463.2 |
| XM_006711995.3 | XM_021848943.1 |
| XM_006712021.3 | XM_021839619.1 |
| XM_006712030.4 | XM_021844223.1 |
| XM_006712094.3 | XM_021857385.1 |
| XM_006712182.3 | XM_001654420.2 |
| XM_006712196.3 | XM_021847296.1 |
| XM_006712208.3 | XM_001651946.2 |
| XM_006712286.3 | XM_021843566.1 |
| XM_006712301.2 | XR_002500445.1 |
| XM_006712336.3 | XM_021857432.1 |
| XM_006712417.1 | XM_021840491.1 |
| XM_006712521.4 | XM_001658261.2 |
| XM_006712551.1 | XM_011494873.2 |
| XM_006712639.3 | XM_021843168.1 |
| XM_006712797.3 | XM_021849506.1 |
| XM_006712799.3 | XR_002499491.1 |
| XM_006712802.1 | XM_021846047.1 |
| XM_006712946.3 | XM_021847096.1 |
| XM_006713041.1 | XM_021854687.1 |
| XM_006713204.3 | XM_021853799.1 |
| XM_006713318.3 | XM_021851193.1 |
| XM_006713417.3 | XM_021850288.1 |
| XM_006713435.3 | XM_021845267.1 |
| XM_006713472.1 | XM_011495268.2 |
| XM_006713603.2 | XM_021853199.1 |
| XM_006713684.3 | XM_021848116.1 |
| XM_006713760.4 | XM_021843439.1 |
| XM_006713831.4 | XM_001661372.2 |
| XM_006713986.3 | XM_021854547.1 |
| XM_006714116.4 | XM_021839899.1 |
| XM_006714158.1 | XM_021841015.1 |
| XM_006714165.3 | XM_001656555.2 |
| XM_006714177.3 | XM_021840757.1 |
| XM_006714231.2 | XM_001655654.2 |
| XM_006714306.4 | XM_021857511.1 |
| XM_006714420.3 | XM_001656102.2 |
| XM_006714464.3 | XM_021857592.1 |
| XM_006714532.3 | XM_001649091.2 |

| | |
|---|---|
| XM_006714587.4 | XM_001656675.2 |
| XM_006714675.4 | XM_001651525.2 |
| XM_006714741.2 | XM_021841719.1 |
| XM_006714792.2 | XM_021838054.1 |
| XM_006714921.3 | XM_021841413.1 |
| XM_006714975.1 | XR_002500754.1 |
| XM_006715052.3 | XM_021850127.1 |
| XM_006715268.2 | XM_021840756.1 |
| XM_006715270.1 | XM_001660287.2 |
| XM_006715279.2 | XM_021851641.1 |
| XM_006715349.4 | XM_021840784.1 |
| XM_006715426.3 | XM_021844547.1 |
| XM_006715586.3 | XM_021851443.1 |
| XM_006715612.3 | XM_001652605.2 |
| XM_006715642.2 | XR_002499121.1 |
| XM_006715645.3 | XM_021854117.1 |
| XM_006715655.2 | XM_021843812.1 |
| XM_006715659.1 | XM_021843299.1 |
| XM_006715667.3 | XM_021852441.1 |
| XM_006715839.3 | XM_021843100.1 |
| XM_006716093.3 | XM_021856520.1 |
| XM_006716126.3 | XM_021854549.1 |
| XM_006716256.4 | XM_001653441.2 |
| XM_006716333.3 | XM_021848247.1 |
| XM_006716585.1 | XM_001652227.2 |
| XM_006716698.3 | XM_021843152.1 |
| XM_006716706.2 | XM_021847533.1 |
| XM_006716936.2 | XM_021842447.1 |
| XM_006716944.4 | XM_001651790.2 |
| XM_006717014.2 | XM_021851881.1 |
| XM_006717096.4 | XM_021842926.1 |
| XM_006717110.3 | XM_001657205.2 |
| XM_006717141.3 | XM_021847930.1 |
| XM_006717170.4 | XM_021845298.1 |
| XM_006717191.1 | XM_021845299.1 |
| XM_006717253.1 | XM_001657538.2 |
| XM_006717273.4 | XM_021840908.1 |
| XM_006717399.3 | XM_021849049.1 |
| XM_006717519.4 | XM_021843806.1 |
| XM_006717631.4 | XM_021851191.1 |
| XM_006717634.3 | XM_001663097.2 |
| XM_006717720.4 | XM_021839684.1 |
| XM_006717851.3 | XM_021850188.1 |
| XM_006717884.4 | XM_021844595.1 |
| XM_006717891.4 | XR_002499918.1 |
| XM_006717974.3 | XM_001654934.2 |
| XM_006718047.2 | XM_021840787.1 |
| XM_006718141.4 | XM_021843807.1 |
| XM_006718191.3 | XM_021851237.1 |
| XM_006718346.2 | XM_021853488.1 |

| | |
|---|---|
| XM_006718374.2 | XM_001650344.2 |
| XM_006718384.2 | XM_021843816.1 |
| XM_006718488.4 | XR_002501361.1 |
| XM_006718517.2 | XM_021843620.1 |
| XM_006718556.4 | XM_021841188.1 |
| XM_006718699.3 | XM_001652715.2 |
| XM_006718707.3 | XM_021843368.1 |
| XM_006718739.2 | XM_021854289.1 |
| XM_006718741.2 | XM_021849981.1 |
| XM_006718797.3 | XM_021842133.1 |
| XM_006718823.2 | XM_021843274.1 |
| XM_006718849.4 | XM_001648021.2 |
| XM_006718945.3 | XM_021848131.1 |
| XM_006719025.4 | XM_021846234.1 |
| XM_006719093.1 | XM_021838136.1 |
| XM_006719114.3 | XM_021855944.1 |
| XM_006719163.4 | XM_021844224.1 |
| XM_006719181.2 | XM_001659187.2 |
| XM_006719186.4 | XM_001664054.2 |
| XM_006719218.1 | XR_002501220.1 |
| XM_006719359.1 | XM_021847237.1 |
| XM_006719400.4 | XM_021837718.1 |
| XM_006719408.4 | XM_021853704.1 |
| XM_006719460.4 | XM_021848451.1 |
| XM_006719478.2 | XM_021840214.1 |
| XM_006719564.2 | XM_021851497.1 |
| XM_006719593.3 | XM_021856629.1 |
| XM_006719644.2 | XM_021841192.1 |
| XM_006719706.2 | XM_021854861.1 |
| XM_006719707.4 | XM_021855162.1 |
| XM_006719713.4 | XR_002499686.1 |
| XM_006719735.1 | XM_001662362.2 |
| XM_006719784.3 | XR_002502144.1 |
| XM_006719786.3 | XM_021851662.1 |
| XM_006719846.3 | XM_021852149.1 |
| XM_006719882.1 | XM_021850318.1 |
| XM_006719894.3 | XM_021837432.1 |
| XM_006719900.2 | XM_021845272.1 |
| XM_006719986.2 | XM_001649240.2 |
| XM_006719991.4 | XM_021850991.1 |
| XM_006720105.3 | XM_021843479.1 |
| XM_006720116.4 | XM_021845673.1 |
| XM_006720178.1 | XM_021842994.1 |
| XM_006720185.3 | XM_021844024.1 |
| XM_006720196.2 | XM_021843298.1 |
| XM_006720204.1 | XM_021842348.1 |
| XM_006720206.4 | XM_021843826.1 |
| XM_006720226.3 | XM_001649349.2 |
| XM_006720301.3 | XM_021856601.1 |
| XM_006720309.1 | XM_021839255.1 |

| | |
|---|---|
| XM_006720355.3 | XM_021852151.1 |
| XM_006720420.1 | XM_021840141.1 |
| XM_006720459.2 | XR_002499762.1 |
| XM_006720539.3 | XM_001659996.2 |
| XM_006720626.3 | XM_021851949.1 |
| XM_006720751.3 | XM_021845623.1 |
| XM_006720832.2 | XM_021837629.1 |
| XM_006720867.3 | XM_021844556.1 |
| XM_006720908.4 | XM_021840166.1 |
| XM_006720954.4 | XM_021838032.1 |
| XM_006720982.3 | XM_021849074.1 |
| XM_006721067.4 | XM_021853934.1 |
| XM_006721074.2 | XM_021847756.1 |
| XM_006721084.1 | XR_002501680.1 |
| XM_006721236.4 | XM_021857460.1 |
| XM_006721295.3 | XM_021846441.1 |
| XM_006721471.3 | XM_021840207.1 |
| XM_006721515.2 | XM_001649687.2 |
| XM_006721546.3 | XM_001661519.2 |
| XM_006721573.3 | XM_021856727.1 |
| XM_006721601.4 | XM_021846245.1 |
| XM_006721671.4 | XM_021846785.1 |
| XM_006721707.3 | XM_021854983.1 |
| XM_006721750.4 | XM_021840946.1 |
| XM_006721866.3 | XM_021848340.1 |
| XM_006721885.3 | XM_001662728.2 |
| XM_006721893.3 | XM_021849808.1 |
| XM_006721896.3 | XM_021838065.1 |
| XM_006721907.3 | XR_002498684.1 |
| XM_006721916.3 | XM_021850763.1 |
| XM_006721920.2 | XM_021854569.1 |
| XM_006721930.3 | XM_021857421.1 |
| XM_006722018.3 | XR_002498774.1 |
| XM_006722026.2 | XM_021855933.1 |
| XM_006722115.3 | XM_021838525.1 |
| XM_006722291.4 | XM_021847301.1 |
| XM_006722425.3 | XM_021843872.1 |
| XM_006722435.3 | XM_021842502.1 |
| XM_006722521.2 | XM_021846479.1 |
| XM_006722547.3 | XM_001654482.2 |
| XM_006722557.2 | XM_001652204.2 |
| XM_006722597.4 | XM_021848210.1 |
| XM_006722616.1 | XM_021838647.1 |
| XM_006722647.3 | XM_021847292.1 |
| XM_006722706.3 | XM_001657653.2 |
| XM_006722741.3 | XM_021843447.1 |
| XM_006722782.4 | XM_021841665.1 |
| XM_006722855.4 | XM_021839983.1 |
| XM_006722866.2 | XM_001656051.2 |
| XM_006722899.4 | XM_021844025.1 |

| | |
|---|---|
| XM_006722999.2 | XM_001654508.2 |
| XM_006723009.3 | XM_021843875.1 |
| XM_006723092.4 | XM_021840554.1 |
| XM_006723097.4 | XM_001662429.2 |
| XM_006723102.4 | XR_002498791.1 |
| XM_006723219.3 | XM_021842452.1 |
| XM_006723382.3 | XM_021840591.1 |
| XM_006723489.1 | XM_021851939.1 |
| XM_006723565.3 | XM_001657662.2 |
| XM_006723598.1 | XM_021848149.1 |
| XM_006723603.2 | XM_001648353.2 |
| XM_006723646.3 | XM_021851111.1 |
| XM_006723655.2 | XM_021849408.1 |
| XM_006723727.3 | XM_021840556.1 |
| XM_006723806.3 | XM_021839449.1 |
| XM_006723961.4 | XM_021842340.1 |
| XM_006723989.2 | XM_001654021.2 |
| XM_006724030.3 | XM_021855518.1 |
| XM_006724038.3 | XM_021838315.1 |
| XM_006724051.3 | XM_001652900.2 |
| XM_006724151.2 | XM_001655239.2 |
| XM_006724180.2 | XR_002499550.1 |
| XM_006724289.4 | XM_021848954.1 |
| XM_006724559.1 | XM_021848604.1 |
| XM_006724592.4 | XM_001648264.2 |
| XM_006724619.1 | XM_011495371.2 |
| XM_006724691.2 | XM_021853374.1 |
| XM_006724763.1 | XR_002501823.1 |
| XM_006724774.3 | XM_021857411.1 |
| XM_006724779.2 | XM_021857518.1 |
| XM_006724783.2 | XM_021849625.1 |
| XM_006724802.4 | XM_021853033.1 |
| XM_006724833.2 | XM_021842451.1 |
| XM_006724855.3 | XM_021857134.1 |
| XM_011509079.3 | XM_021851799.1 |
| XM_011509133.2 | XM_021855575.1 |
| XM_011509183.2 | XM_001649285.2 |
| XM_011509201.2 | XM_021856955.1 |
| XM_011509247.1 | XM_021844005.1 |
| XM_011509265.3 | XM_021839521.1 |
| XM_011509269.2 | XM_021854379.1 |
| XM_011509301.3 | XM_021847865.1 |
| XM_011509339.3 | XM_021843845.1 |
| XM_011509386.2 | XM_021856591.1 |
| XM_011509417.2 | XM_021855670.1 |
| XM_011509453.2 | XM_021850214.1 |
| XM_011509459.2 | XR_002499956.1 |
| XM_011509475.2 | XR_002500393.1 |
| XM_011509549.1 | XM_021854946.1 |
| XM_011509644.3 | XR_002502434.1 |

| | |
|---|---|
| XM_011509667.2 | XM_021842045.1 |
| XM_011509668.2 | XM_021844579.1 |
| XM_011509681.1 | XM_021845927.1 |
| XM_011509707.2 | XM_021837623.1 |
| XM_011509739.2 | XM_021848844.1 |
| XM_011509748.2 | XM_021846052.1 |
| XM_011509759.3 | XM_001661901.2 |
| XM_011509762.3 | XM_021846412.1 |
| XM_011509786.1 | XM_021849011.1 |
| XM_011509804.1 | XM_021839333.1 |
| XM_011509813.2 | XM_021855936.1 |
| XM_011509819.1 | XR_002500593.1 |
| XM_011509820.2 | XR_002501546.1 |
| XM_011509844.2 | XM_021837625.1 |
| XM_011509872.2 | XM_001652556.2 |
| XM_011509881.2 | XR_002500369.1 |
| XM_011509935.1 | XM_021839820.1 |
| XM_011510012.1 | XM_001652880.3 |
| XM_011510152.2 | XM_021850217.1 |
| XM_011510162.2 | XM_021856025.1 |
| XM_011510181.2 | XM_021839317.1 |
| XM_011510229.3 | XM_001651243.2 |
| XM_011510232.2 | XR_002501415.1 |
| XM_011510235.2 | XM_001654206.2 |
| XM_011510236.1 | XR_002499444.1 |
| XM_011510348.1 | XM_021845820.1 |
| XM_011510399.2 | XM_001657638.2 |
| XM_011510461.3 | XM_021849101.1 |
| XM_011510497.2 | XM_001658994.2 |
| XM_011510520.1 | XM_021850410.1 |
| XM_011510527.3 | XM_001664225.2 |
| XM_011510556.2 | XM_021855205.1 |
| XM_011510562.2 | XM_021853660.1 |
| XM_011510589.1 | XM_021847585.1 |
| XM_011510597.3 | XM_001648106.2 |
| XM_011510650.3 | XR_002502893.1 |
| XM_011510655.3 | XM_001659974.2 |
| XM_011510689.3 | XM_021856322.1 |
| XM_011510691.2 | XM_021842024.1 |
| XM_011510694.2 | XM_001659192.2 |
| XM_011510781.3 | XM_021857365.1 |
| XM_011510789.2 | XM_021839063.1 |
| XM_011510919.1 | XM_021857061.1 |
| XM_011510938.1 | XM_021849464.1 |
| XM_011511090.3 | XM_021837359.1 |
| XM_011511196.3 | XM_021839130.1 |
| XM_011511249.3 | XM_021849102.1 |
| XM_011511285.2 | XM_021840427.1 |
| XM_011511325.3 | XM_021846523.1 |
| XM_011511358.2 | XR_002501481.1 |

| | |
|---|---|
| XM_011511369.3 | XM_021839808.1 |
| XM_011511379.2 | XM_021851162.1 |
| XM_011511434.3 | XM_001652467.2 |
| XM_011511445.2 | XM_021857239.1 |
| XM_011511489.2 | XM_021857517.1 |
| XM_011511494.3 | XM_021843260.1 |
| XM_011511535.3 | XM_021843396.1 |
| XM_011511738.3 | XM_021839842.1 |
| XM_011511827.2 | XM_021845891.1 |
| XM_011511933.1 | XM_001654684.2 |
| XM_011511937.1 | XM_021850863.1 |
| XM_011511958.3 | XM_021846210.1 |
| XM_011511982.1 | XR_002498858.1 |
| XM_011511993.2 | XM_021843732.1 |
| XM_011511996.2 | XM_021854290.1 |
| XM_011512005.2 | XM_021856729.1 |
| XM_011512026.3 | XM_001658900.2 |
| XM_011512148.2 | XM_021855953.1 |
| XM_011512213.2 | XM_021854805.1 |
| XM_011512226.2 | XM_021849664.1 |
| XM_011512274.1 | XM_001662595.2 |
| XM_011512277.2 | XM_021837974.1 |
| XM_011512315.1 | XM_021849426.1 |
| XM_011512323.2 | XR_002499242.1 |
| XM_011512331.2 | XM_021846908.1 |
| XM_011512424.2 | XM_021843810.1 |
| XM_011512432.2 | XM_021848817.1 |
| XM_011512488.2 | XM_001655499.2 |
| XM_011512561.2 | XM_021853990.1 |
| XM_011512615.3 | XM_021843562.1 |
| XM_011512627.3 | XM_021857271.1 |
| XM_011512660.2 | XR_002501444.1 |
| XM_011512671.2 | XM_021856666.1 |
| XM_011512711.2 | XM_021839815.1 |
| XM_011512795.3 | XR_002500301.1 |
| XM_011512807.2 | XM_021856222.1 |
| XM_011512823.2 | XM_021845537.1 |
| XM_011512842.2 | XM_001660084.2 |
| XM_011512857.2 | XR_002499068.1 |
| XM_011512891.2 | XM_001656600.2 |
| XM_011512912.2 | XM_021846469.1 |
| XM_011512958.3 | XM_001648426.2 |
| XM_011513042.3 | XM_001663436.2 |
| XM_011513050.2 | XM_001661316.2 |
| XM_011513353.3 | XM_021845978.1 |
| XM_011513356.3 | XM_001658017.2 |
| XM_011513427.2 | XM_021839644.1 |
| XM_011513454.2 | XM_021846784.1 |
| XM_011513590.2 | XM_021838321.1 |
| XM_011513682.3 | XM_021853056.1 |

| | |
|---|---|
| XM_011513719.2 | XM_021839796.1 |
| XM_011513753.2 | XM_001651589.2 |
| XM_011513811.2 | XM_021853952.1 |
| XM_011513821.3 | XM_021843814.1 |
| XM_011513835.2 | XM_021844534.1 |
| XM_011513840.3 | XM_021843533.1 |
| XM_011513856.3 | XM_001649994.2 |
| XM_011513865.2 | XM_001652354.2 |
| XM_011513908.2 | XM_001651256.2 |
| XM_011513909.2 | XM_001660053.3 |
| XM_011513975.2 | XR_002501399.1 |
| XM_011513979.2 | XM_021854404.1 |
| XM_011514025.2 | XM_001660734.2 |
| XM_011514067.1 | XM_021851532.1 |
| XM_011514094.2 | XM_021857189.1 |
| XM_011514152.2 | XM_021838681.1 |
| XM_011514225.1 | XM_001661543.2 |
| XM_011514248.3 | XM_011495417.2 |
| XM_011514264.1 | XM_021853765.1 |
| XM_011514335.2 | XM_021845773.1 |
| XM_011514346.3 | XM_001658227.2 |
| XM_011514395.2 | XM_001648969.2 |
| XM_011514420.2 | XM_021856417.1 |
| XM_011514425.1 | XM_001651346.2 |
| XM_011514428.1 | XM_021847042.1 |
| XM_011514437.3 | XM_021855906.1 |
| XM_011514459.2 | XM_021855615.1 |
| XM_011514506.2 | XM_021843760.1 |
| XM_011514615.2 | XM_021841026.1 |
| XM_011514624.2 | XM_021854878.1 |
| XM_011514663.1 | XM_021857600.1 |
| XM_011514665.1 | XM_021841247.1 |
| XM_011514726.2 | XM_021852948.1 |
| XM_011514767.2 | XM_001657693.2 |
| XM_011514770.1 | XM_021839141.1 |
| XM_011514780.1 | XM_021843563.1 |
| XM_011514812.2 | XM_001651827.2 |
| XM_011514854.2 | XM_021842647.1 |
| XM_011514859.1 | XM_021842753.1 |
| XM_011514873.2 | XM_021855393.1 |
| XM_011514917.2 | XM_001663465.2 |
| XM_011514937.2 | XM_021854502.1 |
| XM_011514938.2 | XM_001649646.2 |
| XM_011514953.3 | XM_021842788.1 |
| XM_011515081.2 | XM_001650392.2 |
| XM_011515087.1 | XM_001648941.2 |
| XM_011515151.3 | XM_001654834.2 |
| XM_011515154.2 | XM_021852968.1 |
| XM_011515213.2 | XM_021850319.1 |
| XM_011515223.2 | XM_021847492.1 |

| | |
|---|---|
| XM_011515231.2 | XM_021842448.1 |
| XM_011515241.1 | XM_021853355.1 |
| XM_011515402.3 | XM_021845408.1 |
| XM_011515407.2 | XM_021843606.1 |
| XM_011515443.2 | XM_021842083.1 |
| XM_011515444.2 | XM_021847975.1 |
| XM_011515458.2 | XM_021851190.1 |
| XM_011515527.3 | XM_021850961.1 |
| XM_011515555.1 | XM_001649754.2 |
| XM_011515556.1 | XM_021850306.1 |
| XM_011515572.1 | XM_001651480.2 |
| XM_011515582.3 | XM_001650614.2 |
| XM_011515583.2 | XM_021850875.1 |
| XM_011515654.2 | XM_021845565.1 |
| XM_011515778.1 | XM_021845850.1 |
| XM_011515795.2 | XM_001654278.2 |
| XM_011515821.2 | XM_001660085.2 |
| XM_011515826.1 | XM_001656929.2 |
| XM_011515921.1 | XM_001657318.2 |
| XM_011515964.2 | XM_021845604.1 |
| XM_011515969.1 | XM_021841583.1 |
| XM_011515990.2 | XM_021839871.1 |
| XM_011515994.1 | XM_021845555.1 |
| XM_011516031.1 | XM_001657792.2 |
| XM_011516053.2 | XR_002498701.1 |
| XM_011516066.3 | XM_021847549.1 |
| XM_011516076.2 | XM_021856314.1 |
| XM_011516102.2 | XM_021843819.1 |
| XM_011516159.3 | XM_021846254.1 |
| XM_011516190.2 | XM_021837687.1 |
| XM_011516259.2 | XR_002498837.1 |
| XM_011516283.1 | XM_021850307.1 |
| XM_011516290.2 | XM_001655220.2 |
| XM_011516609.2 | XR_002498999.1 |
| XM_011516640.2 | XM_001661169.2 |
| XM_011516669.3 | XM_021845536.1 |
| XM_011516672.2 | XM_021849390.1 |
| XM_011516737.1 | XM_021838824.1 |
| XM_011516798.1 | XM_001661387.2 |
| XM_011516926.2 | XM_001662181.2 |
| XM_011516939.3 | XM_001653333.2 |
| XM_011516944.2 | XM_021840330.1 |
| XM_011517022.2 | XM_021853955.1 |
| XM_011517048.2 | XM_021843867.1 |
| XM_011517136.2 | XM_021845925.1 |
| XM_011517202.2 | XM_021838537.1 |
| XM_011517214.1 | XM_021847580.1 |
| XM_011517219.1 | XM_021842485.1 |
| XM_011517239.2 | XM_021853800.1 |
| XM_011517257.2 | XM_021851165.1 |

| | |
|---|---|
| XM_011517259.2 | XM_001660971.2 |
| XM_011517269.1 | XM_021848841.1 |
| XM_011517274.2 | XM_021855883.1 |
| XM_011517312.2 | XM_021856746.1 |
| XM_011517361.2 | XR_002500238.1 |
| XM_011517363.3 | XM_021845601.1 |
| XM_011517495.2 | XR_002501578.1 |
| XM_011517509.3 | XM_001656595.2 |
| XM_011517539.3 | XM_001662561.2 |
| XM_011517570.2 | XM_001662238.2 |
| XM_011517588.3 | XM_021851643.1 |
| XM_011517592.3 | XM_021841844.1 |
| XM_011517616.2 | XM_001648539.2 |
| XM_011517625.2 | XM_021846704.1 |
| XM_011517635.1 | XM_001656358.2 |
| XM_011517676.2 | XR_002502126.1 |
| XM_011517736.3 | XM_021839553.1 |
| XM_011517744.1 | XM_021838667.1 |
| XM_011517861.2 | XM_021846786.1 |
| XM_011517863.3 | XM_021840351.1 |
| XM_011517963.1 | XM_011495400.2 |
| XM_011518067.1 | XM_001660843.2 |
| XM_011518070.2 | XM_021850121.1 |
| XM_011518088.2 | XM_021847319.1 |
| XM_011518153.1 | XM_021852591.1 |
| XM_011518168.2 | XM_001663903.2 |
| XM_011518189.3 | XM_021844553.1 |
| XM_011518209.3 | XM_021848815.1 |
| XM_011518233.2 | XM_001647859.2 |
| XM_011518257.2 | XM_021847899.1 |
| XM_011518277.2 | XM_021849406.1 |
| XM_011518294.3 | XM_001655888.2 |
| XM_011518341.3 | XM_001654831.2 |
| XM_011518345.3 | XM_021851034.1 |
| XM_011518352.1 | XM_021838788.1 |
| XM_011518353.1 | XM_021853348.1 |
| XM_011518361.2 | XM_021853045.1 |
| XM_011518366.3 | XM_021852785.1 |
| XM_011518374.2 | XM_021851784.1 |
| XM_011518406.2 | XR_002500065.1 |
| XM_011518488.2 | XM_021841488.1 |
| XM_011518573.3 | XM_021846460.1 |
| XM_011518584.1 | XM_001663395.2 |
| XM_011518600.1 | XM_021854788.1 |
| XM_011518639.1 | XM_021854164.1 |
| XM_011518650.3 | XM_021848068.1 |
| XM_011518770.2 | XM_021842786.1 |
| XM_011518838.2 | XM_021843448.1 |
| XM_011518875.2 | XM_021854405.1 |
| XM_011518903.3 | XR_002501607.1 |

| | |
|---|---|
| XM_011518907.2 | XM_021856352.1 |
| XM_011518909.2 | XM_011495081.2 |
| XM_011518925.1 | XM_001655284.2 |
| XM_011518950.2 | XM_021856795.1 |
| XM_011519016.1 | XM_001663420.2 |
| XM_011519089.3 | XM_021852868.1 |
| XM_011519091.3 | XM_021847884.1 |
| XM_011519114.2 | XM_021852316.1 |
| XM_011519134.3 | XM_001659450.2 |
| XM_011519152.3 | XM_021850590.1 |
| XM_011519208.2 | XR_002499778.1 |
| XM_011519214.2 | XM_021837263.1 |
| XM_011519224.1 | XM_021856220.1 |
| XM_011519230.2 | XM_001659538.2 |
| XM_011519241.2 | XR_002502405.1 |
| XM_011519264.2 | XM_021847789.1 |
| XM_011519297.1 | XM_021850302.1 |
| XM_011519306.2 | XM_021841209.1 |
| XM_011519330.2 | XM_021840399.1 |
| XM_011519337.2 | XM_001656861.2 |
| XM_011519372.2 | XM_021837367.1 |
| XM_011519398.2 | XM_021850614.1 |
| XM_011519405.2 | XM_001663245.2 |
| XM_011519503.1 | XM_021848701.1 |
| XM_011519519.3 | XM_021845347.1 |
| XM_011519538.2 | XR_002503000.1 |
| XM_011519564.2 | XR_002501732.1 |
| XM_011519604.2 | XM_021847581.1 |
| XM_011519628.1 | XM_021852835.1 |
| XM_011519672.1 | XM_021849080.1 |
| XM_011519674.1 | XM_021841816.1 |
| XM_011519710.2 | XM_021849418.1 |
| XM_011519720.2 | XM_021851096.1 |
| XM_011519743.1 | XM_001660863.2 |
| XM_011519830.3 | XR_002501412.1 |
| XM_011519863.1 | XM_021854696.1 |
| XM_011519873.3 | XM_021846472.1 |
| XM_011519890.1 | XM_001659383.3 |
| XM_011519898.3 | XM_021848117.1 |
| XM_011519907.2 | XM_021850284.1 |
| XM_011519921.2 | XM_021840282.1 |
| XM_011519941.2 | XM_021848938.1 |
| XM_011520048.1 | XM_021848875.1 |
| XM_011520209.3 | XM_021843345.1 |
| XM_011520295.2 | XM_021856049.1 |
| XM_011520302.3 | XM_001656876.2 |
| XM_011520338.1 | XM_021839232.1 |
| XM_011520373.3 | XM_001656354.2 |
| XM_011520379.2 | XM_001654500.3 |
| XM_011520406.2 | XM_021839074.1 |

| | |
|---|---|
| XM_011520476.2 | XM_001657220.2 |
| XM_011520485.2 | XM_021854378.1 |
| XM_011520493.2 | XM_021853298.1 |
| XM_011520524.1 | XM_021856095.1 |
| XM_011520560.2 | XR_002501063.1 |
| XM_011520623.3 | XM_021840880.1 |
| XM_011520747.2 | XM_021837998.1 |
| XM_011520751.3 | XM_021848816.1 |
| XM_011520760.2 | XR_002499536.1 |
| XM_011520802.2 | XM_021857722.1 |
| XM_011520815.3 | XM_021841150.1 |
| XM_011520934.3 | XM_001656090.3 |
| XM_011520997.3 | XM_021841804.1 |
| XM_011521100.2 | XM_021839290.1 |
| XM_011521166.2 | XM_011494692.2 |
| XM_011521186.2 | XM_001649186.2 |
| XM_011521243.3 | XM_021847152.1 |
| XM_011521249.2 | XM_021855909.1 |
| XM_011521262.2 | XM_001647840.2 |
| XM_011521271.2 | XM_021847328.1 |
| XM_011521320.1 | XM_021849013.1 |
| XM_011521345.3 | XM_001656139.2 |
| XM_011521351.2 | XM_021846449.1 |
| XM_011521363.2 | XM_021855396.1 |
| XM_011521366.2 | XM_001653872.2 |
| XM_011521381.2 | XM_021846780.1 |
| XM_011521396.2 | XM_001658537.2 |
| XM_011521403.2 | XM_021847424.1 |
| XM_011521497.2 | XM_001651344.3 |
| XM_011521505.2 | XM_001650219.2 |
| XM_011521556.2 | XM_021838040.1 |
| XM_011521587.2 | XM_001652154.2 |
| XM_011521591.2 | XM_001648885.2 |
| XM_011521611.3 | XM_001661830.2 |
| XM_011521620.3 | XM_021845794.1 |
| XM_011521656.3 | XM_021846946.1 |
| XM_011521660.3 | XM_021855790.1 |
| XM_011521664.2 | XM_001658944.2 |
| XM_011521665.2 | XM_021846950.1 |
| XM_011521668.2 | XM_021846429.1 |
| XM_011521670.2 | XM_021850507.1 |
| XM_011521690.2 | XM_021855816.1 |
| XM_011521780.3 | XM_001661359.2 |
| XM_011521805.2 | XM_001661816.2 |
| XM_011521852.1 | XM_021845837.1 |
| XM_011521884.1 | XM_021841145.1 |
| XM_011521893.1 | XM_021841965.1 |
| XM_011521951.3 | XM_021842488.1 |
| XM_011521976.3 | XM_021843627.1 |
| XM_011522009.2 | XM_001662643.2 |

| | |
|---|---|
| XM_011522020.1 | XM_021839407.1 |
| XM_011522023.1 | XM_021840032.1 |
| XM_011522028.1 | XM_001664198.2 |
| XM_011522034.2 | XM_001651358.2 |
| XM_011522037.2 | XM_021848874.1 |
| XM_011522044.2 | XM_021840329.1 |
| XM_011522071.1 | XM_021852141.1 |
| XM_011522076.2 | XM_021857287.1 |
| XM_011522088.1 | XM_021845988.1 |
| XM_011522125.3 | XM_021846951.1 |
| XM_011522192.2 | XM_021847873.1 |
| XM_011522204.3 | XR_002502557.1 |
| XM_011522232.2 | XM_001650616.2 |
| XM_011522234.3 | XM_021848244.1 |
| XM_011522244.1 | XR_002500357.1 |
| XM_011522351.2 | XM_021846859.1 |
| XM_011522402.1 | XM_021854417.1 |
| XM_011522516.3 | XM_021838629.1 |
| XM_011522580.2 | XM_011494999.2 |
| XM_011522608.2 | XR_002503088.1 |
| XM_011522727.3 | XM_021847441.1 |
| XM_011522783.2 | XM_021850419.1 |
| XM_011522823.2 | XM_001649512.2 |
| XM_011522836.2 | XM_021837976.1 |
| XM_011522914.2 | XM_001661854.2 |
| XM_011522921.2 | XM_021843216.1 |
| XM_011523002.2 | XM_001658139.2 |
| XM_011523023.2 | XM_021842132.1 |
| XM_011523041.2 | XM_001648293.2 |
| XM_011523095.2 | XM_021847543.1 |
| XM_011523176.2 | XM_021840720.1 |
| XM_011523178.2 | XM_021837738.1 |
| XM_011523214.2 | XM_001662600.2 |
| XM_011523222.2 | XM_021857475.1 |
| XM_011523229.1 | XM_021854443.1 |
| XM_011523230.2 | XM_021846240.1 |
| XM_011523232.2 | XM_021849499.1 |
| XM_011523243.1 | XM_021838706.1 |
| XM_011523289.1 | XM_021846943.1 |
| XM_011523294.2 | XR_002498744.1 |
| XM_011523321.1 | XM_021843597.1 |
| XM_011523326.1 | XM_001658376.2 |
| XM_011523352.1 | XM_021842920.1 |
| XM_011523353.1 | XM_021854526.1 |
| XM_011523541.2 | XM_021840930.1 |
| XM_011523597.2 | XM_021846484.1 |
| XM_011523691.2 | XM_021851514.1 |
| XM_011523713.2 | XM_021845897.1 |
| XM_011523724.1 | XM_001648859.2 |
| XM_011523738.2 | XM_021838879.1 |

| | |
|---|---|
| XM_011523752.3 | XM_021857368.1 |
| XM_011523774.2 | XM_021849729.1 |
| XM_011523851.2 | XM_021852966.1 |
| XM_011523902.3 | XR_002501617.1 |
| XM_011523910.2 | XM_001652999.2 |
| XM_011523945.2 | XM_021846534.1 |
| XM_011523963.1 | XM_021849578.1 |
| XM_011523966.1 | XM_021844294.1 |
| XM_011523992.3 | XM_021853238.1 |
| XM_011524045.2 | XM_021848939.1 |
| XM_011524057.2 | XM_021849784.1 |
| XM_011524077.3 | XM_021851261.1 |
| XM_011524080.2 | XM_001651707.2 |
| XM_011524102.3 | XM_021844958.1 |
| XM_011524104.2 | XM_021853688.1 |
| XM_011524109.3 | XM_021857614.1 |
| XM_011524114.3 | XM_021846410.1 |
| XM_011524173.1 | XM_021856076.1 |
| XM_011524194.1 | XM_021854154.1 |
| XM_011524234.1 | XM_001661964.2 |
| XM_011524258.1 | XM_001662776.2 |
| XM_011524291.1 | XM_001649616.2 |
| XM_011524296.2 | XM_021846161.1 |
| XM_011524339.2 | XM_021841509.1 |
| XM_011524390.2 | XM_021846783.1 |
| XM_011524448.2 | XM_021839190.1 |
| XM_011524453.1 | XM_021853801.1 |
| XM_011524467.2 | XM_021844954.1 |
| XM_011524493.1 | XM_021842206.1 |
| XM_011524496.2 | XM_021852862.1 |
| XM_011524521.2 | XR_002499043.1 |
| XM_011524555.2 | XM_021837979.1 |
| XM_011524667.2 | XM_021842492.1 |
| XM_011524688.3 | XM_021844820.1 |
| XM_011524701.1 | XM_021841582.1 |
| XM_011524846.3 | XM_001660356.2 |
| XM_011524896.2 | XM_001652774.2 |
| XM_011524929.2 | XM_021843854.1 |
| XM_011524950.2 | XM_021853450.1 |
| XM_011524957.2 | XM_021850144.1 |
| XM_011525065.2 | XM_021845836.1 |
| XM_011525082.3 | XM_001663485.2 |
| XM_011525102.3 | XR_002502335.1 |
| XM_011525118.2 | XM_021842003.1 |
| XM_011525139.2 | XM_021847583.1 |
| XM_011525162.2 | XM_021844119.1 |
| XM_011525183.2 | XM_021848571.1 |
| XM_011525211.3 | XM_021853180.1 |
| XM_011525216.1 | XM_021844039.1 |
| XM_011525242.2 | XM_021841510.1 |

| | |
|---|---|
| XM_011525453.2 | XM_021842029.1 |
| XM_011525454.3 | XM_021841392.1 |
| XM_011525475.3 | XM_021844935.1 |
| XM_011525478.2 | XM_001652219.2 |
| XM_011525480.2 | XM_001659116.2 |
| XM_011525495.2 | XM_021848702.1 |
| XM_011525504.3 | XM_001661307.2 |
| XM_011525525.1 | XM_021853183.1 |
| XM_011525542.1 | XM_021850321.1 |
| XM_011525569.2 | XM_021857115.1 |
| XM_011525603.2 | XM_021853993.1 |
| XM_011525757.1 | XM_021841273.1 |
| XM_011525804.2 | XM_021848550.1 |
| XM_011525917.3 | XM_021843954.1 |
| XM_011525932.1 | XM_021848238.1 |
| XM_011525933.2 | XM_021838245.1 |
| XM_011526007.2 | XM_021854004.1 |
| XM_011526099.2 | XR_002499191.1 |
| XM_011526132.2 | XM_021848261.1 |
| XM_011526142.1 | XM_021846033.1 |
| XM_011526150.2 | XM_001662356.2 |
| XM_011526169.3 | XM_021839827.1 |
| XM_011526201.2 | XM_021837837.1 |
| XM_011526290.2 | XM_021841251.1 |
| XM_011526335.2 | XM_021853558.1 |
| XM_011526371.2 | XM_021844889.1 |
| XM_011526391.1 | XM_001650283.2 |
| XM_011526484.2 | XM_001649955.2 |
| XM_011526517.2 | XM_021842878.1 |
| XM_011526555.3 | XM_021843989.1 |
| XM_011526603.2 | XM_001658954.2 |
| XM_011526611.2 | XM_021842386.1 |
| XM_011526664.2 | XM_021838474.1 |
| XM_011526842.1 | XM_021838936.1 |
| XM_011526846.1 | XM_021846778.1 |
| XM_011526861.3 | XM_021841266.1 |
| XM_011526867.1 | XM_021853643.1 |
| XM_011526874.2 | XM_021840553.1 |
| XM_011526973.2 | XM_021854235.1 |
| XM_011526980.3 | XM_001662100.2 |
| XM_011526990.2 | XM_021848515.1 |
| XM_011527048.3 | XM_001656034.2 |
| XM_011527071.1 | XM_021855006.1 |
| XM_011527085.2 | XM_021838201.1 |
| XM_011527087.2 | XM_021838183.1 |
| XM_011527116.1 | XM_021848245.1 |
| XM_011527119.1 | XM_021853093.1 |
| XM_011527134.1 | XM_001660472.2 |
| XM_011527156.2 | XR_002500131.1 |
| XM_011527162.2 | XM_001660001.2 |

| | |
|---|---|
| XM_011527170.1 | XM_001654790.2 |
| XM_011527215.1 | XR_002498976.1 |
| XM_011527228.3 | XM_021851909.1 |
| XM_011527297.2 | XM_001659872.2 |
| XM_011527341.2 | XM_021837893.1 |
| XM_011527368.2 | XM_021846594.1 |
| XM_011527370.2 | XM_001653246.2 |
| XM_011527397.2 | XM_021846475.1 |
| XM_011527424.3 | XM_021843275.1 |
| XM_011527428.1 | XM_021839098.1 |
| XM_011527432.3 | XM_001657561.2 |
| XM_011527436.1 | XM_021844665.1 |
| XM_011527560.2 | XM_001654457.2 |
| XM_011527682.2 | XM_021849065.1 |
| XM_011527710.3 | XM_001660293.2 |
| XM_011527831.2 | XM_021840226.1 |
| XM_011527861.1 | XM_001650735.2 |
| XM_011527885.3 | XM_021850317.1 |
| XM_011527897.3 | XM_021838916.1 |
| XM_011527903.3 | XM_001654933.2 |
| XM_011527909.3 | XM_021845319.1 |
| XM_011527917.1 | XR_002500735.1 |
| XM_011527934.1 | XM_021847579.1 |
| XM_011527967.2 | XM_001650583.2 |
| XM_011527974.2 | XM_001652268.2 |
| XM_011527982.3 | XM_011495114.2 |
| XM_011528042.2 | XM_021840496.1 |
| XM_011528069.2 | XM_021848923.1 |
| XM_011528090.2 | XM_021848749.1 |
| XM_011528094.1 | XM_021854632.1 |
| XM_011528136.1 | XM_021837236.1 |
| XM_011528167.2 | XM_001650125.2 |
| XM_011528177.2 | XM_001655126.2 |
| XM_011528272.2 | XR_002499770.1 |
| XM_011528278.2 | XM_021850650.1 |
| XM_011528279.2 | XM_021846965.1 |
| XM_011528281.2 | XM_021853973.1 |
| XM_011528293.2 | XM_021853086.1 |
| XM_011528442.2 | XM_001658471.2 |
| XM_011528449.3 | XM_021841244.1 |
| XM_011528481.3 | XM_021839527.1 |
| XM_011528526.2 | XM_001647723.2 |
| XM_011528527.2 | XM_021848898.1 |
| XM_011528529.3 | XM_021839243.1 |
| XM_011528545.1 | XM_001654514.2 |
| XM_011528566.2 | XR_002499475.1 |
| XM_011528586.2 | XM_021853893.1 |
| XM_011528747.2 | XM_021840288.1 |
| XM_011528748.2 | XM_021853815.1 |
| XM_011528768.3 | XM_001660114.2 |

| | |
|---|---|
| XM_011528994.2 | XM_021857536.1 |
| XM_011528997.3 | XM_021838187.1 |
| XM_011529049.1 | XM_021841231.1 |
| XM_011529085.2 | XM_021841156.1 |
| XM_011529100.2 | XM_021842941.1 |
| XM_011529140.2 | XM_001648401.2 |
| XM_011529182.2 | XM_001658414.2 |
| XM_011529263.2 | XM_001650180.2 |
| XM_011529277.2 | XM_021845305.1 |
| XM_011529342.2 | XM_021850280.1 |
| XM_011529358.3 | XM_021849574.1 |
| XM_011529467.1 | XM_021844666.1 |
| XM_011529471.1 | XM_021848087.1 |
| XM_011529507.3 | XM_001659135.2 |
| XM_011529510.2 | XM_001663443.2 |
| XM_011529514.3 | XM_021855334.1 |
| XM_011529544.2 | XM_021847241.1 |
| XM_011529609.2 | XM_001652735.2 |
| XM_011529637.2 | XM_021840717.1 |
| XM_011529644.1 | XM_021841835.1 |
| XM_011529662.2 | XM_001657634.2 |
| XM_011529766.2 | XM_021842694.1 |
| XM_011529768.2 | XR_002502196.1 |
| XM_011529841.1 | XM_021838221.1 |
| XM_011529900.2 | XM_021841806.1 |
| XM_011529908.2 | XM_021855294.1 |
| XM_011529914.2 | XM_021849082.1 |
| XM_011529921.3 | XR_002500873.1 |
| XM_011529938.2 | XM_021857145.1 |
| XM_011529954.2 | XM_021838653.1 |
| XM_011529955.1 | XM_021851808.1 |
| XM_011529965.2 | XM_021837516.1 |
| XM_011530012.1 | XM_001658338.2 |
| XM_011530038.2 | XM_021854634.1 |
| XM_011530050.2 | XM_021841508.1 |
| XM_011530106.1 | XM_001650488.2 |
| XM_011530200.2 | XM_021848171.1 |
| XM_011530204.1 | XM_001650950.2 |
| XM_011530208.2 | XM_021843591.1 |
| XM_011530260.3 | XM_021851588.1 |
| XM_011530290.2 | XM_021842977.1 |
| XM_011530311.1 | XM_021837655.1 |
| XM_011530377.2 | XM_021855071.1 |
| XM_011530432.2 | XR_002502662.1 |
| XM_011530435.2 | XM_001657378.2 |
| XM_011530472.2 | XM_021840711.1 |
| XM_011530557.2 | XM_021849814.1 |
| XM_011530696.1 | XM_021845456.1 |
| XM_011530697.1 | XM_021857324.1 |
| XM_011530716.2 | XM_021854787.1 |

| | |
|---|---|
| XM_011530717.2 | XM_001658546.2 |
| XM_011530735.2 | XM_001652505.2 |
| XM_011530813.1 | XM_021837902.1 |
| XM_011530848.3 | XM_021837731.1 |
| XM_011530884.1 | XM_021852275.1 |
| XM_011530912.2 | XR_002499697.1 |
| XM_011530950.2 | XM_021841254.1 |
| XM_011530957.3 | XM_021857309.1 |
| XM_011530977.1 | XM_021850338.1 |
| XM_011530994.1 | XM_021839922.1 |
| XM_011531032.1 | XR_002499213.1 |
| XM_011531140.2 | XM_021848390.1 |
| XM_011531171.1 | XM_021857288.1 |
| XM_011531311.2 | XM_001649521.2 |
| XM_011531317.3 | XM_001658275.2 |
| XM_011531407.3 | XM_021845271.1 |
| XM_011531412.3 | XM_021845963.1 |
| XM_011531486.2 | XM_001654422.2 |
| XM_011531509.3 | XR_002501834.1 |
| XM_011531532.2 | XM_021843170.1 |
| XM_011531533.2 | XM_021848752.1 |
| XM_011531563.2 | XM_021855192.1 |
| XM_011531587.3 | XM_021853582.1 |
| XM_011531663.3 | XM_021851723.1 |
| XM_011531671.2 | XM_021847239.1 |
| XM_011531679.3 | XM_021857215.1 |
| XM_011531693.2 | XM_001652577.2 |
| XM_011531695.3 | XM_021839100.1 |
| XM_011531716.3 | XM_021852287.1 |
| XM_011531725.1 | XR_002501160.1 |
| XM_011531775.1 | XM_021839019.1 |
| XM_011531868.3 | XM_021856416.1 |
| XM_011531895.2 | XM_021847929.1 |
| XM_011531903.2 | XM_001655931.2 |
| XM_011531912.2 | XM_001658501.2 |
| XM_011532004.1 | XM_021838052.1 |
| XM_011532022.2 | XM_021843637.1 |
| XM_011532075.1 | XM_021841239.1 |
| XM_011532088.2 | XR_002499548.1 |
| XM_011532102.1 | XM_021850736.1 |
| XM_011532118.3 | XM_021857686.1 |
| XM_011532128.2 | XM_021846894.1 |
| XM_011532130.1 | XM_001663692.2 |
| XM_011532162.2 | XM_021854233.1 |
| XM_011532233.3 | XM_021846833.1 |
| XM_011532267.2 | XM_021850273.1 |
| XM_011532358.2 | XR_002503161.1 |
| XM_011532383.2 | XR_002503006.1 |
| XM_011532386.2 | XM_021853100.1 |
| XM_011532399.2 | XR_002499472.1 |

| | |
|---|---|
| XM_011532400.2 | XM_021857701.1 |
| XM_011532505.2 | XM_021844577.1 |
| XM_011532551.2 | XM_021846589.1 |
| XM_011532566.2 | XM_021854667.1 |
| XM_011532571.2 | XM_021852723.1 |
| XM_011532623.2 | XM_021855079.1 |
| XM_011532642.2 | XM_021843155.1 |
| XM_011532716.3 | XM_001658787.2 |
| XM_011532721.2 | XM_021846838.1 |
| XM_011532730.3 | XM_021844662.1 |
| XM_011532736.2 | XR_002500683.1 |
| XM_011532766.1 | XM_021843276.1 |
| XM_011532819.2 | XM_021845123.1 |
| XM_011532919.3 | XM_021857196.1 |
| XM_011532963.3 | XM_021846434.1 |
| XM_011533045.2 | XM_001650222.2 |
| XM_011533150.3 | XM_001654370.2 |
| XM_011533154.2 | XM_021848863.1 |
| XM_011533166.3 | XR_002501149.1 |
| XM_011533268.1 | XM_021842921.1 |
| XM_011533286.2 | XM_021851030.1 |
| XM_011533371.1 | XM_021845600.1 |
| XM_011533394.3 | XM_021850722.1 |
| XM_011533433.2 | XR_002499456.1 |
| XM_011533445.2 | XR_002500793.1 |
| XM_011533478.2 | XM_001664240.2 |
| XM_011533547.3 | XM_021851167.1 |
| XM_011533576.2 | XM_021856105.1 |
| XM_011533615.2 | XM_021848737.1 |
| XM_011533625.3 | XM_001657053.3 |
| XM_011533747.3 | XM_021838913.1 |
| XM_011533774.1 | XM_021855057.1 |
| XM_011533782.2 | XM_021855236.1 |
| XM_011533834.1 | XM_021839467.1 |
| XM_011533928.1 | XR_002502529.1 |
| XM_011533980.1 | XM_021854162.1 |
| XM_011533990.3 | XM_021847449.1 |
| XM_011533991.2 | XM_001661094.2 |
| XM_011534070.1 | XM_021848600.1 |
| XM_011534079.1 | XM_021856287.1 |
| XM_011534097.2 | XM_021851791.1 |
| XM_011534099.2 | XM_001649179.2 |
| XM_011534113.2 | XM_021852830.1 |
| XM_011534152.2 | XM_021845413.1 |
| XM_011534192.2 | XM_001651697.2 |
| XM_011534200.3 | XM_021843949.1 |
| XM_011534307.2 | XM_021857006.1 |
| XM_011534382.2 | XM_021851374.1 |
| XM_011534478.3 | XM_021839049.1 |
| XM_011534485.1 | XM_021845491.1 |

| | |
|---|---|
| XM_011534487.1 | XM_021848264.1 |
| XM_011534568.2 | XM_021850428.1 |
| XM_011534637.2 | XM_021840721.1 |
| XM_011534699.2 | XM_011494733.2 |
| XM_011534780.2 | XM_021852079.1 |
| XM_011534793.2 | XM_021854773.1 |
| XM_011534836.3 | XM_021853974.1 |
| XM_011534898.2 | XM_021846261.1 |
| XM_011534905.1 | XM_001658364.2 |
| XM_011534924.3 | XM_001661757.2 |
| XM_011534931.1 | XM_021839340.1 |
| XM_011534932.2 | XM_021848118.1 |
| XM_011534956.2 | XM_001648077.2 |
| XM_011535032.3 | XM_021848211.1 |
| XM_011535144.2 | XM_021856046.1 |
| XM_011535158.2 | XM_021852969.1 |
| XM_011535206.1 | XR_002499461.1 |
| XM_011535231.2 | XR_002499395.1 |
| XM_011535239.3 | XM_021840347.1 |
| XM_011535266.2 | XM_001652046.2 |
| XM_011535393.3 | XM_021852588.1 |
| XM_011535438.2 | XM_021847900.1 |
| XM_011535439.2 | XM_021849056.1 |
| XM_011535520.1 | XM_021856801.1 |
| XM_011535524.2 | XM_021852946.1 |
| XM_011535563.2 | XR_002502068.1 |
| XM_011535575.3 | XM_021839354.1 |
| XM_011535604.3 | XM_021847850.1 |
| XM_011535632.2 | XM_021842040.1 |
| XM_011535679.3 | XM_011495126.2 |
| XM_011535685.3 | XM_021853207.1 |
| XM_011535820.2 | XM_001651457.2 |
| XM_011535863.1 | XM_021842067.1 |
| XM_011535868.2 | XM_021846814.1 |
| XM_011535877.3 | XM_001654428.2 |
| XM_011535918.3 | XM_001653632.3 |
| XM_011535921.2 | XM_021854821.1 |
| XM_011535939.2 | XM_021846415.1 |
| XM_011535977.2 | XM_021854755.1 |
| XM_011536005.3 | XR_002500104.1 |
| XM_011536057.3 | XM_021841497.1 |
| XM_011536066.1 | XM_001658924.2 |
| XM_011536106.2 | XM_001654982.2 |
| XM_011536176.2 | XM_021847407.1 |
| XM_011536196.3 | XM_021852949.1 |
| XM_011536227.2 | XM_021843701.1 |
| XM_011536365.2 | XM_001650154.2 |
| XM_011536383.2 | XM_021855080.1 |
| XM_011536406.2 | XM_021846702.1 |
| XM_011536475.2 | XM_001653035.2 |

| | |
|---|---|
| XM_011536522.3 | XM_021847008.1 |
| XM_011536542.3 | XM_021842716.1 |
| XM_011536618.2 | XM_021840831.1 |
| XM_011536651.3 | XM_001656959.2 |
| XM_011536690.3 | XM_021850684.1 |
| XM_011536720.3 | XM_021837611.1 |
| XM_011536778.2 | XM_001657106.2 |
| XM_011536846.2 | XM_021851952.1 |
| XM_011536872.1 | XM_021846502.1 |
| XM_011536886.2 | XM_021841968.1 |
| XM_011536946.3 | XM_021844984.1 |
| XM_011536957.1 | XM_021849647.1 |
| XM_011536966.2 | XM_021839994.1 |
| XM_011537009.1 | XM_021839017.1 |
| XM_011537078.2 | XM_021838915.1 |
| XM_011537081.2 | XM_001656655.3 |
| XM_011537084.2 | XM_021851299.1 |
| XM_011537179.2 | XM_001654501.3 |
| XM_011537182.2 | XM_021842584.1 |
| XM_011537203.3 | XM_001650374.2 |
| XM_011537310.1 | XR_002499460.1 |
| XM_011537332.2 | XM_001659695.2 |
| XM_011537338.1 | XM_021849920.1 |
| XM_011537343.1 | XM_021851263.1 |
| XM_011537362.2 | XM_021854032.1 |
| XM_011537397.1 | XM_001651439.2 |
| XM_011537417.2 | XM_021841516.1 |
| XM_011537462.2 | XM_021847121.1 |
| XM_011537484.1 | XM_021853237.1 |
| XM_011537504.2 | XM_021847535.1 |
| XM_011537509.2 | XM_021852169.1 |
| XM_011537556.2 | XM_021842187.1 |
| XM_011537615.1 | XM_021846428.1 |
| XM_011537616.1 | XM_001662892.2 |
| XM_011537696.2 | XM_021852223.1 |
| XM_011537730.3 | XM_021838264.1 |
| XM_011537846.2 | XM_021841332.1 |
| XM_011537912.2 | XM_021852886.1 |
| XM_011537972.3 | XM_021844842.1 |
| XM_011537995.2 | XM_001660079.2 |
| XM_011538080.2 | XM_001658544.2 |
| XM_011538184.2 | XM_021852078.1 |
| XM_011538186.3 | XM_021857199.1 |
| XM_011538209.2 | XM_021844679.1 |
| XM_011538213.2 | XM_001650853.2 |
| XM_011538361.3 | XM_021854287.1 |
| XM_011538393.2 | XR_002503280.1 |
| XM_011538446.3 | XM_021853234.1 |
| XM_011538477.2 | XM_021841135.1 |
| XM_011538525.3 | XM_021846812.1 |

| | |
|---|---|
| XM_011538598.2 | XR_002498907.1 |
| XM_011538631.2 | XM_001660234.2 |
| XM_011538643.3 | XM_021850336.1 |
| XM_011538688.2 | XM_021847151.1 |
| XM_011538692.2 | XM_021855747.1 |
| XM_011538700.2 | XM_021841206.1 |
| XM_011538802.1 | XM_021837793.1 |
| XM_011538813.2 | XM_001660580.2 |
| XM_011538828.3 | XM_021857322.1 |
| XM_011538829.3 | XM_001660693.3 |
| XM_011538833.2 | XM_021856990.1 |
| XM_011538838.2 | XM_021841234.1 |
| XM_011538844.2 | XM_021847480.1 |
| XM_011538926.1 | XM_001651531.2 |
| XM_011538971.2 | XM_021856338.1 |
| XM_011538993.3 | XM_021838919.1 |
| XM_011539000.2 | XM_021845846.1 |
| XM_011539125.2 | XM_021845158.1 |
| XM_011539132.2 | XM_021841240.1 |
| XM_011539173.3 | XR_002501147.1 |
| XM_011539199.3 | XM_021847430.1 |
| XM_011539279.1 | XM_021854406.1 |
| XM_011539300.2 | XM_021840598.1 |
| XM_011539302.2 | XM_021839057.1 |
| XM_011539308.2 | XM_021842911.1 |
| XM_011539355.2 | XM_021839501.1 |
| XM_011539387.2 | XM_021848495.1 |
| XM_011539492.3 | XM_001654747.2 |
| XM_011539537.2 | XM_001655396.2 |
| XM_011539592.3 | XR_002502313.1 |
| XM_011539640.1 | XM_021856280.1 |
| XM_011539673.2 | XM_021843765.1 |
| XM_011539744.3 | XM_021851638.1 |
| XM_011539746.3 | XM_021847039.1 |
| XM_011539753.2 | XM_001662050.2 |
| XM_011539805.1 | XM_001649154.2 |
| XM_011539826.3 | XM_021854833.1 |
| XM_011539851.3 | XM_021853985.1 |
| XM_011539961.2 | XM_021847460.1 |
| XM_011539987.2 | XM_021840846.1 |
| XM_011540104.2 | XM_021848095.1 |
| XM_011540154.2 | XM_021842769.1 |
| XM_011540160.2 | XM_001654622.2 |
| XM_011540198.2 | XM_021851094.1 |
| XM_011540207.2 | XM_001662995.2 |
| XM_011540221.3 | XM_001652368.2 |
| XM_011540238.2 | XM_021851590.1 |
| XM_011540325.3 | XM_001656635.2 |
| XM_011540327.2 | XM_021838229.1 |
| XM_011540337.1 | XM_021853059.1 |

| | |
|---|---|
| XM_011540338.1 | XR_002500007.1 |
| XM_011540340.3 | XM_001653516.2 |
| XM_011540428.1 | XM_021855671.1 |
| XM_011540466.3 | XM_021844896.1 |
| XM_011540521.3 | XM_021849466.1 |
| XM_011540596.2 | XM_021847711.1 |
| XM_011540603.2 | XM_021853308.1 |
| XM_011540606.2 | XM_021856141.1 |
| XM_011540617.3 | XM_021840244.1 |
| XM_011540639.3 | XM_021845915.1 |
| XM_011540726.3 | XM_001660077.2 |
| XM_011540730.1 | XM_001649072.2 |
| XM_011540779.3 | XM_021855706.1 |
| XM_011540797.2 | XR_002501336.1 |
| XM_011540798.1 | XM_021843438.1 |
| XM_011540903.2 | XM_021841564.1 |
| XM_011540906.3 | XR_002498681.1 |
| XM_011540925.2 | XM_021853969.1 |
| XM_011540932.2 | XM_001648296.2 |
| XM_011540945.2 | XM_001661521.2 |
| XM_011540957.1 | XM_001650622.2 |
| XM_011540960.1 | XM_011495053.2 |
| XM_011540996.2 | XM_021856804.1 |
| XM_011541011.2 | XM_021841796.1 |
| XM_011541051.2 | XM_001649622.2 |
| XM_011541154.2 | XM_021852261.1 |
| XM_011541213.1 | XR_002499056.1 |
| XM_011541285.1 | XM_021843868.1 |
| XM_011541290.2 | XM_021855752.1 |
| XM_011541384.2 | XM_021852706.1 |
| XM_011541459.2 | XR_002501854.1 |
| XM_011541498.1 | XM_001654729.2 |
| XM_011541503.2 | XR_002503223.1 |
| XM_011541587.3 | XM_001658000.2 |
| XM_011541631.3 | XM_021846096.1 |
| XM_011541691.2 | XM_021840978.1 |
| XM_011541708.3 | XM_021843950.1 |
| XM_011541739.2 | XM_021848599.1 |
| XM_011541763.1 | XM_021842742.1 |
| XM_011541766.2 | XM_021855123.1 |
| XM_011541768.1 | XM_021842183.1 |
| XM_011541872.3 | XM_001647755.2 |
| XM_011541962.2 | XR_002500968.1 |
| XM_011541963.2 | XM_001657520.2 |
| XM_011541971.2 | XM_001663696.2 |
| XM_011541974.2 | XM_021839479.1 |
| XM_011542035.3 | XM_021848098.1 |
| XM_011542063.2 | XM_021846813.1 |
| XM_011542072.1 | XM_021838918.1 |
| XM_011542077.2 | XM_021846598.1 |

| | |
|---|---|
| XM_011542212.2 | XM_011494908.2 |
| XM_011542265.3 | XM_021841241.1 |
| XM_011542351.1 | XR_002500663.1 |
| XM_011542383.2 | XM_021837244.1 |
| XM_011542399.2 | XM_001652695.2 |
| XM_011542522.2 | XM_021854003.1 |
| XM_011542567.2 | XM_021846416.1 |
| XM_011542632.3 | XM_021846974.1 |
| XM_011542633.2 | XM_021857457.1 |
| XM_011542648.2 | XM_021837964.1 |
| XM_011542739.2 | XM_021846098.1 |
| XM_011542740.2 | XM_021853434.1 |
| XM_011542790.3 | XM_021840597.1 |
| XM_011542792.1 | XM_001648951.2 |
| XM_011542857.2 | XM_021849129.1 |
| XM_011542870.2 | XM_021840017.1 |
| XM_011542901.2 | XM_021840878.1 |
| XM_011542903.3 | XR_002501937.1 |
| XM_011542907.1 | XM_021840055.1 |
| XM_011542917.2 | XM_021837801.1 |
| XM_011543000.2 | XM_021854094.1 |
| XM_011543048.1 | XM_021849128.1 |
| XM_011543055.2 | XM_001657472.2 |
| XM_011543086.2 | XM_021840704.1 |
| XM_011543131.3 | XM_001656392.2 |
| XM_011543143.2 | XM_001653346.2 |
| XM_011543150.1 | XM_001658122.2 |
| XM_011543167.3 | XM_021844597.1 |
| XM_011543232.2 | XM_021847971.1 |
| XM_011543255.3 | XM_021850345.1 |
| XM_011543308.3 | XM_021854684.1 |
| XM_011543334.2 | XM_021854005.1 |
| XM_011543340.2 | XM_001648603.2 |
| XM_011543349.3 | XM_021846921.1 |
| XM_011543366.2 | XM_001654598.2 |
| XM_011543385.3 | XM_021842134.1 |
| XM_011543531.3 | XM_021842535.1 |
| XM_011543533.2 | XM_021837246.1 |
| XM_011543538.2 | XM_001654722.2 |
| XM_011543559.2 | XR_002500941.1 |
| XM_011543570.2 | XM_021842570.1 |
| XM_011543574.3 | XM_021840712.1 |
| XM_011543586.2 | XM_021852271.1 |
| XM_011543607.2 | XM_021837795.1 |
| XM_011543608.2 | XM_021848341.1 |
| XM_011543609.2 | XM_021839971.1 |
| XM_011543643.2 | XM_021855793.1 |
| XM_011543680.2 | XM_021856841.1 |
| XM_011543735.3 | XM_021851095.1 |
| XM_011543758.2 | XM_021842326.1 |

| | |
|---|---|
| XM_011543766.1 | XM_001662634.2 |
| XM_011543786.2 | XM_021840913.1 |
| XM_011543811.2 | XM_021853314.1 |
| XM_011543821.2 | XM_021838705.1 |
| XM_011543830.3 | XM_021848745.1 |
| XM_011543843.3 | XM_021841660.1 |
| XM_011543885.2 | XR_002498600.1 |
| XM_011544035.2 | XM_021855378.1 |
| XM_011544048.2 | XR_002502832.1 |
| XM_011544063.2 | XM_001658074.2 |
| XM_011544076.1 | XR_002501465.1 |
| XM_011544087.2 | XM_021837548.1 |
| XM_011544108.3 | XM_001658587.2 |
| XM_011544132.2 | XM_021847688.1 |
| XM_011544161.3 | XR_002500474.1 |
| XM_011544164.2 | XM_001649070.2 |
| XM_011544169.2 | XM_021850695.1 |
| XM_011544206.3 | XM_021839421.1 |
| XM_011544220.2 | XM_021839958.1 |
| XM_011544227.1 | XM_021842246.1 |
| XM_011544305.2 | XM_001650752.3 |
| XM_011544342.3 | XM_021848890.1 |
| XM_011544360.2 | XM_021851583.1 |
| XM_011544364.3 | XM_021852950.1 |
| XM_011544368.3 | XM_021847228.1 |
| XM_011544426.2 | XM_021848017.1 |
| XM_011544456.2 | XM_021838799.1 |
| XM_011544481.2 | XM_021853212.1 |
| XM_011544482.2 | XM_001662843.2 |
| XM_011544505.2 | XM_001656295.2 |
| XM_011544518.2 | XM_021856815.1 |
| XM_011544526.2 | XM_001653506.2 |
| XM_011544550.2 | XM_021854234.1 |
| XM_011544569.1 | XM_001656152.2 |
| XM_011544632.2 | XM_021838907.1 |
| XM_011544679.3 | XM_021841248.1 |
| XM_011544680.3 | XM_021855306.1 |
| XM_011544687.1 | XM_001659505.2 |
| XM_011544713.2 | XM_021855051.1 |
| XM_011544860.3 | XM_021849807.1 |
| XM_011544861.1 | XM_021843365.1 |
| XM_011544884.2 | XM_021846250.1 |
| XM_011544888.2 | XM_001648626.2 |
| XM_011544899.1 | XM_021840735.1 |
| XM_011544933.3 | XM_021843362.1 |
| XM_011544955.3 | XM_021853229.1 |
| XM_011545002.2 | XM_021843360.1 |
| XM_011545044.2 | XR_002502729.1 |
| XM_011545063.3 | XM_001655267.2 |
| XM_011545073.1 | XM_021847983.1 |

| | |
|---|---|
| XM_011545110.1 | XM_021843529.1 |
| XM_011545165.3 | XM_001653570.2 |
| XM_011545240.2 | XM_021837798.1 |
| XM_011545293.2 | XM_021842684.1 |
| XM_011545308.3 | XM_021847776.1 |
| XM_011545403.2 | XM_021841004.1 |
| XM_011545468.2 | XM_021857118.1 |
| XM_011545472.3 | XM_021843148.1 |
| XM_011545478.2 | XM_001649954.2 |
| XM_011545564.3 | XM_021838909.1 |
| XM_011545585.2 | XM_021838797.1 |
| XM_011545711.2 | XM_001652872.3 |
| XM_011545713.1 | XM_001657198.2 |
| XM_011545738.1 | XM_021838182.1 |
| XM_011545764.1 | XM_021854151.1 |
| XM_011545770.1 | XM_021843229.1 |
| XM_011545784.3 | XM_021842245.1 |
| XM_011545824.3 | XM_001660773.2 |
| XM_011545836.3 | XM_021838262.1 |
| XM_011545913.2 | XM_001655992.2 |
| XM_011546001.3 | XM_021848083.1 |
| XM_011546149.2 | XR_002502077.1 |
| XM_011546186.1 | XM_021855523.1 |
| XM_011547058.2 | XM_021847784.1 |
| XM_011547362.3 | XM_021853872.1 |
| XM_017000046.1 | XM_021850239.1 |
| XM_017000054.1 | XM_021841535.1 |
| XM_017000095.2 | XM_021850066.1 |
| XM_017000148.2 | XM_021846722.1 |
| XM_017000212.2 | XM_021849064.1 |
| XM_017000215.2 | XM_021845450.1 |
| XM_017000216.1 | XM_021853674.1 |
| XM_017000236.2 | XM_021837906.1 |
| XM_017000256.2 | XM_001663439.2 |
| XM_017000272.1 | XM_001658937.2 |
| XM_017000284.1 | XM_021850825.1 |
| XM_017000307.1 | XM_021838224.1 |
| XM_017000339.1 | XM_021838868.1 |
| XM_017000395.1 | XM_021845708.1 |
| XM_017000451.1 | XM_021848508.1 |
| XM_017000466.1 | XM_021847469.1 |
| XM_017000495.1 | XM_021837915.1 |
| XM_017000509.2 | XM_021846377.1 |
| XM_017000544.2 | XM_021846321.1 |
| XM_017000597.1 | XM_001649468.2 |
| XM_017000622.1 | XM_021852721.1 |
| XM_017000623.2 | XM_021848199.1 |
| XM_017000624.2 | XM_021841003.1 |
| XM_017000665.1 | XM_021850069.1 |
| XM_017000691.1 | XR_002498777.1 |

| | |
|---|---|
| XM_017000712.1 | XM_021843363.1 |
| XM_017000717.1 | XM_021848601.1 |
| XM_017000758.1 | XM_021845748.1 |
| XM_017000769.2 | XM_001654074.2 |
| XM_017000777.1 | XM_021852511.1 |
| XM_017000781.1 | XM_001657405.2 |
| XM_017000805.2 | XM_021841243.1 |
| XM_017000836.1 | XM_021839411.1 |
| XM_017000954.2 | XM_001657568.2 |
| XM_017000976.1 | XM_001651296.3 |
| XM_017000977.1 | XM_021857609.1 |
| XM_017000994.2 | XM_001648911.2 |
| XM_017001000.2 | XM_021851576.1 |
| XM_017001044.1 | XR_002499818.1 |
| XM_017001047.1 | XM_021850347.1 |
| XM_017001051.1 | XM_021855052.1 |
| XM_017001053.1 | XM_021851155.1 |
| XM_017001061.2 | XM_001653437.2 |
| XM_017001074.1 | XM_021852319.1 |
| XM_017001087.2 | XM_021850889.1 |
| XM_017001192.2 | XM_001662418.2 |
| XM_017001206.1 | XM_001662326.2 |
| XM_017001238.1 | XM_021843224.1 |
| XM_017001245.2 | XM_001651234.2 |
| XM_017001254.1 | XM_001661956.2 |
| XM_017001294.1 | XM_001663878.2 |
| XM_017001306.2 | XM_021853027.1 |
| XM_017001307.2 | XM_021841829.1 |
| XM_017001317.1 | XM_021848129.1 |
| XM_017001333.1 | XM_021855661.1 |
| XM_017001383.1 | XM_001660960.2 |
| XM_017001392.2 | XM_001662295.2 |
| XM_017001407.1 | XM_021853453.1 |
| XM_017001428.1 | XM_021851494.1 |
| XM_017001514.1 | XM_021849811.1 |
| XM_017001527.1 | XM_021854947.1 |
| XM_017001534.1 | XM_021840923.1 |
| XM_017001541.2 | XM_021839237.1 |
| XM_017001567.1 | XM_021849613.1 |
| XM_017001618.1 | XM_021855927.1 |
| XM_017001626.1 | XM_021840722.1 |
| XM_017001630.1 | XM_021842722.1 |
| XM_017001690.1 | XM_001655238.2 |
| XM_017001694.1 | XM_021850234.1 |
| XM_017001705.1 | XM_021842714.1 |
| XM_017001751.1 | XM_021850542.1 |
| XM_017001788.1 | XM_001654404.2 |
| XM_017001869.1 | XM_021842495.1 |
| XM_017001905.1 | XM_001655387.2 |
| XM_017001939.1 | XM_001654754.2 |

| | |
|---|---|
| XM_017002089.2 | XM_021849413.1 |
| XM_017002094.2 | XM_021838287.1 |
| XM_017002098.2 | XM_021837239.1 |
| XM_017002099.2 | XM_021847417.1 |
| XM_017002177.1 | XM_001661489.2 |
| XM_017002185.1 | XM_021857128.1 |
| XM_017002241.1 | XM_021846307.1 |
| XM_017002252.2 | XM_001651159.2 |
| XM_017002260.2 | XR_002502909.1 |
| XM_017002264.2 | XM_021844576.1 |
| XM_017002302.1 | XM_001655469.2 |
| XM_017002479.1 | XM_021842789.1 |
| XM_017002550.2 | XM_021842779.1 |
| XM_017002685.2 | XM_021852269.1 |
| XM_017002767.1 | XM_001648651.2 |
| XM_017002774.1 | XM_021839392.1 |
| XM_017002785.2 | XM_021840738.1 |
| XM_017002928.2 | XM_021848862.1 |
| XM_017002929.1 | XR_002500881.1 |
| XM_017002933.2 | XM_021840710.1 |
| XM_017002946.2 | XM_021847137.1 |
| XM_017002955.1 | XM_021849949.1 |
| XM_017002997.1 | XM_021839924.1 |
| XM_017003042.2 | XM_021847856.1 |
| XM_017003093.1 | XM_001650002.2 |
| XM_017003105.2 | XM_001656181.2 |
| XM_017003113.1 | XM_021837991.1 |
| XM_017003116.1 | XM_021845386.1 |
| XM_017003159.2 | XM_001657485.2 |
| XM_017003196.2 | XM_021838956.1 |
| XM_017003220.1 | XM_001658400.2 |
| XM_017003264.2 | XR_002499551.1 |
| XM_017003297.1 | XM_021846082.1 |
| XM_017003309.1 | XM_001655489.2 |
| XM_017003314.2 | XM_021843219.1 |
| XM_017003365.1 | XM_001651123.2 |
| XM_017003382.2 | XM_021854811.1 |
| XM_017003396.1 | XR_002499097.1 |
| XM_017003399.2 | XM_021847413.1 |
| XM_017003402.1 | XM_021839048.1 |
| XM_017003410.1 | XM_001654484.2 |
| XM_017003426.2 | XM_001647518.2 |
| XM_017003432.2 | XM_001650865.2 |
| XM_017003447.1 | XM_021838792.1 |
| XM_017003455.2 | XM_021851207.1 |
| XM_017003475.2 | XM_021839134.1 |
| XM_017003479.1 | XM_001663856.2 |
| XM_017003480.2 | XM_021853339.1 |
| XM_017003488.2 | XM_021839138.1 |
| XM_017003503.1 | XM_021837797.1 |

| | |
|---|---|
| XM_017003526.1 | XM_021842947.1 |
| XM_017003527.1 | XM_001652474.2 |
| XM_017003528.1 | XM_021856418.1 |
| XM_017003548.1 | XM_021856740.1 |
| XM_017003549.2 | XM_021852914.1 |
| XM_017003576.1 | XM_001661737.2 |
| XM_017003604.1 | XM_021854770.1 |
| XM_017003606.1 | XM_021854932.1 |
| XM_017003622.1 | XM_001650152.2 |
| XM_017003659.1 | XM_021842499.1 |
| XM_017003664.1 | XM_021850002.1 |
| XM_017003667.1 | XM_021838112.1 |
| XM_017003670.1 | XM_021846452.1 |
| XM_017003683.2 | XM_021856768.1 |
| XM_017003700.1 | XM_001654510.2 |
| XM_017003716.1 | XM_021856844.1 |
| XM_017003722.1 | XM_021841096.1 |
| XM_017003771.1 | XM_021855842.1 |
| XM_017003818.1 | XM_021855050.1 |
| XM_017003833.2 | XM_021855841.1 |
| XM_017003834.1 | XM_021847621.1 |
| XM_017003838.1 | XM_001648453.2 |
| XM_017003890.1 | XM_001649489.2 |
| XM_017003897.1 | XM_001661230.2 |
| XM_017003925.1 | XM_021852388.1 |
| XM_017003930.1 | XR_002501038.1 |
| XM_017003996.1 | XM_021856926.1 |
| XM_017004007.1 | XM_021837978.1 |
| XM_017004009.1 | XM_021848023.1 |
| XM_017004034.1 | XM_021857351.1 |
| XM_017004062.1 | XM_001664202.2 |
| XM_017004082.2 | XM_021847895.1 |
| XM_017004086.2 | XM_021846256.1 |
| XM_017004174.1 | XR_002501860.1 |
| XM_017004236.1 | XM_021848411.1 |
| XM_017004276.1 | XR_002501252.1 |
| XM_017004312.2 | XR_002498742.1 |
| XM_017004343.2 | XM_001655843.2 |
| XM_017004366.1 | XM_021838478.1 |
| XM_017004379.2 | XM_021839335.1 |
| XM_017004401.2 | XM_021843395.1 |
| XM_017004403.1 | XM_021850235.1 |
| XM_017004432.2 | XM_021856677.1 |
| XM_017004447.1 | XM_021849533.1 |
| XM_017004492.2 | XM_001657766.2 |
| XM_017004514.2 | XM_021844296.1 |
| XM_017004557.1 | XM_021844944.1 |
| XM_017004575.2 | XM_001663566.2 |
| XM_017004624.2 | XM_021847896.1 |
| XM_017004656.1 | XM_021846255.1 |

| | |
|---|---|
| XM_017004664.1 | XM_021848682.1 |
| XM_017004669.1 | XM_021852357.1 |
| XM_017004701.1 | XR_002499156.1 |
| XM_017004738.1 | XM_021842417.1 |
| XM_017004824.1 | XM_021855857.1 |
| XM_017004866.1 | XM_021852132.1 |
| XM_017004891.1 | XM_021839398.1 |
| XM_017004920.2 | XM_021857004.1 |
| XM_017004956.2 | XR_002501861.1 |
| XM_017004979.2 | XM_021853021.1 |
| XM_017004999.1 | XM_001654238.2 |
| XM_017005076.2 | XM_021842027.1 |
| XM_017005088.1 | XM_021851534.1 |
| XM_017005092.2 | XM_021846251.1 |
| XM_017005146.1 | XM_021847035.1 |
| XM_017005160.2 | XM_021847255.1 |
| XM_017005187.2 | XM_021839962.1 |
| XM_017005219.2 | XM_001655268.2 |
| XM_017005235.1 | XM_001651748.2 |
| XM_017005385.1 | XM_001659059.2 |
| XM_017005469.2 | XM_021857269.1 |
| XM_017005476.1 | XM_001659938.2 |
| XM_017005522.1 | XR_002501972.1 |
| XM_017005653.1 | XM_021847231.1 |
| XM_017005690.1 | XM_001655713.2 |
| XM_017005694.2 | XM_001650175.2 |
| XM_017005699.1 | XM_021847897.1 |
| XM_017005702.1 | XM_021839928.1 |
| XM_017005731.1 | XM_021856375.1 |
| XM_017005744.2 | XM_021843142.1 |
| XM_017005756.1 | XM_021842668.1 |
| XM_017005762.2 | XM_001661814.2 |
| XM_017005766.2 | XM_021855064.1 |
| XM_017005785.1 | XM_001652195.2 |
| XM_017005796.1 | XM_021842092.1 |
| XM_017005811.2 | XM_021850142.1 |
| XM_017005813.1 | XM_021843411.1 |
| XM_017005823.2 | XM_021838800.1 |
| XM_017005861.1 | XR_002502309.1 |
| XM_017005894.1 | XM_021847464.1 |
| XM_017005907.2 | XM_021838080.1 |
| XM_017005927.1 | XM_021851864.1 |
| XM_017005946.2 | XM_001654117.2 |
| XM_017005993.1 | XM_021838894.1 |
| XM_017006035.1 | XM_021847342.1 |
| XM_017006041.2 | XM_021855193.1 |
| XM_017006048.1 | XM_001659233.2 |
| XM_017006071.2 | XM_001650571.2 |
| XM_017006080.2 | XM_021853431.1 |
| XM_017006083.2 | XM_021849371.1 |

| | |
|---|---|
| XM_017006091.1 | XM_021857339.1 |
| XM_017006103.1 | XM_001656795.2 |
| XM_017006122.1 | XM_001661715.3 |
| XM_017006123.1 | XM_021840458.1 |
| XM_017006139.1 | XM_021850547.1 |
| XM_017006144.1 | XR_002500716.1 |
| XM_017006147.1 | XM_001650577.2 |
| XM_017006157.1 | XM_021840854.1 |
| XM_017006172.2 | XM_021852082.1 |
| XM_017006199.1 | XM_021841352.1 |
| XM_017006205.1 | XM_021851730.1 |
| XM_017006218.1 | XM_021853673.1 |
| XM_017006257.1 | XM_021855828.1 |
| XM_017006285.1 | XM_021840985.1 |
| XM_017006299.1 | XM_001652537.2 |
| XM_017006368.1 | XR_002502105.1 |
| XM_017006386.1 | XM_021847857.1 |
| XM_017006484.2 | XM_001659332.2 |
| XM_017006520.1 | XM_021846953.1 |
| XM_017006612.2 | XM_001664043.2 |
| XM_017006624.1 | XM_021848010.1 |
| XM_017006629.1 | XM_021842657.1 |
| XM_017006632.1 | XM_021838884.1 |
| XM_017006645.2 | XM_021837305.1 |
| XM_017006709.1 | XM_001657157.2 |
| XM_017006724.1 | XM_021845454.1 |
| XM_017006746.2 | XM_021851981.1 |
| XM_017006759.2 | XM_001659374.2 |
| XM_017006776.1 | XM_021855859.1 |
| XM_017006782.1 | XM_001659267.2 |
| XM_017006787.2 | XM_021853491.1 |
| XM_017006797.2 | XM_021856309.1 |
| XM_017006818.2 | XM_001656627.2 |
| XM_017006825.2 | XM_021852987.1 |
| XM_017006827.2 | XM_021855295.1 |
| XM_017006878.2 | XM_021850025.1 |
| XM_017006929.2 | XM_021847894.1 |
| XM_017006932.2 | XM_021849279.1 |
| XM_017006951.2 | XM_021840984.1 |
| XM_017006963.2 | XM_021847775.1 |
| XM_017006982.1 | XM_021845455.1 |
| XM_017007015.1 | XM_001661096.2 |
| XM_017007017.1 | XM_021855061.1 |
| XM_017007062.1 | XM_021848856.1 |
| XM_017007141.1 | XR_002501475.1 |
| XM_017007158.2 | XM_021854768.1 |
| XM_017007161.2 | XM_001652785.2 |
| XM_017007201.1 | XM_021846577.1 |
| XM_017007208.1 | XM_021855141.1 |
| XM_017007300.1 | XM_021840800.1 |

| | |
|---|---|
| XM_017007347.1 | XM_001651839.2 |
| XM_017007375.1 | XM_021844678.1 |
| XM_017007379.2 | XM_021841468.1 |
| XM_017007436.1 | XM_021838479.1 |
| XM_017007444.1 | XM_021844905.1 |
| XM_017007523.2 | XM_021849275.1 |
| XM_017007526.2 | XM_001659177.2 |
| XM_017007534.1 | XM_021851285.1 |
| XM_017007552.2 | XR_002502218.1 |
| XM_017007571.1 | XM_021837658.1 |
| XM_017007589.1 | XM_021855843.1 |
| XM_017007639.2 | XM_021844396.1 |
| XM_017007648.2 | XM_021849929.1 |
| XM_017007709.2 | XM_021854969.1 |
| XM_017007720.1 | XR_002500255.1 |
| XM_017007722.1 | XM_021853752.1 |
| XM_017007741.2 | XM_021856562.1 |
| XM_017007798.1 | XM_021855662.1 |
| XM_017007799.1 | XM_001649030.2 |
| XM_017007805.2 | XM_021850767.1 |
| XM_017007809.1 | XM_021848982.1 |
| XM_017007892.2 | XM_021851633.1 |
| XM_017007903.1 | XM_001658168.2 |
| XM_017007911.1 | XM_001664160.2 |
| XM_017007913.2 | XM_021854277.1 |
| XM_017007938.1 | XM_001660480.2 |
| XM_017007944.1 | XM_021850513.1 |
| XM_017007958.1 | XM_001661427.2 |
| XM_017007985.1 | XM_021842078.1 |
| XM_017007994.1 | XM_021851760.1 |
| XM_017008009.1 | XM_021838698.1 |
| XM_017008016.2 | XM_021840497.1 |
| XM_017008030.2 | XM_011494732.2 |
| XM_017008130.1 | XM_021850826.1 |
| XM_017008152.2 | XM_021847246.1 |
| XM_017008189.1 | XM_021849383.1 |
| XM_017008199.1 | XM_001651722.2 |
| XM_017008210.2 | XM_021842093.1 |
| XM_017008224.1 | XM_021849683.1 |
| XM_017008295.2 | XR_002500720.1 |
| XM_017008319.1 | XM_021845795.1 |
| XM_017008341.1 | XM_021847770.1 |
| XM_017008353.1 | XR_002500041.1 |
| XM_017008371.1 | XM_021842903.1 |
| XM_017008402.1 | XM_021849823.1 |
| XM_017008474.1 | XM_021842847.1 |
| XM_017008491.1 | XM_021839054.1 |
| XM_017008508.1 | XM_021848172.1 |
| XM_017008511.2 | XM_021853960.1 |
| XM_017008514.2 | XM_021839476.1 |

| | |
|---|---|
| XM_017008515.2 | XM_001660235.2 |
| XM_017008544.2 | XM_001659211.2 |
| XM_017008563.1 | XM_001663685.2 |
| XM_017008568.1 | XM_001653766.2 |
| XM_017008587.1 | XM_021856304.1 |
| XM_017008593.2 | XM_021851421.1 |
| XM_017008594.2 | XM_021843953.1 |
| XM_017008612.2 | XM_021846278.1 |
| XM_017008630.1 | XM_021854779.1 |
| XM_017008699.1 | XM_001654945.2 |
| XM_017008717.2 | XM_021854904.1 |
| XM_017008735.1 | XM_021840807.1 |
| XM_017008736.1 | XM_021852767.1 |
| XM_017008737.1 | XM_021841689.1 |
| XM_017008773.2 | XM_021847921.1 |
| XM_017008820.2 | XM_021852996.1 |
| XM_017008834.1 | XM_021849948.1 |
| XM_017008864.2 | XM_021856646.1 |
| XM_017008884.1 | XM_021838492.1 |
| XM_017008888.1 | XM_021848378.1 |
| XM_017008929.2 | XM_021852652.1 |
| XM_017008947.2 | XM_001649964.2 |
| XM_017008980.2 | XM_021855827.1 |
| XM_017008984.2 | XM_001658236.2 |
| XM_017008985.2 | XM_001655297.2 |
| XM_017009007.1 | XM_021851896.1 |
| XM_017009014.1 | XM_021854302.1 |
| XM_017009019.2 | XM_021848167.1 |
| XM_017009037.2 | XR_002500488.1 |
| XM_017009043.2 | XM_021840709.1 |
| XM_017009085.1 | XM_021842574.1 |
| XM_017009097.2 | XM_021842681.1 |
| XM_017009112.2 | XM_001658956.2 |
| XM_017009126.2 | XM_021857164.1 |
| XM_017009134.1 | XM_021854225.1 |
| XM_017009149.2 | XR_002499184.1 |
| XM_017009161.2 | XM_021849372.1 |
| XM_017009168.1 | XR_002500049.1 |
| XM_017009172.1 | XM_021844845.1 |
| XM_017009186.1 | XM_021841390.1 |
| XM_017009204.1 | XM_021837971.1 |
| XM_017009241.2 | XM_021847691.1 |
| XM_017009250.2 | XM_021842442.1 |
| XM_017009263.1 | XM_021840705.1 |
| XM_017009265.1 | XR_002499259.1 |
| XM_017009293.2 | XM_001651712.2 |
| XM_017009315.2 | XM_001649882.3 |
| XM_017009322.2 | XM_021841491.1 |
| XM_017009363.1 | XM_021848473.1 |
| XM_017009439.2 | XM_001658581.2 |

| | |
|---|---|
| XM_017009449.1 | XM_021846272.1 |
| XM_017009460.1 | XM_001655335.2 |
| XM_017009469.1 | XM_021845237.1 |
| XM_017009470.2 | XM_021846808.1 |
| XM_017009473.1 | XM_021857356.1 |
| XM_017009491.1 | XM_021857473.1 |
| XM_017009497.2 | XM_001651052.2 |
| XM_017009544.2 | XM_021849209.1 |
| XM_017009621.2 | XM_021848293.1 |
| XM_017009630.1 | XM_021847685.1 |
| XM_017009657.1 | XM_001659810.2 |
| XM_017009664.1 | XM_021847854.1 |
| XM_017009725.2 | XM_001661276.2 |
| XM_017009740.1 | XM_021847576.1 |
| XM_017009766.1 | XM_001652555.2 |
| XM_017009792.2 | XM_021842318.1 |
| XM_017009814.1 | XM_021854278.1 |
| XM_017009858.2 | XR_002502574.1 |
| XM_017009869.1 | XM_021838319.1 |
| XM_017009872.1 | XM_021853613.1 |
| XM_017009886.2 | XM_021841504.1 |
| XM_017010000.1 | XM_021838699.1 |
| XM_017010009.1 | XM_021849127.1 |
| XM_017010061.1 | XR_002499640.1 |
| XM_017010072.2 | XM_021856644.1 |
| XM_017010087.2 | XM_021845336.1 |
| XM_017010099.2 | XM_021846279.1 |
| XM_017010115.2 | XM_001657083.2 |
| XM_017010164.2 | XM_021841812.1 |
| XM_017010199.2 | XM_001663329.2 |
| XM_017010202.1 | XR_002501786.1 |
| XM_017010205.2 | XR_002499253.1 |
| XM_017010217.2 | XM_021848919.1 |
| XM_017010278.2 | XM_021857476.1 |
| XM_017010300.2 | XM_021849675.1 |
| XM_017010339.1 | XR_002499533.1 |
| XM_017010374.2 | XM_021853665.1 |
| XM_017010409.1 | XM_001660854.2 |
| XM_017010440.2 | XR_002499260.1 |
| XM_017010441.2 | XM_021843842.1 |
| XM_017010472.1 | XR_002499543.1 |
| XM_017010588.1 | XM_021848360.1 |
| XM_017010598.2 | XM_021855053.1 |
| XM_017010629.2 | XM_021850462.1 |
| XM_017010632.2 | XM_001649426.2 |
| XM_017010636.2 | XM_001650588.2 |
| XM_017010663.2 | XM_021855890.1 |
| XM_017010712.1 | XR_002499441.1 |
| XM_017010714.2 | XM_021843380.1 |
| XM_017010721.1 | XM_021842846.1 |

| | |
|---|---|
| XM_017010736.2 | XR_002502360.1 |
| XM_017010744.2 | XM_021845120.1 |
| XM_017010752.1 | XM_001653113.2 |
| XM_017010809.2 | XM_001662276.2 |
| XM_017010812.1 | XM_021840983.1 |
| XM_017010830.1 | XM_021840464.1 |
| XM_017010882.1 | XM_021837839.1 |
| XM_017010912.2 | XM_021841065.1 |
| XM_017010929.2 | XM_001656182.2 |
| XM_017010949.2 | XM_021837842.1 |
| XM_017010985.1 | XM_021849175.1 |
| XM_017011000.1 | XM_001659936.2 |
| XM_017011001.1 | XM_021839929.1 |
| XM_017011046.1 | XM_021839238.1 |
| XM_017011050.1 | XM_021853666.1 |
| XM_017011052.1 | XM_021857660.1 |
| XM_017011054.1 | XM_021848474.1 |
| XM_017011105.2 | XM_021857454.1 |
| XM_017011139.2 | XM_021857684.1 |
| XM_017011174.2 | XM_021855054.1 |
| XM_017011227.1 | XM_021855926.1 |
| XM_017011235.2 | XM_021846324.1 |
| XM_017011243.2 | XM_021851959.1 |
| XM_017011248.2 | XM_021841176.1 |
| XM_017011278.2 | XM_021850149.1 |
| XM_017011305.2 | XM_021855060.1 |
| XM_017011321.1 | XM_001661331.2 |
| XM_017011348.1 | XM_021841759.1 |
| XM_017011403.1 | XM_021840364.1 |
| XM_017011426.2 | XM_021855140.1 |
| XM_017011448.2 | XM_001654688.2 |
| XM_017011450.2 | XM_021852270.1 |
| XM_017011462.1 | XM_001662653.2 |
| XM_017011466.2 | XM_021837834.1 |
| XM_017011503.1 | XM_021841580.1 |
| XM_017011560.2 | XM_021843780.1 |
| XM_017011562.2 | XM_021840821.1 |
| XM_017011662.2 | XM_021841534.1 |
| XM_017011722.1 | XM_021837539.1 |
| XM_017011739.1 | XM_001655554.2 |
| XM_017011757.1 | XR_002502907.1 |
| XM_017011761.2 | XM_021853425.1 |
| XM_017011795.1 | XM_021840319.1 |
| XM_017011813.1 | XM_021838493.1 |
| XM_017011864.1 | XM_021848921.1 |
| XM_017011874.1 | XM_001657407.2 |
| XM_017011888.1 | XM_021839338.1 |
| XM_017011905.2 | XM_021844675.1 |
| XM_017011929.2 | XM_021846160.1 |
| XM_017011937.1 | XR_002501447.1 |

| | |
|---|---|
| XM_017011972.1 | XM_021846796.1 |
| XM_017011974.1 | XM_021853353.1 |
| XM_017012006.2 | XM_021847613.1 |
| XM_017012016.2 | XM_001658119.2 |
| XM_017012038.2 | XR_002503217.1 |
| XM_017012040.2 | XM_021854967.1 |
| XM_017012060.2 | XM_021850456.1 |
| XM_017012076.1 | XM_021840304.1 |
| XM_017012080.1 | XM_001658746.2 |
| XM_017012114.1 | XM_021837450.1 |
| XM_017012136.2 | XR_002502075.1 |
| XM_017012182.1 | XM_001652304.2 |
| XM_017012226.1 | XM_021842217.1 |
| XM_017012227.1 | XM_021842332.1 |
| XM_017012230.1 | XR_002500408.1 |
| XM_017012294.1 | XM_021855102.1 |
| XM_017012298.2 | XM_001648484.2 |
| XM_017012335.2 | XM_021849260.1 |
| XM_017012388.1 | XM_021857358.1 |
| XM_017012452.1 | XM_021845589.1 |
| XM_017012492.2 | XM_021852536.1 |
| XM_017012506.2 | XM_001655132.2 |
| XM_017012540.1 | XM_021847855.1 |
| XM_017012616.2 | XM_001656766.2 |
| XM_017012659.1 | XM_001662583.2 |
| XM_017012738.1 | XR_002502146.1 |
| XM_017012746.1 | XM_021843347.1 |
| XM_017012750.2 | XM_001654267.2 |
| XM_017012760.2 | XM_001648588.3 |
| XM_017012802.1 | XM_021838630.1 |
| XM_017012820.1 | XM_001661368.2 |
| XM_017012845.2 | XR_002502380.1 |
| XM_017012878.2 | XM_001655326.2 |
| XM_017012938.1 | XM_021844603.1 |
| XM_017012944.1 | XM_021851154.1 |
| XM_017012959.1 | XR_002500655.1 |
| XM_017012994.2 | XM_001659696.2 |
| XM_017012998.2 | XM_021853161.1 |
| XM_017013021.2 | XM_021838837.1 |
| XM_017013048.1 | XM_021852074.1 |
| XM_017013171.1 | XM_021846995.1 |
| XM_017013263.1 | XM_021847277.1 |
| XM_017013268.2 | XM_021841269.1 |
| XM_017013289.1 | XM_021857002.1 |
| XM_017013306.1 | XM_001652431.3 |
| XM_017013310.2 | XM_001653853.2 |
| XM_017013323.1 | XM_021849907.1 |
| XM_017013373.1 | XM_021849132.1 |
| XM_017013376.2 | XR_002498740.1 |
| XM_017013415.2 | XM_021837961.1 |

| | |
|---|---|
| XM_017013416.1 | XM_021850135.1 |
| XM_017013428.2 | XM_021850349.1 |
| XM_017013467.2 | XM_021854344.1 |
| XM_017013516.1 | XM_021847006.1 |
| XM_017013535.1 | XM_021856630.1 |
| XM_017013567.2 | XM_021843745.1 |
| XM_017013576.1 | XM_021854080.1 |
| XM_017013604.2 | XM_001658131.2 |
| XM_017013645.1 | XM_021844589.1 |
| XM_017013656.2 | XM_021845351.1 |
| XM_017013693.1 | XM_021852492.1 |
| XM_017013705.2 | XR_002502518.1 |
| XM_017013755.1 | XM_021846842.1 |
| XM_017013780.1 | XM_021855499.1 |
| XM_017013824.2 | XM_021854079.1 |
| XM_017013856.2 | XM_021847113.1 |
| XM_017013905.1 | XM_021856133.1 |
| XM_017013934.1 | XM_021843531.1 |
| XM_017013942.1 | XM_021852816.1 |
| XM_017013979.2 | XM_001657762.2 |
| XM_017013984.1 | XM_021841581.1 |
| XM_017013995.2 | XM_021852997.1 |
| XM_017014001.2 | XM_021838809.1 |
| XM_017014028.2 | XM_021837364.1 |
| XM_017014091.1 | XM_021857388.1 |
| XM_017014117.2 | XM_001651905.2 |
| XM_017014171.1 | XM_021838980.1 |
| XM_017014182.1 | XM_021843539.1 |
| XM_017014194.1 | XR_002502393.1 |
| XM_017014222.1 | XM_021849133.1 |
| XM_017014237.2 | XM_021845951.1 |
| XM_017014244.2 | XM_021856026.1 |
| XM_017014268.1 | XM_021846423.1 |
| XM_017014269.2 | XM_021844886.1 |
| XM_017014279.1 | XM_021839286.1 |
| XM_017014300.1 | XM_021851208.1 |
| XM_017014340.1 | XM_021856868.1 |
| XM_017014428.1 | XM_021837616.1 |
| XM_017014447.1 | XM_001650388.2 |
| XM_017014501.2 | XM_021838652.1 |
| XM_017014508.1 | XR_002499829.1 |
| XM_017014517.2 | XM_021853924.1 |
| XM_017014548.2 | XR_002502368.1 |
| XM_017014557.1 | XM_021856620.1 |
| XM_017014568.1 | XM_021839933.1 |
| XM_017014571.1 | XR_002503208.1 |
| XM_017014586.1 | XM_001650147.2 |
| XM_017014588.1 | XM_001651459.2 |
| XM_017014594.2 | XM_021853003.1 |
| XM_017014597.2 | XM_001650268.2 |

| | |
|---|---|
| XM_017014666.2 | XM_021847924.1 |
| XM_017014671.1 | XM_021852538.1 |
| XM_017014720.1 | XM_001662585.2 |
| XM_017014753.2 | XM_021839240.1 |
| XM_017014792.1 | XM_021848475.1 |
| XM_017014803.1 | XM_021841075.1 |
| XM_017014804.1 | XM_021841853.1 |
| XM_017014943.2 | XM_021840533.1 |
| XM_017014951.2 | XM_001657073.2 |
| XM_017014969.2 | XM_001661150.2 |
| XM_017015139.2 | XM_021853344.1 |
| XM_017015213.1 | XM_001657231.2 |
| XM_017015225.1 | XM_001650445.2 |
| XM_017015227.1 | XM_021844606.1 |
| XM_017015240.2 | XM_021842333.1 |
| XM_017015241.2 | XM_021851446.1 |
| XM_017015253.1 | XM_001660052.2 |
| XM_017015312.2 | XM_021843377.1 |
| XM_017015316.1 | XM_021853594.1 |
| XM_017015355.2 | XM_021841683.1 |
| XM_017015360.2 | XM_021844590.1 |
| XM_017015380.1 | XM_021840826.1 |
| XM_017015419.1 | XM_021839053.1 |
| XM_017015453.1 | XM_021846195.1 |
| XM_017015454.1 | XM_021837807.1 |
| XM_017015512.1 | XM_021849024.1 |
| XM_017015540.1 | XM_021847227.1 |
| XM_017015572.1 | XM_021849684.1 |
| XM_017015574.2 | XM_021845733.1 |
| XM_017015584.2 | XR_002500734.1 |
| XM_017015604.1 | XM_021849131.1 |
| XM_017015617.1 | XR_002501333.1 |
| XM_017015623.1 | XM_021848996.1 |
| XM_017015630.1 | XR_002498841.1 |
| XM_017015674.1 | XM_001658212.2 |
| XM_017015687.2 | XM_021845058.1 |
| XM_017015703.2 | XM_021838411.1 |
| XM_017015712.1 | XM_001660552.3 |
| XM_017015740.2 | XM_001649559.2 |
| XM_017015831.2 | XM_021837228.1 |
| XM_017015849.1 | XM_021847321.1 |
| XM_017015855.1 | XM_021847103.1 |
| XM_017015878.1 | XM_021845958.1 |
| XM_017015928.1 | XR_002499104.1 |
| XM_017015982.1 | XM_021841204.1 |
| XM_017016042.1 | XM_021848964.1 |
| XM_017016073.1 | XM_001662419.2 |
| XM_017016100.1 | XM_021840360.1 |
| XM_017016130.1 | XM_021851616.1 |
| XM_017016164.1 | XM_001649410.2 |

| | |
|---|---|
| XM_017016170.1 | XM_021842099.1 |
| XM_017016180.1 | XM_001663131.2 |
| XM_017016182.1 | XM_001652741.2 |
| XM_017016209.2 | XR_002498897.1 |
| XM_017016210.2 | XM_021854476.1 |
| XM_017016225.2 | XM_021855333.1 |
| XM_017016328.2 | XM_001662970.2 |
| XM_017016345.2 | XM_021853751.1 |
| XM_017016381.2 | XM_001662901.2 |
| XM_017016402.1 | XM_021849208.1 |
| XM_017016438.2 | XM_021848986.1 |
| XM_017016453.1 | XM_021856163.1 |
| XM_017016480.1 | XM_001661522.2 |
| XM_017016496.2 | XM_011495107.2 |
| XM_017016498.1 | XM_001660323.2 |
| XM_017016511.1 | XM_021854676.1 |
| XM_017016546.2 | XR_002500021.1 |
| XM_017016622.1 | XM_001651045.2 |
| XM_017016632.2 | XM_021849571.1 |
| XM_017016637.1 | XM_021852760.1 |
| XM_017016687.1 | XM_021840358.1 |
| XM_017016715.2 | XM_021857451.1 |
| XM_017016789.2 | XM_021852196.1 |
| XM_017016840.1 | XM_021845735.1 |
| XM_017016853.2 | XM_021841193.1 |
| XM_017016868.1 | XM_001663790.2 |
| XM_017016912.2 | XM_021839664.1 |
| XM_017016941.1 | XM_021838982.1 |
| XM_017016949.2 | XR_002501765.1 |
| XM_017016962.1 | XM_001662516.2 |
| XM_017016973.1 | XR_002499810.1 |
| XM_017017067.1 | XM_021848152.1 |
| XM_017017082.2 | XM_001661829.2 |
| XM_017017087.1 | XM_021847478.1 |
| XM_017017095.1 | XM_021848966.1 |
| XM_017017112.2 | XM_021840529.1 |
| XM_017017137.1 | XM_021845186.1 |
| XM_017017144.2 | XM_021838949.1 |
| XM_017017146.2 | XM_001653614.2 |
| XM_017017212.1 | XM_021839096.1 |
| XM_017017241.2 | XM_021845971.1 |
| XM_017017280.2 | XM_021837970.1 |
| XM_017017361.1 | XM_021853711.1 |
| XM_017017367.1 | XM_021848959.1 |
| XM_017017373.2 | XM_021853580.1 |
| XM_017017399.1 | XM_021847923.1 |
| XM_017017402.1 | XM_001657778.2 |
| XM_017017412.1 | XM_001651506.2 |
| XM_017017420.1 | XM_021842776.1 |
| XM_017017431.1 | XM_021838984.1 |

| | |
|---|---|
| XM_017017462.2 | XM_021852258.1 |
| XM_017017512.1 | XM_021843491.1 |
| XM_017017520.1 | XM_001649224.2 |
| XM_017017574.1 | XM_021846227.1 |
| XM_017017644.1 | XM_021854810.1 |
| XM_017017672.1 | XM_021850750.1 |
| XM_017017723.1 | XM_021846152.1 |
| XM_017017738.2 | XM_021854435.1 |
| XM_017017781.1 | XR_002499174.1 |
| XM_017017840.1 | XM_021838722.1 |
| XM_017017843.2 | XM_021856356.1 |
| XM_017017894.1 | XM_021840359.1 |
| XM_017017912.1 | XM_021838455.1 |
| XM_017017923.1 | XM_021849979.1 |
| XM_017017926.1 | XR_002501466.1 |
| XM_017017932.1 | XM_021840649.1 |
| XM_017017949.2 | XM_001648338.2 |
| XM_017018004.1 | XM_021849742.1 |
| XM_017018045.1 | XM_001656155.2 |
| XM_017018101.2 | XM_021843541.1 |
| XM_017018128.1 | XM_021845433.1 |
| XM_017018133.1 | XM_021854922.1 |
| XM_017018139.1 | XM_021845874.1 |
| XM_017018142.1 | XM_021840534.1 |
| XM_017018145.1 | XM_021853709.1 |
| XM_017018146.1 | XM_021838633.1 |
| XM_017018180.1 | XM_021852769.1 |
| XM_017018199.1 | XM_021839428.1 |
| XM_017018206.1 | XM_021855218.1 |
| XM_017018212.1 | XM_021845348.1 |
| XM_017018215.1 | XM_021855224.1 |
| XM_017018238.1 | XM_001649472.2 |
| XM_017018303.1 | XM_021856365.1 |
| XM_017018345.1 | XM_021849349.1 |
| XM_017018367.1 | XM_021847101.1 |
| XM_017018376.2 | XM_001657686.2 |
| XM_017018415.2 | XM_021855824.1 |
| XM_017018458.1 | XM_021846886.1 |
| XM_017018474.2 | XR_002499197.1 |
| XM_017018537.1 | XM_021853430.1 |
| XM_017018566.1 | XM_021842752.1 |
| XM_017018570.1 | XR_002502148.1 |
| XM_017018666.2 | XM_021844408.1 |
| XM_017018708.1 | XM_021840220.1 |
| XM_017018798.2 | XM_021851615.1 |
| XM_017018836.1 | XM_021857129.1 |
| XM_017018905.1 | XM_021845868.1 |
| XM_017018937.1 | XM_021838031.1 |
| XM_017018956.1 | XM_021850352.1 |
| XM_017018963.2 | XM_021850148.1 |

| | |
|---|---|
| XM_017018973.1 | XM_021851490.1 |
| XM_017018981.2 | XM_021857383.1 |
| XM_017019010.1 | XM_021837963.1 |
| XM_017019020.1 | XM_021842796.1 |
| XM_017019037.2 | XM_021840366.1 |
| XM_017019057.1 | XR_002499074.1 |
| XM_017019071.1 | XM_021856361.1 |
| XM_017019130.2 | XM_021848007.1 |
| XM_017019152.2 | XM_001656747.2 |
| XM_017019172.1 | XM_021841685.1 |
| XM_017019217.1 | XM_021852750.1 |
| XM_017019229.2 | XM_021845519.1 |
| XM_017019237.1 | XM_001655348.2 |
| XM_017019240.2 | XM_001657075.2 |
| XM_017019248.1 | XM_021848958.1 |
| XM_017019252.2 | XM_021845818.1 |
| XM_017019270.1 | XM_021853133.1 |
| XM_017019276.1 | XM_021848369.1 |
| XM_017019288.2 | XM_001657650.2 |
| XM_017019297.1 | XM_021846578.1 |
| XM_017019322.1 | XM_021841268.1 |
| XM_017019335.1 | XM_011494742.2 |
| XM_017019352.1 | XM_001652603.2 |
| XM_017019442.2 | XM_021840979.1 |
| XM_017019463.2 | XM_021839750.1 |
| XM_017019466.2 | XM_001660992.2 |
| XM_017019469.1 | XM_021849642.1 |
| XM_017019473.2 | XM_021837941.1 |
| XM_017019535.1 | XM_021839386.1 |
| XM_017019572.1 | XM_021843714.1 |
| XM_017019591.1 | XM_001659291.2 |
| XM_017019606.1 | XM_021840437.1 |
| XM_017019626.2 | XM_021851516.1 |
| XM_017019644.1 | XM_001655286.2 |
| XM_017019652.2 | XM_021849602.1 |
| XM_017019739.2 | XM_021857080.1 |
| XM_017019767.1 | XM_021843639.1 |
| XM_017019775.2 | XR_002501280.1 |
| XM_017019782.1 | XM_001656480.2 |
| XM_017019783.2 | XM_021855311.1 |
| XM_017019800.1 | XM_021839135.1 |
| XM_017019815.1 | XM_001653194.2 |
| XM_017019836.1 | XM_021840990.1 |
| XM_017019962.2 | XM_001661224.3 |
| XM_017019969.2 | XM_001662166.2 |
| XM_017019991.2 | XM_021848881.1 |
| XM_017020016.2 | XM_021840970.1 |
| XM_017020021.1 | XM_021855893.1 |
| XM_017020023.1 | XM_001660526.2 |
| XM_017020052.2 | XM_021857548.1 |

| | |
|---|---|
| XM_017020069.2 | XM_021847348.1 |
| XM_017020092.2 | XM_021841761.1 |
| XM_017020141.1 | XM_021856536.1 |
| XM_017020145.1 | XM_001654760.2 |
| XM_017020151.1 | XM_021841426.1 |
| XM_017020185.2 | XM_001659729.2 |
| XM_017020207.1 | XM_021837550.1 |
| XM_017020218.1 | XM_021845900.1 |
| XM_017020219.1 | XM_001651413.2 |
| XM_017020241.2 | XM_001656292.2 |
| XM_017020243.1 | XM_021841549.1 |
| XM_017020255.2 | XM_001652877.3 |
| XM_017020262.1 | XM_021837549.1 |
| XM_017020266.1 | XM_021837759.1 |
| XM_017020298.1 | XM_001655070.2 |
| XM_017020317.2 | XM_021846984.1 |
| XM_017020331.1 | XM_001661910.2 |
| XM_017020349.1 | XM_021840181.1 |
| XM_017020362.2 | XM_021854220.1 |
| XM_017020363.2 | XM_021846853.1 |
| XM_017020376.1 | XM_001654716.2 |
| XM_017020429.1 | XM_021838985.1 |
| XM_017020436.2 | XM_001661385.2 |
| XM_017020489.1 | XR_002499277.1 |
| XM_017020493.1 | XM_021852651.1 |
| XM_017020508.1 | XM_001658657.2 |
| XM_017020527.1 | XR_002503185.1 |
| XM_017020544.1 | XM_021849207.1 |
| XM_017020549.1 | XM_021840151.1 |
| XM_017020593.2 | XR_002500076.1 |
| XM_017020604.1 | XM_021847381.1 |
| XM_017020609.1 | XM_021847223.1 |
| XM_017020647.1 | XM_021851465.1 |
| XM_017020680.2 | XR_002502610.1 |
| XM_017020684.1 | XM_021841679.1 |
| XM_017020704.2 | XM_001656722.2 |
| XM_017020733.2 | XM_021838270.1 |
| XM_017020799.2 | XM_021839755.1 |
| XM_017020815.1 | XM_021851724.1 |
| XM_017020829.2 | XM_021838806.1 |
| XM_017020833.1 | XM_021854434.1 |
| XM_017020846.1 | XM_021850791.1 |
| XM_017020853.2 | XM_021852661.1 |
| XM_017020859.2 | XM_001648958.2 |
| XM_017020875.1 | XM_021854468.1 |
| XM_017020926.1 | XM_021840981.1 |
| XM_017020976.1 | XM_021854322.1 |
| XM_017020988.1 | XM_001652778.2 |
| XM_017020993.1 | XM_001663665.2 |
| XM_017021035.2 | XM_021855729.1 |

| | |
|---|---|
| XM_017021070.1 | XM_021842197.1 |
| XM_017021071.1 | XM_021838811.1 |
| XM_017021079.1 | XM_001658502.2 |
| XM_017021104.2 | XM_021839427.1 |
| XM_017021113.2 | XM_021855585.1 |
| XM_017021121.2 | XM_021853089.1 |
| XM_017021147.1 | XM_021849731.1 |
| XM_017021156.2 | XM_021842193.1 |
| XM_017021177.1 | XM_021842192.1 |
| XM_017021203.1 | XM_021847116.1 |
| XM_017021212.1 | XM_021851924.1 |
| XM_017021216.2 | XM_001657484.2 |
| XM_017021232.1 | XM_021857653.1 |
| XM_017021249.2 | XM_021838996.1 |
| XM_017021253.2 | XM_001656197.2 |
| XM_017021271.1 | XR_002501393.1 |
| XM_017021293.1 | XM_021849835.1 |
| XM_017021307.2 | XM_021851915.1 |
| XM_017021313.1 | XM_021846213.1 |
| XM_017021353.1 | XM_021852535.1 |
| XM_017021373.1 | XM_001650066.2 |
| XM_017021390.2 | XM_001659607.2 |
| XM_017021421.2 | XM_021837551.1 |
| XM_017021431.1 | XM_021850987.1 |
| XM_017021433.1 | XR_002500935.1 |
| XM_017021437.2 | XM_001657208.2 |
| XM_017021452.1 | XM_021843905.1 |
| XM_017021455.1 | XM_001651158.2 |
| XM_017021476.1 | XM_021857449.1 |
| XM_017021517.1 | XM_001655055.2 |
| XM_017021576.1 | XM_021842149.1 |
| XM_017021662.1 | XM_021851040.1 |
| XM_017021668.2 | XM_001659550.2 |
| XM_017021678.2 | XM_021846145.1 |
| XM_017021690.2 | XR_002501655.1 |
| XM_017021701.1 | XM_021849020.1 |
| XM_017021702.1 | XM_021855889.1 |
| XM_017021727.2 | XM_021857724.1 |
| XM_017021734.1 | XM_001651047.2 |
| XM_017021735.1 | XM_001659714.2 |
| XM_017021755.2 | XM_021848993.1 |
| XM_017021774.1 | XM_001654826.2 |
| XM_017021799.2 | XM_021837411.1 |
| XM_017021820.2 | XM_001658878.2 |
| XM_017021867.2 | XM_001663011.2 |
| XM_017021897.2 | XM_001657179.2 |
| XM_017021927.1 | XM_021840650.1 |
| XM_017021933.1 | XM_021838407.1 |
| XM_017021937.2 | XM_001660912.2 |
| XM_017021940.2 | XM_021851781.1 |

| | |
|---|---|
| XM_017022007.1 | XM_021838533.1 |
| XM_017022031.2 | XM_021837752.1 |
| XM_017022073.1 | XM_021840695.1 |
| XM_017022099.1 | XM_021848899.1 |
| XM_017022110.1 | XR_002501327.1 |
| XM_017022137.1 | XM_021840980.1 |
| XM_017022160.2 | XM_021837846.1 |
| XM_017022198.2 | XM_021838581.1 |
| XM_017022201.1 | XM_001663983.2 |
| XM_017022226.1 | XM_021853898.1 |
| XM_017022233.1 | XM_021842229.1 |
| XM_017022236.2 | XM_001650712.2 |
| XM_017022259.1 | XM_021852369.1 |
| XM_017022276.1 | XM_021841681.1 |
| XM_017022286.1 | XM_001662644.3 |
| XM_017022298.1 | XM_021838398.1 |
| XM_017022302.1 | XM_021850313.1 |
| XM_017022310.2 | XM_001651507.2 |
| XM_017022349.1 | XM_001660082.2 |
| XM_017022370.1 | XM_021849739.1 |
| XM_017022374.2 | XM_021852481.1 |
| XM_017022384.2 | XM_021838404.1 |
| XM_017022387.2 | XM_021843385.1 |
| XM_017022450.1 | XM_021850341.1 |
| XM_017022463.2 | XM_021838807.1 |
| XM_017022540.1 | XM_021853163.1 |
| XM_017022570.1 | XM_021839306.1 |
| XM_017022575.2 | XM_021840527.1 |
| XM_017022657.1 | XR_002501758.1 |
| XM_017022659.1 | XM_021845694.1 |
| XM_017022673.2 | XM_021852852.1 |
| XM_017022685.1 | XR_002500264.1 |
| XM_017022712.2 | XM_021840239.1 |
| XM_017022761.1 | XM_021854433.1 |
| XM_017022778.1 | XM_001661157.2 |
| XM_017022815.2 | XM_021847224.1 |
| XM_017022822.2 | XM_021853074.1 |
| XM_017022824.2 | XR_002499731.1 |
| XM_017022876.1 | XM_001652468.2 |
| XM_017022912.1 | XR_002502727.1 |
| XM_017022921.2 | XM_021844770.1 |
| XM_017022925.1 | XM_001656833.2 |
| XM_017022928.1 | XM_021851838.1 |
| XM_017022960.2 | XM_001661928.2 |
| XM_017022968.1 | XR_002500324.1 |
| XM_017023019.1 | XM_001656505.2 |
| XM_017023077.1 | XM_021841762.1 |
| XM_017023098.1 | XR_002502079.1 |
| XM_017023099.1 | XM_001656716.3 |
| XM_017023190.2 | XR_002500862.1 |

| | |
|---|---|
| XM_017023193.2 | XM_021857399.1 |
| XM_017023266.2 | XM_021838737.1 |
| XM_017023277.1 | XM_021848106.1 |
| XM_017023281.1 | XM_021839849.1 |
| XM_017023297.2 | XM_021837894.1 |
| XM_017023303.2 | XM_021844578.1 |
| XM_017023350.1 | XM_021837823.1 |
| XM_017023383.2 | XM_021856986.1 |
| XM_017023403.1 | XM_001656309.2 |
| XM_017023424.2 | XM_021847879.1 |
| XM_017023477.1 | XM_021848892.1 |
| XM_017023527.1 | XM_021853398.1 |
| XM_017023532.1 | XM_021857361.1 |
| XM_017023550.1 | XM_021839756.1 |
| XM_017023554.1 | XM_021840997.1 |
| XM_017023625.1 | XM_001648434.2 |
| XM_017023634.2 | XM_021855381.1 |
| XM_017023707.2 | XM_021857270.1 |
| XM_017023715.1 | XM_021848466.1 |
| XM_017023723.2 | XM_001661478.2 |
| XM_017023751.1 | XM_021857679.1 |
| XM_017023757.1 | XM_021842539.1 |
| XM_017023770.1 | XM_021840140.1 |
| XM_017023786.2 | XM_001664232.2 |
| XM_017023808.1 | XM_001657293.2 |
| XM_017023853.2 | XR_002498607.1 |
| XM_017023859.2 | XM_021845993.1 |
| XM_017023966.2 | XM_021837622.1 |
| XM_017023974.1 | XM_021850458.1 |
| XM_017023991.1 | XM_021845429.1 |
| XM_017024006.1 | XM_021838047.1 |
| XM_017024027.1 | XM_021853189.1 |
| XM_017024038.2 | XM_001649726.2 |
| XM_017024040.2 | XM_021857681.1 |
| XM_017024057.1 | XM_021837523.1 |
| XM_017024075.2 | XM_021838635.1 |
| XM_017024079.1 | XM_021855548.1 |
| XM_017024102.1 | XM_021850226.1 |
| XM_017024122.1 | XM_021853124.1 |
| XM_017024149.2 | XM_001652347.2 |
| XM_017024200.1 | XM_021838803.1 |
| XM_017024209.1 | XM_001660791.2 |
| XM_017024228.1 | XM_021841078.1 |
| XM_017024273.1 | XM_021852051.1 |
| XM_017024277.1 | XM_021850447.1 |
| XM_017024289.2 | XM_001663742.2 |
| XM_017024297.1 | XR_002503284.1 |
| XM_017024325.1 | XM_021839114.1 |
| XM_017024338.1 | XM_021841847.1 |
| XM_017024371.1 | XM_001658024.2 |

| | |
|---|---|
| XM_017024409.1 | XM_001657276.2 |
| XM_017024434.1 | XM_021839115.1 |
| XM_017024468.2 | XR_002498857.1 |
| XM_017024484.1 | XR_002498868.1 |
| XM_017024543.2 | XR_002502657.1 |
| XM_017024574.1 | XM_021840988.1 |
| XM_017024618.1 | XR_002501529.1 |
| XM_017024641.2 | XM_021855586.1 |
| XM_017024642.1 | XM_021846004.1 |
| XM_017024730.2 | XM_021856368.1 |
| XM_017024731.2 | XM_021855379.1 |
| XM_017024736.1 | XM_021855138.1 |
| XM_017024740.1 | XM_021857525.1 |
| XM_017024764.2 | XM_021838222.1 |
| XM_017024777.1 | XM_001663386.2 |
| XM_017024808.1 | XM_001662851.2 |
| XM_017024825.1 | XM_001653316.2 |
| XM_017024841.1 | XM_021848686.1 |
| XM_017024844.2 | XM_021849743.1 |
| XM_017024845.2 | XM_021849073.1 |
| XM_017024846.2 | XM_001657465.2 |
| XM_017024921.2 | XM_001652306.2 |
| XM_017024955.1 | XM_001647878.2 |
| XM_017024989.1 | XM_001652231.2 |
| XM_017025036.2 | XM_011494870.2 |
| XM_017025061.1 | XM_001650481.2 |
| XM_017025095.1 | XM_021855923.1 |
| XM_017025096.1 | XM_021854906.1 |
| XM_017025113.1 | XM_021842175.1 |
| XM_017025132.2 | XM_001660253.2 |
| XM_017025146.1 | XM_021850665.1 |
| XM_017025159.2 | XM_021851995.1 |
| XM_017025166.2 | XM_021854895.1 |
| XM_017025170.1 | XM_021841538.1 |
| XM_017025231.1 | XM_021848236.1 |
| XM_017025233.1 | XM_021842012.1 |
| XM_017025305.1 | XM_001656366.2 |
| XM_017025350.1 | XM_021851564.1 |
| XM_017025374.2 | XM_001651438.2 |
| XM_017025377.1 | XM_021838134.1 |
| XM_017025397.2 | XM_001659743.2 |
| XM_017025409.2 | XM_001656502.2 |
| XM_017025441.1 | XM_001663051.2 |
| XM_017025451.1 | XM_021842572.1 |
| XM_017025453.1 | XM_021849108.1 |
| XM_017025454.2 | XM_021838781.1 |
| XM_017025514.2 | XM_021850476.1 |
| XM_017025530.1 | XM_021851843.1 |
| XM_017025561.1 | XM_021844018.1 |
| XM_017025574.1 | XM_001650801.2 |

| | |
|---|---|
| XM_017025599.2 | XM_001652367.2 |
| XM_017025663.2 | XM_021849304.1 |
| XM_017025669.2 | XM_021837615.1 |
| XM_017025684.1 | XM_021838790.1 |
| XM_017025716.1 | XM_001661129.2 |
| XM_017025728.2 | XR_002503147.1 |
| XM_017025787.1 | XM_021855918.1 |
| XM_017025803.2 | XM_021845979.1 |
| XM_017025808.2 | XM_021844840.1 |
| XM_017025894.1 | XM_021853164.1 |
| XM_017025927.2 | XR_002500500.1 |
| XM_017025967.1 | XR_002501319.1 |
| XM_017025993.1 | XM_021847790.1 |
| XM_017026007.1 | XM_021851839.1 |
| XM_017026035.2 | XM_001658610.2 |
| XM_017026086.1 | XM_021853598.1 |
| XM_017026099.1 | XM_001648855.2 |
| XM_017026150.1 | XM_021855182.1 |
| XM_017026170.1 | XM_021845435.1 |
| XM_017026172.2 | XM_021851644.1 |
| XM_017026191.1 | XR_002500798.1 |
| XM_017026200.1 | XM_021843980.1 |
| XM_017026202.2 | XM_021849837.1 |
| XM_017026203.2 | XR_002501965.1 |
| XM_017026210.1 | XM_001663809.2 |
| XM_017026213.1 | XM_021847221.1 |
| XM_017026216.1 | XM_021852853.1 |
| XM_017026243.2 | XM_021853672.1 |
| XM_017026246.1 | XM_021843536.1 |
| XM_017026262.2 | XM_021852540.1 |
| XM_017026269.1 | XM_001657109.2 |
| XM_017026324.1 | XM_021853129.1 |
| XM_017026354.1 | XM_021842074.1 |
| XM_017026411.2 | XM_021853791.1 |
| XM_017026445.1 | XM_021850744.1 |
| XM_017026456.1 | XM_001655545.2 |
| XM_017026470.2 | XM_001659214.2 |
| XM_017026523.1 | XM_021845547.1 |
| XM_017026530.2 | XM_001650645.2 |
| XM_017026541.1 | XM_021856031.1 |
| XM_017026551.1 | XM_021844796.1 |
| XM_017026558.1 | XR_002499315.1 |
| XM_017026573.1 | XM_001650118.2 |
| XM_017026601.1 | XM_021848363.1 |
| XM_017026643.1 | XM_021843384.1 |
| XM_017026711.1 | XM_021846215.1 |
| XM_017026726.1 | XM_021849430.1 |
| XM_017026746.1 | XM_021855373.1 |
| XM_017026752.1 | XM_001658980.2 |
| XM_017026771.1 | XR_002498578.1 |

| | |
|---|---|
| XM_017026798.2 | XM_021838758.1 |
| XM_017026855.1 | XM_021842803.1 |
| XM_017026857.2 | XM_021849636.1 |
| XM_017026928.1 | XM_021838282.1 |
| XM_017026942.1 | XM_021857355.1 |
| XM_017026983.1 | XM_001663411.2 |
| XM_017027020.1 | XR_002500082.1 |
| XM_017027040.1 | XM_021852818.1 |
| XM_017027058.1 | XM_001660394.2 |
| XM_017027074.1 | XM_001653536.2 |
| XM_017027077.1 | XM_021837259.1 |
| XM_017027094.1 | XM_001658778.2 |
| XM_017027096.1 | XM_021849652.1 |
| XM_017027113.2 | XM_021857286.1 |
| XM_017027139.1 | XM_001660279.2 |
| XM_017027153.1 | XM_021848855.1 |
| XM_017027204.2 | XR_002498725.1 |
| XM_017027220.2 | XM_021839906.1 |
| XM_017027231.1 | XM_021838905.1 |
| XM_017027258.1 | XM_001654414.2 |
| XM_017027297.2 | XM_021853036.1 |
| XM_017027305.1 | XM_021852488.1 |
| XM_017027312.1 | XM_021837423.1 |
| XM_017027316.1 | XM_021843545.1 |
| XM_017027327.1 | XM_021837413.1 |
| XM_017027334.1 | XM_021842797.1 |
| XM_017027358.1 | XM_021851639.1 |
| XM_017027362.1 | XM_001652110.2 |
| XM_017027374.2 | XR_002498995.1 |
| XM_017027411.2 | XM_011494914.2 |
| XM_017027428.1 | XR_002502337.1 |
| XM_017027430.1 | XM_021841131.1 |
| XM_017027451.1 | XM_001649703.2 |
| XM_017027453.1 | XM_021851201.1 |
| XM_017027461.1 | XM_021840287.1 |
| XM_017027464.1 | XM_001648178.2 |
| XM_017027481.1 | XM_001656842.2 |
| XM_017027522.2 | XM_021853701.1 |
| XM_017027529.1 | XM_021851914.1 |
| XM_017027532.1 | XM_021841500.1 |
| XM_017027554.1 | XM_021848730.1 |
| XM_017027565.1 | XM_001659344.2 |
| XM_017027592.2 | XM_001649222.2 |
| XM_017027600.2 | XM_021849437.1 |
| XM_017027602.1 | XM_021841744.1 |
| XM_017027628.1 | XM_001662619.2 |
| XM_017027646.1 | XM_021841351.1 |
| XM_017027647.2 | XM_021847936.1 |
| XM_017027667.1 | XM_021847193.1 |
| XM_017027678.1 | XM_021856905.1 |

| | |
|---|---|
| XM_017027692.2 | XM_021839140.1 |
| XM_017027736.2 | XM_021845526.1 |
| XM_017027744.2 | XM_021843543.1 |
| XM_017027754.2 | XR_002501068.1 |
| XM_017027784.2 | XM_001658307.2 |
| XM_017027803.2 | XM_021854062.1 |
| XM_017027827.2 | XM_021841881.1 |
| XM_017027894.2 | XM_021847069.1 |
| XM_017027915.2 | XM_021847071.1 |
| XM_017027951.2 | XM_001657604.2 |
| XM_017027969.2 | XM_021842421.1 |
| XM_017027980.2 | XM_021838372.1 |
| XM_017027982.1 | XM_001654041.2 |
| XM_017027998.1 | XM_021845442.1 |
| XM_017027999.1 | XM_021839566.1 |
| XM_017028064.1 | XM_001660072.2 |
| XM_017028096.1 | XM_021845576.1 |
| XM_017028101.1 | XM_021849404.1 |
| XM_017028109.1 | XM_021853702.1 |
| XM_017028155.1 | XM_021843661.1 |
| XM_017028160.1 | XM_021838777.1 |
| XM_017028258.1 | XM_021838269.1 |
| XM_017028271.1 | XM_021850486.1 |
| XM_017028344.1 | XM_001652585.2 |
| XM_017028362.2 | XM_021852714.1 |
| XM_017028415.2 | XM_021838090.1 |
| XM_017028440.2 | XM_001661396.2 |
| XM_017028451.2 | XR_002499058.1 |
| XM_017028459.2 | XM_021841353.1 |
| XM_017028474.1 | XM_021838989.1 |
| XM_017028485.2 | XM_021852489.1 |
| XM_017028510.1 | XM_021839244.1 |
| XM_017028546.1 | XM_021851513.1 |
| XM_017028556.1 | XM_021852046.1 |
| XM_017028560.1 | XM_021854839.1 |
| XM_017028580.1 | XM_021852899.1 |
| XM_017028612.1 | XM_001653774.2 |
| XM_017028637.1 | XM_001661887.2 |
| XM_017028643.2 | XM_021856380.1 |
| XM_017028667.2 | XM_021843540.1 |
| XM_017028698.1 | XR_002502468.1 |
| XM_017028747.2 | XM_021854413.1 |
| XM_017028769.2 | XM_021855171.1 |
| XM_017028774.1 | XM_001658057.2 |
| XM_017028788.1 | XM_021837241.1 |
| XM_017028809.2 | XM_021849516.1 |
| XM_017028814.1 | XM_021838006.1 |
| XM_017028825.1 | XM_001657295.2 |
| XM_017028836.1 | XM_001653378.2 |
| XM_017028839.1 | XM_021856150.1 |

| | |
|---|---|
| XM_017028896.2 | XM_021849757.1 |
| XM_017028920.1 | XM_021855353.1 |
| XM_017028955.2 | XM_021852062.1 |
| XM_017028957.2 | XM_021852493.1 |
| XM_017028971.2 | XM_001656760.2 |
| XM_017028975.2 | XM_021841952.1 |
| XM_017028976.1 | XM_021837613.1 |
| XM_017029015.1 | XM_021853986.1 |
| XM_017029035.2 | XM_021854412.1 |
| XM_017029053.1 | XM_021853623.1 |
| XM_017029099.2 | XM_021851330.1 |
| XM_017029119.1 | XM_001661501.2 |
| XM_017029130.1 | XR_002502325.1 |
| XM_017029221.1 | XM_021837408.1 |
| XM_017029228.1 | XM_001662327.2 |
| XM_017029364.1 | XM_001656284.2 |
| XM_017029391.1 | XM_021848507.1 |
| XM_017029416.1 | XM_021846331.1 |
| XM_017029418.1 | XR_002502667.1 |
| XM_017029450.2 | XM_021855911.1 |
| XM_017029464.2 | XM_001647574.2 |
| XM_017029469.1 | XM_021842552.1 |
| XM_017029479.1 | XM_021837440.1 |
| XM_017029491.2 | XM_021839514.1 |
| XM_017029513.1 | XM_001651723.2 |
| XM_017029523.1 | XM_021853122.1 |
| XM_017029531.1 | XM_021847734.1 |
| XM_017029571.1 | XM_001650885.2 |
| XM_017029575.1 | XM_021848460.1 |
| XM_017029578.1 | XM_001659416.2 |
| XM_017029585.2 | XM_021857391.1 |
| XM_017029588.2 | XR_002501395.1 |
| XM_017029590.2 | XM_001653009.2 |
| XM_017029602.1 | XM_001652157.2 |
| XM_017029619.2 | XM_021847867.1 |
| XM_017029631.1 | XM_021847251.1 |
| XM_017029647.2 | XM_001650794.2 |
| XM_017029662.1 | XM_021848724.1 |
| XM_017029675.1 | XM_021844771.1 |
| XM_017029679.1 | XM_001661840.3 |
| XM_017029681.1 | XM_021850666.1 |
| XM_017029689.2 | XM_021846124.1 |
| XM_017029708.1 | XM_021837370.1 |
| XM_017029726.2 | XM_021849433.1 |
| XM_017029740.1 | XR_002500035.1 |
| XM_017029762.1 | XM_021840762.1 |
| XM_017029770.1 | XM_021853080.1 |
| XM_017029783.2 | XM_021838369.1 |
| XM_017029817.1 | XM_021849926.1 |
| XM_017029821.1 | XM_021855363.1 |

| | |
|---|---|
| XM_017029872.1 | XR_002498688.1 |
| XM_017029873.1 | XM_011494794.2 |
| XM_017029890.2 | XR_002499010.1 |
| XM_017029893.2 | XR_002502479.1 |
| XM_017029911.1 | XM_021838991.1 |
| XM_017029916.1 | XM_021841141.1 |
| XM_017029920.1 | XR_002501830.1 |
| XM_017029968.1 | XM_001656672.2 |
| XM_017030103.1 | XM_001657162.2 |
| XM_017030116.1 | XM_001655281.2 |
| XM_017030122.1 | XM_021849273.1 |
| XM_017030301.2 | XM_021841263.1 |
| XM_024446078.1 | XR_002498626.1 |
| XM_024446080.1 | XM_021848071.1 |
| XM_024446127.1 | XM_021854070.1 |
| XM_024446137.1 | XR_002502916.1 |
| XM_024446138.1 | XM_021849252.1 |
| XM_024446148.1 | XM_021847959.1 |
| XM_024446149.1 | XM_021848221.1 |
| XM_024446155.1 | XR_002498848.1 |
| XM_024446163.1 | XM_001656286.2 |
| XM_024446181.1 | XM_021849283.1 |
| XM_024446216.1 | XM_021845747.1 |
| XM_024446243.1 | XM_021837606.1 |
| XM_024446265.1 | XM_021847067.1 |
| XM_024446333.1 | XM_021850271.1 |
| XM_024446354.1 | XM_021838954.1 |
| XM_024446367.1 | XM_021850187.1 |
| XM_024446378.1 | XM_001660173.2 |
| XM_024446385.1 | XM_021839031.1 |
| XM_024446424.1 | XM_001651536.2 |
| XM_024446428.1 | XM_021844799.1 |
| XM_024446430.1 | XR_002499374.1 |
| XM_024446431.1 | XM_021852971.1 |
| XM_024446468.1 | XM_021839145.1 |
| XM_024446524.1 | XM_021853275.1 |
| XM_024446529.1 | XM_021855385.1 |
| XM_024446533.1 | XM_001652374.2 |
| XM_024446618.1 | XM_021853626.1 |
| XM_024446636.1 | XM_021839241.1 |
| XM_024446655.1 | XM_021838046.1 |
| XM_024446664.1 | XM_021856985.1 |
| XM_024446671.1 | XM_021841395.1 |
| XM_024446692.1 | XM_021850977.1 |
| XM_024446709.1 | XM_001650312.2 |
| XM_024446726.1 | XM_021851845.1 |
| XM_024446748.1 | XM_021843583.1 |
| XM_024446783.1 | XM_001653977.2 |
| XM_024446786.1 | XM_021853928.1 |
| XM_024446817.1 | XR_002499833.1 |

| | |
|---|---|
| XM_024446820.1 | XM_021852464.1 |
| XM_024446838.1 | XM_021840309.1 |
| XM_024446876.1 | XM_021850831.1 |
| XM_024446881.1 | XM_021841769.1 |
| XM_024446888.1 | XM_021849137.1 |
| XM_024446896.1 | XM_021845744.1 |
| XM_024446963.1 | XM_021848409.1 |
| XM_024446969.1 | XM_021842736.1 |
| XM_024446980.1 | XM_001655096.2 |
| XM_024446986.1 | XM_021849670.1 |
| XM_024447053.1 | XM_021843713.1 |
| XM_024447054.1 | XR_002500671.1 |
| XM_024447055.1 | XM_001654743.2 |
| XM_024447057.1 | XM_021849140.1 |
| XM_024447086.1 | XM_001654657.2 |
| XM_024447113.1 | XM_001655050.2 |
| XM_024447120.1 | XM_021854783.1 |
| XM_024447136.1 | XM_021840975.1 |
| XM_024447140.1 | XM_021846695.1 |
| XM_024447141.1 | XM_021856832.1 |
| XM_024447228.1 | XM_021849203.1 |
| XM_024447266.1 | XM_021853376.1 |
| XM_024447270.1 | XM_021848516.1 |
| XM_024447272.1 | XM_021857471.1 |
| XM_024447295.1 | XM_021849353.1 |
| XM_024447308.1 | XM_021848721.1 |
| XM_024447311.1 | XM_001656546.2 |
| XM_024447315.1 | XM_021837783.1 |
| XM_024447354.1 | XR_002501320.1 |
| XM_024447367.1 | XM_001655604.2 |
| XM_024447377.1 | XR_002500127.1 |
| XM_024447380.1 | XM_001648896.2 |
| XM_024447384.1 | XM_021843880.1 |
| XM_024447434.1 | XM_021853784.1 |
| XM_024447506.1 | XM_021842697.1 |
| XM_024447538.1 | XR_002499477.1 |
| XM_024447549.1 | XM_021839156.1 |
| XM_024447573.1 | XM_021852898.1 |
| XM_024447630.1 | XM_021856353.1 |
| XM_024447632.1 | XM_021854396.1 |
| XM_024447681.1 | XM_001654227.2 |
| XM_024447699.1 | XM_021856839.1 |
| XM_024447735.1 | XM_001656267.2 |
| XM_024447736.1 | XM_001658837.2 |
| XM_024447763.1 | XM_021840105.1 |
| XM_024447765.1 | XM_021852042.1 |
| XM_024447767.1 | XM_021837792.1 |
| XM_024447784.1 | XM_021842674.1 |
| XM_024447830.1 | XM_021850479.1 |
| XM_024447865.1 | XM_001662010.2 |

| | |
|---|---|
| XM_024447867.1 | XM_001660419.2 |
| XM_024447868.1 | XM_021852733.1 |
| XM_024447902.1 | XM_021843902.1 |
| XM_024447967.1 | XR_002501547.1 |
| XM_024447981.1 | XM_001653543.2 |
| XM_024447986.1 | XM_021842526.1 |
| XM_024448026.1 | XM_021849195.1 |
| XM_024448038.1 | XM_021844607.1 |
| XM_024448081.1 | XM_021837646.1 |
| XM_024448083.1 | XM_021853320.1 |
| XM_024448086.1 | XM_021845419.1 |
| XM_024448100.1 | XM_021855356.1 |
| XM_024448120.1 | XR_002502800.1 |
| XM_024448143.1 | XM_021853342.1 |
| XM_024448156.1 | XM_021853351.1 |
| XM_024448170.1 | XM_021847034.1 |
| XM_024448223.1 | XM_021856373.1 |
| XM_024448241.1 | XM_021837899.1 |
| XM_024448243.1 | XR_002501879.1 |
| XM_024448257.1 | XR_002499227.1 |
| XM_024448284.1 | XM_021842128.1 |
| XM_024448286.1 | XR_002501099.1 |
| XM_024448287.1 | XR_002500701.1 |
| XM_024448303.1 | XM_021851447.1 |
| XM_024448319.1 | XM_021842038.1 |
| XM_024448321.1 | XM_021846243.1 |
| XM_024448322.1 | XM_001653256.2 |
| XM_024448341.1 | XM_021854373.1 |
| XM_024448342.1 | XM_001660112.2 |
| XM_024448358.1 | XR_002501276.1 |
| XM_024448381.1 | XM_001658127.2 |
| XM_024448385.1 | XM_021842980.1 |
| XM_024448408.1 | XM_021840841.1 |
| XM_024448426.1 | XM_001650482.2 |
| XM_024448492.1 | XM_021855482.1 |
| XM_024448494.1 | XM_001650817.2 |
| XM_024448501.1 | XM_021856907.1 |
| XM_024448510.1 | XM_021848725.1 |
| XM_024448536.1 | XM_021853321.1 |
| XM_024448572.1 | XM_001664087.2 |
| XM_024448689.1 | XM_021837811.1 |
| XM_024448694.1 | XM_021850983.1 |
| XM_024448739.1 | XM_001663784.2 |
| XM_024448740.1 | XM_001658162.2 |
| XM_024448741.1 | XM_001660267.2 |
| XM_024448767.1 | XM_021856972.1 |
| XM_024448772.1 | XM_001659220.2 |
| XM_024448776.1 | XM_021840627.1 |
| XM_024448806.1 | XR_002499695.1 |
| XM_024448832.1 | XR_002499488.1 |

| | |
|---|---|
| XM_024448861.1 | XR_002498850.1 |
| XM_024448869.1 | XM_021852897.1 |
| XM_024448885.1 | XM_021857394.1 |
| XM_024448954.1 | XM_021849580.1 |
| XM_024449008.1 | XM_021845590.1 |
| XM_024449013.1 | XM_001647845.2 |
| XM_024449022.1 | XM_021841276.1 |
| XM_024449024.1 | XM_021855376.1 |
| XM_024449047.1 | XM_001651917.2 |
| XM_024449052.1 | XM_021856854.1 |
| XM_024449059.1 | XM_021844856.1 |
| XM_024449073.1 | XM_021851422.1 |
| XM_024449085.1 | XM_021852418.1 |
| XM_024449096.1 | XR_002502819.1 |
| XM_024449112.1 | XM_021841885.1 |
| XM_024449130.1 | XM_021837588.1 |
| XM_024449134.1 | XR_002503018.1 |
| XM_024449152.1 | XM_021837393.1 |
| XM_024449190.1 | XM_021851286.1 |
| XM_024449213.1 | XR_002499598.1 |
| XM_024449229.1 | XM_021843733.1 |
| XM_024449252.1 | XM_021855939.1 |
| XM_024449270.1 | XM_021841502.1 |
| XM_024449275.1 | XM_021839951.1 |
| XM_024449278.1 | XM_001648394.2 |
| XM_024449290.1 | XM_021849010.1 |
| XM_024449300.1 | XM_021847217.1 |
| XM_024449312.1 | XM_021853596.1 |
| XM_024449326.1 | XM_021850072.1 |
| XM_024449373.1 | XM_021842566.1 |
| XM_024449390.1 | XM_021851210.1 |
| XM_024449420.1 | XM_021847092.1 |
| XM_024449425.1 | XM_001657186.2 |
| XM_024449470.1 | XM_021856845.1 |
| XM_024449477.1 | XM_001648705.2 |
| XM_024449495.1 | XR_002499105.1 |
| XM_024449520.1 | XM_021840043.1 |
| XM_024449532.1 | XM_001647895.2 |
| XM_024449600.1 | XM_001653340.2 |
| XM_024449618.1 | XM_021842126.1 |
| XM_024449636.1 | XR_002502550.1 |
| XM_024449646.1 | XM_021849669.1 |
| XM_024449649.1 | XM_021840635.1 |
| XM_024449659.1 | XM_021837416.1 |
| XM_024449699.1 | XM_021848753.1 |
| XM_024449704.1 | XM_021854254.1 |
| XM_024449717.1 | XM_001653734.2 |
| XM_024449721.1 | XM_021847347.1 |
| XM_024449739.1 | XM_001650987.2 |
| XM_024449744.1 | XM_021838290.1 |

| | |
|---|---|
| XM_024449748.1 | XM_021837787.1 |
| XM_024449781.1 | XM_021845984.1 |
| XM_024449813.1 | XM_021846388.1 |
| XM_024449823.1 | XM_021837270.1 |
| XM_024449889.1 | XM_021842747.1 |
| XM_024449896.1 | XM_001653317.2 |
| XM_024449912.1 | XM_021850902.1 |
| XM_024449930.1 | XM_021838307.1 |
| XM_024449946.1 | XM_021856765.1 |
| XM_024449957.1 | XM_001661906.2 |
| XM_024449959.1 | XM_001660295.2 |
| XM_024449977.1 | XM_001662877.2 |
| XM_024449992.1 | XM_021838133.1 |
| XM_024449995.1 | XR_002500452.1 |
| XM_024450008.1 | XM_021843284.1 |
| XM_024450013.1 | XM_021851009.1 |
| XM_024450055.1 | XM_021837455.1 |
| XM_024450070.1 | XM_021853920.1 |
| XM_024450075.1 | XM_021848893.1 |
| XM_024450112.1 | XM_021843153.1 |
| XM_024450134.1 | XM_021857360.1 |
| XM_024450147.1 | XM_021855955.1 |
| XM_024450151.1 | XR_002499165.1 |
| XM_024450153.1 | XM_021840736.1 |
| XM_024450213.1 | XM_021856230.1 |
| XM_024450219.1 | XM_021844280.1 |
| XM_024450233.1 | XM_021854479.1 |
| XM_024450271.1 | XM_001663233.2 |
| XM_024450280.1 | XM_021855711.1 |
| XM_024450289.1 | XM_001657096.2 |
| XM_024450306.1 | XM_021838125.1 |
| XM_024450308.1 | XR_002499677.1 |
| XM_024450328.1 | XM_021840768.1 |
| XM_024450356.1 | XM_021849752.1 |
| XM_024450374.1 | XM_021842110.1 |
| XM_024450409.1 | XM_001661825.2 |
| XM_024450414.1 | XR_002499164.1 |
| XM_024450440.1 | XM_001656246.2 |
| XM_024450441.1 | XM_021838312.1 |
| XM_024450455.1 | XM_021852482.1 |
| XM_024450463.1 | XM_021854465.1 |
| XM_024450468.1 | XM_021837768.1 |
| XM_024450471.1 | XM_021849944.1 |
| XM_024450476.1 | XM_021854242.1 |
| XM_024450477.1 | XM_021840763.1 |
| XM_024450512.1 | XM_021851844.1 |
| XM_024450553.1 | XR_002500390.1 |
| XM_024450568.1 | XM_021851694.1 |
| XM_024450586.1 | XR_002499166.1 |
| XM_024450595.1 | XM_021837737.1 |

| | |
|---|---|
| XM_024450624.1 | XM_021844561.1 |
| XM_024450671.1 | XM_021856139.1 |
| XM_024450698.1 | XM_021846608.1 |
| XM_024450710.1 | XM_021853167.1 |
| XM_024450719.1 | XM_001653123.2 |
| XM_024450722.1 | XM_021840617.1 |
| XM_024450729.1 | XM_021840843.1 |
| XM_024450757.1 | XM_021838381.1 |
| XM_024450798.1 | XM_021849271.1 |
| XM_024450817.1 | XM_021857569.1 |
| XM_024450818.1 | XM_001648579.2 |
| XM_024450823.1 | XM_021841775.1 |
| XM_024450875.1 | XM_001661418.2 |
| XM_024450885.1 | XM_001660740.2 |
| XM_024450890.1 | XM_001653195.2 |
| XM_024450930.1 | XM_001660109.2 |
| XM_024450995.1 | XM_021840299.1 |
| XM_024451026.1 | XM_021847138.1 |
| XM_024451045.1 | XM_001662382.2 |
| XM_024451050.1 | XM_021837763.1 |
| XM_024451061.1 | XM_001658906.2 |
| XM_024451070.1 | XM_001659516.2 |
| XM_024451076.1 | XM_001652395.2 |
| XM_024451114.1 | XM_001651025.2 |
| XM_024451116.1 | XM_001662411.2 |
| XM_024451160.1 | XM_021837556.1 |
| XM_024451249.1 | XM_021846253.1 |
| XM_024451268.1 | XM_021847617.1 |
| XM_024451275.1 | XM_021840072.1 |
| XM_024451281.1 | XM_021853771.1 |
| XM_024451308.1 | XM_021837587.1 |
| XM_024451331.1 | XM_021855973.1 |
| XM_024451338.1 | XM_021840702.1 |
| XM_024451380.1 | XM_021841208.1 |
| XM_024451397.1 | XM_021853319.1 |
| XM_024451399.1 | XM_021845732.1 |
| XM_024451405.1 | XM_021856249.1 |
| XM_024451450.1 | XM_021838559.1 |
| XM_024451453.1 | XM_021854314.1 |
| XM_024451462.1 | XR_002499291.1 |
| XM_024451473.1 | XM_021849841.1 |
| XM_024451487.1 | XM_021843180.1 |
| XM_024451499.1 | XR_002498895.1 |
| XM_024451527.1 | XM_021839568.1 |
| XM_024451531.1 | XM_001661775.2 |
| XM_024451533.1 | XM_021854892.1 |
| XM_024451534.1 | XM_001656763.2 |
| XM_024451558.1 | XM_001657877.3 |
| XM_024451575.1 | XM_021854066.1 |
| XM_024451600.1 | XM_001651551.2 |

| | |
|---|---|
| XM_024451622.1 | XM_001652052.2 |
| XM_024451625.1 | XM_001649718.2 |
| XM_024451629.1 | XM_021838210.1 |
| XM_024451647.1 | XR_002499722.1 |
| XM_024451726.1 | XM_001661707.2 |
| XM_024451732.1 | XM_021849196.1 |
| XM_024451734.1 | XM_001647968.2 |
| XM_024451764.1 | XM_021847307.1 |
| XM_024451770.1 | XM_021842981.1 |
| XM_024451774.1 | XM_021840596.1 |
| XM_024451779.1 | XM_021846114.1 |
| XM_024451806.1 | XM_021852462.1 |
| XM_024451824.1 | XM_001653747.3 |
| XM_024451827.1 | XM_021849248.1 |
| XM_024451909.1 | XM_001654308.2 |
| XM_024451916.1 | XR_002501914.1 |
| XM_024451922.1 | XM_021846798.1 |
| XM_024451956.1 | XM_001658084.2 |
| XM_024451961.1 | XM_001658246.2 |
| XM_024451974.1 | XM_021842028.1 |
| XM_024451994.1 | XM_021850137.1 |
| XM_024452005.1 | XM_021837385.1 |
| XM_024452032.1 | XR_002501925.1 |
| XM_024452037.1 | XM_021849537.1 |
| XM_024452062.1 | XM_021837643.1 |
| XM_024452067.1 | XM_001652466.2 |
| XM_024452081.1 | XM_021856756.1 |
| XM_024452091.1 | XM_021847063.1 |
| XM_024452131.1 | XM_021850993.1 |
| XM_024452132.1 | XM_021848024.1 |
| XM_024452133.1 | XM_021837789.1 |
| XM_024452180.1 | XM_021837417.1 |
| XM_024452231.1 | XM_021842176.1 |
| XM_024452232.1 | XM_021856570.1 |
| XM_024452268.1 | XM_021855940.1 |
| XM_024452292.1 | XM_021839139.1 |
| XM_024452296.1 | XM_021854131.1 |
| XM_024452311.1 | XM_021848727.1 |
| XM_024452349.1 | XM_001654166.2 |
| XM_024452370.1 | XM_001648857.2 |
| XM_024452372.1 | XM_021842172.1 |
| XM_024452376.1 | XM_001659500.2 |
| XM_024452391.1 | XM_021842143.1 |
| XM_024452439.1 | XM_021848715.1 |
| XM_024452448.1 | XM_021837458.1 |
| XM_024452465.1 | XM_021853638.1 |
| XM_024452470.1 | XM_021855956.1 |
| XM_024452483.1 | XM_021841211.1 |
| XM_024452494.1 | XR_002502616.1 |
| XM_024452531.1 | XR_002500458.1 |

| | |
|---|---|
| XM_024452578.1 | XR_002503109.1 |
| XM_024452662.1 | XM_001648856.2 |
| XM_024452683.1 | XM_021849194.1 |
| XM_024452703.1 | XR_002499223.1 |
| XM_024452713.1 | XM_021848714.1 |
| XM_024452715.1 | XM_021842322.1 |
| XM_024452723.1 | XM_021848944.1 |
| XM_024452733.1 | XM_021840648.1 |
| XM_024452780.1 | XM_001648844.2 |
| XM_024452796.1 | XM_021849923.1 |
| XM_024452806.1 | XM_021854449.1 |
| XM_024452807.1 | XM_021853849.1 |
| XM_024452824.1 | XM_021844642.1 |
| XM_024452832.1 | XM_021837645.1 |
| XM_024452866.1 | XM_021849707.1 |
| XM_024452898.1 | XR_002501910.1 |
| XM_024452926.1 | XM_001656108.2 |
| XM_024452942.1 | XM_021852421.1 |
| XM_024452965.1 | XR_002502002.1 |
| XM_024453032.1 | XM_021840677.1 |
| XM_024453050.1 | XM_021838294.1 |
| XM_024453052.1 | XM_021853323.1 |
| XM_024453056.1 | XM_001660756.2 |
| XM_024453080.1 | XM_021839118.1 |
| XM_024453089.1 | XR_002499681.1 |
| XM_024453106.1 | XM_021847065.1 |
| XM_024453109.1 | XR_002499308.1 |
| XM_024453160.1 | XM_021848177.1 |
| XM_024453184.1 | XM_001647616.2 |
| XM_024453193.1 | XM_021857708.1 |
| XM_024453196.1 | XR_002499561.1 |
| XM_024453213.1 | XM_021854036.1 |
| XM_024453257.1 | XR_002501350.1 |
| XM_024453265.1 | XM_021854371.1 |
| XM_024453303.1 | XM_001650479.2 |
| XM_024453308.1 | XR_002500451.1 |
| XM_024453316.1 | XM_001649602.3 |
| XM_024453319.1 | XM_021851020.1 |
| XM_024453342.1 | XM_021851344.1 |
| XM_024453365.1 | XM_021853442.1 |
| XM_024453380.1 | XM_021850445.1 |
| XM_024453396.1 | XM_021855961.1 |
| XM_024453429.1 | XM_001654759.2 |
| XM_024453437.1 | XM_001661200.3 |
| XM_024453438.1 | XM_021837494.1 |
| XM_024453445.1 | XR_002502167.1 |
| XM_024453451.1 | XM_021839165.1 |
| XM_024453460.1 | XM_021851850.1 |
| XM_024453471.1 | XM_021841020.1 |
| XM_024453490.1 | XM_021848712.1 |

| | |
|---|---|
| XM_024453491.1 | XR_002501575.1 |
| XM_024453532.1 | XM_021849965.1 |
| XM_024453556.1 | XM_021856698.1 |
| XM_024453558.1 | XM_021843133.1 |
| XM_024453563.1 | XR_002502917.1 |
| XM_024453571.1 | XM_021841023.1 |
| XM_024453628.1 | XM_021841271.1 |
| XM_024453654.1 | XM_021846081.1 |
| XM_024453656.1 | XM_021848592.1 |
| XM_024453659.1 | XM_021847887.1 |
| XM_024453667.1 | XM_021843532.1 |
| XM_024453672.1 | XM_001648324.2 |
| XM_024453676.1 | XM_021838279.1 |
| XM_024453682.1 | XM_021842158.1 |
| XM_024453694.1 | XM_021852793.1 |
| XM_024453708.1 | XM_001662393.2 |
| XM_024453737.1 | XM_021845126.1 |
| XM_024453747.1 | XM_021847888.1 |
| XM_024453767.1 | XM_001654157.2 |
| XM_024453775.1 | XR_002501346.1 |
| XM_024453783.1 | XM_021851524.1 |
| XM_024453793.1 | XM_021842215.1 |
| XM_024453796.1 | XM_001658334.2 |
| XM_024453838.1 | XM_001661778.2 |
| XM_024453839.1 | XR_002501457.1 |
| XM_024453908.1 | XM_001649171.2 |
| XM_024453911.1 | XR_002502836.1 |
| XM_024453945.1 | XM_021856939.1 |
| XM_024454028.1 | XM_001655122.2 |
| XM_024454036.1 | XM_021846803.1 |
| XM_024454069.1 | XM_021852730.1 |
| XM_024454118.1 | XM_021849205.1 |
| XM_024454142.1 | XM_021855528.1 |
| XM_024454166.1 | XM_021853820.1 |
| XM_024454167.1 | XM_021855174.1 |
| XM_024454170.1 | XM_021843065.1 |
| XM_024454186.1 | XM_001647924.3 |
| XM_024454206.1 | XM_001662645.2 |
| XM_024454207.1 | XM_021855027.1 |
| XM_024454208.1 | XM_021851281.1 |
| XM_024454218.1 | XM_021855966.1 |
| XM_024454227.1 | XM_001653950.2 |
| XM_024454246.1 | XM_021838482.1 |
| XM_024454251.1 | XM_001651793.2 |
| XM_024454267.1 | XM_001659328.3 |
| XM_024454269.1 | XM_021842483.1 |
| XM_024454277.1 | XM_021845426.1 |
| XM_024454319.1 | XM_021855607.1 |
| XM_024454328.1 | XM_021855203.1 |
| XM_024454368.1 | XM_021857136.1 |

| | |
|---|---|
| XM_024454390.1 | XM_021838480.1 |
| XM_037916392.1 | XM_021840699.1 |
| XR_001736906.2 | XM_001655205.2 |
| XR_001736922.2 | XM_021842008.1 |
| XR_001736973.2 | XM_001653836.2 |
| XR_001736998.1 | XM_021857023.1 |
| XR_001737015.1 | XM_021841134.1 |
| XR_001737023.2 | XM_021844867.1 |
| XR_001737032.2 | XM_021854580.1 |
| XR_001737079.2 | XM_021851518.1 |
| XR_001737080.1 | XM_021846689.1 |
| XR_001737092.1 | XM_021844000.1 |
| XR_001737094.2 | XM_021843130.1 |
| XR_001737098.1 | XM_021837479.1 |
| XR_001737115.1 | XM_001658740.2 |
| XR_001737133.1 | XM_021850446.1 |
| XR_001737134.2 | XM_001652485.2 |
| XR_001737137.2 | XM_021848075.1 |
| XR_001737144.1 | XM_001654205.2 |
| XR_001737165.2 | XM_001651767.2 |
| XR_001737223.2 | XR_002500885.1 |
| XR_001737237.1 | XR_002499031.1 |
| XR_001737277.1 | XM_021842698.1 |
| XR_001737310.2 | XM_021838617.1 |
| XR_001737317.2 | XM_021854361.1 |
| XR_001737323.1 | XM_021843734.1 |
| XR_001737330.1 | XM_001655383.3 |
| XR_001737357.2 | XM_021845232.1 |
| XR_001737361.2 | XM_001653284.2 |
| XR_001737461.1 | XM_001663957.2 |
| XR_001737462.1 | XM_001659455.2 |
| XR_001737482.2 | XM_021853091.1 |
| XR_001737485.2 | XM_021850471.1 |
| XR_001737511.1 | XM_021850390.1 |
| XR_001737517.2 | XM_021855701.1 |
| XR_001737544.1 | XM_021837659.1 |
| XR_001737546.1 | XM_021839874.1 |
| XR_001737579.2 | XM_021841210.1 |
| XR_001737582.2 | XM_021841360.1 |
| XR_001737611.1 | XM_001656178.2 |
| XR_001737624.1 | XR_002502786.1 |
| XR_001737631.1 | XM_021837412.1 |
| XR_001737632.2 | XM_001654758.2 |
| XR_001737668.1 | XM_001655067.2 |
| XR_001737683.1 | XM_021841021.1 |
| XR_001737705.1 | XM_021847126.1 |
| XR_001737710.1 | XM_021848462.1 |
| XR_001737725.2 | XM_021854614.1 |
| XR_001737732.2 | XM_021855858.1 |
| XR_001737771.1 | XM_021855605.1 |

| | |
|---|---|
| XR_001737773.1 | XR_002500950.1 |
| XR_001737798.1 | XM_021841883.1 |
| XR_001737813.1 | XR_002501693.1 |
| XR_001737822.1 | XM_001652777.2 |
| XR_001737842.1 | XM_001662307.2 |
| XR_001737876.1 | XM_021837247.1 |
| XR_001737880.1 | XM_021845248.1 |
| XR_001737887.1 | XM_001661517.2 |
| XR_001737901.1 | XM_001653028.2 |
| XR_001737940.1 | XM_021841539.1 |
| XR_001737962.1 | XM_021840697.1 |
| XR_001737965.1 | XM_021849204.1 |
| XR_001738037.2 | XM_001656297.2 |
| XR_001738063.1 | XM_021856496.1 |
| XR_001738081.1 | XM_021838830.1 |
| XR_001738098.1 | XM_021856177.1 |
| XR_001738100.1 | XR_002499210.1 |
| XR_001738128.1 | XM_021846408.1 |
| XR_001738151.2 | XR_002502791.1 |
| XR_001738163.1 | XM_021855177.1 |
| XR_001738193.1 | XR_002500912.1 |
| XR_001738199.1 | XM_021856284.1 |
| XR_001738207.1 | XM_021840000.1 |
| XR_001738270.1 | XM_001653590.2 |
| XR_001738277.2 | XM_001650564.2 |
| XR_001738296.1 | XM_021837591.1 |
| XR_001738356.1 | XM_021844789.1 |
| XR_001738364.1 | XM_021844927.1 |
| XR_001738366.1 | XM_021847177.1 |
| XR_001738395.1 | XR_002500810.1 |
| XR_001738411.1 | XM_001658487.2 |
| XR_001738415.1 | XM_021846113.1 |
| XR_001738468.1 | XM_021849202.1 |
| XR_001738472.1 | XM_021848630.1 |
| XR_001738477.1 | XM_001662406.2 |
| XR_001738488.1 | XM_021840041.1 |
| XR_001738517.1 | XR_002500811.1 |
| XR_001738553.1 | XR_002501418.1 |
| XR_001738626.2 | XM_021840274.1 |
| XR_001738633.1 | XM_021857578.1 |
| XR_001738636.2 | XM_021842440.1 |
| XR_001738648.1 | XM_001659361.2 |
| XR_001738679.1 | XM_021848104.1 |
| XR_001738688.2 | XM_021847840.1 |
| XR_001738697.1 | XM_021846115.1 |
| XR_001738723.1 | XR_002502213.1 |
| XR_001738738.1 | XR_002502755.1 |
| XR_001738748.1 | XM_001648793.2 |
| XR_001738764.1 | XM_021840001.1 |
| XR_001738822.1 | XM_021841887.1 |

| | |
|---|---|
| XR_001738850.1 | XM_001660691.2 |
| XR_001738864.2 | XR_002500996.1 |
| XR_001738871.1 | XR_002498727.1 |
| XR_001738899.2 | XM_001653587.2 |
| XR_001738927.1 | XR_002502843.1 |
| XR_001738961.1 | XM_021854581.1 |
| XR_001738970.1 | XM_001651415.2 |
| XR_001738979.1 | XM_001650213.2 |
| XR_001738989.2 | XM_001651263.2 |
| XR_001739020.1 | XR_002502109.1 |
| XR_001739021.1 | XM_021852746.1 |
| XR_001739055.2 | XM_021847024.1 |
| XR_001739066.1 | XM_021848103.1 |
| XR_001739068.1 | XM_021843511.1 |
| XR_001739090.1 | XM_001650802.2 |
| XR_001739093.1 | XM_021838088.1 |
| XR_001739109.1 | XM_021849708.1 |
| XR_001739117.1 | XM_001654416.2 |
| XR_001739134.1 | XM_001661841.3 |
| XR_001739169.1 | XM_001652830.2 |
| XR_001739177.1 | XM_001650117.2 |
| XR_001739197.1 | XR_002500710.1 |
| XR_001739201.1 | XM_021842701.1 |
| XR_001739250.1 | XM_021842011.1 |
| XR_001739266.2 | XM_021852534.1 |
| XR_001739285.1 | XM_021840935.1 |
| XR_001739298.1 | XM_021852803.1 |
| XR_001739304.2 | XM_001649480.2 |
| XR_001739354.1 | XM_001658178.2 |
| XR_001739359.1 | XM_001651081.2 |
| XR_001739377.1 | XM_021852110.1 |
| XR_001739414.2 | XM_021837675.1 |
| XR_001739418.2 | XM_001660153.2 |
| XR_001739422.1 | XM_021848760.1 |
| XR_001739445.1 | XM_021853322.1 |
| XR_001739496.1 | XM_011494668.2 |
| XR_001739519.1 | XM_021850575.1 |
| XR_001739530.1 | XM_021849090.1 |
| XR_001739556.1 | XM_001663923.2 |
| XR_001739566.1 | XM_021855611.1 |
| XR_001739576.1 | XM_021850709.1 |
| XR_001739636.1 | XM_001650557.2 |
| XR_001739660.1 | XM_021844965.1 |
| XR_001739664.1 | XM_001656257.2 |
| XR_001739695.2 | XM_001662001.2 |
| XR_001739716.1 | XM_001655482.2 |
| XR_001739755.1 | XR_002501934.1 |
| XR_001739778.1 | XM_021855990.1 |
| XR_001739815.1 | XM_021857313.1 |
| XR_001739823.1 | XM_021852111.1 |

| | |
|---|---|
| XR_001739846.1 | XM_021848968.1 |
| XR_001739852.2 | XM_001655054.2 |
| XR_001739861.1 | XM_021846732.1 |
| XR_001739873.1 | XM_021851024.1 |
| XR_001739889.1 | XM_021844696.1 |
| XR_001739906.1 | XM_021840074.1 |
| XR_001739923.1 | XM_021856270.1 |
| XR_001739979.1 | XM_021845424.1 |
| XR_001739996.2 | XM_001658557.2 |
| XR_001740001.1 | XM_021837465.1 |
| XR_001740047.1 | XM_021855360.1 |
| XR_001740091.1 | XM_001652331.2 |
| XR_001740131.2 | XM_021852741.1 |
| XR_001740138.1 | XM_021846101.1 |
| XR_001740145.2 | XR_002503113.1 |
| XR_001740149.2 | XR_002499890.1 |
| XR_001740164.1 | XM_021840250.1 |
| XR_001740175.1 | XM_021849888.1 |
| XR_001740199.2 | XM_021853324.1 |
| XR_001740200.2 | XM_001657415.2 |
| XR_001740209.2 | XM_021857327.1 |
| XR_001740219.2 | XM_001655620.2 |
| XR_001740232.1 | XM_001649022.2 |
| XR_001740250.1 | XM_001656353.2 |
| XR_001740253.2 | XM_001658614.2 |
| XR_001740257.1 | XM_021841541.1 |
| XR_001740301.2 | XM_021848465.1 |
| XR_001740353.2 | XM_021850425.1 |
| XR_001740396.1 | XM_001664252.2 |
| XR_001740412.1 | XM_001656773.2 |
| XR_001740415.1 | XR_002501003.1 |
| XR_001740436.2 | XM_021852494.1 |
| XR_001740438.1 | XM_021857026.1 |
| XR_001740449.1 | XM_011495358.2 |
| XR_001740465.1 | XM_011495055.2 |
| XR_001740467.1 | XM_021838668.1 |
| XR_001740471.1 | XM_021840381.1 |
| XR_001740490.2 | XM_021846461.1 |
| XR_001740500.1 | XM_021837597.1 |
| XR_001740524.1 | XM_021852207.1 |
| XR_001740547.1 | XM_021837510.1 |
| XR_001740554.1 | XR_002499424.1 |
| XR_001740575.1 | XM_021845528.1 |
| XR_001740599.1 | XM_021846397.1 |
| XR_001740600.1 | XM_001657105.2 |
| XR_001740605.2 | XR_002502480.1 |
| XR_001740609.1 | XM_001658500.2 |
| XR_001740621.1 | XM_021853869.1 |
| XR_001740622.1 | XM_021850632.1 |
| XR_001740634.1 | XM_021847129.1 |

| | |
|---|---|
| XR_001740740.1 | XM_001656940.2 |
| XR_001740741.1 | XR_002499532.1 |
| XR_001740748.2 | XM_021838787.1 |
| XR_001740760.2 | XM_021838646.1 |
| XR_001740761.1 | XM_001655133.2 |
| XR_001740807.1 | XM_021848205.1 |
| XR_001740819.1 | XM_001654940.2 |
| XR_001740824.1 | XM_021848164.1 |
| XR_001740850.2 | XM_021855718.1 |
| XR_001740939.1 | XR_002502987.1 |
| XR_001740980.1 | XM_021849885.1 |
| XR_001741071.1 | XM_021843742.1 |
| XR_001741091.2 | XR_002500706.1 |
| XR_001741131.2 | XM_021837251.1 |
| XR_001741136.1 | XM_021838301.1 |
| XR_001741146.2 | XM_021851672.1 |
| XR_001741185.1 | XM_021854459.1 |
| XR_001741218.1 | XM_001649227.2 |
| XR_001741272.2 | XM_021839865.1 |
| XR_001741277.1 | XM_021856697.1 |
| XR_001741285.2 | XM_001657281.2 |
| XR_001741289.1 | XM_021855997.1 |
| XR_001741311.2 | XM_021844157.1 |
| XR_001741374.1 | XR_002499137.1 |
| XR_001741421.1 | XM_021846066.1 |
| XR_001741425.2 | XM_001656915.2 |
| XR_001741432.1 | XR_002501046.1 |
| XR_001741437.1 | XM_001651884.2 |
| XR_001741438.1 | XM_021846462.1 |
| XR_001741444.2 | XM_021848988.1 |
| XR_001741503.1 | XM_021851203.1 |
| XR_001741530.1 | XM_021849971.1 |
| XR_001741616.1 | XM_021841272.1 |
| XR_001741634.2 | XM_021851734.1 |
| XR_001741656.1 | XM_021839158.1 |
| XR_001741689.1 | XR_002499889.1 |
| XR_001741693.2 | XM_021855596.1 |
| XR_001741694.1 | XM_021847603.1 |
| XR_001741697.1 | XR_002498761.1 |
| XR_001741704.1 | XM_021846612.1 |
| XR_001741729.1 | XM_021856130.1 |
| XR_001741733.1 | XM_021849529.1 |
| XR_001741735.1 | XM_001654328.2 |
| XR_001741738.1 | XM_021839167.1 |
| XR_001741745.2 | XM_021854599.1 |
| XR_001741796.1 | XM_021855421.1 |
| XR_001741816.1 | XM_021840407.1 |
| XR_001741823.2 | XM_021845880.1 |
| XR_001741861.1 | XM_021852681.1 |
| XR_001741884.2 | XR_002499103.1 |

| | |
|---|---|
| XR_001741885.1 | XM_001651085.2 |
| XR_001741907.2 | XR_002502578.1 |
| XR_001741914.1 | XM_021849485.1 |
| XR_001741920.1 | XR_002501595.1 |
| XR_001741929.1 | XM_021855937.1 |
| XR_001741933.1 | XM_021850981.1 |
| XR_001741943.2 | XM_021849337.1 |
| XR_001741949.1 | XM_001657587.2 |
| XR_001741972.2 | XM_021856321.1 |
| XR_001742013.2 | XM_021853939.1 |
| XR_001742017.2 | XR_002499876.1 |
| XR_001742018.1 | XM_021840700.1 |
| XR_001742021.1 | XM_001657816.2 |
| XR_001742039.1 | XM_021856609.1 |
| XR_001742067.2 | XM_001652590.2 |
| XR_001742172.1 | XM_021847881.1 |
| XR_001742177.1 | XM_001654863.2 |
| XR_001742212.1 | XM_021855131.1 |
| XR_001742346.1 | XM_021843080.1 |
| XR_001742390.2 | XM_001655921.2 |
| XR_001742407.1 | XM_021857045.1 |
| XR_001742420.2 | XM_001648242.2 |
| XR_001742426.1 | XM_001648865.2 |
| XR_001742431.1 | XM_021846085.1 |
| XR_001742448.1 | XM_021846313.1 |
| XR_001742489.1 | XM_001649280.2 |
| XR_001742523.1 | XR_002501067.1 |
| XR_001742562.1 | XM_021838665.1 |
| XR_001742570.1 | XM_001653617.2 |
| XR_001742578.1 | XM_021856796.1 |
| XR_001742597.1 | XM_001663732.2 |
| XR_001742607.1 | XM_021854593.1 |
| XR_001742616.1 | XM_001655406.2 |
| XR_001742674.1 | XR_002503157.1 |
| XR_001742688.1 | XM_021853889.1 |
| XR_001742709.1 | XM_001653067.2 |
| XR_001742725.2 | XM_021840395.1 |
| XR_001742804.1 | XM_021847133.1 |
| XR_001742807.1 | XM_001653995.2 |
| XR_001742811.2 | XM_021838515.1 |
| XR_001742822.1 | XM_001649928.2 |
| XR_001742867.1 | XR_002500115.1 |
| XR_001742868.1 | XR_002498720.1 |
| XR_001742905.1 | XM_001650986.2 |
| XR_001742913.1 | XM_021846230.1 |
| XR_001742931.1 | XM_021853000.1 |
| XR_001742941.1 | XM_021853001.1 |
| XR_001742945.1 | XR_002499130.1 |
| XR_001742960.1 | XM_001664089.2 |
| XR_001742987.1 | XM_021847073.1 |

| | |
|---|---|
| XR_001742990.1 | XM_021845657.1 |
| XR_001743034.1 | XM_021838074.1 |
| XR_001743048.1 | XM_021854245.1 |
| XR_001743180.2 | XM_021856071.1 |
| XR_001743200.1 | XM_021844926.1 |
| XR_001743223.2 | XM_021841827.1 |
| XR_001743233.2 | XM_021849840.1 |
| XR_001743243.1 | XM_021847594.1 |
| XR_001743262.2 | XM_021844915.1 |
| XR_001743302.1 | XM_021844618.1 |
| XR_001743308.2 | XM_021855145.1 |
| XR_001743453.1 | XM_021854841.1 |
| XR_001743487.2 | XR_002498632.1 |
| XR_001743504.1 | XR_002499274.1 |
| XR_001743623.2 | XM_021839295.1 |
| XR_001743626.2 | XM_001658271.2 |
| XR_001743696.2 | XR_002503120.1 |
| XR_001743702.1 | XM_001653720.2 |
| XR_001743720.1 | XM_021854442.1 |
| XR_001743723.1 | XM_021849804.1 |
| XR_001743726.2 | XM_021857631.1 |
| XR_001743790.1 | XR_002498824.1 |
| XR_001743815.1 | XM_021846459.1 |
| XR_001743903.1 | XM_021839103.1 |
| XR_001743911.2 | XM_001657699.2 |
| XR_001743912.2 | XM_021841911.1 |
| XR_001743913.1 | XR_002498584.1 |
| XR_001743918.1 | XM_001663070.2 |
| XR_001743956.1 | XM_021844916.1 |
| XR_001743981.1 | XM_021845422.1 |
| XR_001743990.1 | XR_002502919.1 |
| XR_001744027.1 | XM_001658879.2 |
| XR_001744091.1 | XM_021852559.1 |
| XR_001744099.1 | XM_001653305.2 |
| XR_001744157.1 | XM_021849288.1 |
| XR_001744160.1 | XR_002501101.1 |
| XR_001744190.1 | XM_001648078.3 |
| XR_001744216.2 | XM_021854551.1 |
| XR_001744257.1 | XM_021846918.1 |
| XR_001744333.1 | XM_021844920.1 |
| XR_001744360.2 | XM_021851517.1 |
| XR_001744374.1 | XM_021854489.1 |
| XR_001744391.1 | XM_001663754.2 |
| XR_001744393.1 | XM_021841876.1 |
| XR_001744397.2 | XM_021838692.1 |
| XR_001744408.1 | XM_021838815.1 |
| XR_001744418.1 | XR_002500460.1 |
| XR_001744431.1 | XM_021837453.1 |
| XR_001744446.1 | XM_001659468.2 |
| XR_001744482.1 | XM_001653619.2 |

| | |
|---|---|
| XR_001744514.1 | XM_021854639.1 |
| XR_001744601.2 | XR_002499676.1 |
| XR_001744694.1 | XM_021842749.1 |
| XR_001744731.2 | XM_021856619.1 |
| XR_001744750.1 | XM_001661607.2 |
| XR_001744811.1 | XM_021852634.1 |
| XR_001744812.1 | XM_021837524.1 |
| XR_001744823.1 | XM_021853776.1 |
| XR_001744840.1 | XM_021839166.1 |
| XR_001744887.2 | XR_002501263.1 |
| XR_001744889.2 | XM_001664080.2 |
| XR_001744898.2 | XM_021846456.1 |
| XR_001744903.2 | XM_021838502.1 |
| XR_001744938.1 | XM_001653057.2 |
| XR_001744943.2 | XM_021851883.1 |
| XR_001745010.1 | XM_021842262.1 |
| XR_001745019.1 | XM_021851882.1 |
| XR_001745068.1 | XM_021837403.1 |
| XR_001745070.2 | XM_021854842.1 |
| XR_001745074.1 | XM_021847598.1 |
| XR_001745083.2 | XM_021855242.1 |
| XR_001745090.1 | XM_021843594.1 |
| XR_001745096.1 | XR_002499891.1 |
| XR_001745183.1 | XM_021838664.1 |
| XR_001745186.1 | XM_021840514.1 |
| XR_001745257.1 | XR_002499102.1 |
| XR_001745273.1 | XR_002502033.1 |
| XR_001745332.1 | XM_021837681.1 |
| XR_001745352.1 | XM_021849522.1 |
| XR_001745364.1 | XM_001651043.2 |
| XR_001745380.1 | XM_001663668.3 |
| XR_001745384.1 | XM_021847644.1 |
| XR_001745422.1 | XM_021847345.1 |
| XR_001745496.1 | XR_002502808.1 |
| XR_001745506.2 | XM_021838820.1 |
| XR_001745508.2 | XM_001661622.2 |
| XR_001745533.1 | XM_021838349.1 |
| XR_001745554.2 | XM_021854281.1 |
| XR_001745582.1 | XM_001657219.2 |
| XR_001745593.1 | XM_021849605.1 |
| XR_001745611.2 | XM_021856608.1 |
| XR_001745613.2 | XM_001662007.2 |
| XR_001745629.2 | XM_001657181.2 |
| XR_001745651.2 | XM_001654198.2 |
| XR_001745657.1 | XM_001657837.2 |
| XR_001745684.1 | XM_021838434.1 |
| XR_001745705.1 | XM_021856968.1 |
| XR_001745709.1 | XM_021849878.1 |
| XR_001745769.1 | XM_021849706.1 |
| XR_001745779.1 | XR_002501251.1 |

| | |
|---|---|
| XR_001745885.1 | XM_021857099.1 |
| XR_001745914.1 | XM_001653056.2 |
| XR_001745965.1 | XM_021837715.1 |
| XR_001745973.1 | XR_002500503.1 |
| XR_001746017.1 | XM_021845167.1 |
| XR_001746060.1 | XM_021852720.1 |
| XR_001746092.2 | XM_021842007.1 |
| XR_001746128.1 | XM_001658518.2 |
| XR_001746129.1 | XM_021853426.1 |
| XR_001746150.2 | XM_021844029.1 |
| XR_001746162.2 | XM_001649859.2 |
| XR_001746166.2 | XM_021856879.1 |
| XR_001746272.1 | XM_021846237.1 |
| XR_001746310.2 | XM_021855610.1 |
| XR_001746329.2 | XM_021838957.1 |
| XR_001746345.1 | XM_021852871.1 |
| XR_001746417.2 | XM_021838549.1 |
| XR_001746431.1 | XR_002502560.1 |
| XR_001746459.1 | XM_021844963.1 |
| XR_001746478.2 | XM_021840631.1 |
| XR_001746499.1 | XM_021837641.1 |
| XR_001746540.2 | XR_002502292.1 |
| XR_001746542.2 | XM_021851322.1 |
| XR_001746556.2 | XM_021849735.1 |
| XR_001746565.1 | XM_021855343.1 |
| XR_001746574.1 | XR_002502432.1 |
| XR_001746577.1 | XR_002501324.1 |
| XR_001746613.2 | XM_021838081.1 |
| XR_001746656.1 | XR_002502639.1 |
| XR_001746667.1 | XM_021840307.1 |
| XR_001746681.1 | XR_002502677.1 |
| XR_001746684.2 | XM_001653729.2 |
| XR_001746690.1 | XM_021844922.1 |
| XR_001746718.1 | XM_021838538.1 |
| XR_001746727.1 | XM_021837761.1 |
| XR_001746756.1 | XM_021840633.1 |
| XR_001746760.1 | XM_001661354.2 |
| XR_001746839.1 | XM_001659222.2 |
| XR_001746844.1 | XM_021857618.1 |
| XR_001746887.1 | XM_001651280.3 |
| XR_001746907.1 | XM_021846685.1 |
| XR_001746914.1 | XM_001655423.2 |
| XR_001746931.1 | XM_001653857.2 |
| XR_001746932.1 | XM_021848901.1 |
| XR_001746934.1 | XM_021847601.1 |
| XR_001746963.1 | XM_001661839.2 |
| XR_001746986.1 | XM_021857574.1 |
| XR_001747015.1 | XM_021853384.1 |
| XR_001747016.1 | XM_001657776.2 |
| XR_001747096.1 | XM_021845554.1 |

| | |
|---|---|
| XR_001747123.2 | XM_021838289.1 |
| XR_001747173.1 | XM_021845709.1 |
| XR_001747197.2 | XM_021845660.1 |
| XR_001747218.1 | XM_021855405.1 |
| XR_001747221.1 | XM_001648407.2 |
| XR_001747223.1 | XM_021839321.1 |
| XR_001747227.1 | XM_021845359.1 |
| XR_001747229.1 | XM_021837722.1 |
| XR_001747250.1 | XM_021839749.1 |
| XR_001747275.1 | XM_021838359.1 |
| XR_001747280.1 | XR_002499186.1 |
| XR_001747282.1 | XM_021845653.1 |
| XR_001747293.1 | XM_021841843.1 |
| XR_001747319.2 | XR_002502051.1 |
| XR_001747335.1 | XR_002501966.1 |
| XR_001747343.1 | XM_021846524.1 |
| XR_001747348.1 | XM_021840571.1 |
| XR_001747359.1 | XM_021849520.1 |
| XR_001747385.2 | XM_021839723.1 |
| XR_001747416.1 | XM_021840559.1 |
| XR_001747428.1 | XM_021843073.1 |
| XR_001747431.1 | XM_021849594.1 |
| XR_001747434.1 | XM_021849853.1 |
| XR_001747479.1 | XM_021837569.1 |
| XR_001747484.2 | XM_001648240.2 |
| XR_001747488.2 | XM_021839169.1 |
| XR_001747494.1 | XM_021843135.1 |
| XR_001747560.2 | XM_021840394.1 |
| XR_001747561.1 | XM_021837568.1 |
| XR_001747568.1 | XM_021855126.1 |
| XR_001747602.1 | XM_021840173.1 |
| XR_001747603.1 | XM_021844593.1 |
| XR_001747616.1 | XM_021851670.1 |
| XR_001747657.1 | XM_021855322.1 |
| XR_001747753.1 | XR_002502697.1 |
| XR_001747852.1 | XR_002501339.1 |
| XR_001747896.1 | XM_021847616.1 |
| XR_001747911.2 | XR_002498716.1 |
| XR_001747931.1 | XM_001653685.2 |
| XR_001747949.2 | XR_002499313.1 |
| XR_001747953.2 | XM_001650619.2 |
| XR_001748011.2 | XM_001649738.2 |
| XR_001748053.2 | XM_021842750.1 |
| XR_001748059.1 | XR_002499249.1 |
| XR_001748066.1 | XM_021851395.1 |
| XR_001748072.1 | XR_002498753.1 |
| XR_001748092.1 | XM_021855530.1 |
| XR_001748099.2 | XM_021846404.1 |
| XR_001748108.2 | XM_021848074.1 |
| XR_001748163.1 | XM_001648605.2 |

| | |
|---|---|
| XR_001748164.1 | XM_021855792.1 |
| XR_001748167.1 | XM_021837560.1 |
| XR_001748204.2 | XM_021842703.1 |
| XR_001748227.1 | XM_001651988.2 |
| XR_001748242.1 | XM_021840608.1 |
| XR_001748254.1 | XM_001652746.2 |
| XR_001748279.1 | XM_001652244.2 |
| XR_001748294.1 | XM_001651062.2 |
| XR_001748325.1 | XM_001659690.2 |
| XR_001748327.1 | XM_021838496.1 |
| XR_001748341.1 | XM_001648918.2 |
| XR_001748349.1 | XR_002502658.1 |
| XR_001748376.1 | XR_002502265.1 |
| XR_001748382.1 | XM_001662090.2 |
| XR_001748384.1 | XM_021842275.1 |
| XR_001748400.1 | XM_021841130.1 |
| XR_001748439.1 | XM_021842857.1 |
| XR_001748456.1 | XM_021837249.1 |
| XR_001748469.1 | XM_021841374.1 |
| XR_001748486.1 | XM_001661748.2 |
| XR_001748580.2 | XM_021849017.1 |
| XR_001748606.2 | XM_001649842.2 |
| XR_001748628.1 | XR_002499639.1 |
| XR_001748691.1 | XM_001649915.2 |
| XR_001748700.2 | XM_021845656.1 |
| XR_001748726.2 | XM_001658911.2 |
| XR_001748728.1 | XM_021851043.1 |
| XR_001748820.1 | XR_002498862.1 |
| XR_001748839.1 | XR_002502642.1 |
| XR_001748843.1 | XR_002498776.1 |
| XR_001748866.2 | XM_021841869.1 |
| XR_001748897.2 | XM_021848850.1 |
| XR_001748898.2 | XM_021842261.1 |
| XR_001748905.2 | XM_001656746.2 |
| XR_001748933.2 | XM_001649460.2 |
| XR_001748946.1 | XM_021857712.1 |
| XR_001749033.1 | XR_002500673.1 |
| XR_001749039.1 | XM_001661822.2 |
| XR_001749071.1 | XM_021854894.1 |
| XR_001749095.1 | XM_021847111.1 |
| XR_001749102.1 | XM_001655763.2 |
| XR_001749108.1 | XR_002502710.1 |
| XR_001749114.1 | XM_021837599.1 |
| XR_001749127.2 | XM_021840678.1 |
| XR_001749131.1 | XR_002498707.1 |
| XR_001749146.1 | XM_001655511.2 |
| XR_001749251.1 | XM_001658844.2 |
| XR_001749260.1 | XM_021837603.1 |
| XR_001749305.1 | XM_001656499.2 |
| XR_001749346.2 | XM_021851866.1 |

| | |
|---|---|
| XR_001749378.1 | XR_002499743.1 |
| XR_001749394.1 | XR_002501924.1 |
| XR_001749430.1 | XM_021840007.1 |
| XR_001749585.2 | XM_021846000.1 |
| XR_001749614.2 | XM_021845987.1 |
| XR_001749668.2 | XM_021839018.1 |
| XR_001749717.2 | XM_021838497.1 |
| XR_001749854.1 | XM_021843961.1 |
| XR_001749883.1 | XM_001650170.2 |
| XR_001749924.1 | XM_001655193.2 |
| XR_001749952.1 | XM_021840172.1 |
| XR_001749996.1 | XM_001653421.2 |
| XR_001750001.1 | XM_021852801.1 |
| XR_001750008.1 | XM_021851982.1 |
| XR_001750018.1 | XM_021844249.1 |
| XR_001750024.1 | XM_001663866.2 |
| XR_001750181.2 | XM_001656713.2 |
| XR_001750279.1 | XM_001650151.2 |
| XR_001750360.1 | XM_021843171.1 |
| XR_001750400.1 | XM_001657313.2 |
| XR_001750415.1 | XM_021845812.1 |
| XR_001750478.2 | XM_001658363.2 |
| XR_001750482.1 | XM_001664001.2 |
| XR_001750510.2 | XM_021852496.1 |
| XR_001750605.1 | XM_001657576.3 |
| XR_001750621.1 | XM_021838220.1 |
| XR_001750626.1 | XM_021846476.1 |
| XR_001750673.1 | XM_021852280.1 |
| XR_001750696.1 | XM_021841871.1 |
| XR_001750733.1 | XM_001657695.2 |
| XR_001750736.1 | XM_021852160.1 |
| XR_001750758.1 | XM_021851039.1 |
| XR_001750759.1 | XM_021853423.1 |
| XR_001750777.1 | XR_002499229.1 |
| XR_001750780.1 | XM_001652399.2 |
| XR_001750788.2 | XM_001648473.2 |
| XR_001750796.1 | XM_001649528.2 |
| XR_001750818.2 | XR_002499690.1 |
| XR_001750836.2 | XM_021839698.1 |
| XR_001750866.1 | XM_001661405.2 |
| XR_001750871.2 | XR_002501568.1 |
| XR_001750891.1 | XM_001656024.2 |
| XR_001750945.1 | XM_021838077.1 |
| XR_001750963.1 | XM_021849482.1 |
| XR_001751016.1 | XM_001657056.2 |
| XR_001751081.1 | XM_021848317.1 |
| XR_001751098.2 | XM_001651845.2 |
| XR_001751142.1 | XR_002499095.1 |
| XR_001751181.1 | XM_001660686.2 |
| XR_001751221.2 | XM_021837276.1 |

| | |
|---|---|
| XR_001751298.2 | XR_002501921.1 |
| XR_001751311.1 | XM_001653987.2 |
| XR_001751318.1 | XM_021837248.1 |
| XR_001751319.2 | XM_001650112.2 |
| XR_001751333.2 | XR_002503240.1 |
| XR_001751379.2 | XM_021855323.1 |
| XR_001751381.1 | XM_021849859.1 |
| XR_001751400.1 | XM_001658617.2 |
| XR_001751523.1 | XM_021843061.1 |
| XR_001751554.2 | XM_001662359.2 |
| XR_001751587.1 | XR_002499691.1 |
| XR_001751594.1 | XM_021855124.1 |
| XR_001751607.1 | XM_021856020.1 |
| XR_001751613.2 | XM_021841377.1 |
| XR_001751636.1 | XM_021840044.1 |
| XR_001751683.1 | XR_002502528.1 |
| XR_001751694.1 | XM_001652146.2 |
| XR_001751696.1 | XM_021847813.1 |
| XR_001751763.2 | XM_021839867.1 |
| XR_001751767.2 | XM_021845457.1 |
| XR_001751795.1 | XM_001663283.2 |
| XR_001751806.1 | XM_021849470.1 |
| XR_001751818.2 | XM_021853048.1 |
| XR_001751827.2 | XM_021851548.1 |
| XR_001751838.1 | XM_021848836.1 |
| XR_001751881.2 | XM_021851929.1 |
| XR_001751886.1 | XM_021841216.1 |
| XR_001751895.1 | XM_001654936.2 |
| XR_001751907.2 | XR_002501125.1 |
| XR_001751939.1 | XM_021837333.1 |
| XR_001751942.1 | XR_002502170.1 |
| XR_001751948.1 | XM_021845055.1 |
| XR_001751996.1 | XM_021838670.1 |
| XR_001752064.1 | XM_021838293.1 |
| XR_001752066.1 | XM_021845458.1 |
| XR_001752082.1 | XR_002501154.1 |
| XR_001752085.2 | XR_002502925.1 |
| XR_001752103.1 | XM_021841923.1 |
| XR_001752119.2 | XM_021838941.1 |
| XR_001752179.1 | XM_021845950.1 |
| XR_001752213.1 | XM_001653993.2 |
| XR_001752214.1 | XM_021840775.1 |
| XR_001752219.1 | XM_021843129.1 |
| XR_001752224.1 | XM_001655478.2 |
| XR_001752225.1 | XM_021839218.1 |
| XR_001752227.1 | XM_021855150.1 |
| XR_001752232.1 | XM_001652057.2 |
| XR_001752236.1 | XM_021849546.1 |
| XR_001752238.1 | XM_021841441.1 |
| XR_001752250.1 | XM_001662421.2 |

| | |
|---|---|
| XR_001752271.2 | XM_001653580.2 |
| XR_001752336.1 | XM_001663378.2 |
| XR_001752359.1 | XM_021840077.1 |
| XR_001752394.2 | XM_021851232.1 |
| XR_001752486.2 | XM_021845936.1 |
| XR_001752487.1 | XM_001661586.2 |
| XR_001752495.1 | XM_021854594.1 |
| XR_001752509.2 | XM_021856345.1 |
| XR_001752529.1 | XM_021853301.1 |
| XR_001752545.1 | XM_001653777.2 |
| XR_001752558.1 | XM_021857639.1 |
| XR_001752594.1 | XM_021838276.1 |
| XR_001752604.2 | XM_021839618.1 |
| XR_001752626.2 | XM_021849457.1 |
| XR_001752661.2 | XM_021854173.1 |
| XR_001752664.1 | XM_021853276.1 |
| XR_001752677.2 | XM_021846450.1 |
| XR_001752679.1 | XM_001652704.2 |
| XR_001752687.2 | XM_001654156.2 |
| XR_001752688.2 | XR_002502805.1 |
| XR_001752722.2 | XM_001659083.3 |
| XR_001752740.1 | XM_021838292.1 |
| XR_001752749.1 | XM_021838204.1 |
| XR_001752814.2 | XM_021854587.1 |
| XR_001752868.1 | XM_001651800.2 |
| XR_001752870.1 | XM_021845879.1 |
| XR_001752882.2 | XR_002502104.1 |
| XR_001752918.2 | XM_021851044.1 |
| XR_001752972.2 | XM_021841080.1 |
| XR_001752980.2 | XM_021842479.1 |
| XR_001752996.1 | XM_021857634.1 |
| XR_001753013.2 | XM_021844658.1 |
| XR_001753025.1 | XM_021847285.1 |
| XR_001753096.1 | XM_021854641.1 |
| XR_001753115.2 | XM_021854907.1 |
| XR_001753127.1 | XM_001657779.2 |
| XR_001753138.1 | XM_001648520.2 |
| XR_001753166.1 | XM_021846830.1 |
| XR_001753189.1 | XM_021851252.1 |
| XR_001753260.2 | XM_001660368.2 |
| XR_001753324.1 | XM_021856864.1 |
| XR_001753345.1 | XM_021841446.1 |
| XR_001753371.1 | XR_002500262.1 |
| XR_001753374.1 | XM_021838688.1 |
| XR_001753375.1 | XM_001658390.2 |
| XR_001753379.2 | XM_021848076.1 |
| XR_001753382.1 | XM_001659049.2 |
| XR_001753404.1 | XM_001663952.2 |
| XR_001753412.1 | XM_021853419.1 |
| XR_001753427.1 | XR_002502289.1 |

| | |
|---|---|
| XR_001753435.1 | XR_002501325.1 |
| XR_001753442.1 | XR_002499754.1 |
| XR_001753467.1 | XM_021850024.1 |
| XR_001753495.1 | XM_001664123.2 |
| XR_001753503.1 | XM_021849712.1 |
| XR_001753542.1 | XM_001658796.2 |
| XR_001753549.1 | XM_001653334.2 |
| XR_001753552.1 | XM_021856419.1 |
| XR_001753553.1 | XM_021851248.1 |
| XR_001753567.2 | XR_002501806.1 |
| XR_001753614.2 | XM_021842799.1 |
| XR_001753642.1 | XM_021854813.1 |
| XR_001753661.1 | XM_001652895.2 |
| XR_001753662.1 | XM_021839296.1 |
| XR_001753663.2 | XM_021843955.1 |
| XR_001753665.2 | XM_001654611.2 |
| XR_001753675.1 | XM_021856709.1 |
| XR_001753702.1 | XR_002500636.1 |
| XR_001753751.1 | XM_021838191.1 |
| XR_001753840.1 | XM_021839122.1 |
| XR_001753866.1 | XM_001650584.2 |
| XR_001753920.1 | XM_021848860.1 |
| XR_001753961.1 | XM_021854589.1 |
| XR_001754017.1 | XR_002500298.1 |
| XR_001754024.1 | XM_021846688.1 |
| XR_001754058.1 | XR_002501352.1 |
| XR_001754062.1 | XM_001647564.2 |
| XR_001754077.1 | XM_021855348.1 |
| XR_001754085.1 | XM_001652775.2 |
| XR_001754187.1 | XM_021855340.1 |
| XR_001754252.1 | XM_001659833.2 |
| XR_001754294.1 | XR_002501145.1 |
| XR_001754324.1 | XR_002502552.1 |
| XR_001754379.1 | XM_021840008.1 |
| XR_001754412.1 | XM_001656887.2 |
| XR_001754443.2 | XM_021838704.1 |
| XR_001754491.1 | XR_002500724.1 |
| XR_001754493.1 | XM_001648259.2 |
| XR_001754526.1 | XR_002501258.1 |
| XR_001754532.1 | XM_001653646.2 |
| XR_001754556.2 | XM_021850663.1 |
| XR_001754560.1 | XM_021852341.1 |
| XR_001754596.1 | XM_001649218.2 |
| XR_001754618.1 | XM_001654203.2 |
| XR_001754623.1 | XM_001652058.2 |
| XR_001754634.2 | XM_001663791.2 |
| XR_001754691.1 | XM_021838035.1 |
| XR_001754738.1 | XM_001660118.2 |
| XR_001754902.2 | XM_021856282.1 |
| XR_001754920.2 | XM_001662816.2 |

| | |
|---|---|
| XR_001754963.2 | XM_021851250.1 |
| XR_001754973.1 | XM_021851389.1 |
| XR_001754982.1 | XM_021840312.1 |
| XR_001754983.1 | XM_001652166.2 |
| XR_001754994.1 | XM_001648785.2 |
| XR_001754995.1 | XM_021854272.1 |
| XR_001755040.1 | XR_002501072.1 |
| XR_001755082.2 | XM_001655379.2 |
| XR_001755190.1 | XM_021839846.1 |
| XR_001755232.1 | XM_021844962.1 |
| XR_001755233.1 | XM_021841010.1 |
| XR_001755416.2 | XM_021854591.1 |
| XR_001755441.1 | XM_001658930.2 |
| XR_001755448.1 | XM_021846841.1 |
| XR_001755452.1 | XM_021843422.1 |
| XR_001755457.1 | XR_002501890.1 |
| XR_001755490.1 | XM_021853766.1 |
| XR_001755513.1 | XM_021838314.1 |
| XR_001755552.2 | XM_021847431.1 |
| XR_001755623.1 | XM_021843184.1 |
| XR_001755679.1 | XM_001656397.2 |
| XR_001755707.2 | XM_001647970.2 |
| XR_001755714.1 | XR_002499765.1 |
| XR_001755718.2 | XM_021852356.1 |
| XR_001755743.1 | XM_001662860.2 |
| XR_001755765.1 | XM_021848391.1 |
| XR_001755783.1 | XR_002500543.1 |
| XR_001755784.1 | XM_021837716.1 |
| XR_001755790.1 | XM_001654927.2 |
| XR_001755798.1 | XM_001649552.2 |
| XR_001755858.1 | XM_021851921.1 |
| XR_001755861.2 | XM_001663674.2 |
| XR_001755870.1 | XM_021855274.1 |
| XR_001755898.1 | XM_021849685.1 |
| XR_001755913.1 | XM_021847057.1 |
| XR_001755922.1 | XR_002503165.1 |
| XR_001755939.1 | XM_021856389.1 |
| XR_001755967.1 | XR_002501045.1 |
| XR_001755969.1 | XM_021843066.1 |
| XR_001756037.1 | XM_021851689.1 |
| XR_001756056.1 | XM_021841603.1 |
| XR_001756065.1 | XR_002500691.1 |
| XR_001756077.1 | XM_021855871.1 |
| XR_001756087.2 | XM_021852641.1 |
| XR_001756115.1 | XR_002499688.1 |
| XR_001756128.1 | XR_002499519.1 |
| XR_001756135.1 | XM_021854586.1 |
| XR_001756158.2 | XM_001653835.2 |
| XR_001756170.1 | XM_021847058.1 |
| XR_001756188.1 | XM_001662525.2 |

| | |
|---|---|
| XR_001756224.1 | XM_021848079.1 |
| XR_001756227.1 | XM_021851393.1 |
| XR_001756308.1 | XM_021842082.1 |
| XR_001756322.1 | XM_021851919.1 |
| XR_001756339.1 | XR_002502509.1 |
| XR_001756347.1 | XM_021838076.1 |
| XR_001756355.1 | XM_021853278.1 |
| XR_001756371.1 | XM_001662111.2 |
| XR_001756382.1 | XM_001650810.2 |
| XR_001756402.1 | XR_002500995.1 |
| XR_001756421.2 | XM_001656052.2 |
| XR_001756422.2 | XR_002499272.1 |
| XR_001756455.1 | XM_021849262.1 |
| XR_001756475.1 | XR_002498786.1 |
| XR_001756486.1 | XR_002501643.1 |
| XR_001756499.1 | XM_021842654.1 |
| XR_001756508.1 | XM_001650795.2 |
| XR_001756514.1 | XM_001660548.2 |
| XR_001756529.2 | XR_002502082.1 |
| XR_001756547.2 | XR_002498947.1 |
| XR_001756560.1 | XM_021841995.1 |
| XR_001756600.2 | XM_001648849.2 |
| XR_001756614.1 | XM_001654721.2 |
| XR_001756617.1 | XM_021845553.1 |
| XR_001756628.1 | XM_021852467.1 |
| XR_001756639.1 | XM_021845529.1 |
| XR_001756649.1 | XR_002498734.1 |
| XR_001756671.1 | XM_021841957.1 |
| XR_001756770.1 | XM_021844237.1 |
| XR_001756782.2 | XM_021853280.1 |
| XR_001756882.1 | XM_021856319.1 |
| XR_001756883.1 | XM_021843998.1 |
| XR_001756888.2 | XM_021840033.1 |
| XR_001756895.1 | XM_001660603.2 |
| XR_001756897.1 | XM_021846821.1 |
| XR_001756940.1 | XM_021855349.1 |
| XR_001756948.2 | XM_001652105.2 |
| XR_002956186.1 | XM_001663505.2 |
| XR_002956193.1 | XR_002501540.1 |
| XR_002956238.1 | XM_021853277.1 |
| XR_002956299.1 | XR_002499282.1 |
| XR_002956316.1 | XR_002501127.1 |
| XR_002956323.1 | XM_021850438.1 |
| XR_002956327.1 | XM_021855551.1 |
| XR_002956329.1 | XM_021853846.1 |
| XR_002956330.1 | XM_001648602.2 |
| XR_002956338.1 | XM_021840030.1 |
| XR_002956359.1 | XM_001648550.2 |
| XR_002956360.1 | XM_001662491.2 |
| XR_002956377.1 | XM_021838588.1 |

| | |
|---|---|
| XR_002956385.1 | XM_001653222.2 |
| XR_002956402.1 | XM_001651521.2 |
| XR_002956419.1 | XR_002499040.1 |
| XR_002956438.1 | XM_021857575.1 |
| XR_002956472.1 | XR_002499584.1 |
| XR_002956484.1 | XR_002502962.1 |
| XR_002956516.1 | XR_002500919.1 |
| XR_002956522.1 | XM_001655733.2 |
| XR_002956528.1 | XM_001660347.2 |
| XR_002956556.1 | XM_021839219.1 |
| XR_002956570.1 | XR_002500808.1 |
| XR_002956581.1 | XM_021838465.1 |
| XR_002956587.1 | XM_021852166.1 |
| XR_002956616.1 | XM_021849421.1 |
| XR_002956645.1 | XM_001651177.2 |
| XR_002956648.1 | XM_021856055.1 |
| XR_002956664.1 | XR_002501390.1 |
| XR_002956710.1 | XR_002499721.1 |
| XR_002956719.1 | XM_001650281.2 |
| XR_002956749.1 | XM_021850844.1 |
| XR_002956759.1 | XM_021839533.1 |
| XR_002956775.1 | XM_001654282.2 |
| XR_002956808.1 | XM_021846614.1 |
| XR_002956817.1 | XM_011494789.2 |
| XR_002956845.1 | XM_011495041.2 |
| XR_002956869.1 | XM_021855920.1 |
| XR_002956908.1 | XR_002500725.1 |
| XR_002956924.1 | XM_011494996.2 |
| XR_002956954.1 | XM_021837422.1 |
| XR_002956983.1 | XM_001657428.2 |
| XR_002957021.1 | XM_001651388.2 |
| XR_002957043.1 | XR_002499127.1 |
| XR_002957106.1 | XM_021846903.1 |
| XR_002957116.1 | XM_021841771.1 |
| XR_002957174.1 | XM_021838487.1 |
| XR_002957188.1 | XM_021849419.1 |
| XR_002957190.1 | XM_001654857.2 |
| XR_002957204.1 | XM_021857409.1 |
| XR_002957228.1 | XR_002503176.1 |
| XR_002957229.1 | XR_002501522.1 |
| XR_002957230.1 | XM_021846901.1 |
| XR_002957272.1 | XM_021846717.1 |
| XR_002957281.1 | XM_001659513.2 |
| XR_002957294.1 | XM_001653636.3 |
| XR_002957308.1 | XM_021856981.1 |
| XR_002957310.1 | XM_001652993.3 |
| XR_002957337.1 | XM_001652133.2 |
| XR_002957340.1 | XM_021856567.1 |
| XR_002957350.1 | XM_021851083.1 |
| XR_002957360.1 | XR_002498782.1 |

| | |
|---|---|
| XR_002957363.1 | XM_021857404.1 |
| XR_002957384.1 | XM_021842016.1 |
| XR_002957422.1 | XM_021845530.1 |
| XR_002957474.1 | XR_002501179.1 |
| XR_002957476.1 | XM_001658393.2 |
| XR_002957480.1 | XM_021856913.1 |
| XR_002957486.1 | XM_021840615.1 |
| XR_002957489.1 | XM_021850821.1 |
| XR_002957490.1 | XM_021850910.1 |
| XR_002957496.1 | XM_021846843.1 |
| XR_002957501.1 | XM_021845871.1 |
| XR_002957533.1 | XM_021839132.1 |
| XR_002957537.1 | XM_001651270.2 |
| XR_002957577.1 | XM_001655472.2 |
| XR_002957578.1 | XM_021856874.1 |
| XR_002957582.1 | XM_021847729.1 |
| XR_002957618.1 | XM_001648783.2 |
| XR_002957624.1 | XM_001661996.2 |
| XR_002957634.1 | XM_021848080.1 |
| XR_002957679.1 | XM_001655909.2 |
| XR_002957691.1 | XM_021841396.1 |
| XR_002957706.1 | XM_001664212.2 |
| XR_002957746.1 | XM_021856088.1 |
| XR_002957752.1 | XM_001662964.2 |
| XR_002957809.1 | XM_021854695.1 |
| XR_002957853.1 | XM_021841956.1 |
| XR_002957873.1 | XR_002500147.1 |
| XR_002957881.1 | XM_001660006.2 |
| XR_002957885.1 | XR_002501177.1 |
| XR_002957897.1 | XM_001657585.2 |
| XR_002957901.1 | XR_002500213.1 |
| XR_002957911.1 | XM_021841972.1 |
| XR_002957945.1 | XM_021841158.1 |
| XR_002957961.1 | XM_001652048.2 |
| XR_002958000.1 | XM_021857408.1 |
| XR_002958039.1 | XM_001654870.2 |
| XR_002958047.1 | XM_021848057.1 |
| XR_002958065.1 | XM_001654599.2 |
| XR_002958071.1 | XM_021841716.1 |
| XR_002958079.1 | XR_002499096.1 |
| XR_002958082.1 | XM_021847021.1 |
| XR_002958088.1 | XM_021846120.1 |
| XR_002958091.1 | XR_002499316.1 |
| XR_002958096.1 | XM_001663071.2 |
| XR_002958104.1 | XM_021856370.1 |
| XR_002958114.1 | XM_001657642.2 |
| XR_002958116.1 | XM_001659905.2 |
| XR_002958175.1 | XM_021857434.1 |
| XR_002958176.1 | XM_001651862.2 |
| XR_002958182.1 | XM_001647791.2 |

| | |
|---|---|
| XR_002958204.1 | XR_002498981.1 |
| XR_002958206.1 | XM_021838020.1 |
| XR_002958207.1 | XM_021854873.1 |
| XR_002958235.1 | XM_021845404.1 |
| XR_002958244.1 | XM_021844503.1 |
| XR_002958250.1 | XM_021845263.1 |
| XR_002958252.1 | XM_021841393.1 |
| XR_002958257.1 | XM_021855163.1 |
| XR_002958266.1 | XM_001653633.3 |
| XR_002958282.1 | XM_001656339.2 |
| XR_002958330.1 | XM_021839649.1 |
| XR_002958342.1 | XM_021857253.1 |
| XR_002958369.1 | XR_002501651.1 |
| XR_002958373.1 | XR_002501828.1 |
| XR_002958387.1 | XM_001653915.2 |
| XR_002958388.1 | XM_021850511.1 |
| XR_002958393.1 | XM_021839787.1 |
| XR_002958414.1 | XM_001653989.2 |
| XR_002958418.1 | XM_001653884.2 |
| XR_002958423.1 | XM_021851690.1 |
| XR_002958460.1 | XM_001651993.2 |
| XR_002958481.1 | XM_001660592.2 |
| XR_002958482.1 | XM_021855083.1 |
| XR_002958489.1 | XM_021841936.1 |
| XR_002958504.1 | XR_002501364.1 |
| XR_002958524.1 | XM_021849150.1 |
| XR_002958557.1 | XM_001653765.2 |
| XR_002958565.1 | XM_021852894.1 |
| XR_002958575.1 | XM_021848198.1 |
| XR_002958601.1 | XM_021840088.1 |
| XR_002958616.1 | XM_021846015.1 |
| XR_002958625.1 | XM_001649482.2 |
| XR_002958639.1 | XM_021837328.1 |
| XR_002958666.1 | XM_021853132.1 |
| XR_002958698.1 | XM_001656531.2 |
| XR_002958716.1 | XM_021846315.1 |
| XR_002958720.1 | XM_021846139.1 |
| XR_002958749.1 | XM_021838278.1 |
| XR_002958760.1 | XR_002499504.1 |
| XR_002958774.1 | XM_021849051.1 |
| XR_002958806.1 | XR_002502818.1 |
| XR_002958821.1 | XM_021841994.1 |
| XR_002958830.1 | XM_021843222.1 |
| XR_002958865.1 | XM_021852638.1 |
| XR_002958872.1 | XR_002500583.1 |
| XR_002958884.1 | XR_002503206.1 |
| XR_002958894.1 | XM_021846138.1 |
| XR_002958898.1 | XM_021848338.1 |
| XR_002958922.1 | XM_001661763.2 |
| XR_002958941.1 | XM_021851733.1 |

| | |
|---|---|
| XR_002958976.1 | XM_021841442.1 |
| XR_002959024.1 | XM_021845139.1 |
| XR_002959052.1 | XM_001653086.2 |
| XR_002959055.1 | XM_001649092.2 |
| XR_002959065.1 | XM_021847023.1 |
| XR_002959099.1 | XM_021854539.1 |
| XR_002959104.1 | XR_002501946.1 |
| XR_002959124.1 | XM_021853958.1 |
| XR_002959134.1 | XM_021852163.1 |
| XR_002959142.1 | XR_002499365.1 |
| XR_002959146.1 | XM_021856195.1 |
| XR_002959170.1 | XM_021853957.1 |
| XR_002959184.1 | XM_001651272.2 |
| XR_002959197.1 | XM_021847053.1 |
| XR_002959213.1 | XR_002501959.1 |
| XR_002959292.1 | XM_001661123.2 |
| XR_002959317.1 | XR_002500529.1 |
| XR_002959329.1 | XM_021852691.1 |
| XR_002959342.1 | XM_021852640.1 |
| XR_002959350.1 | XR_002502973.1 |
| XR_002959355.1 | XR_002502141.1 |
| XR_002959363.1 | XM_001656806.2 |
| XR_002959364.1 | XM_021840773.1 |
| XR_002959368.1 | XM_001648257.2 |
| XR_002959392.1 | XM_021854156.1 |
| XR_002959403.1 | XR_002501298.1 |
| XR_002959407.1 | XM_001663995.2 |
| XR_002959421.1 | XM_021847901.1 |
| XR_002959424.1 | XM_001655167.2 |
| XR_002959425.1 | XM_001659854.2 |
| XR_002959439.1 | XM_021840512.1 |
| XR_002959446.1 | XM_021845834.1 |
| XR_002959451.1 | XM_001661610.2 |
| XR_002959471.1 | XM_001662036.3 |
| XR_002959489.1 | XR_002498839.1 |
| XR_002959503.1 | XM_021851379.1 |
| XR_002959510.1 | XR_002501961.1 |
| XR_002959518.1 | XM_021844635.1 |
| XR_002959528.1 | XM_001649611.2 |
| XR_002959568.1 | XM_021850032.1 |
| XR_002959592.1 | XM_021850289.1 |
| XR_002959605.1 | XM_021850033.1 |
| XR_002959606.1 | XM_021839790.1 |
| XR_002959616.1 | XM_021857620.1 |
| XR_002959656.1 | XM_021841800.1 |
| XR_002959782.1 | XM_021842000.1 |
| XR_002959813.1 | XM_001660247.2 |
| XR_002959816.1 | XM_021855345.1 |
| XR_002959821.1 | XM_021849362.1 |
| XR_002959824.1 | XR_002501880.1 |

| | |
|---|---|
| XR_004643303.1 | XM_001657420.2 |
| XR_004837503.1 | XM_021841740.1 |
| XR_004837528.1 | XM_001649340.2 |
| XR_004837530.1 | XR_002501382.1 |
| XR_004837531.1 | XM_001657174.2 |
| XR_005228010.1 | XM_001649821.2 |
| XR_005228025.1 | XM_021839459.1 |
| XR_158942.3 | XM_001649696.2 |
| XR_171071.2 | XM_001649289.2 |
| XR_241302.2 | XM_021838370.1 |
| XR_241325.1 | XR_002498770.1 |
| XR_241866.3 | XM_001656916.2 |
| XR_241885.2 | XM_021842251.1 |
| XR_242213.1 | XR_002499527.1 |
| XR_242682.4 | XM_021857621.1 |
| XR_242777.3 | XR_002501186.1 |
| XR_242877.4 | XM_021844228.1 |
| XR_242895.4 | XM_001656164.2 |
| XR_243258.3 | XM_021849809.1 |
| XR_243305.3 | XM_001657961.3 |
| XR_243433.3 | XM_021852694.1 |
| XR_243875.3 | XR_002499011.1 |
| XR_244323.3 | XM_021837349.1 |
| XR_244356.3 | XM_021839271.1 |
| XR_244468.4 | XM_021851916.1 |
| XR_244887.5 | XM_021854601.1 |
| XR_244972.3 | XM_021851189.1 |
| XR_245020.3 | XM_021838382.1 |
| XR_245100.1 | XM_001660426.2 |
| XR_245182.3 | XM_001661618.2 |
| XR_245208.3 | XM_021841932.1 |
| XR_245442.5 | XM_001655564.2 |
| XR_245604.3 | XM_001652020.2 |
| XR_245695.2 | XM_001660270.2 |
| XR_245708.4 | XM_021840311.1 |
| XR_245788.2 | XM_001653769.2 |
| XR_246111.5 | XM_011494982.2 |
| XR_246197.2 | XM_001662377.2 |
| XR_246305.4 | XM_001663852.2 |
| XR_426708.3 | XM_021841567.1 |
| XR_426737.3 | XM_021847160.1 |
| XR_426846.4 | XM_021842683.1 |
| XR_426950.2 | XM_001655743.2 |
| XR_427017.3 | XR_002499292.1 |
| XR_427085.3 | XM_001663394.2 |
| XR_427149.4 | XM_021842006.1 |
| XR_427503.4 | XR_002499604.1 |
| XR_427775.3 | XM_021837284.1 |
| XR_427805.2 | XM_001662581.2 |
| XR_427811.3 | XM_021837767.1 |

| | |
|---|---|
| XR_427883.4 | XM_021842692.1 |
| XR_427919.3 | XM_021852695.1 |
| XR_428154.3 | XM_021851375.1 |
| XR_428319.2 | XM_021854530.1 |
| XR_428359.3 | XR_002501471.1 |
| XR_428520.3 | XR_002501174.1 |
| XR_428583.3 | XM_021852693.1 |
| XR_428650.1 | XM_021850340.1 |
| XR_428922.3 | XM_001652562.2 |
| XR_428971.3 | XM_001650364.2 |
| XR_428993.3 | XM_021852692.1 |
| XR_429016.3 | XR_002500197.1 |
| XR_429135.2 | XM_021840167.1 |
| XR_429192.4 | XM_001654226.2 |
| XR_429333.3 | XM_021839123.1 |
| XR_429360.2 | XR_002499401.1 |
| XR_429740.2 | XM_021837348.1 |
| XR_429875.3 | XR_002499327.1 |
| XR_429913.4 | XM_021856824.1 |
| XR_430031.3 | XM_001654740.2 |
| XR_430124.4 | XM_021857119.1 |
| XR_430202.4 | XM_001663079.2 |
| XR_430301.3 | XM_021854447.1 |
| XR_430308.3 | XR_002501367.1 |
| XR_430821.3 | XM_001661894.2 |
| XR_922018.2 | XM_001653349.2 |
| XR_922067.2 | XR_002499689.1 |
| XR_922140.2 | XM_021855030.1 |
| XR_922148.2 | XM_001655168.2 |
| XR_922160.2 | XM_021839345.1 |
| XR_922213.1 | XM_021843206.1 |
| XR_922299.3 | XM_021857057.1 |
| XR_922306.2 | XM_021853150.1 |
| XR_922317.2 | XM_001662129.2 |
| XR_922341.2 | XM_021855869.1 |
| XR_922454.2 | XM_021843516.1 |
| XR_922459.2 | XM_001660098.2 |
| XR_922472.1 | XM_021841159.1 |
| XR_922508.2 | XM_021840861.1 |
| XR_922530.3 | XM_021856867.1 |
| XR_922599.1 | XM_001657368.2 |
| XR_922623.2 | XR_002500355.1 |
| XR_922667.1 | XM_021845165.1 |
| XR_922705.3 | XM_001654193.2 |
| XR_922815.2 | XR_002500044.1 |
| XR_922879.2 | XR_002503081.1 |
| XR_923022.3 | XM_021852688.1 |
| XR_923082.1 | XM_021850176.1 |
| XR_923085.2 | XM_001649937.2 |
| XR_923090.2 | XM_021848155.1 |

| | |
|---|---|
| XR_923105.2 | XR_002500533.1 |
| XR_923108.2 | XM_001648019.2 |
| XR_923125.1 | XM_021852972.1 |
| XR_923137.2 | XM_021851726.1 |
| XR_923142.2 | XM_001653472.2 |
| XR_923144.2 | XM_021848156.1 |
| XR_923175.2 | XM_021841610.1 |
| XR_923250.2 | XM_021854603.1 |
| XR_923288.1 | XM_001648287.2 |
| XR_923296.1 | XM_021856875.1 |
| XR_923388.1 | XM_021857638.1 |
| XR_923505.2 | XM_021845163.1 |
| XR_923509.2 | XM_001647914.2 |
| XR_923527.1 | XM_021854540.1 |
| XR_923591.3 | XM_001656992.2 |
| XR_923624.2 | XM_021849185.1 |
| XR_923747.2 | XM_001654361.2 |
| XR_923757.2 | XM_001660868.2 |
| XR_923855.2 | XM_001659205.2 |
| XR_923914.2 | XM_021852635.1 |
| XR_923916.2 | XM_021852035.1 |
| XR_923917.2 | XM_021838779.1 |
| XR_923954.1 | XR_002501359.1 |
| XR_924046.2 | XM_001654190.2 |
| XR_924073.3 | XR_002499726.1 |
| XR_924097.2 | XM_021846121.1 |
| XR_924162.3 | XM_021854605.1 |
| XR_924191.3 | XM_001656340.2 |
| XR_924256.2 | XR_002499641.1 |
| XR_924302.2 | XM_021851749.1 |
| XR_924434.1 | XM_001655610.2 |
| XR_924555.2 | XM_021839346.1 |
| XR_924563.2 | XM_001659726.2 |
| XR_924578.2 | XM_001662220.2 |
| XR_924807.2 | XM_001652992.3 |
| XR_924819.2 | XM_021837281.1 |
| XR_925034.2 | XM_021840899.1 |
| XR_925064.2 | XM_021853802.1 |
| XR_925198.1 | XM_021838248.1 |
| XR_925203.2 | XM_021839281.1 |
| XR_925257.3 | XM_021840941.1 |
| XR_925305.2 | XM_021842653.1 |
| XR_925402.2 | XM_021846451.1 |
| XR_925515.2 | XM_021848837.1 |
| XR_925530.2 | XM_001658159.2 |
| XR_925579.2 | XR_002499030.1 |
| XR_925644.2 | XM_021842178.1 |
| XR_925714.1 | XM_021847258.1 |
| XR_925762.2 | XM_021855389.1 |
| XR_925792.2 | XR_002499877.1 |

| | |
|---|---|
| XR_925956.3 | XR_002499955.1 |
| XR_926044.1 | XR_002502781.1 |
| XR_926099.2 | XR_002499187.1 |
| XR_926141.2 | XM_021840483.1 |
| XR_926166.2 | XM_001661339.2 |
| XR_926267.2 | XR_002501224.1 |
| XR_926402.2 | XM_021847950.1 |
| XR_926458.2 | XM_021852687.1 |
| XR_926469.2 | XM_021845851.1 |
| XR_926618.1 | XM_021849180.1 |
| XR_926620.2 | XM_021847161.1 |
| XR_926710.1 | XM_011494657.2 |
| XR_926747.2 | XR_002499053.1 |
| XR_926753.2 | XM_001664022.2 |
| XR_926757.2 | XM_021847306.1 |
| XR_926762.2 | XM_001656288.2 |
| XR_926780.2 | XM_001651391.2 |
| XR_926787.2 | XR_002500332.1 |
| XR_926797.2 | XM_021851335.1 |
| XR_926868.1 | XR_002501001.1 |
| XR_926894.2 | XM_021847935.1 |
| XR_926895.2 | XM_021849373.1 |
| XR_926908.2 | XR_002501960.1 |
| XR_926955.2 | XM_001660047.2 |
| XR_926956.2 | XR_002501317.1 |
| XR_927051.2 | XR_002503168.1 |
| XR_927111.2 | XM_001651276.2 |
| XR_927113.2 | XM_001659158.2 |
| XR_927123.2 | XM_001653861.2 |
| XR_927134.2 | XM_021844808.1 |
| XR_927174.2 | XM_021848204.1 |
| XR_927176.3 | XM_021847257.1 |
| XR_927187.2 | XM_021852838.1 |
| XR_927199.2 | XM_021844629.1 |
| XR_927421.1 | XM_021848498.1 |
| XR_927518.1 | XM_021856872.1 |
| XR_927559.2 | XM_001661968.2 |
| XR_927576.1 | XR_002499832.1 |
| XR_927649.2 | XM_001663324.3 |
| XR_927652.1 | XR_002499977.1 |
| XR_927698.2 | XM_021846361.1 |
| XR_927729.2 | XM_021854604.1 |
| XR_927826.3 | XM_021842685.1 |
| XR_927864.2 | XM_021857028.1 |
| XR_927885.2 | XM_001661855.3 |
| XR_927895.2 | XM_001653515.2 |
| XR_927952.3 | XM_001655804.2 |
| XR_927953.3 | XR_002500019.1 |
| XR_928005.2 | XM_021852612.1 |
| XR_928029.2 | XM_021844207.1 |

| | |
|---|---|
| XR_928033.1 | XR_002499394.1 |
| XR_928047.2 | XM_021857400.1 |
| XR_928059.2 | XM_021841458.1 |
| XR_928066.1 | XM_021842687.1 |
| XR_928171.2 | XM_001649862.2 |
| XR_928242.2 | XM_021838753.1 |
| XR_928260.1 | XM_021843514.1 |
| XR_928309.1 | XM_021849178.1 |
| XR_928312.3 | XR_002503104.1 |
| XR_928340.2 | XR_002501594.1 |
| XR_928399.2 | XR_002502702.1 |
| XR_928407.3 | XM_021853153.1 |
| XR_928448.3 | XM_001655922.2 |
| XR_928452.3 | XM_001655381.2 |
| XR_928461.2 | XM_001661509.2 |
| XR_928491.2 | XM_021853010.1 |
| XR_928650.2 | XM_001648333.2 |
| XR_928665.3 | XM_001656591.2 |
| XR_928727.1 | XM_001649855.2 |
| XR_928855.3 | XM_011495222.2 |
| XR_928863.2 | XM_001657117.2 |
| XR_928868.2 | XM_001649044.2 |
| XR_928896.2 | XM_021854492.1 |
| XR_928899.2 | XM_021843686.1 |
| XR_928910.2 | XM_021839646.1 |
| XR_928913.2 | XM_021845640.1 |
| XR_928934.2 | XM_021849179.1 |
| XR_928959.1 | XM_021852255.1 |
| XR_928962.2 | XM_001656888.2 |
| XR_928965.2 | XR_002501761.1 |
| XR_928967.1 | XM_021852686.1 |
| XR_928972.3 | XM_001657191.2 |
| XR_928993.2 | XR_002502624.1 |
| XR_929013.2 | XM_021850637.1 |
| XR_929027.2 | XM_021857213.1 |
| XR_929040.2 | XM_001654336.2 |
| XR_929113.2 | XM_021846123.1 |
| XR_929119.2 | XR_002501891.1 |
| XR_929159.2 | XR_002502776.1 |
| XR_929407.2 | XM_001661708.2 |
| XR_929462.2 | XM_021852841.1 |
| XR_929464.2 | XR_002502168.1 |
| XR_929529.2 | XM_021852170.1 |
| XR_929580.3 | XR_002502936.1 |
| XR_929605.1 | XR_002498614.1 |
| XR_929627.2 | XM_021841375.1 |
| XR_929731.3 | XR_002501144.1 |
| XR_929748.2 | XM_001648676.3 |
| XR_929767.1 | XM_001655083.2 |
| XR_929939.1 | XM_021844960.1 |

| | |
|---|---|
| XR_929945.1 | XM_001652114.2 |
| XR_929976.2 | XM_021843513.1 |
| XR_929983.2 | XR_002502354.1 |
| XR_929985.2 | XR_002500905.1 |
| XR_930039.2 | XM_001662959.2 |
| XR_930050.2 | XR_002502234.1 |
| XR_930068.2 | XM_021853155.1 |
| XR_930102.2 | XM_021845630.1 |
| XR_930133.2 | XR_002499724.1 |
| XR_930187.2 | XR_002502766.1 |
| XR_930195.1 | XM_021855161.1 |
| XR_930304.2 | XM_001661927.2 |
| XR_930345.1 | XM_001655664.2 |
| XR_930391.2 | XM_001655753.2 |
| XR_930393.2 | XM_021837437.1 |
| XR_930428.2 | XR_002499388.1 |
| XR_930469.2 | XR_002499733.1 |
| XR_930472.1 | XR_002501583.1 |
| XR_930475.2 | XM_021851719.1 |
| XR_930476.2 | XM_021856865.1 |
| XR_930488.1 | XM_021837505.1 |
| XR_930538.2 | XM_021857629.1 |
| XR_930542.2 | XM_021852613.1 |
| XR_930560.2 | XR_002499009.1 |
| XR_930564.2 | XM_021857630.1 |
| XR_930583.3 | XR_002500951.1 |
| XR_930643.1 | XM_021854879.1 |
| XR_930729.2 | XM_001657170.2 |
| XR_930755.3 | XR_002501446.1 |
| XR_930765.2 | XR_002500525.1 |
| XR_930828.2 | XM_001662770.3 |
| XR_930832.2 | XM_021857628.1 |
| XR_930903.3 | XR_002502548.1 |
| XR_930914.2 | XM_021851983.1 |
| XR_931009.2 | XM_001649211.2 |
| XR_931022.2 | XM_021845241.1 |
| XR_931118.2 | XM_001658428.3 |
| XR_931165.2 | XR_002501669.1 |
| XR_931178.2 | XM_001656360.2 |
| XR_931214.2 | XM_021845187.1 |
| XR_931241.2 | XR_002498942.1 |
| XR_931262.1 | XR_002500304.1 |
| XR_931351.2 | XR_002500766.1 |
| XR_931422.2 | XR_002501077.1 |
| XR_931448.2 | XM_021842637.1 |
| XR_931522.2 | XR_002503201.1 |
| XR_931541.2 | XM_001651794.2 |
| XR_931557.3 | XM_001654085.2 |
| XR_931568.2 | XM_021840943.1 |
| XR_931573.2 | XR_002498637.1 |

| | |
|---|---|
| XR_931621.2 | XM_001649048.2 |
| XR_931700.1 | XM_021855595.1 |
| XR_931734.2 | XR_002502627.1 |
| XR_931736.2 | XR_002502863.1 |
| XR_931777.1 | XM_021839414.1 |
| XR_931877.2 | XM_021845939.1 |
| XR_931938.2 | XM_021838317.1 |
| XR_931945.2 | XM_001654730.2 |
| XR_931993.2 | XR_002502814.1 |
| XR_932048.2 | XR_002502825.1 |
| XR_932069.3 | XR_002501532.1 |
| XR_932171.2 | XR_002499432.1 |
| XR_932236.2 | XM_001663924.2 |
| XR_932250.2 | XM_001662063.2 |
| XR_932255.3 | XM_011495298.2 |
| XR_932399.2 | XR_002499008.1 |
| XR_932404.3 | XR_002502346.1 |
| XR_932409.2 | XM_001659710.2 |
| XR_932424.2 | XM_021838776.1 |
| XR_932478.1 | XM_021852227.1 |
| XR_932493.2 | XR_002500858.1 |
| XR_932518.2 | XM_021839952.1 |
| XR_932542.2 | XR_002503057.1 |
| XR_932590.1 | XR_002500333.1 |
| XR_932615.3 | XM_021852172.1 |
| XR_932644.2 | XM_021853850.1 |
| XR_932669.2 | XR_002500207.1 |
| XR_932838.3 | XR_002501993.1 |
| XR_932928.3 | XR_002502204.1 |
| XR_933085.2 | XR_002499110.1 |
| XR_933149.3 | XR_002500004.1 |
| XR_933341.3 | XM_001651106.2 |
| XR_933352.2 | XM_001657076.2 |
| XR_933433.3 | XM_001650350.2 |
| XR_933519.2 | XM_001651505.2 |
| XR_933596.2 | XR_002499967.1 |
| XR_933619.3 | XR_002503150.1 |
| XR_933623.2 | XR_002503083.1 |
| XR_933711.2 | XM_001656141.2 |
| XR_933729.3 | XM_001649043.2 |
| XR_933754.2 | XR_002500260.1 |
| XR_933786.2 | XM_021853868.1 |
| XR_934174.2 | XR_002500436.1 |
| XR_934178.2 | XM_001652520.2 |
| XR_934252.2 | XR_002501159.1 |
| XR_934258.2 | XM_021839816.1 |
| XR_934312.3 | XM_001653027.2 |
| XR_934326.2 | XM_021842689.1 |
| XR_934369.1 | XM_001660285.2 |
| XR_934476.3 | XR_002500686.1 |

| | |
|---|---|
| XR_934502.1 | XM_021849451.1 |
| XR_934535.1 | XM_001660437.2 |
| XR_934637.3 | XR_002500623.1 |
| XR_934687.2 | XM_021840900.1 |
| XR_934751.2 | XM_021856689.1 |
| XR_934753.2 | XM_021846306.1 |
| XR_934771.2 | XR_002499749.1 |
| XR_934811.2 | XM_021854556.1 |
| XR_934874.3 | XM_021856228.1 |
| XR_934976.2 | XM_021854838.1 |
| XR_935089.2 | XM_001657773.2 |
| XR_935157.2 | XM_021839551.1 |
| XR_935171.2 | XM_021853851.1 |
| XR_935195.1 | XM_001647696.2 |
| XR_935200.1 | XM_001658854.2 |
| XR_935202.1 | XR_002503250.1 |
| XR_935325.1 | XM_021842320.1 |
| XR_935327.2 | XM_001653431.2 |
| XR_935378.3 | XM_001650951.2 |
| XR_935432.2 | XR_002498876.1 |
| XR_935456.2 | XM_001649950.2 |
| XR_935559.2 | XR_002499670.1 |
| XR_935599.2 | XR_002499634.1 |
| XR_935690.3 | XR_002502201.1 |
| XR_935712.2 | XM_001657878.2 |
| XR_935741.2 | XM_021842956.1 |
| XR_935812.3 | XR_002502878.1 |
| XR_935823.1 | XM_021857241.1 |
| XR_935866.2 | XM_001661702.2 |
| XR_935896.2 | XR_002499573.1 |
| XR_935942.2 | XR_002500743.1 |
| XR_935966.2 | XM_021857131.1 |
| XR_936006.2 | XM_021839658.1 |
| XR_936015.2 | XM_001662431.2 |
| XR_936052.2 | XR_002499387.1 |
| XR_936074.2 | XR_002500483.1 |
| XR_936091.2 | XR_002503213.1 |
| XR_936111.2 | XM_021857127.1 |
| XR_936137.3 | XR_002501873.1 |
| XR_936169.2 | XM_001649678.2 |
| XR_936174.1 | XR_002500368.1 |
| XR_936272.2 | XR_002498705.1 |
| XR_936283.2 | XR_002499964.1 |
| XR_936472.1 | XM_021852365.1 |
| XR_936554.2 | XM_001649308.2 |
| XR_936626.2 | XM_021846055.1 |
| XR_936728.2 | XM_001655942.2 |
| XR_936739.3 | XM_001657238.3 |
| XR_936750.3 | XR_002501658.1 |
| XR_936764.2 | XR_002502625.1 |

| | |
|---|---|
| XR_936847.2 | XR_002503152.1 |
| XR_936848.2 | XR_002498948.1 |
| XR_936896.2 | XR_002499884.1 |
| XR_936898.2 | XM_021838437.1 |
| XR_936911.2 | XR_002499711.1 |
| XR_936916.2 | XM_021844413.1 |
| XR_936962.1 | XR_002502385.1 |
| XR_936963.1 | XR_002501391.1 |
| XR_936968.2 | XM_021837443.1 |
| XR_936984.2 | XR_002503069.1 |
| XR_937057.1 | XR_002500908.1 |
| XR_937251.2 | XM_001660286.2 |
| XR_937263.2 | XM_001661470.2 |
| XR_937267.2 | XR_002499798.1 |
| XR_937268.2 | XR_002501435.1 |
| XR_937394.2 | XR_002502688.1 |
| XR_937396.2 | XR_002501566.1 |
| XR_937438.2 | XR_002503075.1 |
| XR_937606.3 | XM_001660459.2 |
| XR_937637.2 | XR_002502947.1 |
| XR_937688.3 | XM_001654154.2 |
| XR_937739.1 | XR_002502467.1 |
| XR_937748.3 | XR_002502295.1 |
| XR_937786.3 | XR_002501456.1 |
| XR_937811.2 | XR_002501377.1 |
| XR_937819.1 | XM_001650457.2 |
| XR_937827.3 | XM_001659009.2 |
| XR_937856.1 | XR_002500907.1 |
| XR_937863.2 | XR_002502879.1 |
| XR_937952.2 | XR_002502472.1 |
| XR_938055.2 | XR_002501734.1 |
| XR_938058.2 | XR_002502572.1 |
| XR_938072.2 | XR_002502427.1 |
| XR_938076.1 | XR_002498621.1 |
| XR_938086.2 | XM_011494699.2 |
| XR_938102.2 | XR_002500352.1 |
| XR_938131.2 | XR_002500054.1 |
| XR_938139.2 | XR_002500703.1 |
| XR_938171.2 | XM_001655456.2 |
| XR_938212.2 | XM_001652391.2 |
| XR_938226.2 | XM_021845976.1 |
| XR_938283.2 | XR_002498721.1 |
| XR_938289.2 | XM_021855248.1 |
| XR_938433.1 | XR_002502773.1 |
| XR_938507.2 | XR_002499490.1 |
| XR_938516.1 | XR_002501119.1 |
| XR_938534.1 | XR_002498795.1 |
| XR_938603.3 | XR_002501178.1 |
| XR_938641.2 | XM_021838628.1 |
| XR_938677.3 | |

XR_938679.1
XR_938701.3
XR_938753.2
XR_938771.1
XR_938780.3
XR_938796.3
XR_938854.1
XR_938919.2
XR_938920.2
XR_938931.2
XR_938935.2
XR_938957.3
XR_938977.2
XR_939091.2
XR_939185.2
XR_939242.2
XR_939498.2
XR_939522.2
XR_939620.2
XR_939722.2
XR_939785.2
XR_939834.1
XR_939844.2
XR_939854.2
XR_939873.2
XR_939886.2
XR_939974.2
XR_940002.3
XR_940019.2
XR_940067.2
XR_940083.1
XR_940093.1
XR_940111.2
XR_940183.3
XR_940187.3
XR_940205.2
XR_940228.2
XR_940235.2
XR_940260.1
XR_940281.2
XR_940317.2
XR_940392.3
XR_940455.2
XR_940541.3
XR_940586.1
XR_940609.2
XR_940700.3
XR_940733.1
XR_940782.3
XR_940846.2

XR_940857.2
XR_940859.1
XR_940897.3
XR_940970.2
XR_940974.2
XR_940990.2
XR_940997.2
XR_941024.2
XR_941077.1
XR_941191.2
XR_941198.2
XR_941226.2
XR_941265.2
XR_941266.2
XR_941302.2
XR_941314.2
XR_941352.2
XR_941358.2
XR_941368.2
XR_941401.2
XR_941422.3
XR_941546.1
XR_941752.1
XR_941758.1
XR_941779.3
XR_941849.2
XR_941947.1
XR_941954.2
XR_941972.2
XR_942014.1
XR_942092.2
XR_942116.2
XR_942124.2
XR_942150.1
XR_942152.1
XR_942158.3
XR_942305.3
XR_942460.2
XR_942666.1
XR_942676.2
XR_942693.3
XR_942699.3
XR_942738.3
XR_942765.3
XR_942866.2
XR_942881.2
XR_942901.2
XR_942987.2
XR_943084.3
XR_943085.3

XR_943136.2
XR_943176.1
XR_943270.2
XR_943413.3
XR_943577.1
XR_943586.2
XR_943610.2
XR_943636.2
XR_943640.2
XR_943701.2
XR_943705.2
XR_943724.2
XR_943736.2
XR_943771.2
XR_943864.3
XR_943908.2
XR_943932.2
XR_943953.2
XR_943955.3
XR_943977.2
XR_944008.2
XR_944013.3
XR_944089.2
XR_944155.2
XR_944157.1
XR_944172.1
XR_944174.2
XR_944187.3
XR_944190.2
XR_944191.2
XR_944321.3
XR_944329.3
XR_944386.2
XR_944428.3
XR_944432.2
XR_944515.2
XR_944537.2
XR_944581.2
XR_944864.2
XR_944868.2
XR_944901.2
XR_944904.2
XR_944974.2
XR_945015.2
XR_945054.2
XR_945057.2
XR_945081.2
XR_945188.1
XR_945238.2
XR_945280.2

XR_945284.3
XR_945329.2
XR_945356.1
XR_945365.3
XR_945389.2
XR_945416.1
XR_945429.2
XR_945449.2
XR_945453.2
XR_945520.1
XR_945568.2
XR_945949.2
XR_945957.2
XR_945991.2
XR_946003.2
XR_946152.2
XR_946198.2
XR_946246.2
XR_946309.2
XR_946513.3
XR_946521.3
XR_946523.1
XR_946576.2
XR_946636.2
XR_946642.2
XR_946665.1
XR_946674.2
XR_946682.2
XR_946721.2
XR_946764.1
XR_946802.3
XR_946869.2
XR_946892.2
XR_946952.2
XR_947005.2
XR_947088.2
XR_947141.2
XR_947147.2
XR_947174.2
XR_947255.2
XR_947273.2
XR_947278.2
XR_947280.2
XR_947289.1
XR_947488.2
XR_947518.2
XR_947529.2
XR_947603.2
XR_947608.2
XR_947636.1

XR_947682.2
XR_947727.1
XR_947807.1
XR_947834.2
XR_947851.2
XR_947940.2
XR_948008.2
XR_948039.1
XR_948057.2
XR_948092.3
XR_948352.2
XR_948410.2
XR_948435.2
XR_948528.2
XR_948533.2
XR_948540.2
XR_948581.2
XR_948623.2
XR_948664.2
XR_948692.2
XR_948899.3
XR_948944.3
XR_948956.2
XR_948958.3
XR_948965.1
XR_949045.1
XR_949163.3
XR_949201.3
XR_949269.2
XR_949286.2
XR_949402.3
XR_949563.2
XR_949584.2
XR_949692.2
XR_949693.2
XR_949694.3
XR_949705.1
XR_949706.2
XR_949933.3
XR_949950.2
XR_950008.3
XR_950143.2
XR_950215.2
XR_950235.2
XR_950265.2
XR_950334.1
XR_950580.1
XR_950632.1
XR_950638.2
XR_950767.3

XR_950818.3
XR_950871.2
XR_950891.2
XR_950945.3
XR_951108.2
XR_951291.2
XR_951312.2
XR_951398.2
XR_951426.3
XR_951438.2
XR_951439.2
XR_951443.2
XR_951449.2
XR_951681.2
XR_951862.2
XR_951872.2
XR_951914.2
XR_951920.2
XR_951932.3
XR_951996.2
XR_952154.1
XR_952182.3
XR_952219.2
XR_952247.2
XR_952538.2
XR_952545.2
XR_952686.2
XR_952697.2
XR_952711.2
XR_952881.3
XR_952979.1
XR_953178.3
XR_953253.2
