## Supplement file 2 for "An adaptive compromise - Conflicting evolutionary pressures on arthropod-borne Zika virus dinucleotide composition in mammalian hosts and mosquito vectors"

>Wildtype
ACCGCCTGGGGGTGGGGGGAGGCTGGGGCCCTGATCACAGCCGCAACTTCCACTTTGTGGGAAGGCTCTCCGA
ACAAGTACTGGAACTCCTCTACAGCCACTTCACTGTGTAACATTTTTAGGGGAAGTTACTTGGCTGGAGCTTC
TCTAATCTACACAGTAACAAGAAACGCTGGCTTGGTCAAGAGACGTGGGGGTGGAACAGGAGAGACCCTGGGA
GAGAAATGGAAGGCCCGCTTGAACCAGATGTCGGCCCTGGAGTTCTACTCCTACAAAAAGTCAGGCATCACCG
AGGTGTGCAGAGAAGAGGCCCGCCGCGCCCTCAAGGACGGTGTGGCAACGGGAGGCCATGCTGTGTCCCGAGG
AAGTGCAAAGCTGAGATGGTTGGTGGAGCGGGGATACCTGCAGCCCTATGGAAAGGTCATTGATCTTGGATGT
GGCAGAGGGGGCTGGAGTTACTACGCCGCCACCATCCGCAAAGTTCAAGAAGTGAAAGGATACACAAAAGGAG
GCCCTGGTCATGAAGAACCCGTGTTGGTGCAAAGCTATGGGTGGAACATAGTCCGTCTTAAGAGTGGGGTGGA
CGTCTTTCATATGGCGGCTGAGCCGTGTGACACGCTGCTGTGTGACATAGGTGAGTCATCATCTAGTCCTGAA
GTGGAAGAAGCACGGACGCTCAGAGTCCTCTCCATGGTGGGGGATTGGCTTGAAAAAAGACCAGGAGCCTTTT
GTATAAAAGTGTTGTGCCCATACACCAGCACTATGATGGAAACCCTGGAGCGACTGCAGCGTAGGTATGGGGG
AGGACTGGTCAGAGTGCCACTCTCCCGCAACTCTACACATGAGATGTACTGGGTCTCTGGAGCGAAAAGCAAC
ACCATAAAAAGTGTGTCCACCACGAGCCAGCTCCTCTTGGGGCGCATGGACGGGCCTAGAAGGCCAGTGAAAT
ATGAGGAGGATGTGAATCTCGGCTCTGGCACGCGGGCTGTGGTAAGCTGCGCTGAAGCTCCCAACATGAAGAT
CATTGGTAACCGCATTGAAAGGATCCGCAGTGAGCACGCGGAAACGTGGTTCTTTGACGAGAACCACCCATAT
AGGACATGGGCTTACCATGGAAGCTATGAGGCCCCCACACAAGGGTCAGCGTCCTCTCTAATAAACGGGGTTG
TCAGGCTCCTGTCAAAACCCTGGGATGTGGTGACTGGAGTCACAGGAATAGCCATGACCGACACCACACCGTA
TGGTCAGCAAAGAGTTTTCAAGGAAAAAGTGGACACTAGGGTGCCAGACCCCCAAGAAGGCACTCGTCAGGTT
ATGAGCATGGTCTCTTCCTGGTTGTGGAAAGAGCTAGGCAAACACAAACGGCCACGAGTCTGTACCAAAGAAG
AGTTCATCAACAAGGTTCGTAGCAATGCAGCATTAGGGGCAATATTTGAAGAGGAAAAAGAGTGGAAGACTGC
AGTGGAAGCTGTGAACGATCCAAGGTTCTGGGCTCTAGTGGACAAGGAAAGAGAGCACCACCTGAGAGGAGAG
TGCCAGAGTTGTGTGTATAACATGATGGGAAAAAGAGAAAAGAAACAAGGGGAATTTGGAAAGGCCAAGGGCA
GCCGCGCCATCTGGTATATGTGGCTAGGGGCTAGATTTCTAGAGTTCG

>SCR
ACCGCATGGGGATGGGGAGAGGCAGGGGCACTGATCACCGCCGCAACATCGACATTGTGGGAAGGCTCCCCAA
ACAAGTACTGGAACTCCTCCACCGCCACCTCCCTGTGTAACATCTTCAGAGGCAGCTACTTAGCTGGCGCATC
TCTGATCTACACAGTGACCAGGAACGCCGGTTTGGTTAAGAGGCGAGGGGGTGGAACTGGTGAGACTCTAGGA
GAGAAGTGGAAGGCTCGATTGAACCAGATGTCGGCCCTGGAGTTTTACTCGTACAAAAAGTCAGGTATCACAG
AGGTTTGCAGAGAAGAGGCGCGTCGCGCCCTCAAGGACGGAGTAGCAACGGGTGGCCATGCAGTCTCGCGAGG
TAGTGCAAAACTGAGATGGTTGGTGGAGCGGGGATATCTGCAGCCGTATGGCAAGGTAATTGATCTTGGGTGC
GGCAGAGGGGGTTGGAGCTATTACGCTGCCACCATCCGCAAGGTGCAGGAAGTGAAGGGTTACACGAAAGGGG
GCCCGGGGCATGAAGAACCTGTCTTAGTGCAAAGTTATGGATGGAACATAGTCCGTCTGAAGAGTGGGGTTGA
TGTGTTTCATATGGCTGCTGAACCCTGTGACACACTGCTCTGTGACATTGGAGAGTCGTCCTCCAGCCCAGAA
GTAGAGGAAGCCCGCACCCTCAGAGTGCTTTCCATGGTGGGGGACTGGCTAGAGAAAAGACCAGGAGCATTCT
GTATCAAAGTGTTGTGCCCTTACACCAGCACTATGATGGAGACACTGGAGCGGCTTCAGCGGAGATATGGAGG
GGGTCTGGTCAGAGTGCCTCTCTCCCGGAACTCAACTCATGAGATGTACTGGGTGTCAGGTGCCAAGAGCAAC
ACCATAAAGAGTGTCTCAACGACCAGTCAGCTCCTCTTGGGACGAATGGATGGACCAAGGAGGCCAGTGAAGT
ACGAGGAAGATGTAAACCTGGGCTCAGGCACCCGTGCTGTTGTAAGCTGCGCTGAAGCCCCCAACATGAAGAT
CATTGGCAACCGCATTGAAAGGATTCGCAGTGAGCACGCAGAAACTTGGTTCTTCGACGAGAATCACCCCTAT
AGAACATGGGCTTACCATGGAAGCTATGAGGCCCCCACACAAGGCTCAGCCTCATCTCTGATCAACGGAGTTG
TCAGACTCCTGTCAAAGCCTTGGGATGTGGTGACGGGAGTCACAGGAATTGCCATGACTGACACCACCCCCTA
TGGACAGCAGAGGGTGTTCAAGGAAAAAGTGGACACAAGGGTCCCGGACCCCCAAGAGGGAACACGCCAAGTC
ATGAGTATGGTGTCTTCTTGGTTGTGGAAAGAGCTTGGCAAGCACAAACGGCCTCGAGTTTGTACCAAAGAAG
AGTTCATTAACAAGGTGCGAAGCAATGCAGCATTAGGGGCAATTTTTGAAGAGGAAAAAGAGTGGAAAACGGC
AGTGGAAGCTGTAAACGATCCAAGATTCTGGGCGCTGGTGGACAAGGAAAGAGAGCATCATCTGAGAGGAGAG
TGCCAGAGTTGTGTTTATAACATGATGGGCAAAAGGGAGAAAAAGCAAGGAGAATTTGGAAAGGCCAAGGGCA
GCCGAGCCATCTGGTATATGTGGCTAGGGGCAAGATTCCTAGAGTTCG

>UpA_max TpA^Target_ratio=2.500^
ACCGCCTGGGGGTGGGGGGAGGCTGGGGCCCTAATAACAGCCGCTACTTCTACTTTATGGGAAGGCTCTCCGA
ATAAGTACTGGAACTCCTCTACAGCTACTTCACTATGTAATATATTTAGGGGTAGTTACTTAGCTGGAGCTTC
TCTAATATATACAGTAACTAGAAACGCTGGCTTAGTTAAGAGACGTGGGGGTGGTACAGGAGAGACCCTAGGA
GAGAAATGGAAGGCCCGCTTAAACCAGATGTCGGCCCTAGAGTTCTACTCCTATAAAAAGTCAGGTATTACCG
AGGTATGTAGAGAAGAGGCCCGCCGCGCCCTTAAGGACGGTGTAGCTACGGGAGGCCATGCTGTATCCCGAGG
TAGTGCTAAGCTTAGATGGTTAGTAGAGCGGGGATACCTACAGCCCTATGGTAAGGTTATAGATCTAGGATGT
GGTAGAGGGGGCTGGAGTTACTACGCCGCTACTATACGTAAAGTACAAGAAGTTAAAGGATATACTAAAGGAG

GCCCTGGTCATGAAGAACCCGTATTAGTACAAAGCTATGGGTGGAATATAGTACGTCTTAAGAGTGGGGTAGA
CGTATTTCATATGGCGGCTGAGCCGTGTGATACGCTACTATGTGATATAGGTGAGTCATCATCTAGTCCTGAA
GTAGAAGAAGCACGTACGCTTAGAGTACTATCTATGGTAGGGGATTGGCTAGAAAAAAGACCAGGAGCCTTTT
GTATAAAAGTATTATGCCCATATACTAGTACTATGATGGAAACCCTAGAGCGACTACAGCGTAGGTATGGGGG
AGGACTAGTAAGAGTACCACTATCCCGTAACTCTACACATGAGATGTACTGGGTATCTGGAGCGAAAAGTAAT
ACTATAAAAAGTGTATCTACTACGAGCCAGCTACTATTAGGGCGTATGGACGGGCCTAGAAGGCCAGTTAAAT
ATGAGGAGGATGTAAATCTCGGCTCTGGTACGCGGGCTGTAGTAAGCTGCGCTGAAGCTCCTAATATGAAGAT
AATAGGTAACCGTATAGAAAGGATACGTAGTGAGCACGCGGAAACGTGGTTCTTTGACGAGAACCACCCATAT
AGGACATGGGCTTACCATGGTAGCTATGAGGCCCCTACACAAGGGTCAGCGTCCTCTCTAATAAACGGGGTAG
TAAGGCTACTATCTAAACCCTGGGATGTAGTTACTGGAGTAACAGGTATAGCTATGACCGATACTACACCGTA
TGGTCAGCAAAGAGTATTTAAGGAAAAAGTAGATACTAGGGTACCAGACCCCCAAGAAGGTACTCGTCAGGTT
ATGAGTATGGTATCTTCCTGGTTATGGAAAGAGCTAGGTAAACATAAACGGCCACGAGTATGTACTAAAGAAG
AGTTTATTAATAAGGTACGTAGTAATGCAGCATTAGGGGCTATATTTGAAGAGGAAAAAGAGTGGAAGACTGC
AGTAGAAGCTGTAAACGATCCTAGGTTCTGGGCTCTAGTAGATAAGGAAAGAGAGCACCACCTTAGAGGAGAG
TGCCAGAGTTGTGTATATAATATGATGGGTAAAAGAGAAAAGAAACAAGGGGAATTTGGTAAGGCTAAGGGTA
GCCGCGCTATATGGTATATGTGGCTAGGGGCTAGATTTCTAGAGTTCG

>CpG_max hi CpG^Target_ratio=2.500^
ACCGCCTGGGGGTGGGGCGAGGCCGGCGCGCTGATCACCGCCGCGACGTCGACGTTGTGGGAAGGCTCGCCGA
ACAAGTACTGGAACTCGTCTACCGCGACGTCGCTGTGTAACATTTTTAGGGGAAGTTACTTGGCCGGCGCGTC
GCTAATCTACACGGTAACGCGAAACGCCGGCTTGGTCAAGCGACGCGGCGGCGGAACCGGCGAGACGCTCGGC
GAGAAATGGAAGGCGCGCTTGAACCAGATGTCGGCGCTCGAGTTCTACTCGTACAAAAAGTCGGGCATCACCG
AGGTGTGCCGCGAAGAGGCGCGCCGCGCGCTCAAGGACGGCGTCGCGACGGGCGGCCACGCGGTGTCGCGCGG
AAGCGCGAAGCTGCGATGGTTGGTCGAGCGCGGATACCTGCAGCCGTACGGAAAGGTCATCGATCTCGGATGC
GGCCGCGGCGGCTGGAGTTACTACGCCGCGACGATCCGCAAAGTTCAAGAAGTGAAAGGATACACGAAAGGCG
GCCCCGGTCACGAAGAACCCGTGTTGGTGCAAAGCTACGGGTGGAACATAGTCCGTCTTAAGAGCGGCGTCGA
CGTCTTTCATATGGCGGCGGAGCCGTGCGACACGCTGCTGTGCGACATAGGCGAGTCGTCGTCTAGTCCGGAA
GTCGAAGAAGCGCGGACGCTCCGCGTCCTCTCGATGGTCGGCGATTGGCTCGAAAAACGACCGGGCGCGTTTT
GTATAAAAGTGTTGTGCCCGTACACGAGCACTATGATGGAAACGCTCGAGCGCCTGCAGCGTAGGTACGGCGG
CGGACTCGTCCGCGTGCCGCTCTCGCGCAACTCTACGCACGAGATGTACTGGGTCTCCGGCGCGAAAAGCAAC
ACGATAAAAAGCGTGTCGACGACGAGCCAGCTCCTCTTGGGGCGCATGGACGGGCCTAGACGGCCGGTGAAAT
ACGAGGAGGACGTGAATCTCGGCTCCGGCACGCGCGCCGTCGTAAGCTGCGCCGAAGCGCCGAACATGAAGAT
CATCGGTAACCGCATCGAACGAATCCGCAGCGAGCACGCGGAAACGTGGTTCTTCGACGAGAACCACCCGTAT
AGGACGTGGGCGTACCACGGAAGCTACGAGGCGCCGACGCAAGGGTCCGCGTCGTCGCTAATAAACGGCGTCG
TCCGGCTCCTGTCGAAACCGTGGGACGTCGTGACCGGCGTCACCGGAATAGCGATGACCGACACGACGCCGTA
CGGTCAGCAACGCGTTTTCAAGGAAAAAGTCGACACTAGGGTGCCGGACCCGCAAGAAGGCACGCGTCAGGTT
ATGAGCATGGTCTCGTCGTGGTTGTGGAAAGAGCTAGGCAAACACAAACGGCCGCGCGTCTGTACGAAAGAAG
AGTTCATCAACAAGGTTCGTAGCAACGCGGCGTTAGGCGCGATATTCGAGGAGGAAAAAGAGTGGAAGACCGC
CGTCGAAGCCGTGAACGATCCGCGGTTCTGGGCGCTAGTCGACAAGGAACGCGAGCACCACCTGCGCGGCGAG
TGCCAGAGTTGCGTGTATAACATGATGGGAAAACGCGAAAAGAAACAAGGCGAATTCGGAAAGGCGAAGGGCA
GCCGCGCGATCTGGTATATGTGGCTAGGCGCTAGATTTCTAGAGTTCG

>CpG_1.0 CpG^Target_ratio=1.000^
ACCGCCTGGGGCTGGGGTGAGGCTGGAGCACTCATCACCGCCGCAACGTCGACGTTGTGGGAAGGCTCGCCGA
ACAAGTACTGGAACTCGTCTACGGCCACGTCACTGTGTAACATTTTTAGGGGAAGTTACTTGGCGGGAGCTTC
GCTAATCTACACGGTAACACGAAACGCGGGCTTGGTCAAGAGAAGGGGCGGTGGAACAGGAGAGACTCTGGGT
GAGAAATGGAAGGCGAGATTGAACCAGATGTCCGCGCTCGAGTTCTACTCCTACAAAAAGTCCGGCATCACGG
AGGTGTGCAGAGAAGAGGCGCGTCGCGCTCTAAAGGACGGTGTCGCAACTGGAGGCCATGCTGTGTCGCGCGG
AAGTGCAAAGCTGAGATGGTTGGTTGAGCGAGGATACCTGCAGCCGTATGGAAAGGTCATTGATCTTGGATGT
GGCAGAGGTGGCTGGAGTTATTATGCCGCAACGATCCGCAAAGTTCAAGAAGTGAAAGGATACACAAAAGGAG
GACCGGGTCATGAAGAACCAGTTTTGGTGCAAAGCTACGGGTGGAACATAGTACGTCTTAAGAGTGGGGTAGA
CGTCTTTCATATGGCTGCCGAGCCGTGTGACACCCTCCTTTGCGACATCGGTGAGTCGTCATCTAGTCCTGAA
GTCGAAGAAGCACGGACTCTCCGCGTCCTCTCGATGGTGGGGGATTGGCTTGAAAAAAGACCGGGCGCCTTCT
GTATAAAAGTGTTGTGCCCATACACCAGCACTATGATGGAAACTCTCGAGAGACTGCAGCGTAGGTATGGCGG
CGGACTCGTCAGAGTGCCACTCTCTCGAAACTCTACACATGAGATGTACTGGGTGTCTGGAGCGAAAAGCAAC
ACGATAAAAAGTGTTTCCACGACAAGCCAGCTCCTATTGGGGAGGATGGATGGTCCTAGAAGGCCAGTGAAAT
ATGAGGAGGATGTAAATCTCGGCTCTGGGACTCGAGCCGTGGTAAGCTGCGCTGAAGCGCCGAACATGAAGAT
CATTGGTAACCGCATTGAACGAATCCGCAGCGAGCACGCTGAAACATGGTTCTTTGACGAGAACCACCCCTAT

AGGACATGGGCGTACCATGGAAGCTATGAGGCGCCGACACAAGGTTCTGCGTCCTCTCTAATAAACGGGGTCG
TCCGACTCCTGTCAAAACCCTGGGATGTGGTGACGGGGGTCACGGGAATAGCGATGACGGACACCACACCATA
CGGTCAGCAACGAGTTTTCAAGGAAAAAGTCGACACTCGCGTGCCAGACCCGCAAGAAGGCACTCGTCAGGTT
ATGAGCATGGTCTCGTCGTGGTTGTGGAAAGAACTAGGCAAACACAAAAGACCACGAGTCTGTACGAAAGAAG
AGTTCATCAACAAGGTTAGAAGCAACGCCGCATTAGGCGCAATATTTGAAGAGGAAAAAGAGTGGAAGACTGC
CGTGGAAGCTGTGAACGATCCACGGTTCTGGGCTCTCGTCGACAAGGAAAGAGAGCATCACCTCAGAGGAGAG
TGCCAGAGTTGTGTGTATAACATGATGGGAAAAAGAGAAAAGAAACAAGGCGAATTTGGAAAGGCCAAGGGCA
GCCGCGCGATCTGGTATATGTGGCTCGGCGCGCGATTTCTAGAGTTCG
